# Structural and mechanistic views of enzymatic, heme-dependent nitrogen-nitrogen bond formation

**DOI:** 10.1101/2023.12.15.571702

**Authors:** Melanie A. Higgins, Xinjie Shi, Jordi Soler, Jill B. Harland, Taylor Parkkila, Nicolai Lehnert, Marc Garcia-Borràs, Yi-Ling Du, Katherine S. Ryan

## Abstract

Molecules with nitrogen-nitrogen (N-N) bonds constitute a large group of clinically important drugs, and various synthetic approaches have been developed to construct functional groups like hydrazines, diazos, pyrazoles, and N-nitrosos. While hundreds of N-N-containing specialized natural metabolites have also been discovered, little is known about the underlying enzymatic mechanisms that have evolved for N-N bond formation. In order to directly form a single N(sp^3^)-N(sp^3^) bond, enzymes must reverse the typical nucleophilicity of one nitrogen. Here, we report structural and mechanistic interrogations of the piperazate synthase PipS, a heme-dependent enzyme that catalyzes an N-N bond forming cyclization of *N*^5^-OH-L-ornithine to give the non-proteinogenic amino acid L-piperazic acid. We show that PipS can process a variety of *N*-substituted hydroxylamines, to give either an imine or an N-N bond, in a substrate-specific manner. Using a combination of structural and biochemical experiments, computational studies, and spectroscopic characterization, we propose that heme-dependent dehydration and N-N bond formation in PipS proceed through divergent pathways, which may stem from a shared nitrenoid intermediate that effectively reverses the nucleophilicity of the hydroxylamine nitrogen. Our work expands the current knowledge of enzymatic N-N bond formation, and delineates the catalytic versatility of a heme cofactor, paving the way for future development of genetically encoded biocatalysts for N-N bond formation.

## Main Text

Natural products with a nitrogen-nitrogen (N-N) bond are a group of rare and often bioactive natural products (**Fig. 1a**), and molecules with an N-N bond include critical intermediates in the nitrogen cycle and important synthetic drugs like Remdesivir and Valsartan^1–5^. However, direct formation of an N(sp^3^)-N(sp^3^) bond presents a conundrum: two typically nucleophilic atoms must unite^6^. To overcome this fundamental barrier to bond formation, diverse biosynthetic strategies have emerged toward N-N bond formation (**Supplementary Fig. 1**)^7–14^. In the biosynthetic pathway to the non-proteinogenic amino acid piperazic acid, the heme-dependent piperazate synthase KtzT reacts with hydroxylamine-containing *N*^5^-OH-L-ornithine (L-**1**) to give the cyclic hydrazine L-piperazic acid (L-**2**) (**Fig. 1b**)^15^. The gene *ktzT* and its homologs are widely distributed in bacterial genomes, often adjacent to non-ribosomal peptide synthetase and polyketide synthase genes^15–18^.

**Fig. 1.**
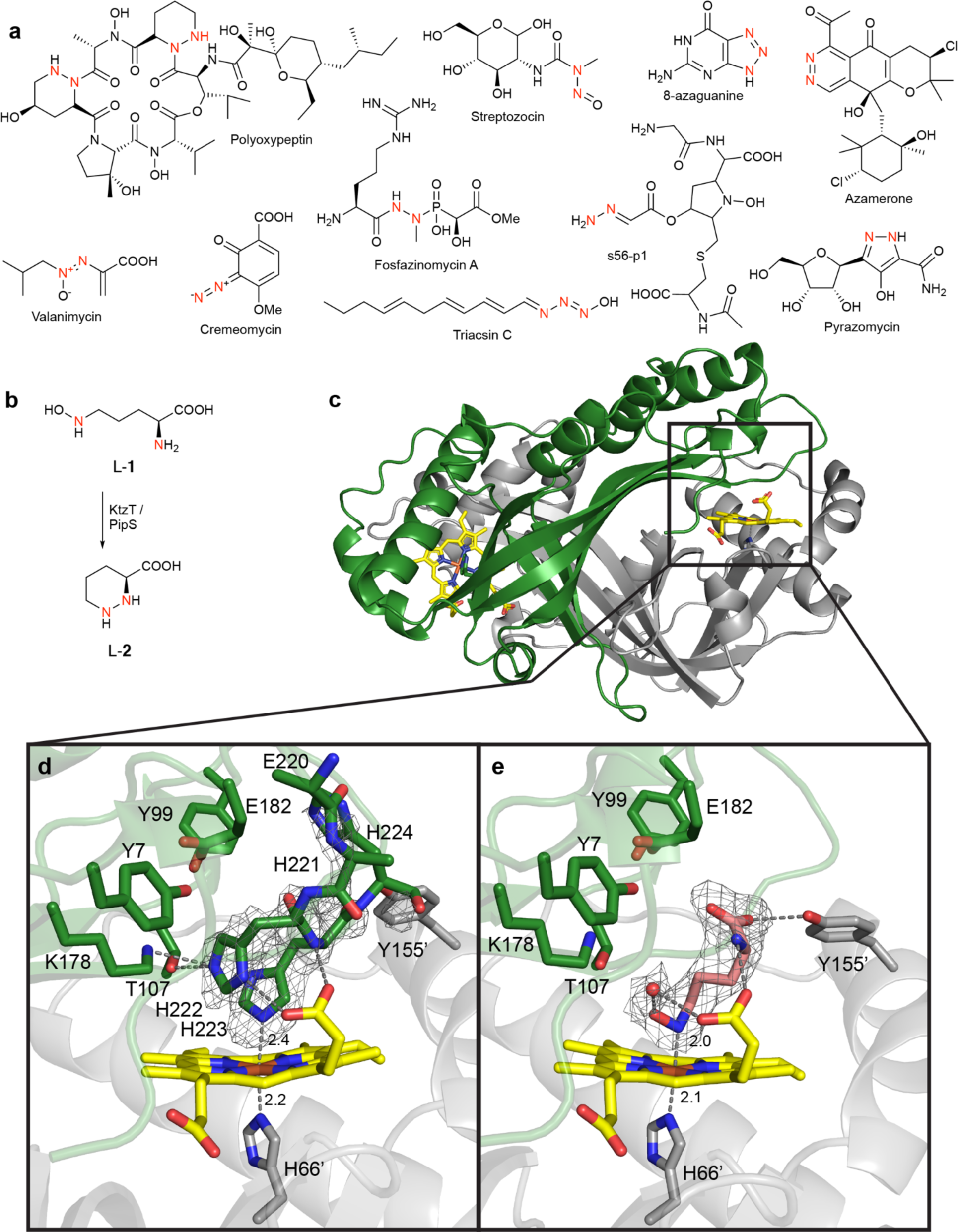
Structure of PipS. **a**, Select natural products with N-N bonds. **b**, Biosynthetic transformation of L-**1** to L-**2**. **c**, Dimeric structure of holo-PipS, where monomer A is green, monomer B is grey, and heme is yellow. **d**, Proximal and distal sites for the heme iron for holo-PipS. The grey mesh represents the F_obs_ – F_calc_ simulated annealing omit map for the partial His_6_-tag contoured to 3 σ. **e**, PipS-L-**1**-complex. The grey mesh represents the F_obs_ – F_calc_ simulated annealing omit map for L-**1** (in pink) and an ordered water, contoured to 3 σ. Nitrogen is blue, oxygen is red, iron is orange, and the water molecule is a red sphere. Key distances are given in angstroms (Å).

To interrogate the mechanism of N-N bond formation by a piperazate synthase, we first set out to determine the binding mode of L-**1** to the enzyme active site, which would provide valuable information regarding the involvement of the heme cofactor and potential catalytic residues. We chose to use X-ray crystallography but found that KtzT was recalcitrant to crystallization. Thus, we purified a new piperazate synthase PipS (Accession number = WP_030722408.1) encoded by a polyoxypeptin-like gene cluster from *Streptomyces griseus* subsp. *griseus* NRRL F-5144 (**Supplementary Table 1**)^19^. PipS has 54 % amino acid identity to KtzT, and as-isolated, C-terminal His_6_-tagged PipS binds heme b with a similar Soret band of 411 nm^15^, and performs the identical reaction to KtzT, converting L-**1** into L-**2** (**Extended Data Fig. 1a-b, Supplementary Fig. 2**). We crystallized holo-PipS and obtained a structure to 2.08 Å resolution (**Supplementary Table 2, Supplementary Fig. 3**). PipS is a dimer with each monomer consisting of a split barrel-like fold flanked by a series of alpha helices (**Fig. 1c**). A single heme molecule is present at an interface of the monomers, with the distal site exposed to monomer A, and a coordinating proximal His66 coming from monomer B. There is also extensive electron density in the distal axial site that we were able to attribute to the C-terminal His_6_-tag by EPR, UV-Vis, and structural analysis (**Fig. 1d, Extended Data Figure 1b-f**).

Next, we set out to obtain a substrate-or product-bound crystal structure. Since the C-terminal His_6_-tag coordinates the heme in our holo-PipS structure, we generated a tag-free version of PipS to ensure we would observe substrate or product binding. EPR studies determined that the tag-free form of the enzyme exhibits an axial HS ferric signal with g-values at g_eff_ = 5.74, 2, indicating that the ferric heme is either five-coordinate or has a water molecule bound (**Extended Data Fig. 1c, Supplementary Fig. 4**).^20–22^ However, we were unable to produce diffraction-quality crystals of PipS without the His_6_-tag, so we aimed to outcompete the His_6_-tag by soaking a holo-PipS crystal with excess L-**1** for 1 hr. The resulting 1.92 Å resolution structure has the same overall fold as the holo-PipS structure (RMSD of 0.588 Å over all Cα atoms^23^); however, the electron density on the distal axial site of the heme iron was significantly different (**Extended Data Fig. 2a**). We found that PipS is active in the crystallization conditions, but significant L-**1** remains after 1 hr (**Extended Data Fig. 2b**), meaning a mixture of molecules could be present. We observed that the electron density best corresponds to the δ-nitrogen of L-**1** coordinating the heme iron (**Extended Data Fig. 2a**). In this case, we observed a 2.0 Å distance between the heme iron and the nitrogen (**Fig. 1e, Extended Data Fig. 2c**). In the structure with L-**1**, the α-amino nitrogen of the substrate is best modelled with a 2.9 - 3.0 Å distance to the heme propionate group, consistent with a hydrogen bond. Collectively, these two interactions place the two nitrogens within 5.1 – 5.2 Å of one another (**Extended Data Fig. 2d**).

To determine which residues are important for N-N bond formation, we created a variant library of tagless PipS variants, targeting conserved residues in the distal axial site in proximity to L-**1** (**Supplementary Fig. 5-8**). We found that most of the variants (Y7F, Y99F, Y155F, and E182A) still produced L-**2**, albeit at reduced levels (**Extended Data Fig. 3, Supplementary Fig. 9a**). The T107A variant only produced ∼3 % of L-**2** when compared to the wild-type, but both the K178A and the T107A/K178A double variant did not produce any L-**2** (**Extended Data Fig. 3, Supplementary Fig. 9a**). To further probe the role of T107 and K178, we generated additional variants T107V, T107S, T107C, and K178M. We observed that all of these variants did not produce any L-**2**, except T107S which was able to generate ∼28 % L-**2** when compared to the wild-type.

One role that a K178-T107 dyad could play in the enzyme catalyzed reaction is to promote loss of the hydroxyl group, which is pointing towards the K178-T107 dyad and is not present in the final product L-**2**. To interrogate whether the K178-T107 dyad is key to early steps in the reaction, we studied the reaction of PipS with *N*^5^-OH-aminovaleric acid (**3**), a natural substrate analog without the α-amine (**Fig. 2a**, **Supplementary Fig. 10**). Because this substrate lacks the α-amine, it should be incapable of N-N bond formation, however, it still contains the hydroxylamine functional group that could interact with the K178-T107 dyad. We found that **3** was completely consumed when incubated with both as-isolated and dithionite-reduced PipS, with concomitant formation of ammonia (**Fig. 2b**, **Supplementary Fig. 11**). In aerobic conditions, this reaction proceeded rapidly for the dithionite-reduced enzyme and more slowly using as-isolated enzyme. Loss of ammonia from **3** could potentially occur through two steps: (1) loss of the hydroxyl as water, giving an imine, and (2) hydrolysis of the imine to release ammonia. To verify that this pathway leads to ammonia production in this way, we used DNPH derivatization to demonstrate that a *m/z* peak consistent with co-product 5-oxopentanoic acid is also produced along with ammonia from PipS + **3** (**Supplementary Fig. 12**). We then incubated all the PipS variants with **3**. Similar to the result with L-**1**, we observed that K178 and T107 are crucial for the dehydration reaction with **3** (**Supplementary Fig. 13**).

**Fig. 2.**
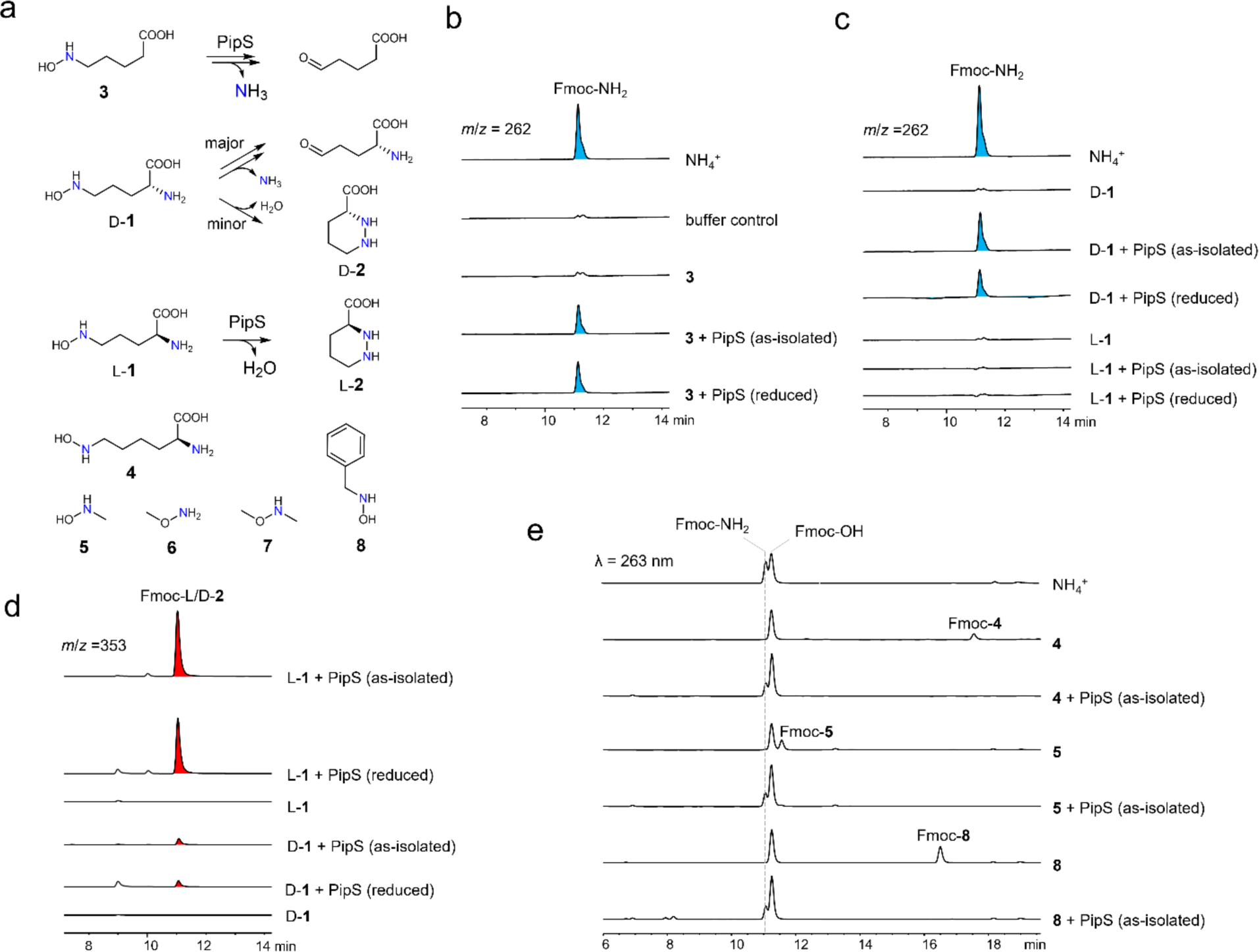
PipS has dehydratase activity on *N*-substituted hydroxylamines. **a**, Substrate analogs used in this study. **b**, LC-MS analysis of the reaction mixtures of PipS with **3** after Fmoc-Cl derivatization. Note: extracted ion chromatogram (EIC) of *m*/*z* 262 ([M+Na]^+^ ion for Fmoc-NH_2_) are shown. **c**, LC-MS analysis (EIC= *m*/*z* 262) of reaction mixtures of PipS with D-**1** or L-**1**. **d**, LC-MS analysis (EIC= *m*/*z* 353, [M+H]^+^ ion for Fmoc-D/L-**2**) of reaction mixtures of PipS with D-**1** or L-**1**. **e**, HPLC analysis of the reaction mixtures of PipS with N-substituted hydroxylamines (**4**, **5**, **8**). Each reaction was performed in triplicate and representative results are shown. Fmoc-OH is a hydroxylated product of Fmoc-Cl.

Intrigued by this result, we investigated the reaction of PipS with another substrate analog D**-1**. Previously, we observed with KtzT that only 10 % of the D**-1** underwent N-N cyclization to generate D-**2**^15^. Now knowing that PipS also possess deamination activity, we analyzed the reaction mixtures of D**-1** with PipS, and successfully detected both D-**2** and ammonium, with ammonium as the major product (**Fig. 2c and 2d**). Additionally, we also observed the production of dehydroproline, the predicted co-product from ammonia in this reaction (**Supplementary Fig. 14**). The observation that D-**1** can undergo both deamination and N-N bond formation suggests that these two reaction routes may share early catalytic intermediates. We further found that incubation of L-*N*^6^-OH-lysine (**4**) with PipS afforded ammonia (**Fig. 2e**). When the reaction of L-**1** with PipS is analyzed, no ammonia production is observed (**Fig. 2c**). Finally, we tested the PipS variants for production of ammonia, and observed only K178A had slightly increased amounts of ammonia present (**Supplementary Fig. 9b**).

To further interrogate this unique reaction with hydroxylamine-containing substrates, we next extended our investigation to another three hydroxylamines, *N*-methylhydroxylamine (**5**), *O*-methylhydroxylamine (**6**) and *N*,*O*-dimethylhydroxylamine (**7**). Among them, we found that only **5** can be processed by PipS, affording ammonium as a product (**Fig. 2e, Extended Data Fig. 4a**), which might indicate that the *N*-hydroxyl group is required and directly involved in the PipS catalyzed reaction mechanism. As the simplest *N*-substituted hydroxylamine, **5** could be converted to methanimine, which would spontaneously hydrolyze to ammonia and formaldehyde. Indeed, we successfully detected production of formaldehyde in the reaction mixture of **5** with PipS, however, we found that when 1000 nmol of **5** was consumed, only 408 ± 10 nmol (mean ± s.d, n = 3) of formaldehyde was detected, indicating that more chemical transformations might occur in the above assay mixture. To get a fuller picture of PipS-mediated turnover of *N*-substituted hydroxylamines, we used *N*-benzylhydroxylamine (**8**) as a model substrate for easier detection of any additional product(s) (**Fig. 2a**). Detailed analysis of this model system further confirmed that PipS catalyzes deamination on unnatural *N*-substituted hydroxylamine substrates (**Extended Data Fig. 4b-f, Supplementary Fig. 15**).

Analysis of the assay mixtures of L-**1** or **8** with four diverse PipS homologs demonstrate that all have N-N bond-forming activity on L-**1** and dehydration activity on **8** (**Supplementary Fig. 16**). Heme-dependent dehydration by PipS is reminiscent of aldoxime dehydratase Oxd, which converts oximes to nitriles and also involves the direct binding of substrate to the heme iron^24^. Previous studies on Oxd have shown that the first step of oxime dehydration involves the protonation of the hydroxyl group by active site residues to release a water molecule forming a Fe-N·intermediate. We deduce that PipS might use similar mechanistic transformations as aldoxime dehydratase for dehydration. We tested the ability of Oxd to react with **8**, and observed that as-isolated Oxd successfully mediated the conversion of **8** to ammonium and **9** (**Supplementary Fig. 17**). Furthermore, we found that dithionite-reduced PipS can also mediate oxime dehydration. Altogether, these data support the conclusion that PipS and Oxd share the reaction route for the dehydration reaction.

Based on previous results for Oxd-related systems and considering our current experimental data, we performed computational modelling to explore a possible reaction mechanism of PipS and the role of the T107/K178 dyad, first considering the ferrous-heme PipS. We analyzed in detail the arrangement of T107 and K178 using MD simulations when L-**1** is bound to the ferrous-heme PipS (**Fig. 3b**, **Extended Data Fig. 5**, and **Supplementary Figs. 18-21**). MD simulations demonstrated that these residues are well-positioned and preorganized for reaction with the hydroxylamine substrate, establishing persistent H-bond interactions. QM/MM calculations (see computational details and **Supplementary Fig. 26**) showed that these two residues can act as a proton shuttle to activate the hydroxylamine group. First, the T107/K178 catalytic dyad acts as proton donor source to activate the *N*-hydroxyl group and promote the loss of a water molecule as a leaving group (PipS-**TS1**, ΔG^‡^ = 21.2 kcal·mol^−1^, lowest in energy open-shell singlet (OSS) electronic state), to generate a protonated Fe-nitrenoid species (PipS**-Int1**, **Figs. 3a,c,e** and **Supplementary Figs. 26** and **30**). Deprotonation at the FeN-H position of the nitrenoid PipS-**Int1** can subsequently take place by a proton shuttle formed by the released water molecule and the T107/K178 catalytic dyad (PipS-**TS2**, ΔG^‡^ = 7.3 kcal·mol^−2^, lowest in energy triplet (T) electronic state), restoring the initial protonation states of the catalytic residues and generating a Fe-nitrene intermediate species, PipS-**Int2**, with a triplet ground state and a radical centered on the nitrogen atom (**Figs. 3a,c,e**, and **Supplementary Fig. 28**). On the other hand, QM/MM calculations considering a ferric-heme oxidation state describe a much higher barrier for the first water release step (PipS-ferric-**TS1**, ΔG^‡^ = 42.1 kcal·mol^−2^, **Extended Data Fig. 6**, and **Supplementary Figs. 31-33**) leading to a rather high in energy nitrenoid intermediate PipS-ferric-**Int2**. This result suggests that ferric PipS is not likely catalytically active toward the hydroxylamine substrate through this proposed mechanism. These results are in line with previous computational predictions for the related aldoxime dehydratase activity of Oxd in its ferrous vs. ferric oxidation states.^25^

**Figure 3.**
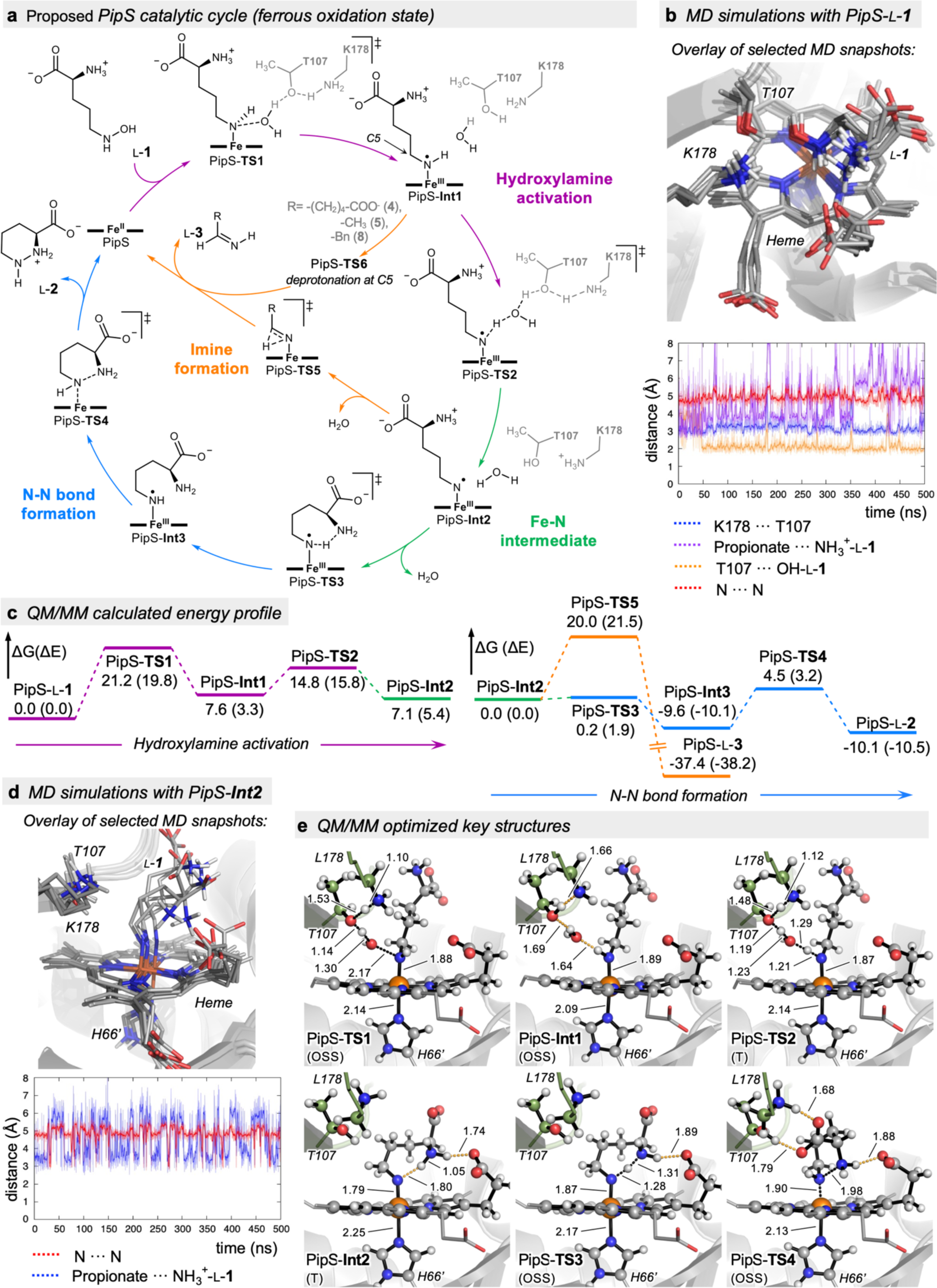
PipS proposed catalytic cycle and role of K178 and T107 in the PipS reaction. **a**, Proposed PipS catalytic cycle for N-N bond formation and dehydration of hydroxylamines. **b**, Conformational analysis of K178/T107 catalytic dyad and L-**1** substrate bound complex (PipS-L-**1**) from MD simulations. K108 - T107 - OH(L-**1**) H-bond interactions are found to be highly persistent along MD trajectories (see computational methods section in the SI and **Supplementary Figs. 18-21**). Interactions between L-**1** and proximal residues of PipS active site are analyzed in **Extended Data Fig. 5**. **c**, QM/MM calculated energy profile for the proposed hydroxylamine activation, and competing N-N bond forming and dehydration pathways. Only the energies of the lowest in energy electronic states for each stationary point are shown. A complete description of all the studied electronic states is reported in **Extended Data Fig. 6** and **Supplementary Figs. 26, 27**, and **30**). **d**, Conformational analysis of key Fe-nitrenoid intermediate PipS-**Int2** bound complex from MD simulations. The Fe-nitrenoid intermediate bound in PipS explores alternative conformations during MD trajectories, where the distance between the α-amine and the *N*-nitrene can be reduced up to ca. 3Å (see **Supplementary Figs. 22-25**). **e**, Lowest in energy QM/MM optimized key structures (transition states (TSs) and intermediates) from proposed PipS catalyzed hydroxylamine L-**1** activation and N-N bond formation pathways (OSS: Open-shell singlet; T: Triplet electronic states. See computational methods section in the SI, and **Supplementary Figs. 26** and **27**).

At the same time, hydroxylamines are well-known to catalyze reduction of ferric porphyrins in model studies^26^. To further characterize whether the heme cofactor in PipS can be reduced by hydroxylamine-containing substrates, we focused our analysis of the inactive T107A/K178A double variant with L-**1**. We observed that in aerobic conditions in the presence of L-**1**, the Soret band shifts from 407 nm to 422 nm (**Extended Data Fig. 7, Supplementary Figure 34**). To further characterize this species, we undertook EPR and rRaman studies on the reaction of T107A/K178A-PipS with L-**1**. The EPR signal decreases upon addition of L-**1**(**Extended Data Fig. 7**) and the rRaman spectrum exhibits a band at 1377 cm^−2^ (**Supplementary Figure 35a**), implying that L-**1** reduces the ferric iron to ferrous iron. To further probe the nature of this species, reactions of PipS WT with isopropylhydroxylamine and 2-nitropropane were analyzed with rRaman and MCD spectroscopy (**Supplementary Figure 36**), which allowed us to directly compare with previous studies on other heme enzymes that used these substrates^27–30^, revealing that the species with the 422 nm Soret band corresponds to an alkyl-nitroso complex of the heme. This species is generated from an initial single-electron transfer from the hydroxylamine to the ferric heme, followed by O_2_-dependent formation of a ferrous-nitrosoalkane complex.^28^ Combined with the findings from our previous kinetic studies on KtzT showing that the reaction rate for the reduced enzyme is significantly faster than that of the ferric enzyme,^15^ we suggest the genuine *in vivo* PipS-catalyzed reaction may be the reaction between L-1 and ferrous PipS, whereas the ferric PipS-catalyzed reaction may be an *in vitro* side reaction, albeit one that is potentially more useful in biocatalysis. Whether and how substrate facilitates conversion of the ferric, wildtype PipS will have to be interrogated in future mechanistic studies.

We next computationally explored how N-N bond formation could take place from PipS-**Int2** Fe-nitrenoid intermediate. MD simulations revealed that PipS-**Int2** intermediate can explore conformations where the distance between the α-amine and the *N*-nitrene is reduced up to ca. 3 Å. This is triggered by the strong interaction between the protonated α-amine and one propionate group of the heme that is strategically placed pointing inwards to the active site (**Fig. 3a,d** and **Supplementary Fig. 22**). QM/MM calculations indicate that, when PipS-**Int2** is formed, a quick proton transfer from the protonated α-amine to the nitrene N-Fe center can take place when they are close enough (PipS-**TS3**, ΔG^‡^ < 1.0 kcal·mol^−2^ for the lowest in energy OSS and T states, **Fig. 3c**, and **Supplementary Fig. 27**). This proton transfer activates the nucleophilic α-amine and enhances the electrophilic character of the protonated nitrene, favoring an efficient N-N bond formation through a nucleophilic attack (PipS-**TS4**, ΔG^‡^ = 14.1 kcal·mol^−2^ for the lowest in energy OSS state, although it is barrierless in the closed-shell singlet CSS state, **Fig. 3a,c** and **Supplementary Figs. 27**). The overall rate-limiting step of the proposed mechanism corresponds to the initial hydroxylamine activation and loss of a water molecule, further supporting the capture of L-**1** in the active site of PipS by crystallography, rather than a more reactive nitrenoid-type intermediate.

The existence of a Fe-N nitrenoid intermediate in the PipS catalytic cycle is in line with the formation of a new N-N bond in L-**1** or the formation of the corresponding imine when different unnatural substrates are considered (**Fig. 3a,c**, **Supplementary Figs. 27-30**). Once the nitrenoid PipS-**Int2** intermediate is formed, a tautomerization of the Fe-nitrenoid to a Fe-imine species could complete the hydroxylamine dehydration pathway. This tautomerization is proposed to be rate-limiting in related abiological heme-nitrene mediated biocatalytic cycles^31^. Computations indicate that the direct formation of the Fe-imine species from PipS-**Int2** via a direct H-migration from C5 to N5-Fe is energetically feasible, although it is less favorable than N-N bond formation (PipS-**TS5**, ca. ΔG^‡^ = 20.0 kcal·mol^−2^ for the lowest in energy CSS, **Fig. 3c** and **Supplementary Figs. 27** and **30**). On the other hand, during the course of PipS-**Int2** formation when the PipS-**Int1** intermediate is generated, the H_2_O-T107-K178 catalytic machinery could deprotonate at the C5 position instead at NH-Fe to assist to the formation of the PipS-L-**3** imine species (PipS-**TS6**, **Fig. 3a**); however, an unfavorable conformational rearrangement of the intermediate would be required to approach the H-C5 position to the H_2_O-T107-K178 proton shuttle (**Extended Data Fig. 8, and Supplementary Fig. 29**). Nonetheless, these dehydration imine formation pathways become possible after the hydroxylamine is activated when no additional amine groups are present in the substrate molecule or when substrates are more flexible in the binding pocket. These proposed plausible mechanisms reconcile our observation that imine formation predominates for substrates D**-1, 4**, **5**, and **8**.

In this work we unveiled the structure of PipS and interrogated the role of the hydroxyl in the substrate L-**1** and its analogs, to reveal that the enzyme can catalyze both dehydration reactions and the N-N bond-forming reaction. Both reactions critically rely on the role of a Lys/Thr pair, which is widely conserved among the PipS homologs and proposed to activate hydroxylamine substrates to form a key iron-nitrenoid intermediate species. A plausible mechanism for N-N formation is also proposed, solving the mystery of how nature can bring two nucleophilic atoms into a bond. Specifically, the nucleophilic character of the hydroxylamine nitrogen can be reversed via formation of an electrophilic nitrenoid, which is conformationally preorganized by the protein active site to bring the nucleophilic α-amine close enough to react forming a new N-N bond. Recently, a new cytochrome P450 has been reported to catalyze a nitrene transfer reaction in the biosynthesis of benzastatin from an *N*-acetoxy substrate^32^. At the same time, directed evolution has been used to engineer several heme^31,33–36^ and non-heme^34,37^ proteins, and artificial cofactor-based metalloenzymes^38^ for abiological nitrene-mediated reactions, which are currently limited to azide derivatives and *N*-acetoxy substrates as nitrene precursors (**Extended Data Fig. 9**). We suggest that PipS and related family of enzymes could also be nitrene transferases that utilize a newly proposed activation mechanism of hydroxylamines that allows their direct use as nitrene precursors in aerobic conditions, expanding the substrate scope of biocatalytic nitrene transfer reactions and catalytic repertoire toward N-N bond formation, paving the way for further reaction development and protein engineering.

## Acknowledgements

This work was supported by financial support from the Michael Smith Foundation for Health Research (16006 to M.A.H. and 16776 to K.S.R.), the Natural Sciences and Engineering Council of Canada (RPGIN-2021-02626 to K.S.R.), the National Natural Science Foundation of China (32122005, 31872625) to Y.-L.D., the Zhejiang Provincial Natural Science Foundation (LR19C010001) to Y.-L.D., the Generalitat de Catalunya AGAUR for a Beatriu de Pinós H2020 MSCA-Cofund 2018-BP-00204 project (to M.G.B.), the Spanish MICINN (*Ministerio de Ciencia e Innovación*) for PID2019-111300GA-I00 project (to M.G.B) and the Ramón y Cajal program via the RYC2020-028628-I fellowship (to M.G.B), the Spanish MIU (*Ministerio de Universidades*) for a predoctoral FPU fellowship FPU18/02380 (to J.S.), and the National Science Foundation of the USA (CHE-2002885 to N.L.). Crystallographic data were collected at the Canadian Light Source and at the Stanford Synchrotron Radiation Laboratory. The authors thankfully acknowledge the computer resources at *MinoTauro* and the technical support provided by Barcelona Supercomputing Center BSC-RES (RES-QSB-2019-3-0009 and RES-QSB-2020-2-0016).

## Author Contributions

M.A.H. carried out structural, mutagenesis, and biochemical studies. X.S. and Y.-L.D. carried out biochemical studies on substrate analogs. J.S. and M.G.B. carried out DFT, QM/MM and MD computational studies. J.H. carried out by EPR, rRaman, and MCD analysis. X.S., Y.-L.D. and T.P. carried out synthetic work. N.L. supervised and analyzed EPR, rRaman, and MCD work. M.G.B. supervised and analyzed computational studies. Y.-L.D. supervised and analyzed biochemical studies. K.S.R. supervised and analyzed structural and biochemical studies. M.A.H., M.G.B., Y.-L.D. and K.S.R. wrote the paper with contributions from all authors.

---

**Extended Data Figure 1.**
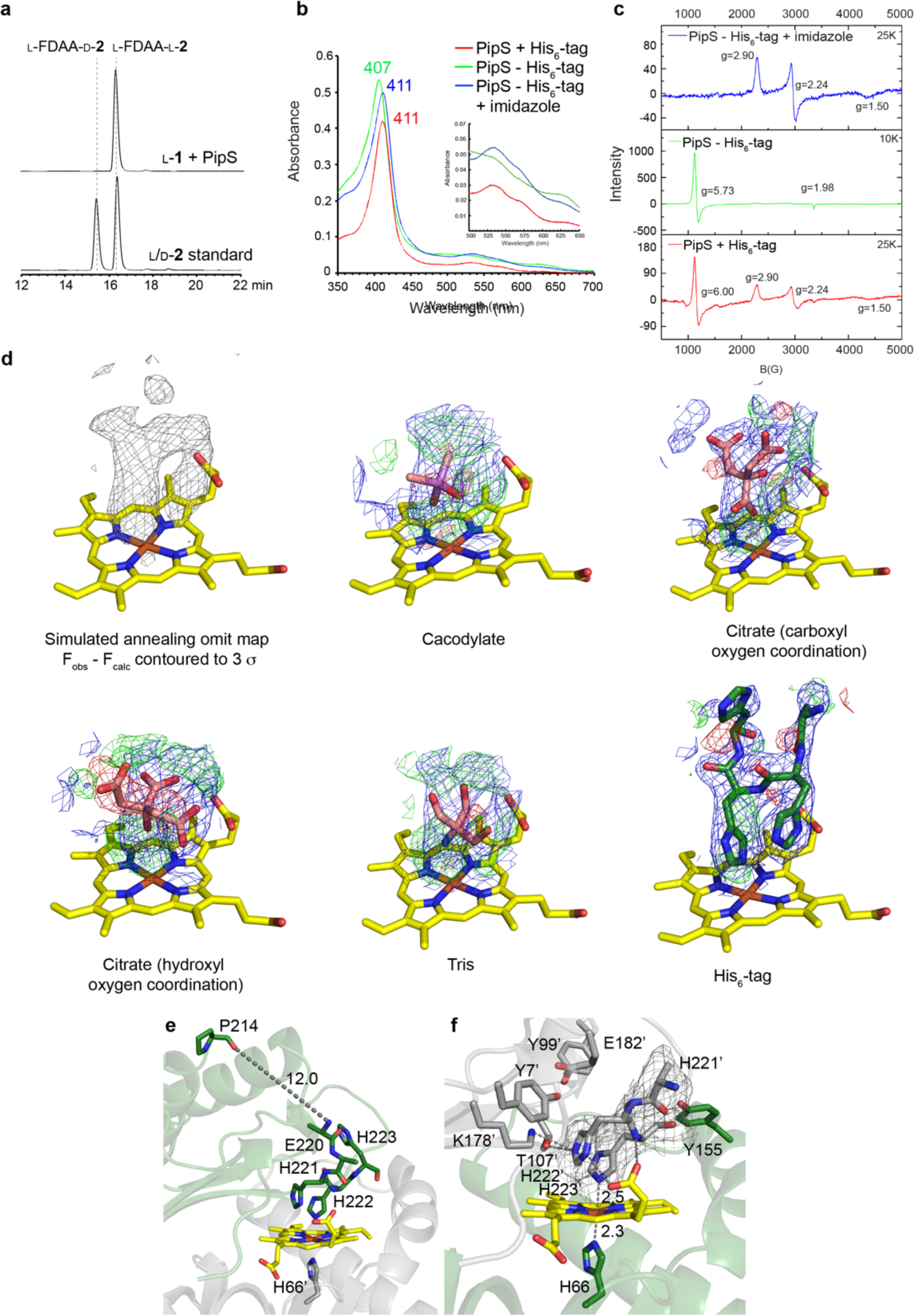
In vitro biochemical characterization of PipS and the C-terminal His_6_-tag coordinates the distal site of the heme iron. **a**, PipS reaction with L**-1** gives L-**2**. LC-MS analysis (EIC for *m/z* of 383, corresponding to L-FDAA-L/D**-2**) of 1-fluoro-2,4-dinitrophenyl-5-L-alanine amide (L-FDAA) derivatization of the PipS reaction and the L/D**-2** standard. **b**, UV-Vis and **c**, EPR analysis of PipS with (+) and without (–) the His_6_-tag and without the His_6_-tag + 2mM imidazole. In the presence of the His_6_-tag, the EPR spectrum shows a mixture of axial HS ferric signals and a new species with g values of 2.90, 2.24, 1.50, typical for LS bis-His coordinated ferric heme,^39–41^ indicative of the His_6_-tag coordinating to the ferric heme. Further evidence for His coordination to the heme in this ferric LS species was obtained by addition of free imidazole to the holo-enzyme, which converts all of the heme into the same LS ferric species with g = 2.90, 2.24, 1.50. Further spectroscopic analysis of PipS with His_6_-tag can be found in SI Figure X. **d**, Modeling the His_6_-tag in the distal site of the PipS holo-structure. Different compounds from the crystallization solution or protein buffer modeled into the distal site of the heme iron with one round of refinement performed. Blue mesh represents the 2F_obs_ – F_calc_ density map contoured to 1 σ while the green and red mesh represent the F_obs_ – F_calc_ map contoured to +3 and –3 σ, respectively. **e**, Proximity of the C-terminus to the distal site of the heme iron. **f**, Active site architecture for the opposite monomer (heme coordinating H66 from monomer A) from the PipS dimer for the holo-PipS-His_6_ structure. The grey mesh represents the F_obs_ – F_calc_ simulated annealing omit map for the partial His_6_-tag contoured to 3 σ. PipS monomer A is shown in green, monomer B in grey, heme in yellow, and potential ligands in pink with nitrogen in blue, oxygen in red, iron in orange, and arsenic in light purple. Key distances are given in angstroms (Å).

**Extended Data Figure 2.**
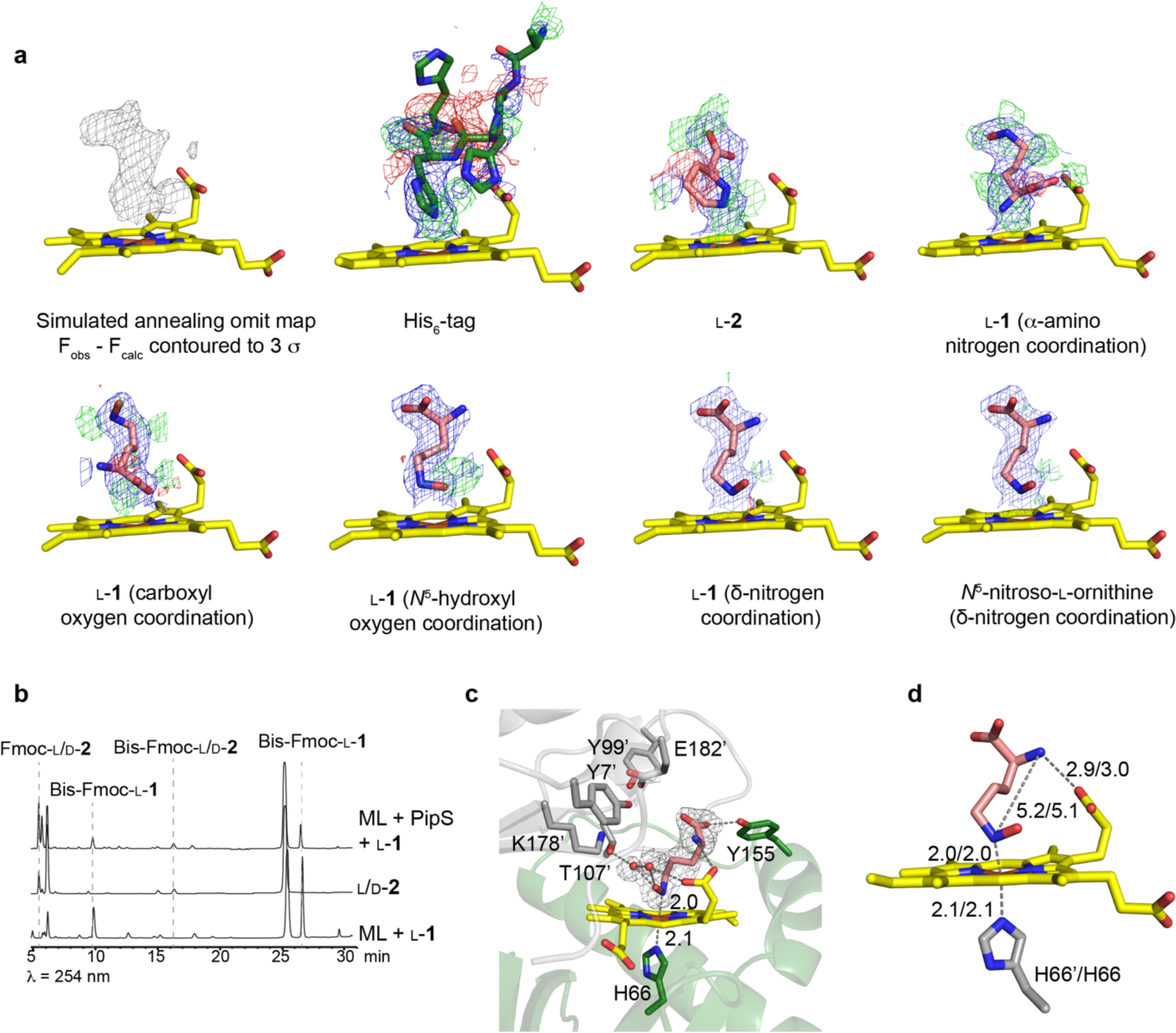
PipS-L-1-structure. **a**, Different compounds were modeled into the distal site of the heme iron and one round of refinement performed. Blue mesh represents the 2F_obs_ – F_calc_ density map contoured to 1 0 while the green and red mesh represent the F_obs_ – F_calc_ density map contoured to +3 and –3 0, respectively. **b,** PipS reaction in crystallization mother liquor (ML). A 1:1 ratio of 4 mg mL^−2^ PipS (equivalent to one large crystal in a 1 uL drop) and ML (0.75M sodium citrate, 0.1 M sodium cacodylate pH 6.5) was mixed. Then, 2 μL of the PipS:ML was added to 2 μL of 1 M L-**1** and incubated at room temperature for 1 hr. The reaction was diluted to 100 μL with sodium phosphate buffer and immediately quenched with 500 μL acetonitrile then derivatized with Fmoc-Cl and analyzed by HPLC. The reactions were performed in triplicate with and representative reaction shown. **c**, Active site architecture for the opposite monomer (heme coordinating H66 from monomer A) from the PipS dimer for the PipS-L-**1**-structure. The grey mesh represents the F_obs_ – F_calc_ simulated annealing omit map for L-**1** and both ordered waters contoured to 3 0. **d**, Heme and ligand distances for L-**1** for both active sites in the PipS-L-**1**-structure along with the distances between the α- and δ-nitrogens. PipS monomer A is shown in green, monomer B in grey, heme in yellow, and L-**1** and other potential ligands in pink with ordered waters as red spheres. Nitrogen is blue, oxygen is red, and iron is orange. Key distances are given in angstroms (Å).

**Extended Data Fig. 3.**
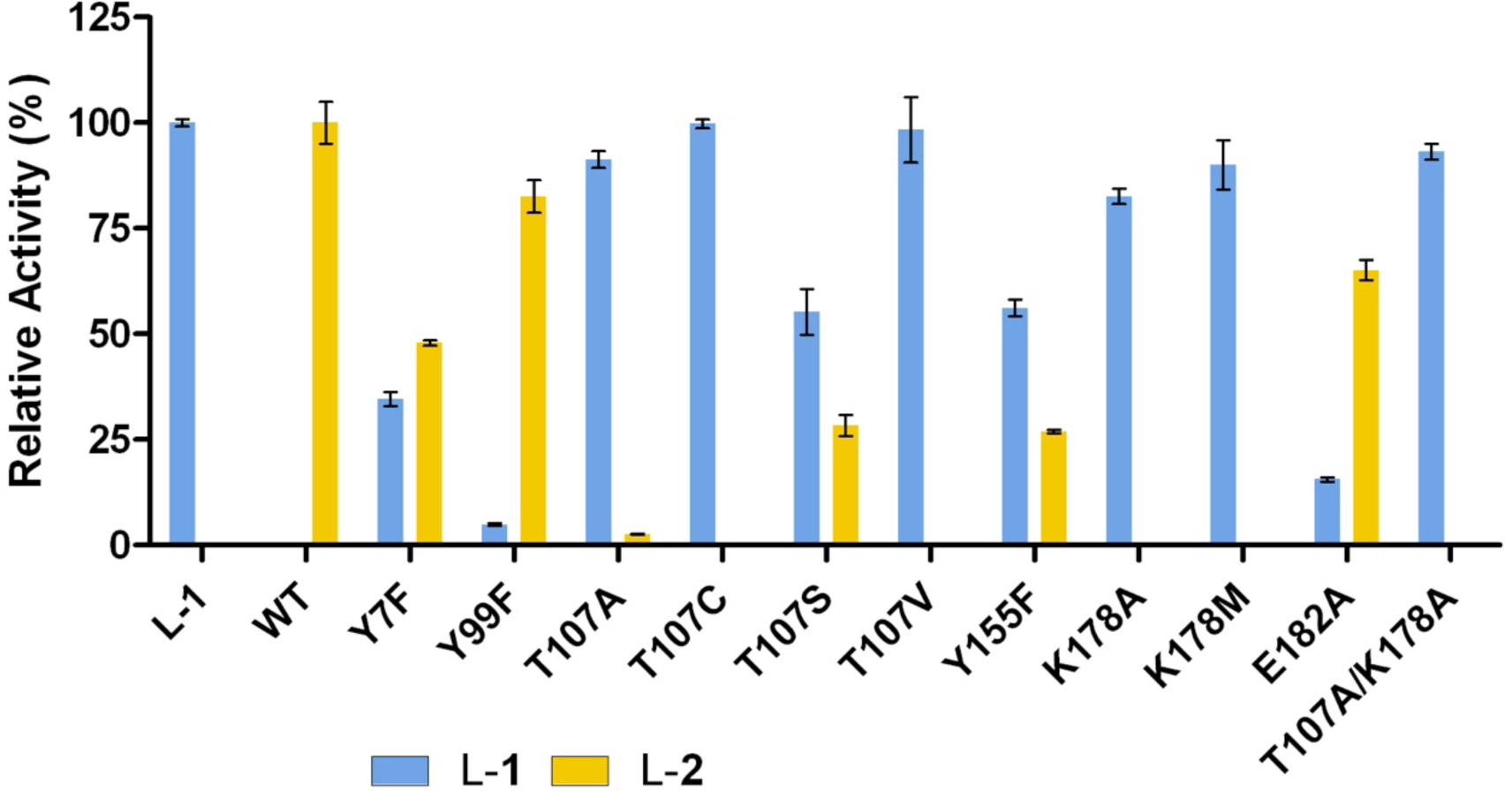
Enzyme activities for PipS variants. **a**, Reactions of L-**1** with PipS and its variants. Relative activities when compared to L-**1** with no enzyme or L-**2** produced by PipS WT. Each reaction was performed in triplicate.

**Extended Data Fig. 4.**
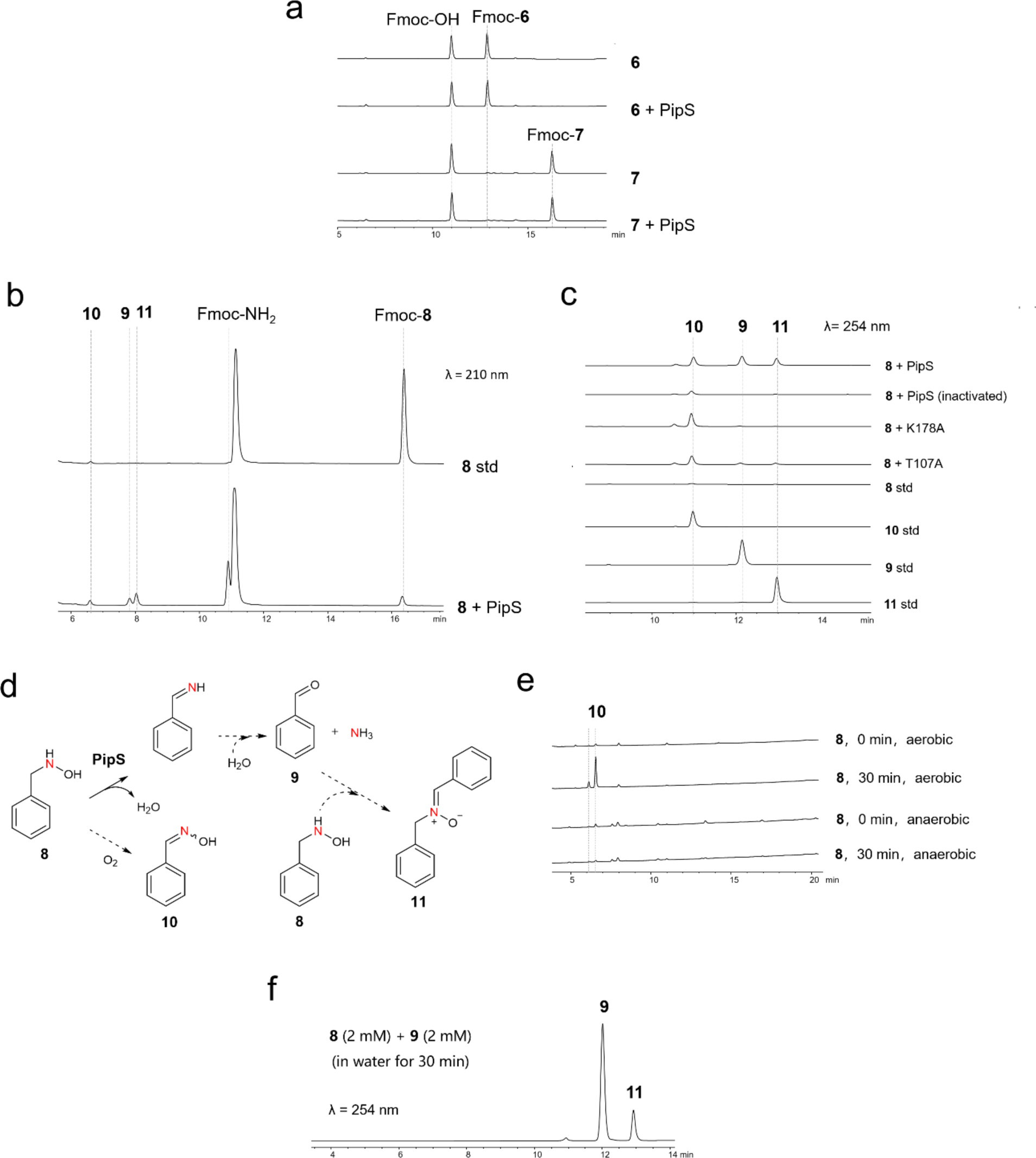
Analysis of PipS reactions with various substrates (6, 7 and 8). **a,** HPLC analysis (λ=263 nm) of the reaction mixture of PipS with *O*-methylhydroxylamine (**6**) and *N*,*O*-dimethylhydroxylamine (**7**) after Fmoc-Cl derivatization. **b,** HPLC analysis (λ=210 nm) of the reaction mixture of PipS with *N*-benzylhydroxylamine (**8**) after Fmoc-Cl derivatization revealed the production of ammonia, **9**, **10** and **11**. **c**, HPLC analysis (λ=254 nm) of the reaction mixtures of **8** with PipS and its variants (T107A and K178A). Product **9** was identified as benzaldehyde and **10** as benzaldehyde oxime by comparison with standard compounds. Compound **11** was prepared through scale-up enzymatic synthesis and characterized as *N*-benzyl-α-phenylnitrone by extensive NMR analysis (see **Supplementary Fig. 8**). **d**, Reaction routes in the reaction of PipS with **8**. Note: We found that **10** derived from an O_2_-dependent non-enzymatic oxidation (**e**) and **11** can be generated by direct incubation of the substrate **8** with product **9** (**f**). Together, these data demonstrate that PipS catalyzes deamination on **8** to give **9**. **e**, non-enzymatic generation of **10**. **f**, non-enzymatic generation of **11**. Production of **11** was observed upon incubation of **8** and **9** in water.

**Extended Data Fig. 5.**
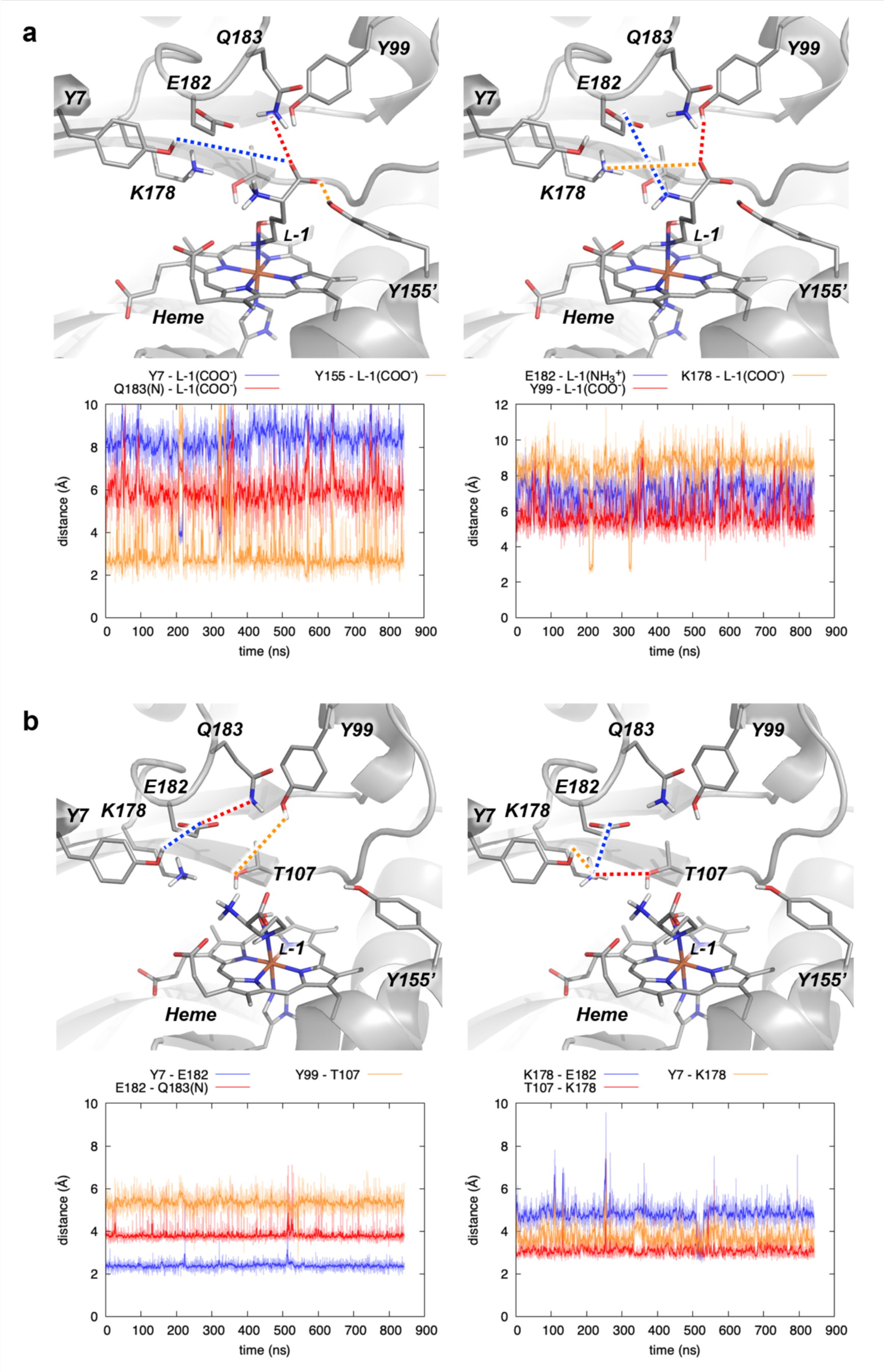
Interactions between L-1 and proximal residues of PipS active site explored by MD simulations. **a**, Key distances between Y7, Y99, Y155’, E182, Q183, and K178 polar residues and L-**1** substrate when bound in PipS (Replica 1, active site from monomer 1). **b**, H-bond network of interactions between these proximal residues (Replica 1, active site from monomer 1). Distances are given in angstrom (Å). Although there exists a stable H-bond network between these residues, only Y155’ can establish persistent polar interactions with the carboxylic group of L-1. See **Supplementary Figs. 18** and **19** for additional MD replicas and analysis of these interactions in active site from monomer 2.

**Extended Data Fig. 6.**
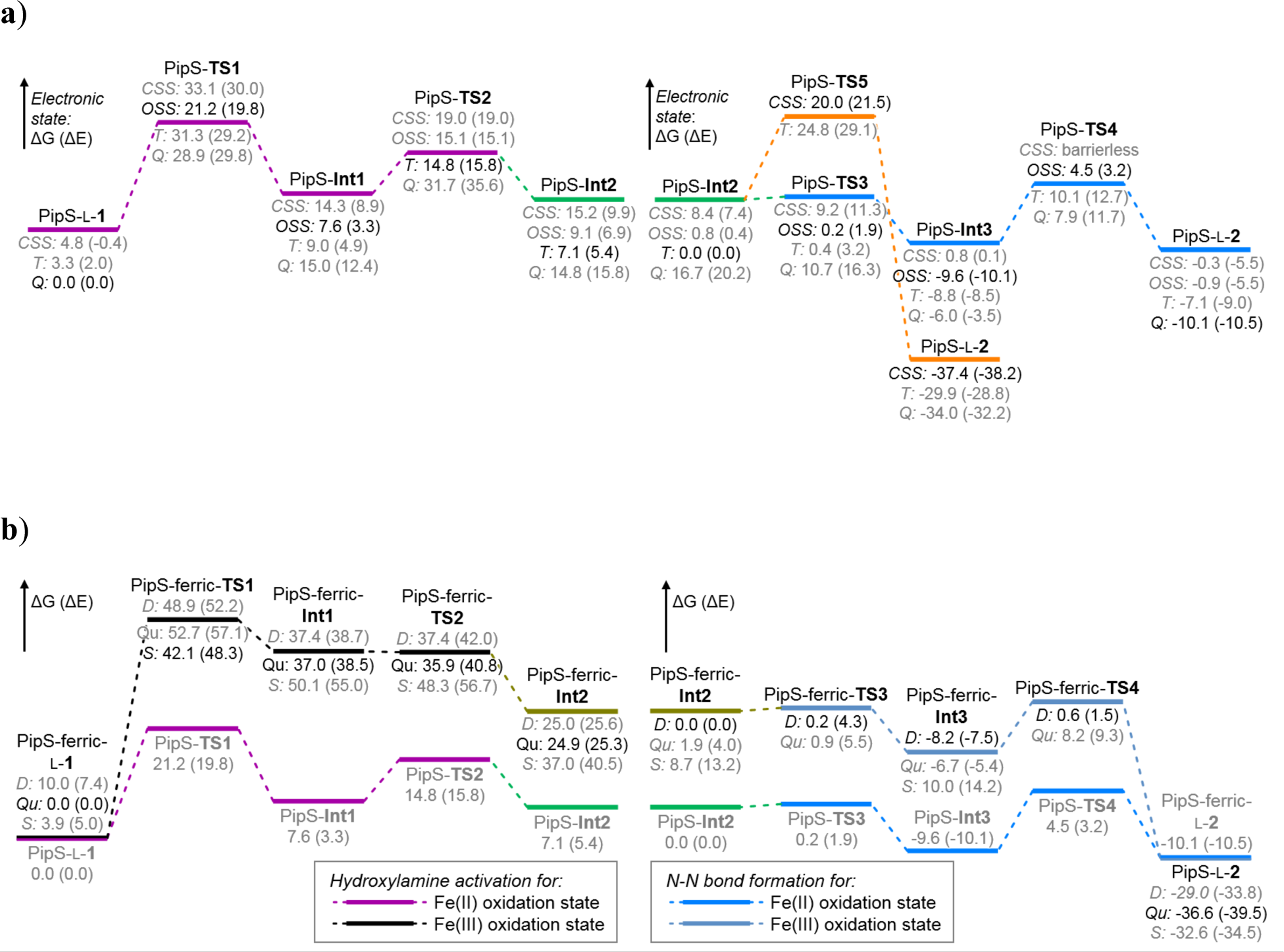
QM/MM calculated energy profile for PipS-catalyzed hydroxylamine activation (left) and N-N bond formation (right) considering a) ferrous (Fe(II)) and b) ferric (Fe(III)) oxidation states. Potential Energy Surface (PES) for PipS-catalyzed hydroxylamine activation and N-N bond formation considering PipS **a**) ferrous (Fe(II)) and **b**) ferric (Fe(III)) as resting oxidation states, respectively. For each stationary point different energetically accessible electronic states have been considered: **a**) closed-shell singlet (CSS), open-shell singlet (OSS), triplet (T), and quintet (Q); and **b**) doublet (D), quartet (QU), and sextet (s). See **Supplementary Figs. a) 26-30** and **b) 31-33** for additional details on the QM/MM optimized structures. The equivalent PES profile for the ferrous (Fe(II)) oxidation state is provided for comparison next to the Fe(III) one.

**Extended Data Figure 7.**
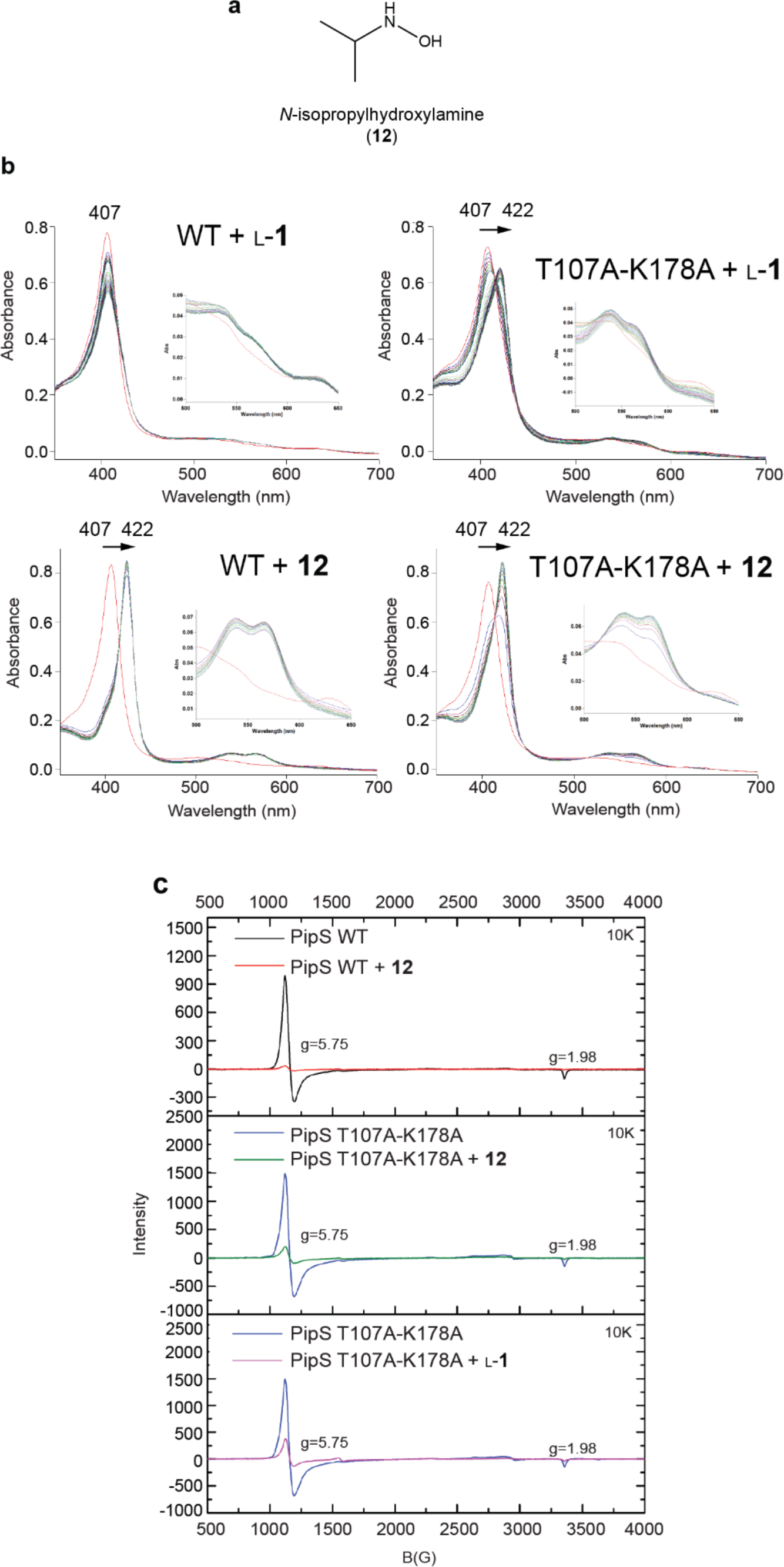
Characterization of the 422 nm peak. **a**, Structure of *N*-isopropylhydroxylamine. **b**, UV-Vis analysis of reactions with 6 μM PipS WT or T107A-K178A + 5 mM **12** for 1 h or L-**1** for 4 h. **c**, EPR analysis of 1472 μM PipS WT or PipS-T107A-K178A incubated with 20 mM **12** or L-**1** overnight at 4 °C.

**Extended Data Fig. 8.**
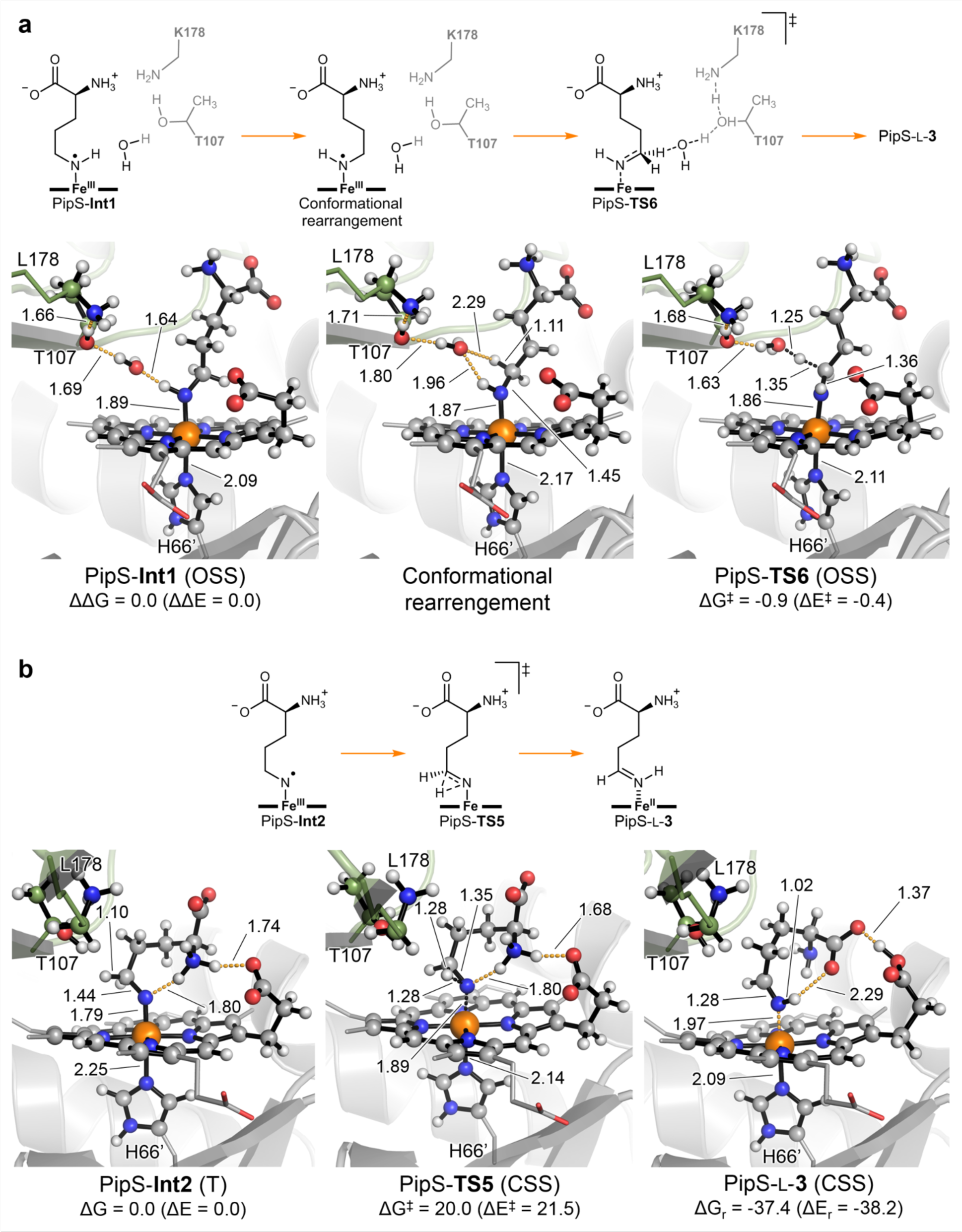
Possible mechanistic pathways for imine PipS-L-3 formation explored using QM/MM calculations. Two possible routes for imine formation are explored: **a**, Imine formation via direct deprotonation of nitrenoid intermediate PipS-**Int1** at C5 position by T107-K178 and the previously released water molecule from PipS-**TS1**. In this case, nitrenoid intermediate PipS-**Int1** should undergo a large and energetically unfavorable conformational rearrangement (see **Supplementary Fig. 29**) before C5 deprotonation can take place through PipS-**TS6**. Although C5 deprotonation reaction barrier is lower than competing nitrenoid deprotonation through PipS-**TS2**, the large conformational rearrangement of PipS-**Int1** required for deprotonation at C5 contrasts with the high preorganization of the H_2_O-T107-K178 catalytic machinery towards deprotonation at NH position, thus favouring PipS-**TS2** to take place instead of PipS-**TS6**. **b**, Imine formation through a direct 1,2-H migration from C5 to N-nitrene center in intermediate PipS-**Int2**. Distances are given in angstrom (Å). Only lowest in energy optimized structures are shown, see **Supplementary Figures 26** and **29** for a complete QM/MM analysis considering different spin states and further details.

**Extended Data Fig. 9.**
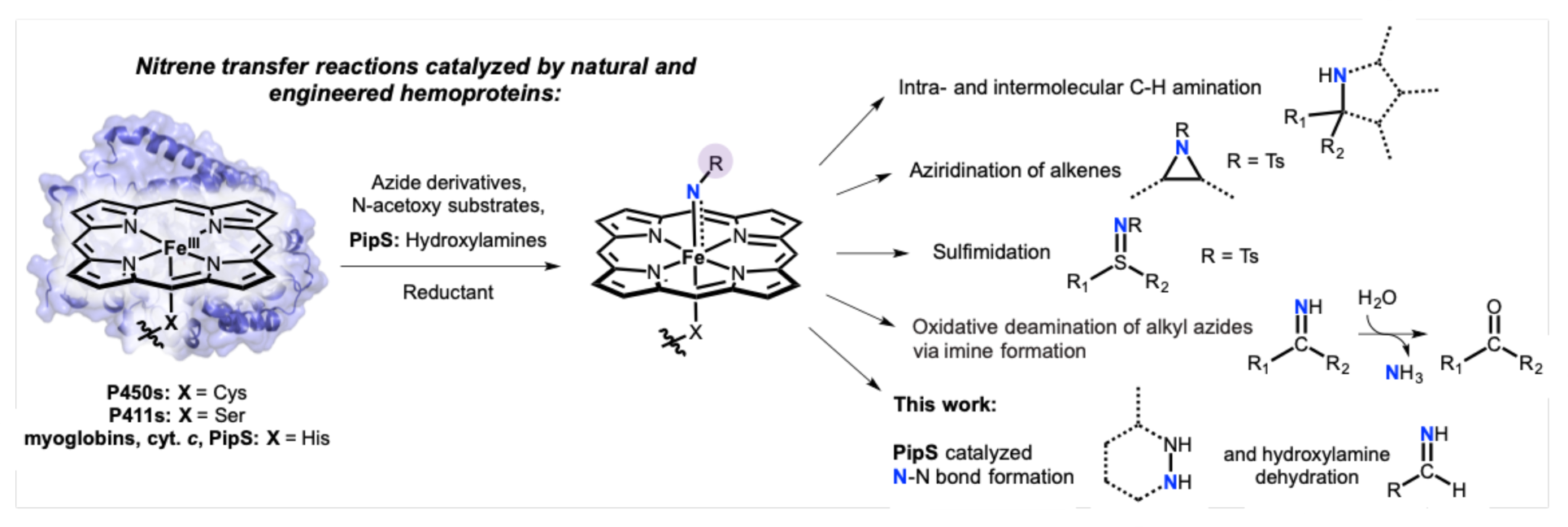
Enzymatic nitrene transfer reactions catalyzed by natural and engineered hemoproteins.

