## Supplemental Data for "Structural and mechanistic views of enzymatic, heme-dependent nitrogen-nitrogen bond formation"

This PDF file includes:

**Part I**: page 2
- Experimental Materials and Methods
- Supplementary Tables 1 to 3
- Supplementary Figures 1 to 17, and Figures 34 to 36

**Part II**: page 42
- Computational Methods and Protocols
- Supplementary Figures 18 to 33
- Supplementary Tables 4 to 7

**References** page 114

Supplementary Tables 8 to 11 containing QM/MM (only QM atoms) Cartesian Coordinates are reported as separate files.

### Part I

#### Materials and Methods:

*General Methods*:

Primers were ordered from Integrated DNA Technologies. DNA sequencing was performed by the NAPS Unit DNA Sequencing Facility at The University of British Columbia. Reagents were purchased from Sigma-Aldrich, Bio Basic Canada, Inc., Bio-Rad and New England Biolabs (NEB). *Streptomyces griseus* subsp. *griseus* NRRL F-5144 was obtained from the Agriculture Research Service (ARS) Culture Collection (NRRL). $N^5$-OH-aminovaleric acid and D-$N^5$-OH-ornithine were synthesized similarly as previously described or as described in the SI Methods.[1–3]

*Polyoxypeptin gene cluster identification*:

A multi-gene BLAST search[4] was used to identify KtzT homologs that contained a gene encoding for a KtzI homolog in the genomic neighbourhood. DNA sequences from *Streptomyces griseus* subsp. *griseus* NRRL F-5144 were downloaded from NCBI Assembly database (Accession number ASM71935v1). Contigs were aligned using Sequencher (www.genecodes.com). Overlapping sequences were observed for contigs 41 with 59 and 59 with 32. These contigs were assembled and DNA sequencing with primers 5′-GGATCAGCTCGGCCAGATA-3′ and 5′-CCCTGTTCCAGGTGATGTTC-3′ was used to validate this assembly. Not all of the polyoxypeptin gene homologs were contained in this contig assembly. Contig 40 also contained polyoxypeptin cluster gene homologs but did not have any overlapping sequence with contig 41 or 32. The 3′ end of contig 40 and 5′ end of contig 41 encoded for a *plyF* homolog suggesting there was missing sequence between contigs 40 and 41. Colony PCR with primers 5′-CGACGACCACCGCGTCGTGCA-3′ and 5′-GACTATCTGGCCTGGCTCTCCAC-3′ as well as internal primers 5′-CACCGATACGTCGAAGCTC-3′ and 5′-TGGAGTGTTTGGTGGGAGAT-3′ was used to determine the missing sequence and complete the contig assembly. A complete list of genes for this cluster is in **Supplementary Table 1**.

*L-$N^5$-OH-ornithine synthesis:*

Boc-ornithine (1.00 g, 4.3 mmol, 1.0 equiv) and 3 Å molecular sieves were added to a solution of potassium hydroxide (0.23 g, 4.5 mmol, 1.05 equiv) in methanol (10 mL). The reaction mixture was stirred at room temperature for 16 h, after which it was filtered and the residue washed with a further 5 mL of methanol. The filtrate was concentrated under reduced pressure, and the resulting off-white solid was used without further purification.

The residue was taken up in methanol (10 mL) and cooled to 0˚C over an ice-water bath. A solution of 3-chloroperbenzoic acid (1.50 g, 6.7 mmol, 1.5 equiv) in methanol (7 mL) was added to the reaction mixture over a period of 1 h, after which the reaction mixture was stirred at 0 °C for a further 1 h. The reaction mixture was filtered, the precipitate washed with methanol, and the combined filtrate concentrated under reduced pressure. The resulting solid was taken up in water (25 mL) and ethyl acetate (25 mL). The aqueous layer was acidified with 1 M HCl (15 mL), after which the layers were separated. The aqueous layer was extracted with ethyl acetate (2 × 25 mL) and the combined organics were washed with brine (50 mL), dried over $MgSO_4$, filtered, and concentrated under reduced pressure to yield a pale yellow solid, which was used without further purification.

The residue was taken up in dichloromethane (6 mL) and trifluoroacetic acid (6 mL) and stirred at room temperature for 1 h. The reaction mixture was concentrated under reduced pressure and the residue was taken up in ethyl acetate (15 mL). 10% DIPEA in THF was added until the solution had an approximate pH of 9. The mixture was cooled to 0 °C for 1 h, after which it was filtered to produce a yellow solid. The crude product was recrystallized in MeCN/$H_2O$ to yield a pale yellow solid, which was then taken up in TFA (13 mL) and water (2 mL). To this solution was added $CH_2Cl_2$ (15 mL), after which the mixture was stirred at 40 °C for 15 min, and then concentrated under reduced pressure. The residue was taken up in 1 M HCl (25 mL) and $CH_2Cl_2$ (15 mL) and stirred at room temperature for 1 h. The layers were separated and the aqueous phase was washed with further $CH_2Cl_2$ (30 mL) and hexanes (30 mL), after which it was lyophilized to yield L-**1** as a beige solid (0.265 g, 1.8 mmol, 40% overall yield). $^1H$ NMR ($D_2O$, 600 MHz): $\delta$ 4.09 (t, $J$ = 6 Hz, 1 H), 3.33 (t, $J$ = 8 Hz, 2 H), 2.11–1.82 (comp m, 4 H). MS calcd for $C_5H_{13}N_2O_3$ ($[M+H]^+$) 149.1, found 149.2.

*Cloning, Expression, and Purification*:

*S. griseus* subsp. *griseus* NRRL F5144 was used for colony PCR to amplify *pipS* using 5′-GGCAGCCATATGTTCGTTCCCAGTTACTACCGAGAG-3′ and 5′-GTGGTGCTCGAGGGATTCCGTCTGAGGCACTCG-3′ primers and subsequently cloned into pET22b using NdeI and XhoI restriction sites and confirmed by DNA sequencing. The protein was produced in *E. coli* BL21 (DE3) cells using autoinduction media (20 g/L tryptone, 10 g/L yeast extract, 50 mM $NH_4Cl$, 2 mM $MgSO_4$, 0.5% glycerol, 17 mM $KH_2PO_4$, 72 mM $K_2HPO_4$, 0.05% glucose, and 0.2% lactose) supplemented with 200 μg $ml^{-1}$ ampicillin and incubated for 1 h at 37 °C then 16 °C for 68-74 h. Cells were harvested by centrifugation, disrupted using sonication (5 rounds of 3 s/3 s on/off cycles for 2.5 min at 25% amplitude), and purified using nickel-nitrilotriacetic acid (Ni-NTA) resin (GE Healthcare) with binding buffer (20 mM Tris pH 8, 500 mM NaCl, 5 mM imidazole) and elution buffer (binding buffer plus 250 mM imidazole). Further purification was carried out using a HiLoad 16/600 Superdex 200pg size exclusion chromatography column (GE Healthcare) equilibrated in 20mM Tris pH 8, 50 mM NaCl. PipS-$His_6$ was then reconstituted with hemin chloride (Calbiochem) at a 2:1 molar ratio (hemin:protein) and incubated overnight at 4 °C. Excess heme was removed with a HiTrap Desalting Column (GE Healthcare) in 20 mM Tris pH 8.0, 50 mM NaCl buffer. Purified protein was then concentrated and used fresh for crystallization or flash frozen with the addition of 10% glycerol for further assays.

The *pipS* gene was also synthesized and codon optimized for *E. coli* by GenScript using their standard methods. The primers 5′-GGCAGCCATATGTTTGTGCCGAGCTATTATCGT-3′ and 5′-GTGGTGCTCGAGGCCTTGGAAGTAGAGGTTCTCGCTTTCGGTTTGCGGCAC-3′, with an engineered TEV cleavage site in the reverse primer, were used to amplify codon optimized *pipS*. *pipS* was cloned into pET22b using NdeI and XhoI restriction sites (construct tagless PipS) and confirmed by DNA sequencing. Expression and purification with Ni-NTA was the same as for PipS-$His_6$-tag (described above). After Ni-NTA purification, PipS was dialyzed into 20 mM Tris pH 8, 150 mM NaCl for 2 h. Dithiothreitol was added to 1.5 mM and TEV was added to 1:50 mg ratio (TEV:protein) and incubated at 4°C overnight to cleave off the $His_6$-tag. TEV-cleaved protein was further purified using a HiLoad 16/600 or 26/600 Superdex 200pg size exclusion column run in 20 mM Tris pH 8, 50 mM NaCl or sodium phosphate pH 8, 100 mM NaCl. PipS was then reconstituted with hemin chloride with a 1:1 molar ratio and incubated at 4°C overnight. Excess heme was removed using either a HiTrap Desalting Column or a PD-10 column (GE

Healthcare) in either 20 mM Tris pH 8, 50 mM NaCl or 20 mM Sodium phosphate pH 8, 100 mM NaCl. Protein was concentrated down and flash frozen with the addition of 10 % glycerol.

Protein concentration was determined by UV absorbance at 280 nM using calculated extinction coefficient 34,950 $M^{-1}$ $cm^{-1}$ [5]. Heme concentration was determined using the hemachromagen assay [6].

*UV-Vis Spectroscopy*:

All scans were carried out on a Cary 100 Bio UV-Visible Spectrophotometer (Agilent) and measured from 700 to 350 nm.

*In vitro biochemical analysis of PipS with* L-**1**:

Reactions were carried out using 2 mM $N^5$-OH-L-ornithine with 4 μM enzyme in 50 mM Tris pH 8.0 or 20 mM sodium phosphate, 100 mM NaCl pH 8.0 for 1 h at room temperature, unless otherwise stated. 9-fluorenylmethoxycarbonyl chloride (Fmoc-Cl) derivatization was performed as previously described [1]. Briefly, 100 μL of reaction mixture was quenched with 200 μl acetonitrile and incubated at −20°C for 5 min followed by centrifugation. 150 μl of the supernatant was removed and mixed with 10 μl borate buffer (0.2 M, pH 8) and 20 μl of Fmoc-Cl (20 mM in acetonitrile) for 5 minutes at room temperature. 20 μl 1-aminoadamantane (0.1 M) was then added for another 10 minutes before HPLC and/or LC-MS analysis.

For HPLC analysis, 20 μl of derivatized reaction mixture was loaded onto a 1260 HPLC apparatus (Agilent) using a Phenomenex Luna C18(2) column (5 μm, 4.6 mm ID x 250 mm). Elution was performed at 1 ml $min^{-1}$ with a mobile-phase mixture consisting of a linear gradient of water and acetonitrile both of which contains 0.05 % (v/v) trifluoroacetic acid ((v/v): 35:60 to 20:80, 0 to 20 min; 20:80 to 0:100, 20 to 25 min; 0:100, 25 to 27 min) with a detection wavelength of 280 and 254 nm.

*Absolute configuration of Piz*

The absolute configuration of Piz was determined as previously described [1]. Briefly, a 100 μL reaction mixture consisting of 2 μM enzyme and 1 μM L/D-**1** in 50 mM MOPS buffer pH 8.0 was incubated for 1 h. The reaction mixture or L/D-**2** standard was derivatized with 1-fluoro-2-4-dinitrophenyl-5-L-alanine amide (L-FDAA) and subjected to LC-MS analysis. L-FDAA derivatization was performed as follows: 50 μl of samples were treated with 20 μl of 1 M $NaHCO_3$ and 100 μl of 1% L-FDAA dissolved in acetone at 37 °C for 60 min. The mixtures were quenched by the addition of 20 μl of 1 N HCl and diluted with 810 μl of methanol before centrifugation and LC-MS analysis. For LC-MS analysis, 20 μl of derivatized reaction mixture was loaded onto 6120 Quadrupole LC-MS system (Agilent) using a Phenomenex Luna C18(2) column (5 μm, 4.6 mm ID x 250 mm), and operated in selected ion monitoring (SIM) mode with a *m/z* value of 383 monitored, corresponding to FDAA-**2**. Elution was performed at 0.5 ml $min^{-1}$ with a mobile-phase mixture consisting of a linear gradient of water and acetonitrile, both containing 0.1% (v/v) formic acid: 70:30, 0 to 3 min; 70:30 to 30:70, 3 to 18 min; 0:100, 18 to 22 min.

*Crystallization, Data Collection, and Structure Solving*:

PipS-$His_6$, reconstituted with heme, was initially screened using Hampton Crystal Screens 1 and 2 (Hampton Research) and MCSG Crystallization Suite (Anatrace) using sitting drop vapor diffusion at room temperature in the dark. After optimization, the best crystals were grown in 0.74-

0.78 M sodium citrate tribasic dihydrate, 0.1 M sodium cacodylate pH 6.5 using hanging drop vapor diffusion at room temperature in the dark.

Holo crystals were cryoprotected in mother liquor supplemented with 20 % ethylene glycol and flash frozen in liquid nitrogen. To obtain a complex structure, holo crystals were soaked with 500 mM **1** for 1 h then cryoprotected in mother liquor supplemented with 20 % ethylene glycol. Holo PipS-His$_6$ data was collected at the Canadian Light Source (CLS) at beamline 08ID-1, processed using iMOSFLM [7], and scaled with Aimless [8]. Complex structure data was collected at the Stanford Synchrotron Radiation Lightsource (SSRL) beamline 9-2 and processed and scaled using XDS [9].

The holo structure was solved with Phaser-MR in Phenix [10] using the Pai 2 structure (PDB 2OL5) [11] as the starting model followed by AutoBuild [12] to give the initial PipS-His$_6$ model. Several rounds of manual building in COOT [13,14] with refinement using phenix.refine [15], with water molecules added using the FindWaters function in Coot [13,14], led to the finished PipS-His$_6$-tag holo structure. The substrate-bound PipS-His$_6$-tag structure was solved using Phaser-MR with the PipS-His$_6$-tag holo structure as the starting model. Several rounds of manual building with refinement were performed similar to the holo-PipS-His$_6$ structure. Refinement procedures were monitored by flagging 5% of all observations as 'free' [16] and model validation was performed by SFCHECK [17], RAMPAGE [18], and MOLPROBITY [19]. A list of data collection and refinement statistics can be found in **Supplementary Table 2**.

*PipS mutagenesis*:

Quickchange mutagenesis [20] was used to generate single amino acid substitutions in the PipS construct. Primer sequences are listed in **Supplementary Table 3**. Expression and purification were carried out as described above for PipS. For the reaction, 2 mM L-**1** was incubated with 1 μM enzyme in 20 mM sodium phosphate pH 8, 100 mM NaCl for 1.5 h at 30 °C. Reactions were quenched with 2 volumes of acetonitrile, followed by Fmoc-Cl derivatization before HPLC analysis. Peak areas for **2** were integrated for quantification to determine the percent activity when compared to wild-type enzyme. Each reaction was performed in triplicate.

*In vitro biochemical assay of as-isolated PipS with unnatural substrates*

For the reaction of as-isolated PipS with unnatural substrates, the reaction mixtures containing 2 μM enzyme (PipS or its variants) and 2 mM substrates (**4**, **5**, **6**, **7**, or **8**) in 25 mM HEPES buffer (pH 8.0) were incubated at room temperature for 1 h and then quenched by 2 volumes of acetonitrile. HPLC analysis of the reaction mixtures after Fmoc derivatization were conducted as described above. For HPLC analysis of assay mixtures of PipS (or its variants) and **8** without Fmoc derivatization, 20 μl of quenched reaction mixture was loaded onto a 1260 HPLC apparatus (Agilent) using a Phenomenex Luna C18(2) column (5 μm, 4.6 mm ID x 250 mm). Elution was performed at 1 ml min$^{-1}$ with a mobile-phase mixture consisting of a linear gradient of water and acetonitrile both of which contains 0.05% (v/v) trifluoroacetic acid ((v/v): 80:20 to 0:100, 0 to 25 min; 0:100, 25 to 30 min) with a detection wavelength of 254 nm. Formaldehyde production was detected by a Formaldehyde Assay Kit from Sigma-Aldrich.

*EPR analysis of PipS*

Electron paramagnetic resonance (EPR) spectra were recorded on a Bruker X-band EMX spectrometer equipped with an Oxford Instruments 3 S3 liquid helium cryostat. EPR spectra were obtained on frozen solution samples with a glycerol content of 30% in buffer at temperatures of

10 K or 25 K. Holo-PipS and PipS T107A-K178A samples had a concentration of 1472 μM. Isopropylhydroxylamine (**12**) and L-**1** containing samples were prepared by addition of 12 μL of 500 mM stock to 300 μL of holo-PipS and PipS T107A-K178A samples. Isopropylhydroxylamine (**12**) or L-**1** added samples were incubated overnight at 4 °C before freezing. PipS+$His_6$-tag and PipS-$His_6$-tag samples were prepared at a concentration of 360 μM and 50 mM imidazole. The parameters used were 20 mW microwave power and 100kHz field modulation with the amplitude set to 3G. The g-values and signal integrals were obtained using the program SpinCount (by Prof. M. P. Hendrich, Carnegie Mellon University).

*MCD analysis of PipS*

All MCD samples were prepared in 18 mM phosphate, 43 mM NaCl, 50 % glycerol (added as a glassing agent), pH 8.0 buffer. The proteins of interest were injected between two quartz plate windows housed in a custom-made MCD sample holder. The samples were frozen in liquid nitrogen to produce a glass.

An OXFORD SM4000 cryostat and a JASCO J-815-CD spectrometer were used for the MCD setup. The SM4000 cryostat, consisting of a liquid helium-cooled superconducting magnet, provides horizontal magnetic fields of 0-7 T. The J-815 spectrometer uses a gaseous nitrogen cooled xenon lamp and a detector system consisting of two interchangeable photomultiplier tubes in the UV-vis and NIR range. The samples were loaded into a 1.5 - 300 K variable temperature insert (VTI), which offers optical access to the sample via four optical windows made from Suprasil B quartz. The MCD spectra were measured in [θ] = mdeg and manually converted to Δε ($M^{-1}$ $cm^{-1}$ $T^{-1}$) using the conversion factor

$$\Delta\varepsilon = \frac{\theta}{32980 \bullet c \bullet d \bullet B}$$

where c is the concentration, B is the magnetic field, and d is the path length. The product c•d can be substituted by A(MCD)/(UV-vis), where A(MCD) is the absorbance of the sample measured by the CD spectrometer and (UV-vis) is the molar extinction coefficient.[21] Complete spectra were recorded at indicated temperatures and magnetic fields.

*rRaman analysis of PipS*

Samples prepared for EPR were also used directly for the rRaman measurements. The rRaman experiments were performed using an INNOVA-301K* Krypton Ion laser system at an excitation wavelength of 407 nm with 20 mW power. Samples were transferred to a liquid nitrogen bath inside an EPR cold finger to cool the samples during the measurements. The scattered light from the samples was focused onto an Acton two-stage TriVista 555 monochromator and detected by a liquid N2-cooled Princeton Instruments Spec-10:400B/LN CCD camera. The accumulation and exposure times for the samples were 10 acquisitions for 60 seconds. The spectral resolution was 0.3 $cm^{-1}$. Spectra were then plotted and processed using the OriginPro 9.0.0 (64-bit) software for baseline correction

*Monitoring oxygen concentration in aerobic PipS WT and T107A-K178A reactions.*

The oxygen concentration in aerobic reactions with 400 μM PipS (WT or T107A-K178A) and 15 mM L-**1** or *N*-isopropylhydroxylamine (**12**) in 20 mM sodium phosphate buffer pH 8.0 were monitored using a Clark-type electrode coupled to an OXYG1-Oxygraph electrode control unit (Hansatech). The electrode was initially calibrated using air-saturated 20 mM sodium phosphate buffer pH 8.0 at 25 °C. 1 mL of buffer solution and enzyme were equilibrated in the reaction chamber before adding substrate to start the reaction. 15 μL of the reaction was removed every 10 min and diluted to a final enzyme concentration of 3 μM for UV-Vis analysis.

**Supplementary Table 1**. *S. griseus* subsp. *griseus* NRRL F-5144 polyoxypeptin-like biosynthetic gene cluster.

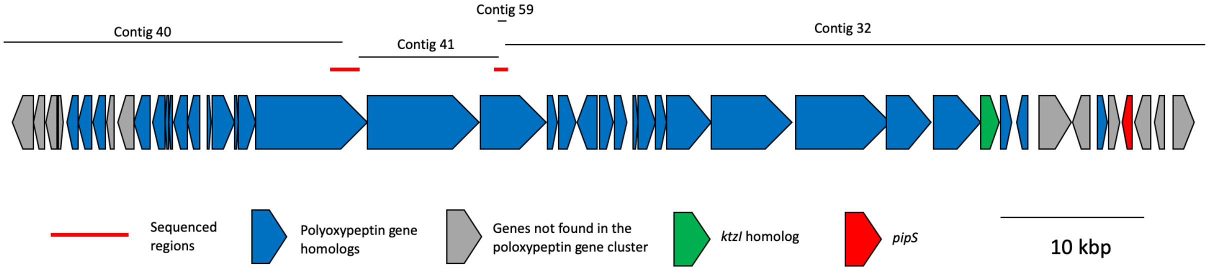

| Contig | Start | Stop | Size | Accession number | Proposed function | Homolog in *ply* cluster | % ID[a] |
|---|---|---|---|---|---|---|---|
| 40 | 2041 | 584 | 578 | WP_030723148.1 | Serine hydrolase | *Not found* | |
| 40 | 2909 | 2085 | 274 | N/A | Transcriptional regulator | *Not found* | |
| 40 | 3709 | 2975 | 260 | WP_030723150.1 | Transcriptional regulator | *Not found* | |
| 40 | 3764 | 4120 | 118 | N/A | Hypothetical protein | *Not found* | |
| 40 | 5173 | 4382 | 263 | WP_030723152.1 | ABC transporter | PlyK | 42 |
| 40 | 6171 | 5170 | 333 | WP_030723153.1 | ABC transporter | PlyJ | 63 |
| 40 | 7075 | 6233 | 280 | WP_030723154.1 | DUF4097 family protein | Orf11 | 41 |
| 40 | 7673 | 7158 | 171 | WP_030723156.1 | Antitoxin | *Not found* | |
| 40 | 9067 | 7910 | 385 | WP_030723159.1 | Acetyl-CoA dehydrogenase | *Not found* | |
| 40 | 10261 | 9128 | 377 | WP_050513873.1 | Zinc-binding dehydrogenase | Orf5 | 86 |
| 40 | 11282 | 10344 | 312 | WP_030723166.1 | Ketoacyl-ACP-synthase III | Orf6 | 84 |
| 40 | 11527 | 11279 | 82 | WP_030723169.1 | Hypothetical protein | Orf7 | 91 |
| 40 | 11805 | 11524 | 93 | WP_050513874.1 | Dihydrolipoamide succinyltransferase | Orf8 | 72 |
| 40 | 12794 | 11823 | 323 | WP_030723175.1 | Alpha-ketoacid hydrogenase | Orf9 | 87 |
| 40 | 13642 | 12803 | 279 | WP_032769773.1 | Pyruvate dehydrogenase | Orf10 | 85 |
| 40 | 14194 | 14409 | 71 | WP_030723182.1 | MbtH-like | PlyA | 83 |
| 40 | 14485 | 16047 | 520 | WP_030723184 | Amino acid adenylation domain | PlyC | 79 |
| 40 | 16044 | 16277 | 76 | WP_032769775.1 | Acyl carrier protein | PlyD | 77 |
| 40 | 16338 | 17525 | 395 | WP_030723189.1 | NAD(P) / FAD-dependent oxidoreductase | PlyE | 82 |
| 40 /seq/ 41[b] | 17570 | 25351 | 2593 | WP_030723191.1[c] | Non-ribosomal peptide synthetase | PlyF | 80 |
| 41 | 25348 | 33114 | 2588 | WP_030723197.1 | Non-ribosomal peptide synthetase | PlyG | 78 |
| 41/59/ 32 | 33150 | 37784 | 1544 | N/A | Non-ribosomal peptide synthetase | PlyH | 74 |
| 32 | 37844 | 38596 | 250 | WP_032769648.1 | Thioesterase | PlyI | 82 |

| | | | | | | | |
|---|---|---|---|---|---|---|---|
| 32 | 38610 | 39842 | 410 | WP_030722355.1 | Cytochrome P450 | PlyM | 83 |
| 32 | 41297 | 39921 | 440 | WP_030722358.1 | PLP-dependent aminotransferase | PlyN | 81 |
| 32 | 41546 | 42472 | 308 | WP_030722361.1 | Phytanoyl-CoA dioxygenase | PlyO | 87 |
| 32 | 42536 | 43402 | 288 | WP_078614552.1 | Proline hydroxylase | PlyP | 80 |
| 32 | 43848 | 44159 | 103 | WP_030722367.1 | Thiolation domain | PlyQ | 74 |
| 32 | 44156 | 45406 | 416 | WP_030722370.1 | Cytochrome P450 | PlyR | 86 |
| 32 | 45403 | 46140 | 245 | WP_030722373.1 | Thioesterase | PlyS | 80 |
| 32 | 46192 | 49269 | 1025 | WP_078614553.1 | Modular polyketide synthase | PlyT | 78 |
| 32 | 49308 | 54908 | 1866 | WP_032769651.1 | Modular polyketide synthase | PlyU | 75 |
| 32 | 54905 | 61483 | 2192 | WP_030722382.1 | Modular polyketide synthase | PlyV | 78 |
| 32 | 61485 | 64601 | 1038 | WP_030722384.1 | Modular polyketide synthase | PlyW | 80 |
| 32 | 64779 | 68012 | 1077 | WP_078614554.1 | NRPS | PlyX | 74 |
| 32 | 68079 | 69386 | 435 | WP_030722388.1 | Lysine oxygenase | *Not found.* | |
| 32 | 69441 | 70181 | 246 | WP_030722390.1 | TE | PlyY | 77 |
| 32 | 71377 | 70532 | 281 | WP_107047173.1 | Hypothetical | Orf11 | 35 |
| 32 | 72130 | 74373 | 747 | WP_107047174.1 | Transporter | *Not found* | |
| 32 | 75611 | 74445 | 388 | WP_050513858.1 | Regulatory protein | *Not found.* | |
| 32 | 76140 | 76826 | 228 | WP_030722402.1 | Antibiotic biosynthesis protein | *PlyB* | 40 |
| 32 | 76974 | 77759 | 261 | WP_078614555.1 | Amidinotransferase | *Not found* | |
| 32 | 78539 | 77883 | 218 | WP_030722408.1 | Piperazate synthase | *Not found* | |
| 32 | 79914 | 78745 | 389 | WP_030722411 | Alkaline D-peptidase | *Not found* | |
| 32 | 80882 | 80136 | 248 | WP_063837377 | 4-phosphopantetheinyl transferase | *Not found* | |
| 32 | 81417 | 82919 | 500 | WP_030722417.1 | Polyketide oxygenase/ hydroxylase | Orf4 | 76 |

[a]Amino acid identity
[b]Missing sequence was determined via sequencing
[c]Truncated sequence

**Supplementary Table 2**. Data collection and refinement statistics.

| | PipS-His$_6$-tag | PipS-L-**1** |
|---|---|---|
| **Data collection** | | |
| Beamline | CMCF-08ID-1 | SSRL-9-2 |
| Wavelength | 0.97949 | 0.97936 |
| Space group | P 61 2 2 | P 61 2 2 |
| Cell dimensions | | |
| *a*, *b*, *c* (Å) | 66.51, 66.51, 478.69 | 67.00, 67.00, 479.00 |
| Resolution (Å) | 79.78 - 2.08 (2.14 – 2.08)[a] | 79.83 – 1.92 (1.97 – 1.92) |
| $R_{pim}$ (within I+/I-)[b] | 0.044 (0.476) | 0.045 (0.506) |
| $I$ / σ$I$ | 15.3 (2.3) | 13.3 (2.0) |
| $CC_{1/2}$[c] | 0.999 (0.882) | 0.998 (0.778) |
| Completeness (%) | 100 (100) | 100 (100) |
| No. unique reflections | 39715 (2987) | 50850 (3343) |
| Redundancy | 33.7 (32.4) | 27.3 (28.0) |
| **Refinement** | | |
| Resolution (Å) | 2.08 | 1.92 |
| $R_{work}$[d] / $R_{free}$[e] | 0.1885/0.2269 | 0.1740/0.2116 |
| No. atoms | | |
| Protein | 3379 | 3337 |
| Solvent | 173 | 260 |
| Heme | 86 | 86 |
| L-**1** | N/A | 20 |
| *B*-factors | | |
| Protein | 49.5 | 41.6 |
| Solvent | 46.9 | 45.8 |
| Heme | 35.6 | 30.4 |
| L-**1** | N/A | 52.2 |
| R.m.s. deviations | | |
| Bond lengths (Å) | 0.009 | 0.007 |
| Bond angles (°) | 1.155 | 0.893 |
| Ramachandran | | |
| Preferred (%) | 97.0 | 97.9 |
| Generously allowed (%) | 3.0 | 2.1 |
| Disallowed (%) | 0.0 | 0.0 |
| TLS groups | 13 | 13 |
| PDB | To be determined | To be determined |

[a]Data from the highest-resolution shell are indicated in parentheses.

[b]$R_{pim} = \frac{\sum_{hki} (\frac{1}{n-1})^{1/2} \sum_{i=1}^{n} |I_i(hkl) - I(hkl)|}{\sum_{hki} \sum_{i=1}^{n} I_i(hkl)}$

[c]$CC_{1/2} = cov \frac{(X,Y)}{\sigma_x \sigma_y} = E \frac{[X-\mu_x][Y-\mu_y]}{\sigma_x \sigma_y}$

[d]$R_{work} = \frac{(\sum_{hkl} ||F_o(hkl)| - |F_c(hkl)||)}{\sum_{hkl} |F_o(hkl)|}$

[e]$R_{free}$ is calculated identically to $R_{work}$, using 5% of reflections omitted from refinement.

**Supplementary Table 3.** Primer sequences for PipS mutagenesis.

| Mutation | Primer sequences (5′ → 3′) |
|---|---|
| Y7F | F = TGTGCCGAGCTATTTCCGTGAGCCGCATGGTAGCT |
| | R = AGCTACCATGCGGCTCACGGAAATAGCTCGGCACA |
| H66A | F = TGCGAACCTGCTGGGCGCAATGAACCGTGCGAACC |
| | R = GGTTCGCACGGTTCATTGCGCCCAGCAGGTTCGCA |
| Y99F | F = ACGTGAGCCCGGCGCTGTTTGGTGTTACCCCGGCG |
| | R = CGCCGGGGTAACACCAAACAGCGCCGGGCTCACGT |
| T107A | F = TTACCCCGGCGGCGCCGGCATGGAACTTTACCAGCGT |
| | R = ACGCTGGTAAAGTTCCATGCCGGCGCCGCCGGGGTAA |
| T107V | F = TTACCCCGGCGGCGCCGGTGTGGAACTTTACCAGCGT |
| | R = ACGCTGGTAAAGTTCCACACCGGCGCCGCCGGGGTAA |
| T107S | F = TTACCCCGGCGGCGCCGAGCTGGAACTTTACCAGCGT |
| | R = ACGCTGGTAAAGTTCCGCTCCGGCGCCGCCGGGGTAA |
| T107C | F = TTACCCCGGCGGCGCCGTGCTGGAACTTTACCAGCGT |
| | R = ACGCTGGTAAAGTTCCGCACCGGCGCCGCCGGGGTAA |
| Y155F | F = GAGCGATAGCATCGACTTCTTCCGTAAAATTGTG |
| | R = CACAATTTTACGGAAGAAGTCGATGCTATCGCTC |
| K178A | F = AGCGCGCACGGCATGTTCGCACTGAGCCAGGAGCA |
| | R = TGCTCCTGGCTCAGTGCGAACATGCCGTGCGCGCT |
| K178M | F = AGCGCGCACGGCATGTTCATGCTGAGCCAGGAGCA |
| | R = TGCTCCTGGCTCAGCATGAACATGCCGTGCGCGCT |
| E182A | F = TTCAAGCTGAGCCAGGCACAACCGGCGGAA |
| | R = TTCCGCCGGTTGTGCCTGGCTCAGCTTGAA |

**Supplementary Figure 1**. **Biosynthesis of selected N-N-bond-containing natural products**. **a**, Piperazic acid, a residue in found diverse non-ribosomal peptides like kutzneride, is biosynthesized through a two-step pathway from L-ornithine[1,22]. **b**, The ω-nitrogens of L-arginine from a single molecule are incorporated together into streptozocin,[23,24] with the streptozotocin biosynthetic enzyme SznF catalyzing an $O_2$-dependent N-nitrosation.[23] **c**, Valanimycin is assembled from L-serine and N-OH-isobutylamine[25,26]. **d**, GTP and arginine are the biosynthetic precursors for 8-azaguanine, with arginine-derived nitric oxide as the biosynthetic source of the central nitrogen in the triazole[27]. **e**, Glycine and N-OH-L-lysine are converted to hydrazino acetic acid by the action of Spb40 and Spb39 in the assembly of s56-p1[28]. **f**, In cremeomycin biosynthesis, CreE and CreD form nitrous acid from aspartate,[29] and CreM is proposed to catalyze cremeomycin assembly from 3-amino-2-hydroxy-4-methoxybenzoic acid and nitrous acid[30]. **g**, PyrN couples glutamate and N-OH-l-lysine to give a linear hydrazine on-pathway to pyrazomycin[31]. **h**, The hydrazine in fosfazinomycin is derived from glutamylhydrazine[32].

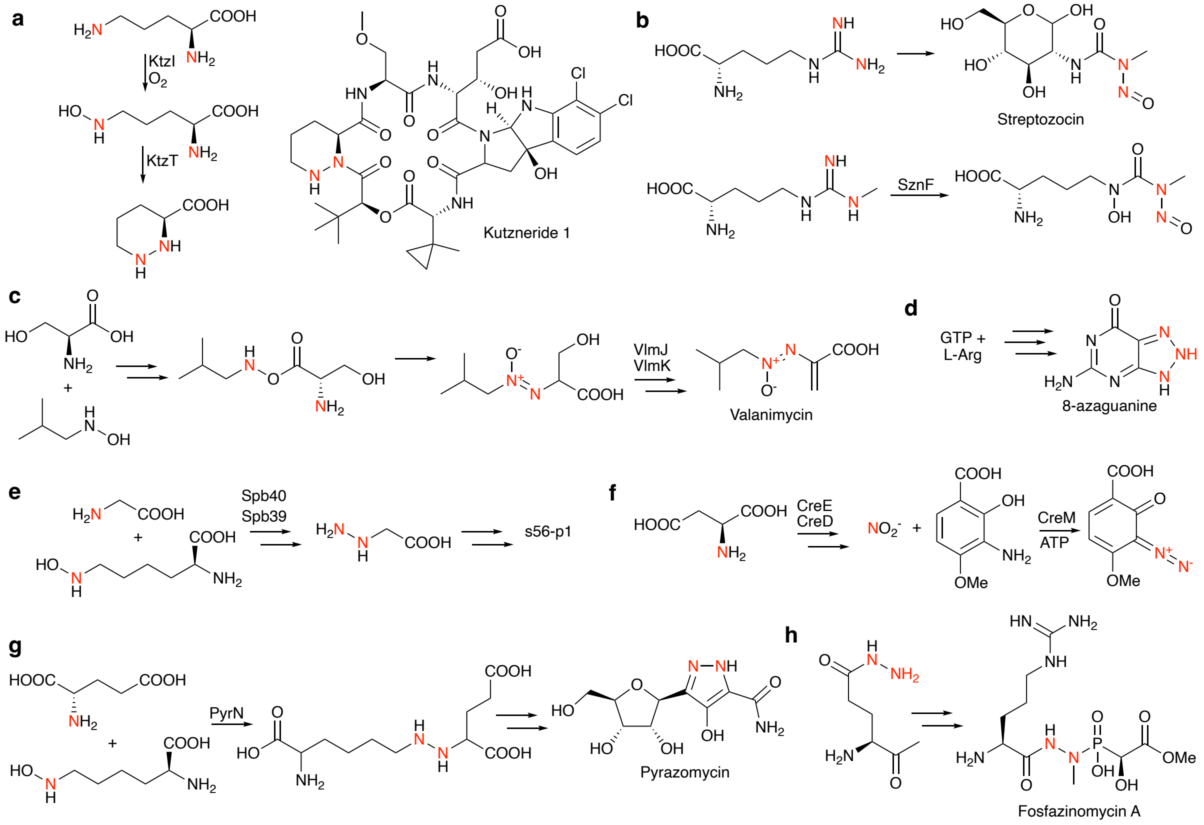

**Supplementary Figure 2**. L-**1** synthesis. Diisopropylethylamine (DIPEA) is used in to partially purify the nitrone by precipitation [1,3]. **a**, Scheme synthesis of L-**1**. **b**, NMR spectra of L-**1**. Peaks due to an DIPEA impurity are labelled with stars. Purity was assessed on an NMR integration basis. 12 protons in the DIPEA impurity correspond to an integration value of 0.36 in the spectrum; one proton therefore integrates for 0.03, giving an approximate 3% DIPEA impurity. **c**, Reactions with 1 mM L-**1**, 300 nM PipS, and increasing concentrations of DIPEA for 10 min at room temperature. Reactions were derivatized with Fmoc-Cl and analyzed via HPLC.

**a**

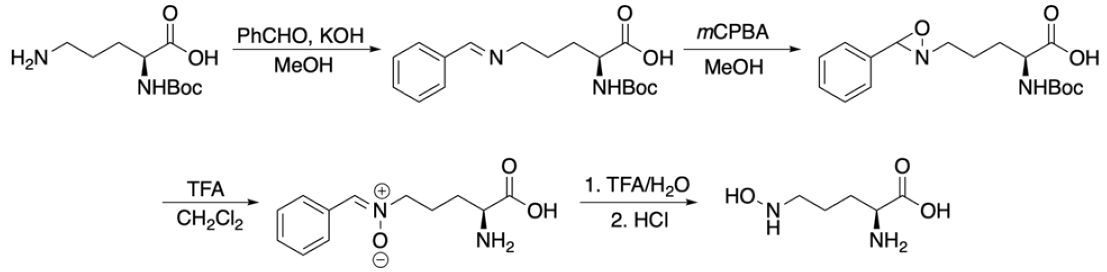

**b**

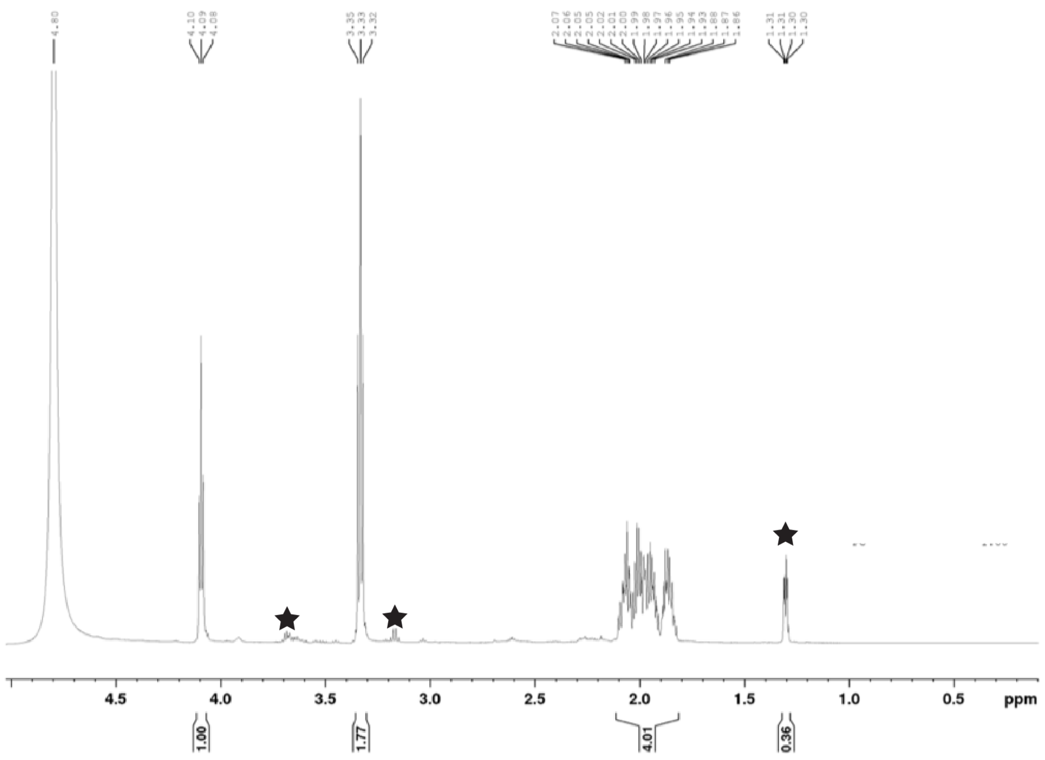

c

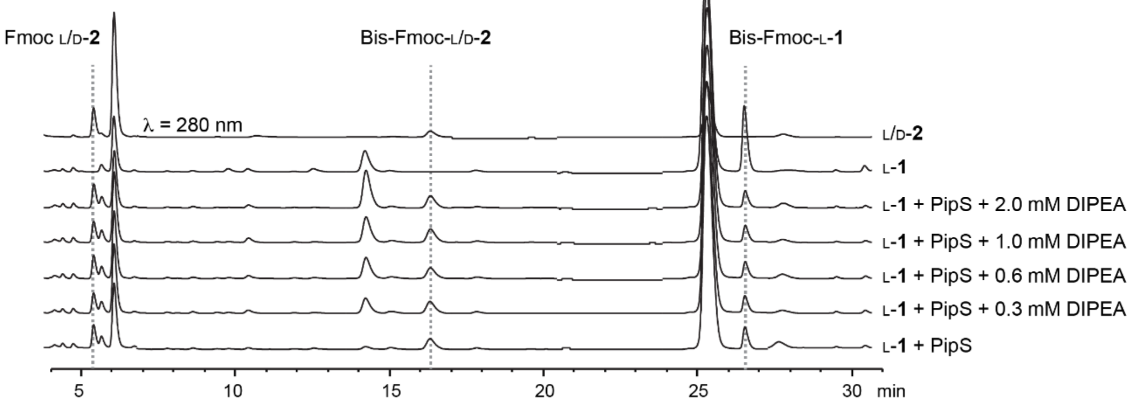

Fmoc L/D-2
Bis-Fmoc-L/D-2
Bis-Fmoc-L-1
λ = 280 nm
L/D-2
L-1
L-1 + PipS + 2.0 mM DIPEA
L-1 + PipS + 1.0 mM DIPEA
L-1 + PipS + 0.6 mM DIPEA
L-1 + PipS + 0.3 mM DIPEA
L-1 + PipS
5
10
15
20
25
30
min

**Supplementary Fig. 3:** Structural comparison of PipS with PaiB (PDB 2OL5) [11] and Anf3 (PDB 6RK0) [33]. Overlay of a cartoon representation of PipS (grey) and PaiB/Anf3 (purple) with the heme from the PipS structure shown in yellow and FAD and heme from Anf3 shown in orange. The inset shows the active site architecture of PipS and PaiB/Anf3. PipS residues are labeled in black while PaiB/Anf3 residues are labeled in red and the N-terminus of PipS is shown as a green line. Oxygen is red, nitrogen is blue, and iron is orange.

To date, the only structural homolog of PipS is PaiB [11]. PipS has the same overall fold to PaiB (RMSD = 1.8 Å over 183 residues [34]). Interestingly, PaiB cannot catalyze the piperazate synthase reaction [1], even though it has a similar active site architecture to PipS and contains the conserved catalytic Thr/Lys dyad. Additionally, both the heme binding site and the His are also conserved with the oxidase Anf3 [33] (RMSD = 3.2 Å over 163 residues [34]) which is part of a larger split β-barrel enzyme superfamily [35]. However, PipS lacks the flavin binding site often found in this enzyme superfamily because the heme propionate and side-chains from the N-terminus of PipS physically block the flavin binding site. The lack of a flavin binding site, together with the observations that no NAD(P)H or flavin is needed for the PipS reaction provides further evidence that PipS undergoes a reaction mechanism different from an oxidase.

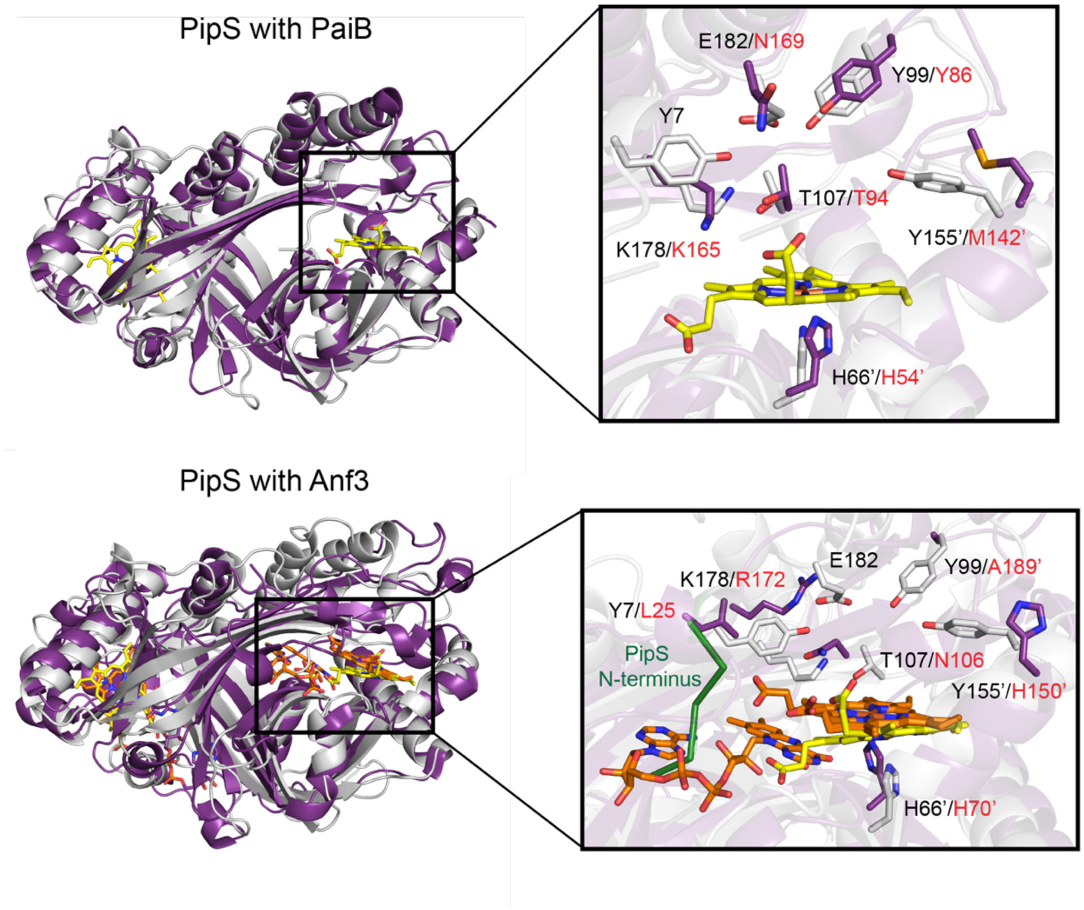

**Supplementary Figure 4**: Spectroscopic analysis of oxidized and reduced tagless PipS. **a**, UV-Vis spectrum of 5μM PipS with and without 10 mM dithionite. **b**, rRaman (407nm wavelength excitation, 20mW power) spectrum of PipS. **c**, rRaman spectrum of PipS with excess dithionite reductant. These data support the presence of a ferric heme, with the ferric marker band observed at 1374 $cm^{-1}$, which shifts to 1357 $cm^{-1}$ upon reduction of the heme with dithionite, corresponding to the ferrous marker band.[36–38] **d**, MCD spectrum of PipS at 2K; magnetic field 1T. These data support a typical HS ferric signal.[39]

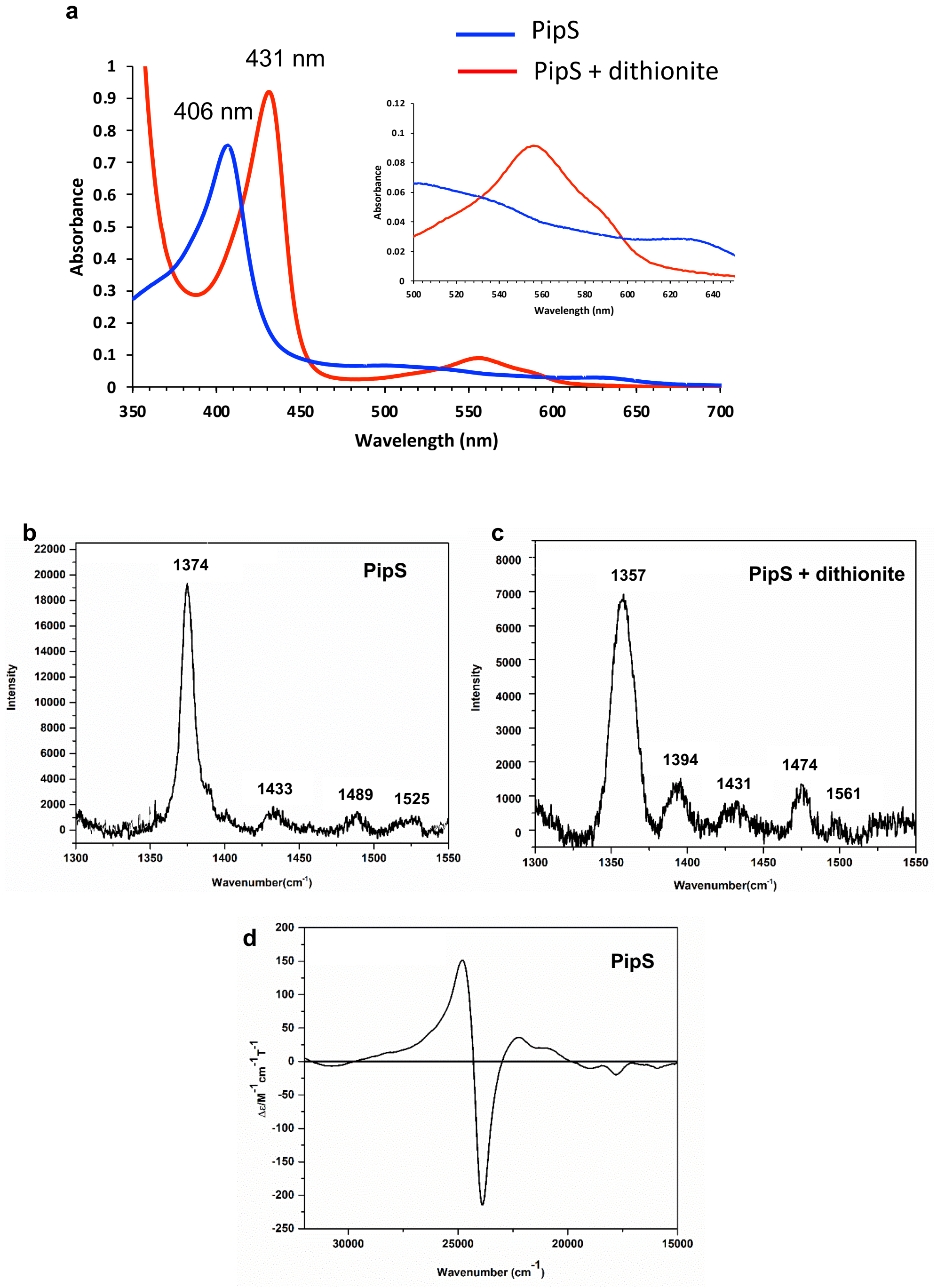

**Supplementary Figure 5**. Amino acid sequence alignment of PipS with homologous enzymes KtzT from *Kutzneria sp*. 744 (WP_043726237.1), CfPipS from *Collimonas fungivorans* (WP_061538620.1), KsPipS from *Kibdelosporangium sp*. MJ126-NF4 (WP_042177946.1), PsPipS from *Pseudonocardia* (WP_060714104.1), and LfPipS from *Lentzea flaviverrucosa* (WP_090066830.1). Residues highlighted in blue are located in the active site while residues highlighted in red are critical for catalysis. Activity assays for these homologs can be found in **Supplementary Fig. 6**.

```
PipS        MFVPSYYREPHGSWMAELIRGNPLAMAVINGST-DDGPFATHLPIIPDPRTTGE--WPDD       57
KtzT        MFVPGPYHAPEDRWLVDLVRGHPLAQLASNGAG-GAAPHITHVPIIVDPELDGP---VDR       56
CfPipS      MYVPEYYRVDE-NTARELVYRHPLALLVCNGN--NGLPWATHLPAIFPPETRKLLDQGES       57
KsPipS      MHVPPMYEAPDPAWIPALIRAHPLATLVTAP---DGIPAASHVPMIIRRTPDD-------       50
PsPipS      MFVPEQYREQDSNWMLDIVRSNPLALMASDGTPEGCGPAATHLPCIPDPSAPHD--WSDG       58
LfPipS      MFVPAQYREPHGHWITDLVRGHPLAQLVSNGPA-GSSPYVTHAPIILDPGHPDP--HPDD       57
            *.**  *.  .      ::  :***  .      .  *  :* * *

PipS        LTGANLLGHMNRANPQWQELETGKVILLAFTGPHAYVSPALYGVTPAAPTWNFTSVHVRG       117
KtzT        LVGITLWGHMNRANPHWAALGGAANVVATFAGPNAYVSPAVYRTAPAAPTWNFTSVQVRG       116
CfPipS      IIGKTMYGHMNRINPHWNALQAGS-ALLIFQGPNSYVSPTVYEVTPAAPTWNFTSTHLRG       116
KsPipS      -ERLTLVGHMNRMNPQFKAIGDGCPALLVFTGPHGYVSPTVYGFTPAAPTWNFAVVHASG       109
PsPipS      PRGAVLLGHMNRANPQWRHLHDGQTVLLVFTGPHAYVSPAVYDTTPAAPTWDFTAVHVHG       118
LfPipS      LHGAVLWGHLNRANPHWAALGDGTEVTAVFTGPGSYVSPTVYERTPAAPTWDFTAVHVRG       117
                 : **:** **::  :  .      * ** .****::*  :*****:*: .:   *

PipS        VVEKIESL---EETLDVVRATAGSFEARFGDDWDPSDSIDYFRKIVPGVGAFRVTVTSAH       174
KtzT        ELRKVESA---DDTLATVRATVAALESRFGAGWDMTGSLDYFRRILPGVGAFRLRVAEAD       173
CfPipS      TLRPIDER---DQILEIVRWTVATFEKEFCTNWDLTESIPYFERIVHGVGAFAFEVESFD       173
KsPipS      TLSPLPAG---PDTLEVIIDTVTALEGQLGNGWQMRDSLEYFDQLLPGVGAFSVQVDRVE       166
PsPipS      VVTKLEPHKAERTTLDVVTDTVTALEGRFGAGWDMTDSIEYFHRLLPGVGAFRVRVGSAE       178
LfPipS      TLRRVLDA---EQTLATVTATVRAFEADHGTGWSMESSLDYFDQLLPGVGAFRLAVTGVD       174
             :  :         *  :  *. ::*     .*.   *: ** ::: ***** . *   .

PipS        GMFKLSQEQPAEVRDRVQKSFSGRGCSRHRETAELMGRVPQTES--------       218
KtzT        GMFKLSQEQQPAIRRRVRHSFGGCEATR--AVAGLMDRLPTE----------       213
CfPipS      SMFKLSQEQPAAIQERVVNSFASSSHCPHKEIADLMQRTNSKNKK-------       218
KsPipS      AMYKLSQEQEPTTRETVAAAFEARSS----DLAAMMRVCLDVERSTLGNRVG       214
PsPipS      GMFKLSQEQPSDIRDRVRCHFAAAQHGRSSEIAHLMTTLDGH----------       220
LfPipS      AMFKLSQEQPPEVRLRVRDHFAGSERTHHCLIAEMMDRLPVAEH--------       218
            .*:******    :  *   * .         * :*
```

**Supplementary Figure 6**. Sequence logos of the PipS active site residues and surrounding regions. The top 250 hits from a Blast search using the PipS sequence were aligned using COBALT.[40] The multiple sequence alignment was then uploaded to WebLogo[41] to generate the sequence logo.

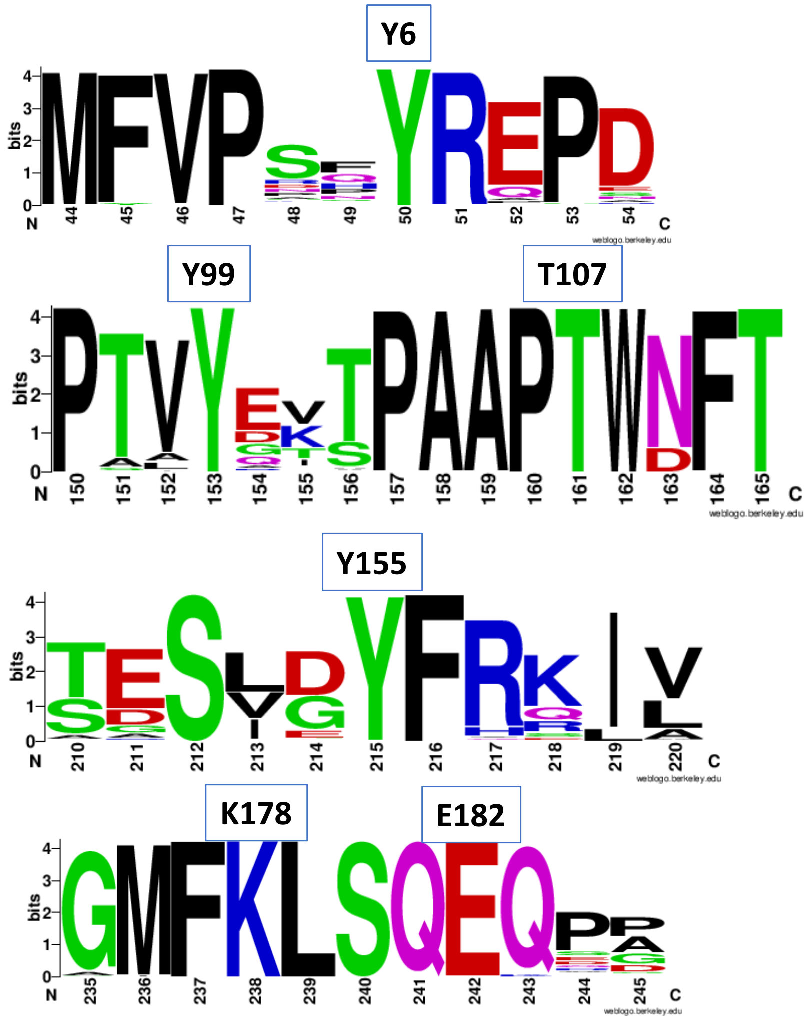

**Supplementary Figure 7**. SDS-PAGE (top) and size exclusion chromatography profiles (lower) for PipS WT and variants. The oligomeric state of PipS is consistent with KtzT.

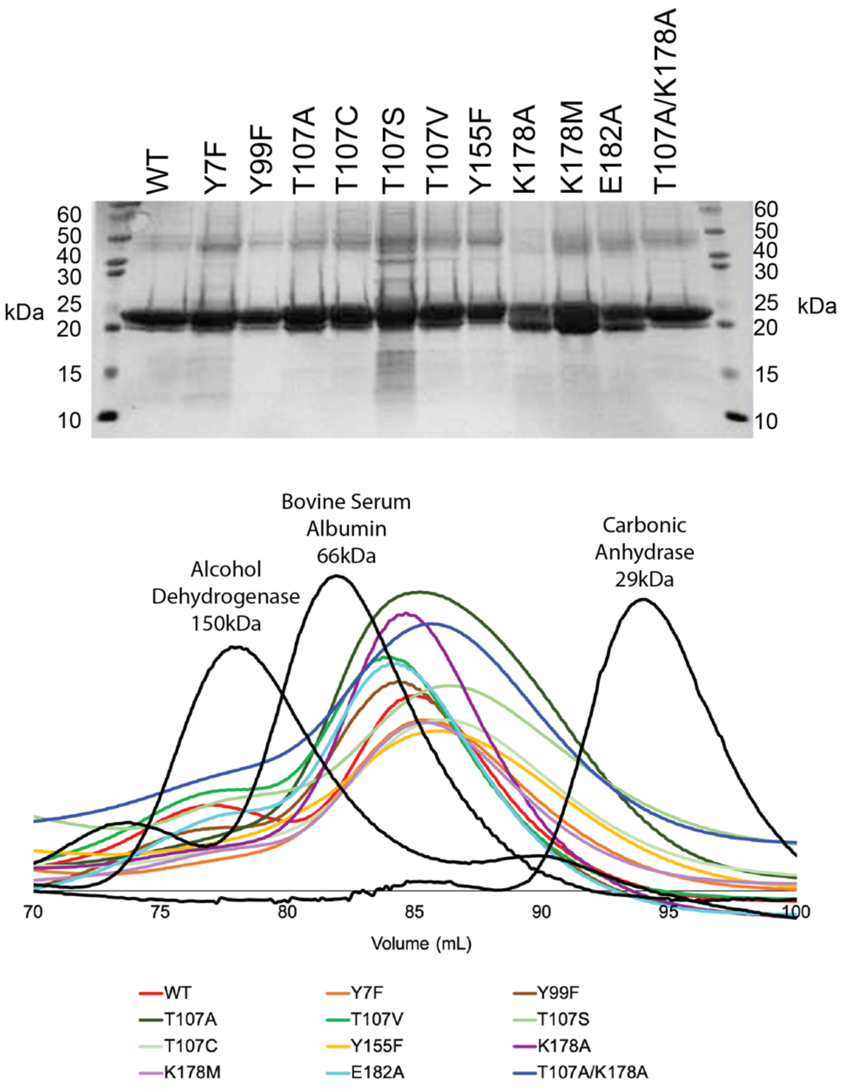

**Supplementary Figure 8**. UV-Vis spectra for PipS wild-type and variant purified enzymes.

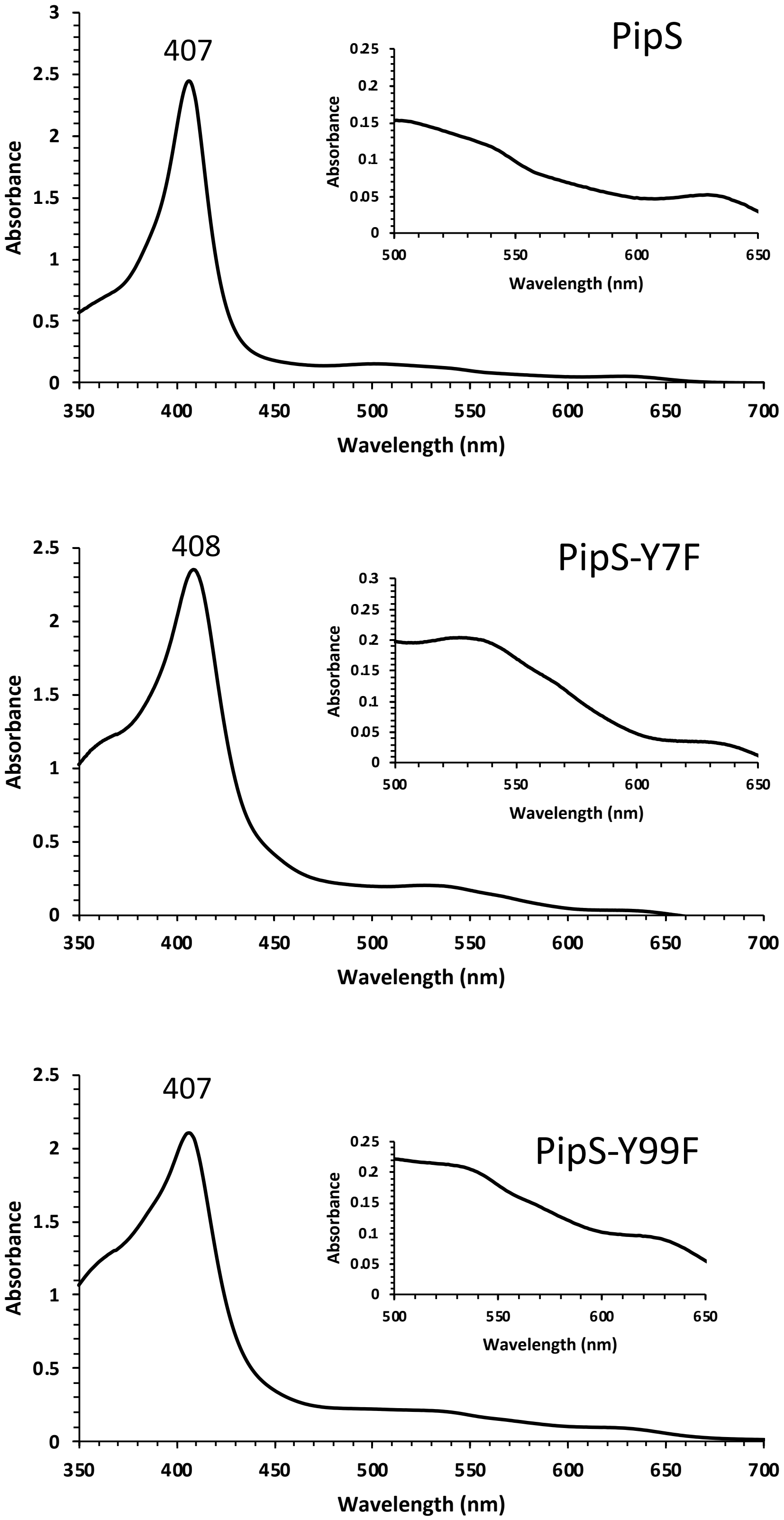

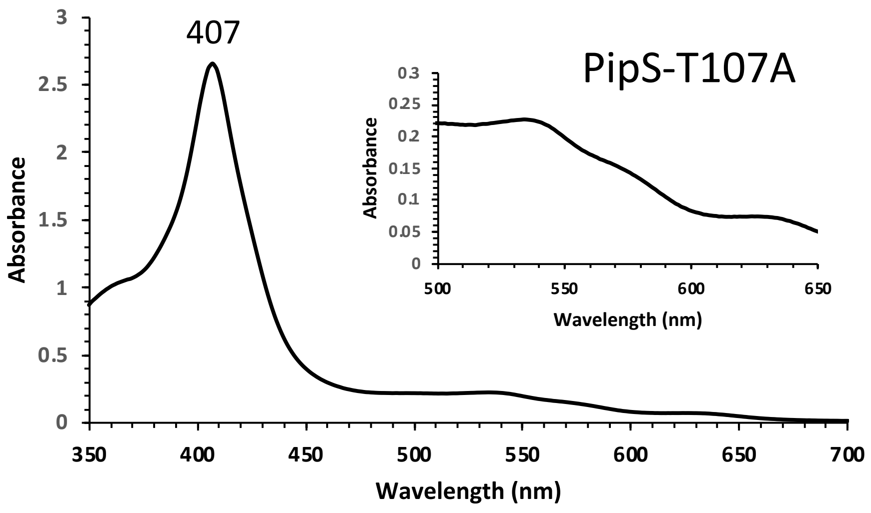
407
PipS-T107A
Absorbance
Wavelength (nm)

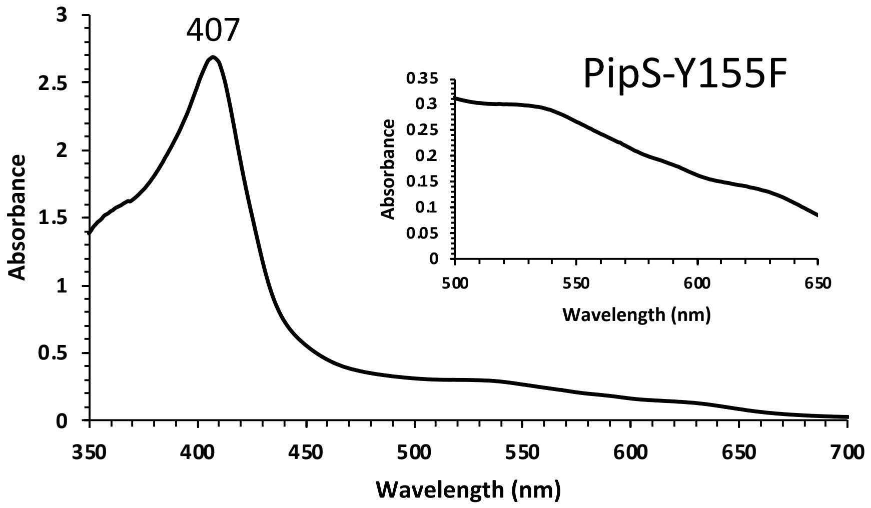
407
PipS-Y155F
Absorbance
Wavelength (nm)

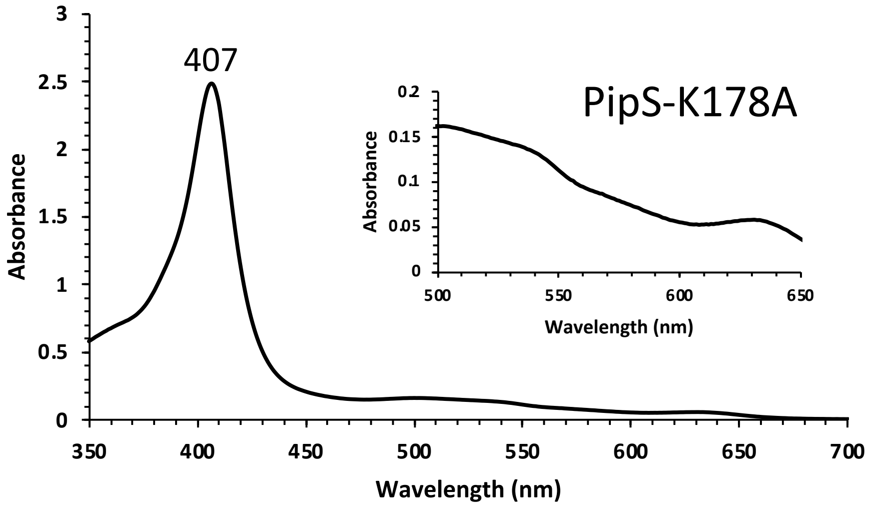
407
PipS-K178A
Absorbance
Wavelength (nm)

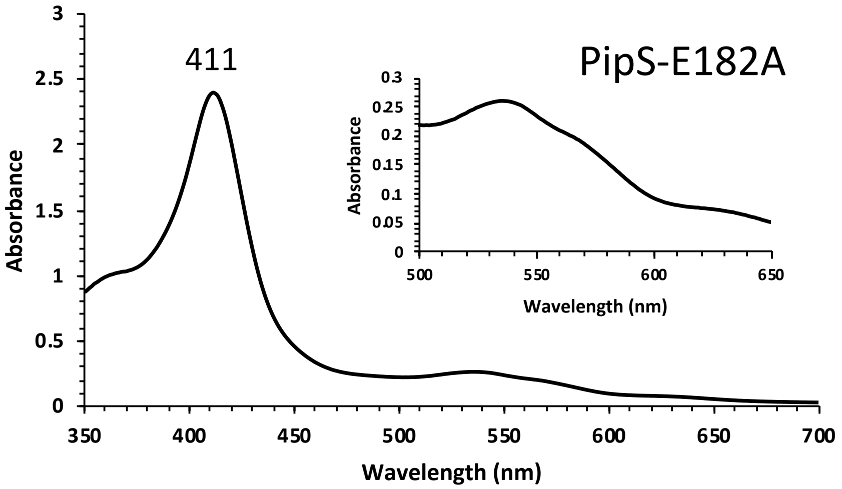
PipS-E182A
411
Absorbance
Wavelength (nm)

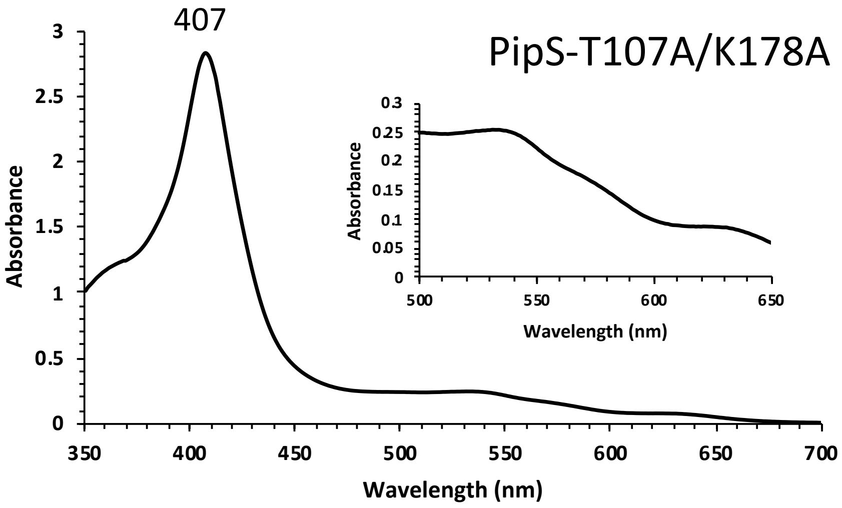
PipS-T107A/K178A
407
Absorbance
Wavelength (nm)

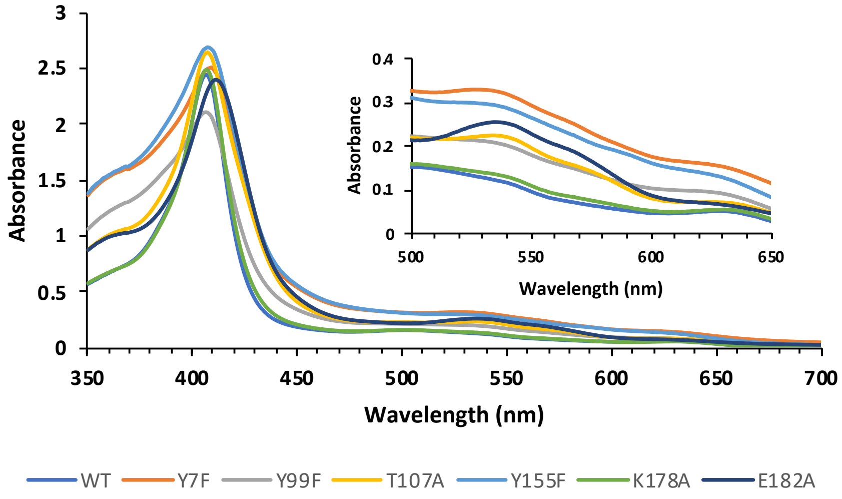
Absorbance
Wavelength (nm)
WT
Y7F
Y99F
T107A
Y155F
K178A
E182A

**Supplementary Figure 9**. Enzyme activities for PipS variants. **a**, HPLC traces of Fmoc-Cl derivatized samples of PipS and its variants. **b**, LC-MS analysis of reactions from (a). Note: *m/z* 262, $[M+Na]^+$ for Fmoc-$NH_2$. The synthetic substrate L-**1** sample contains trace amount of ammonia.

(**a**)

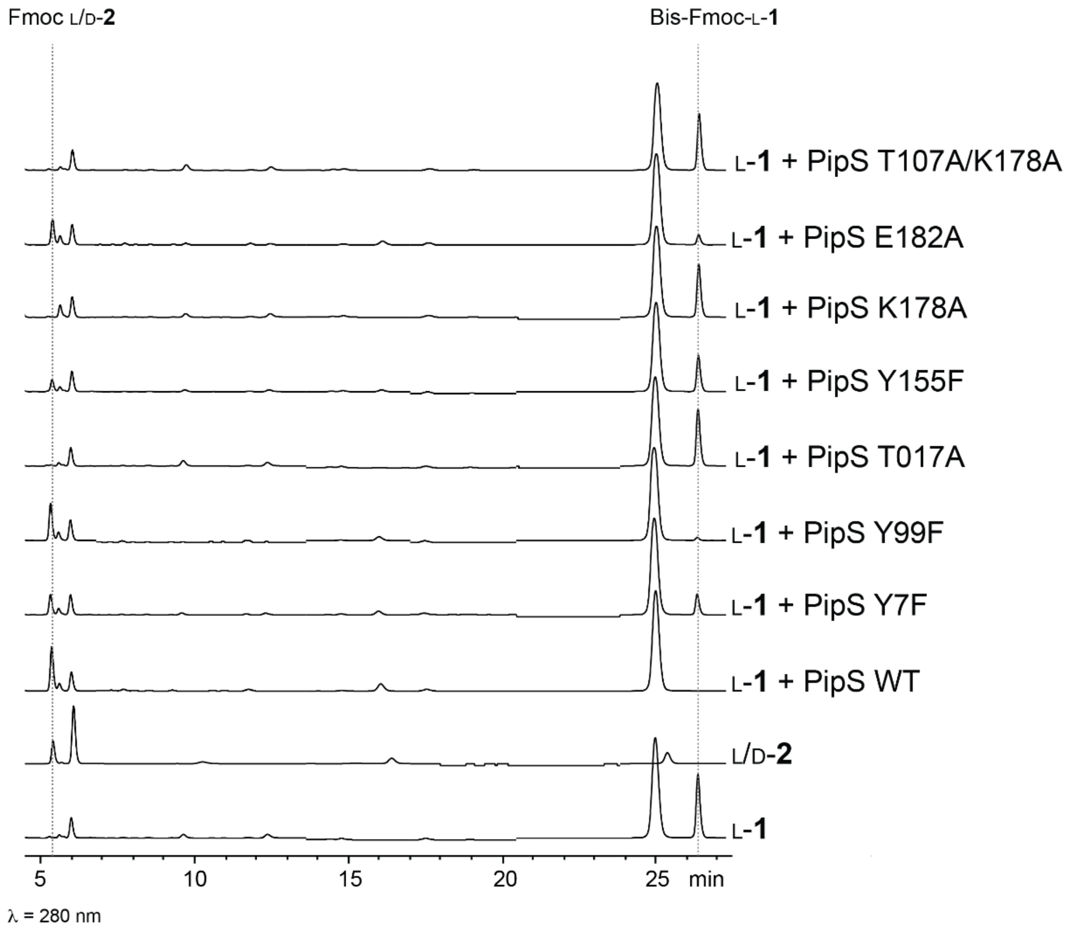

(**b**)

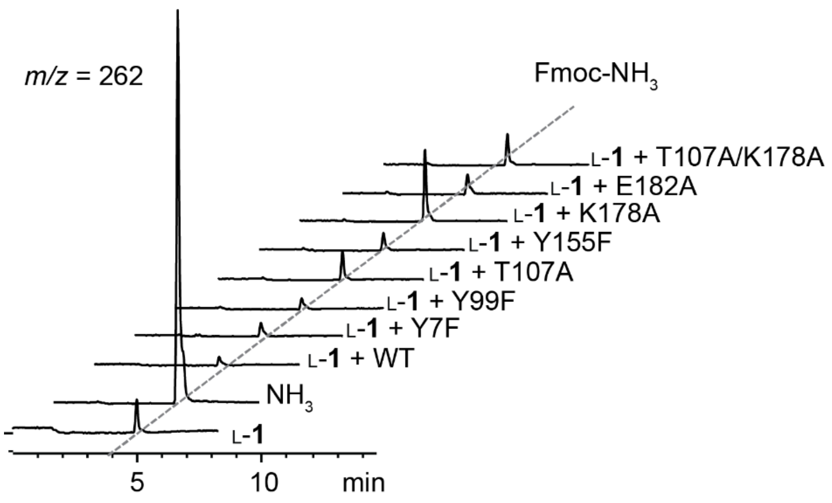

m/z = 262
Fmoc-NH3
L-1 + T107A/K178A
L-1 + E182A
L-1 + K178A
L-1 + Y155F
L-1 + T107A
L-1 + Y99F
L-1 + Y7F
L-1 + WT
NH3
L-1
5
10
min

**Supplementary Figure 10**. Synthesis and $^{1}$H NMR spectrum of **3**. Note: **3** was synthesized similarly as described above for L-**1**, with 5-aminopentanoic acid as the starting material. $^{1}$H NMR of synthetic **3**: $^{1}$H-NMR (400 MHz, $D_2O$) δ3.29 (t, $J$ = 7.5 Hz, 2H), 2.42 (t, $J$ = 7.1 Hz, 2H), 1.75 (m, 2H), 1.69 (m, 2H).

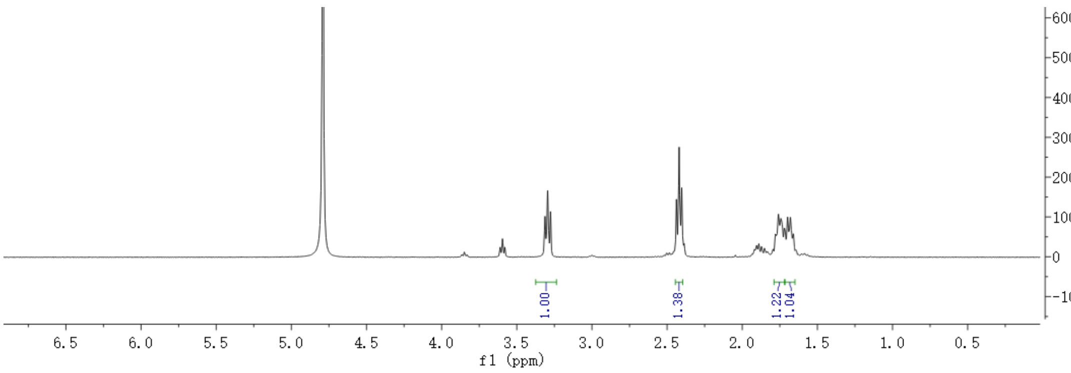

**Supplementary Figure 11**. Time-course studies of the reaction mixtures of **3** with as-isolated PipS (ferric) or dithionite-reduced PipS (ferrous) by HPLC after Fmoc-Cl derivatization. Detection wavelength=263 nm. The reaction mixture contains 2mM of synthesized **3**, 2 μM of PipS. For the reduced PipS, sodium dithionite (2 mM) was included. The results demonstrated that reduced PipS-catalyzed deamination of **3** completed within 10 min, whereas it took ~2h for as-isolated PipS to consume all the **3**.

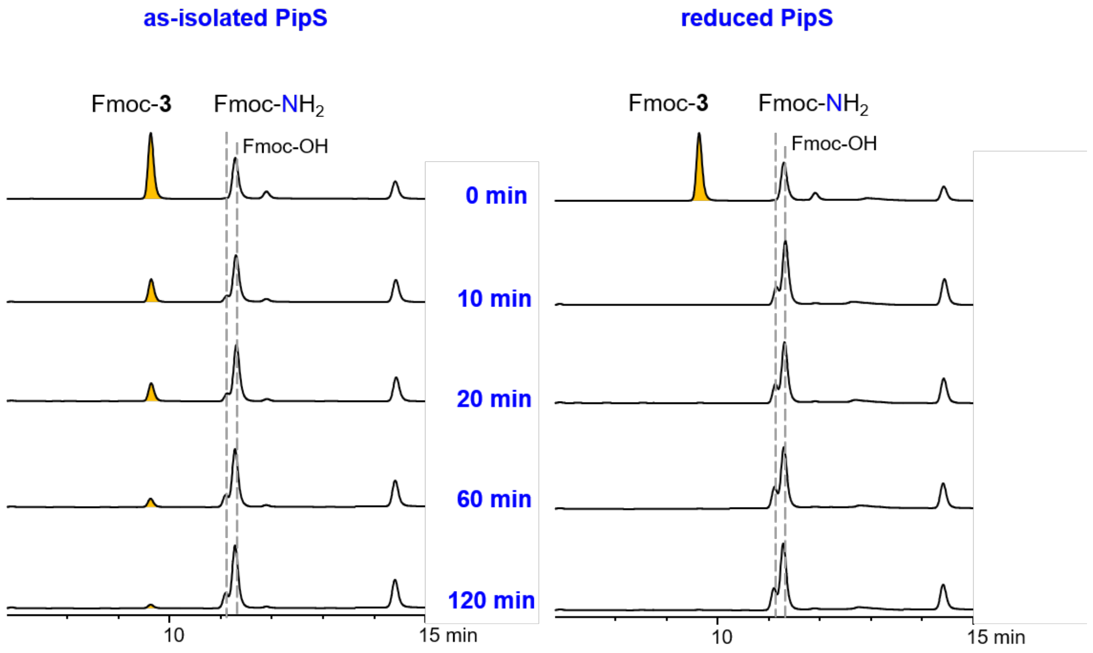

**Supplementary Figure 12**. HPLC analysis of the reaction mixture of PipS + **3** after DNPH (2,4-Dinitrophenylhydrazine) derivatization identified a product that displaying a *m*/*z* peak consistent with 5-oxopentanoic acid.

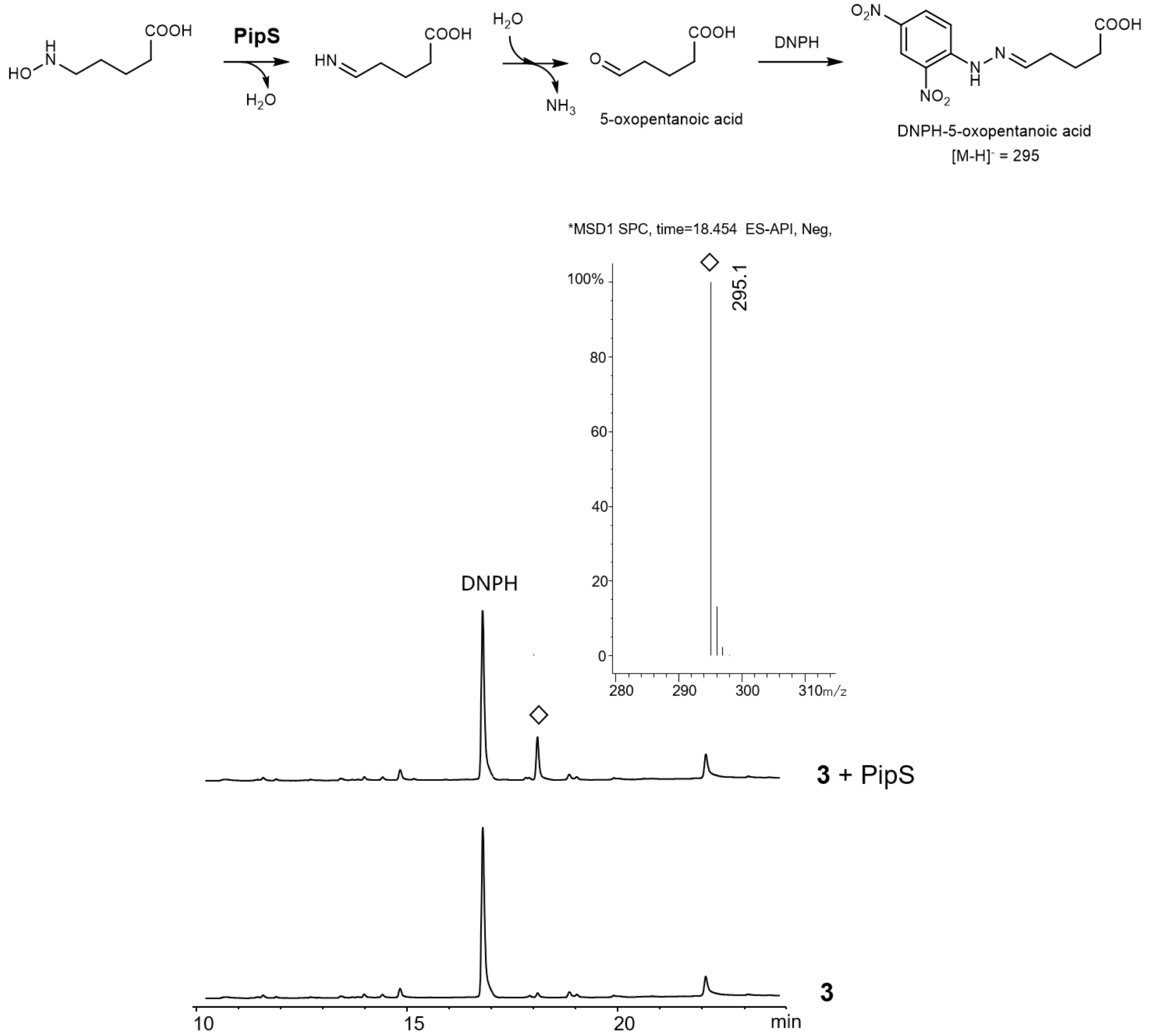

**Supplementary Figure 13**. Relative activities of various PipS variants when compared to **3** with PipS WT. Each reaction was performed in triplicate.

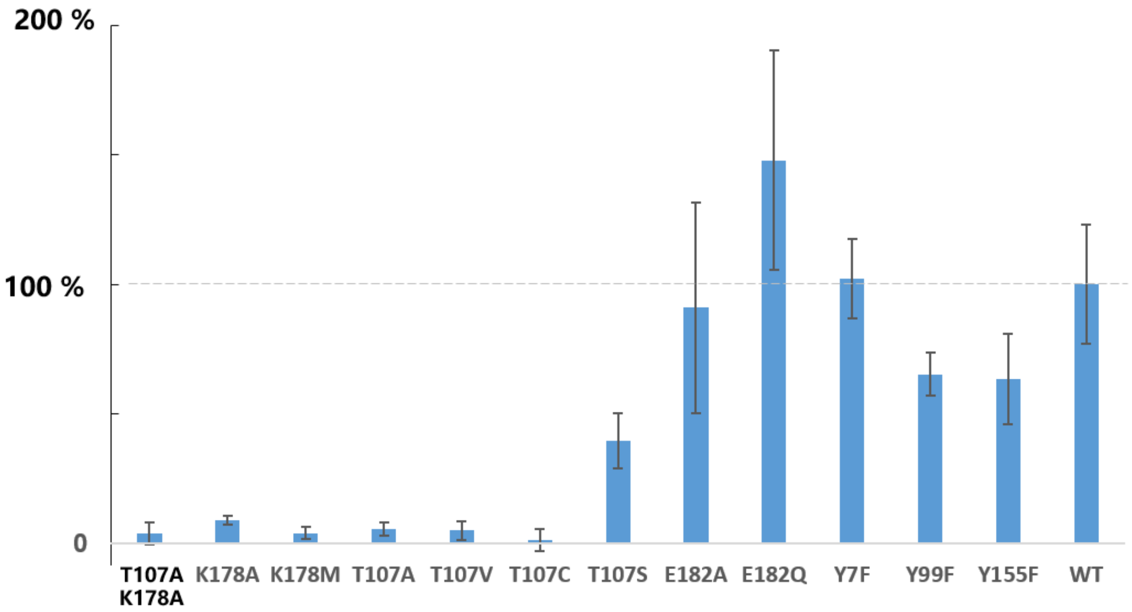

**Supplementary Figure 14**. Detection of the reaction product (dehydroproline) from PipS + D-**1** by chemical reduction by $NaBH_4$ and followed by Fmoc-Cl derivatization. EIC (*m*/*z* 338) from the LC-MS analysis of reaction mixtures were shown.

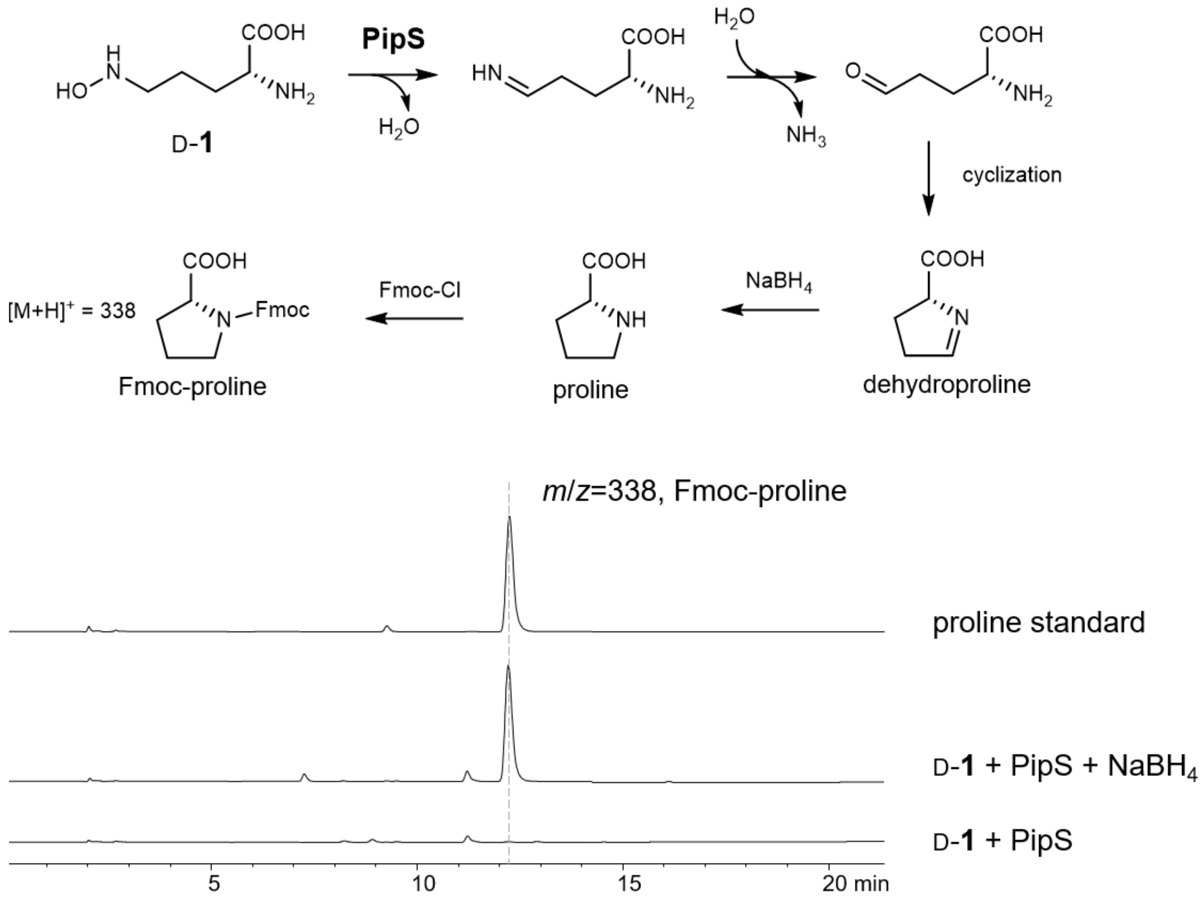

**Supplementary Figure 15**. Identification of *N*-benzyl-α-phenylnitrone (**11**) by NMR analysis. (**a**) NMR signal assignment for **11**. (**b**) $^{1}$H NMR spectrum (500 MHz, d$_6$-DMSO). (**c**) $^{13}$C NMR spectrum (125 MHz, d$_6$-DMSO). (**d**) COSY. (**e**) HSQC. (**f**) HMBC. (**g**) ESI-MS analysis.

**(a)**

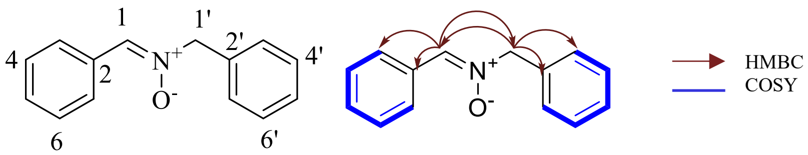

| No. | $^{1}$H-NMR (mult, J in Hz) | $^{13}$C-NMR |
|---|---|---|
| 1 | 8.10 (1H, s) | 133.3 |
| 2 | - | 131.0 |
| 3 | 8.22 (1H, dd, J = 7.55, 2.31 Hz) | 127.9 |
| 4 | 7.42 overlap | 129.9 |
| 5 | 7.38 overlap | 128.4 |
| 6 | 7.42 overlap | 129.9 |
| 7 | 8.22 (1H, dd, J = 7.55, 2.31 Hz) | 127.9 |
| 1′ | 5.07 (2H, s) | 70.2 |
| 2′ | - | 134.7 |
| 3′ | 7.49 (1H, dd, J = 7.95, 1.68 Hz) | 128.9 |
| 4′ | 7.41 overlap | 128.4 |
| 5′ | 7.38 overlap | 128.4 |
| 6′ | 7.41 overlap | 128.4 |
| 7′ | 7.49 (1H, dd, J = 7.95, 1.68 Hz) | 128.9 |

**(b)**

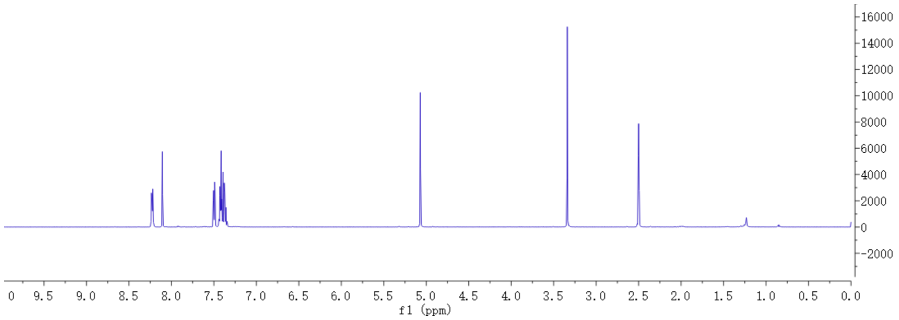

**(c)**

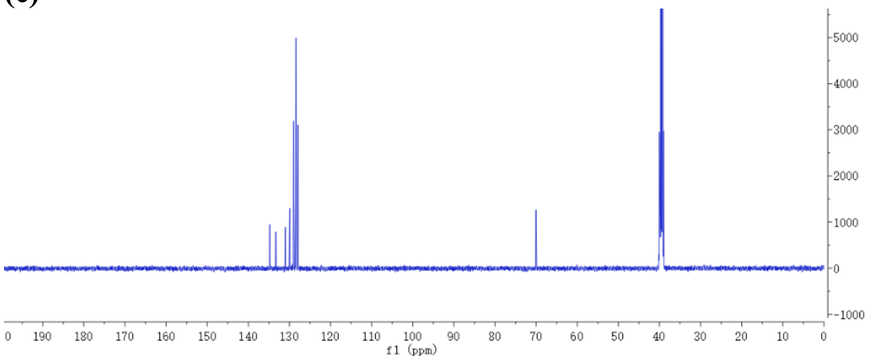

**(d)**

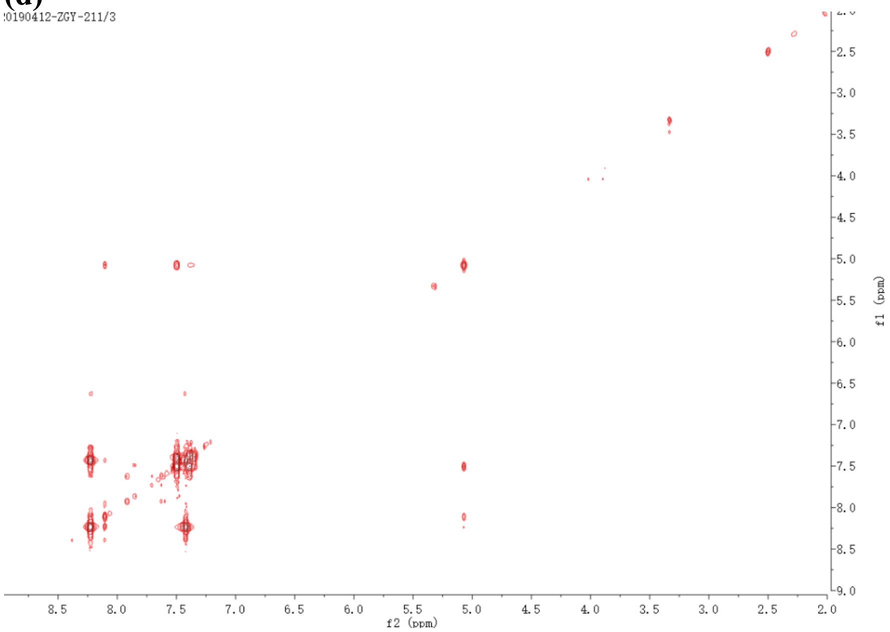

**(e)**

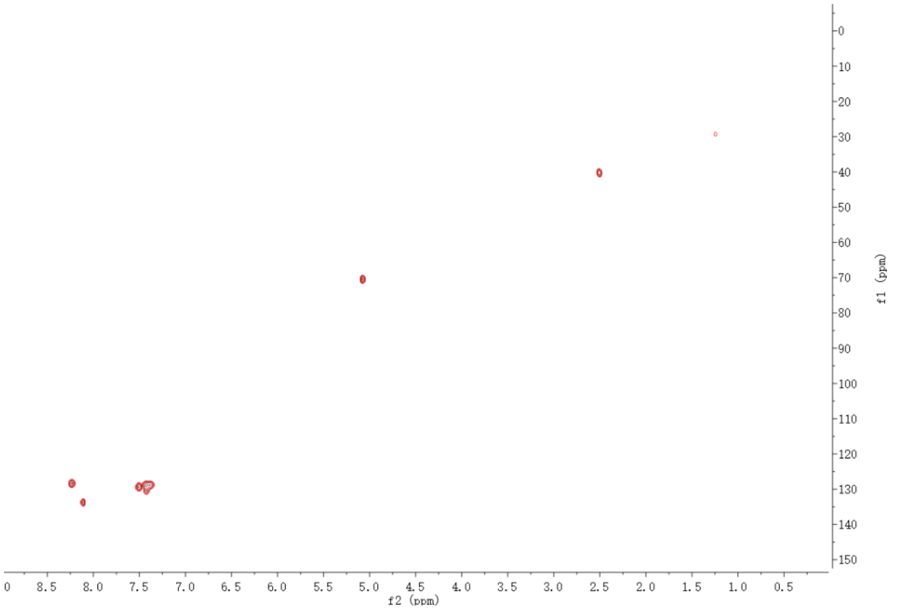

**(f)**

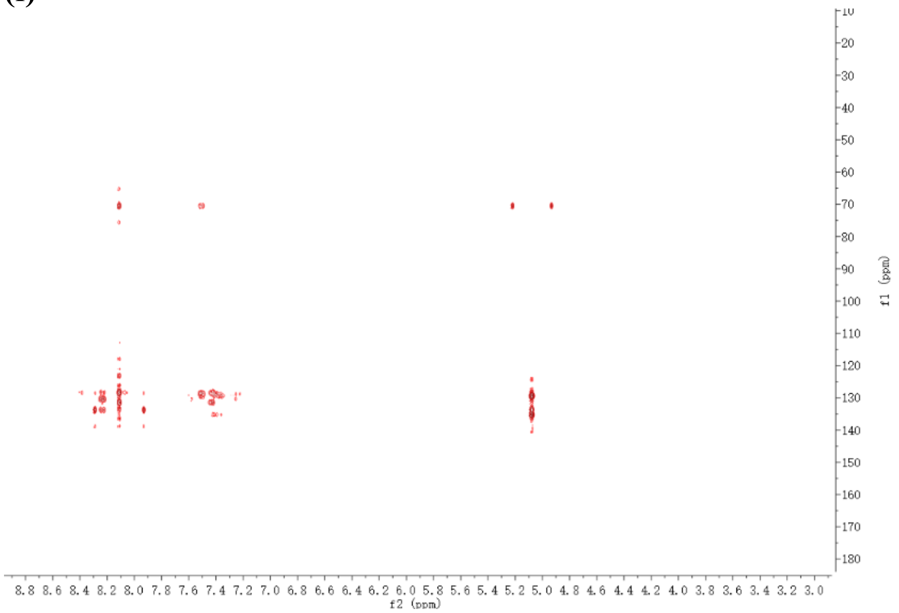

**(g)**

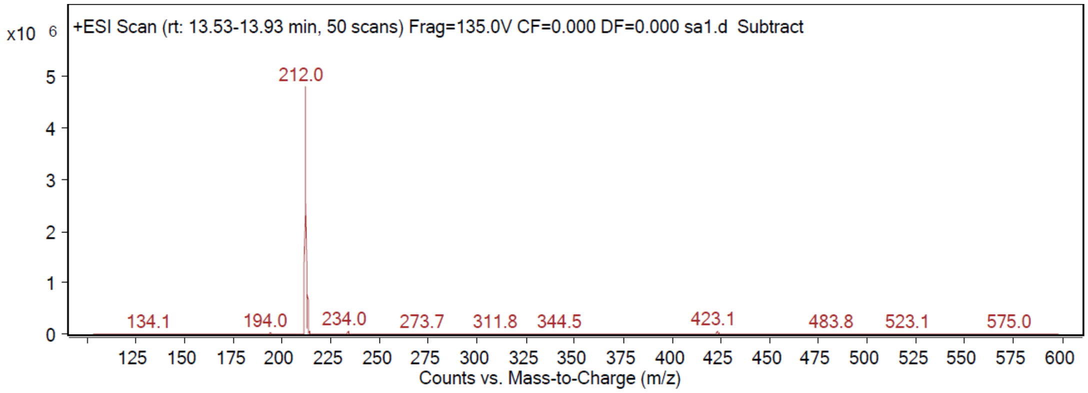

+ESI Scan (rt: 13.53-13.93 min, 50 scans) Frag=135.0V CF=0.000 DF=0.000 sa1.d Subtract
x10 6
212.0
134.1
194.0
234.0
273.7
311.8
344.5
423.1
483.8
523.1
575.0
Counts vs. Mass-to-Charge (m/z)

**Supplementary Figure 16**. HPLC analysis of reaction mixtures of four PipS homologs with **8** (**a**) (HPLC detection λ=254 nm) or L-**1** (**b**) (HPLC detection λ=263 nm). Note: the four codon-optimized PipS homologs (CfPipS from *Collimonas fungivorans* (WP_061538620.1), KsPipS from *Kibdelosporangium sp*. MJ126-NF4 (WP_042177946.1), PsPipS from *Pseudonocardia* (WP_060714104.1), and LfPipS from *Lentzea flaviverrucosa* (WP_090066830.1)) were synthesized, cloned into pET22b with NdeI and XhoI sites, and transformed into *E. coli* BL21 (DE3) for enzyme production. Crude extracts from *E. coli* strains containing these constructs were directly used for in vitro assays with 2 mM substrates.

**(a)**

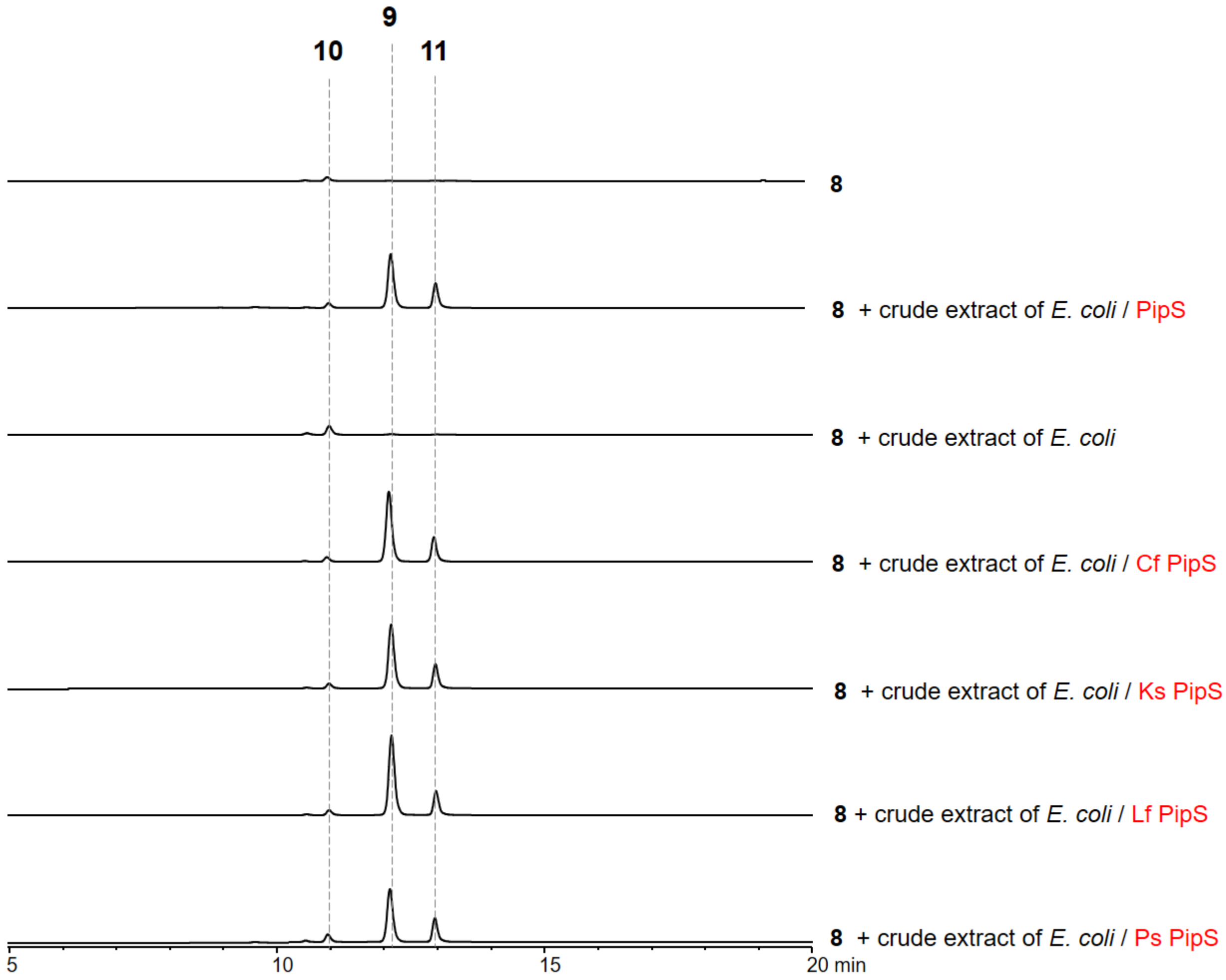

**(b)**

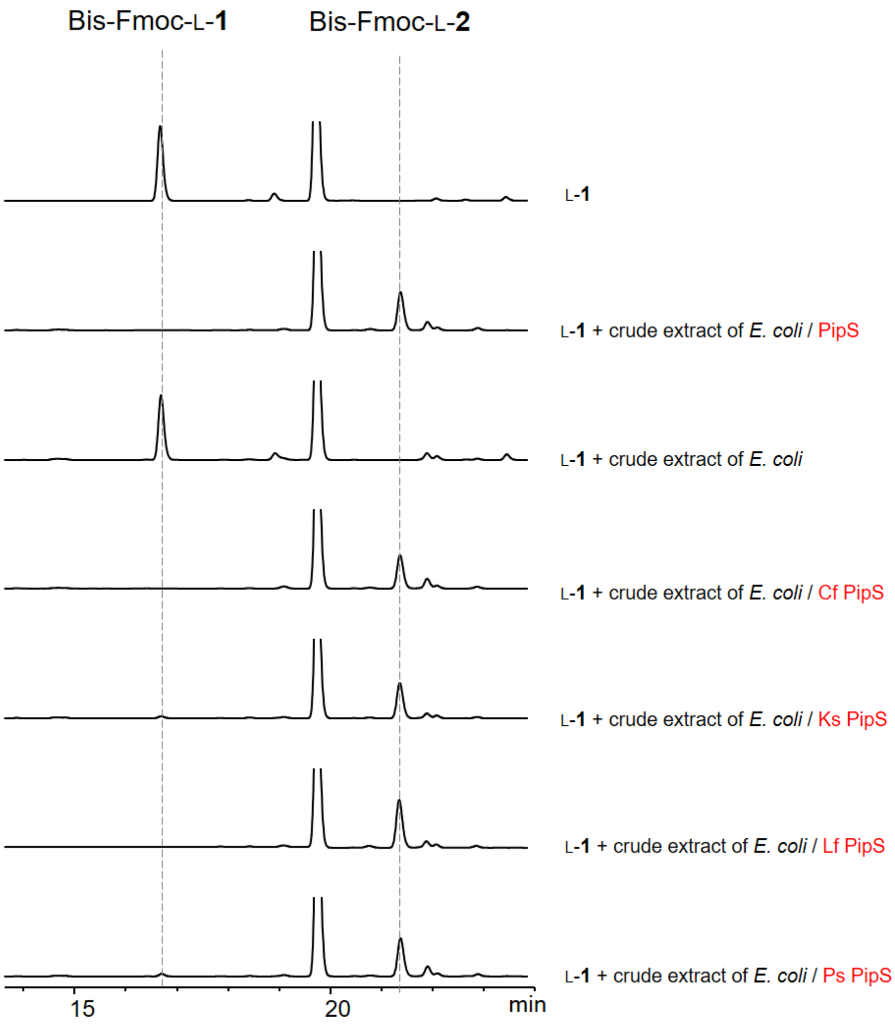
Bis-Fmoc-L-1
Bis-Fmoc-L-2
L-1
L-1 + crude extract of E. coli / PipS
L-1 + crude extract of E. coli
L-1 + crude extract of E. coli / Cf PipS
L-1 + crude extract of E. coli / Ks PipS
L-1 + crude extract of E. coli / Lf PipS
L-1 + crude extract of E. coli / Ps PipS
15
20
min

**Supplementary Figure 17**. Reaction of **8** or benzeneacetaldehyde oxime (BAAHO) with Oxd and PipS. (**a**) HPLC analysis (λ=263 nm) of reaction mixture of Oxd with **8** after Fmoc-Cl derivatization. The reaction mixtures contain 2mM of **8** and 2μM of as-isolated Oxd (ferric). (**b**) HPLC analysis (λ=254 nm) of reaction mixture of PipS with BAAHO. The reaction mixtures contain 2mM of BAAHO and 2μM of PipS (dithionite-reduced or as-isolated). The product (benzyl cyanide, BCN) control was generated in situ by incubation of BAAHO with reduced Oxd.

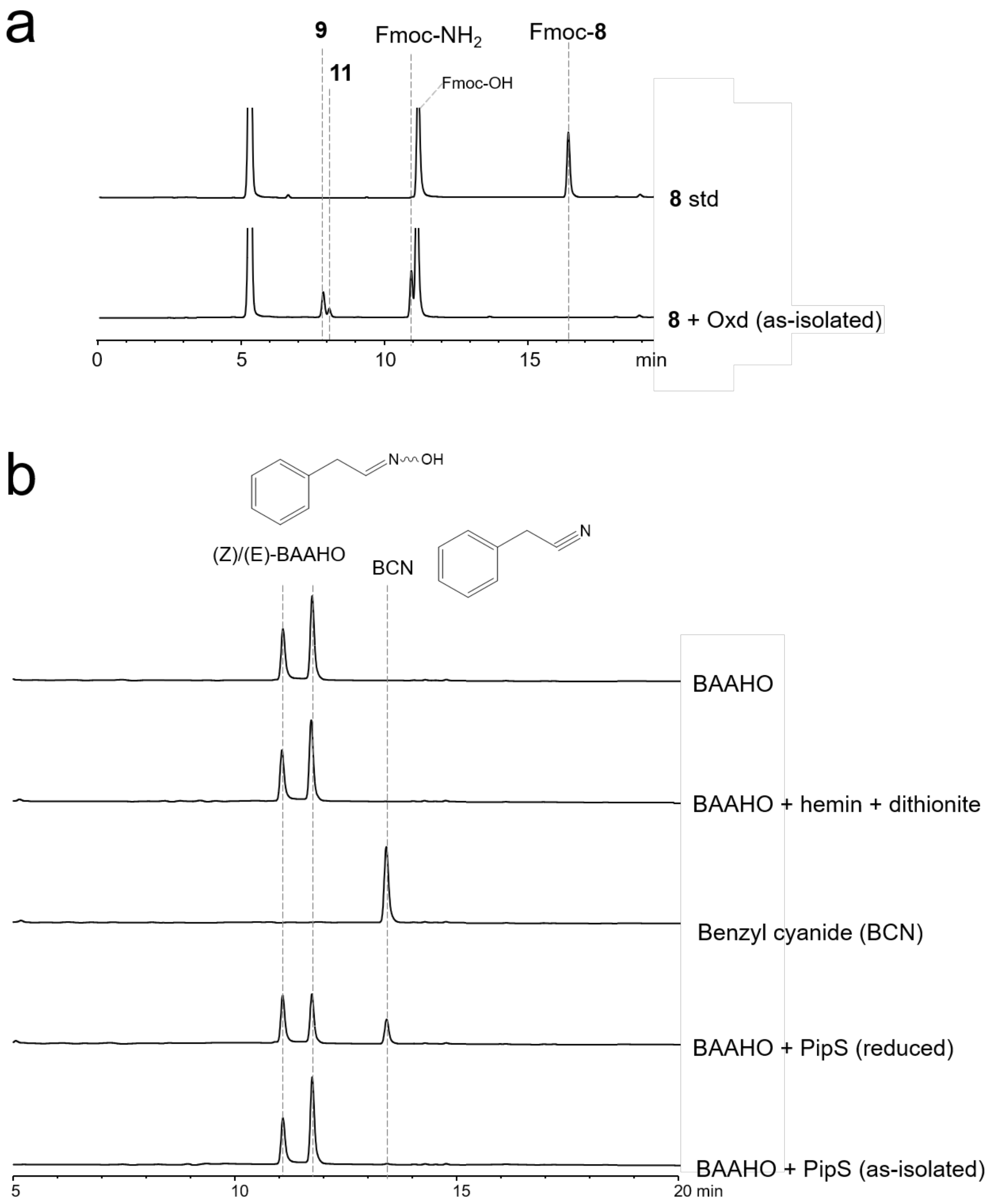

**Supplementary Figure 34**. Additional characterization of the 422nm species. **a**, Oxygraph of PipS WT or T107A-K178A with **12** or L-**1**. Oxygen concentration was monitored when 15 mM **12** or L-**1** was incubated with 400 μM PipS WT or T107A-K178A for 20 or 30 min. Samples from these reactions were removed every 10 min and analyzed by UV-Vis spectroscopy. **b**, PipS T107A-K178A incubated with 30mM L-**1** then desalted on a PD-10 column to remove excess L-**1** and 6 μM subjected to different temperatures and (**f**) oxidizing (100 μM $Fe(CN)_6K_3$) and reducing (5 mM dithionite) agents and monitored using UV-Vis spectroscopy. PipS-T107A-K178A holo or in complex with L-**1** incubated with 5 mM dithionite for 1 h and analyzed by UV-Vis spectroscopy. **C**, Reactions of PipS with L-**1** and **12**. Reactions were performed with 5 μM PipS, 2 mM L-**1**, and 5 mM **12** at room temperature for 1 h, derivatized with Fmoc-Cl and analyzed by HPLC show that **12** does not get consumed when incubated with PipS and does not inhibit a reaction with PipS and L-**1**.

**a**

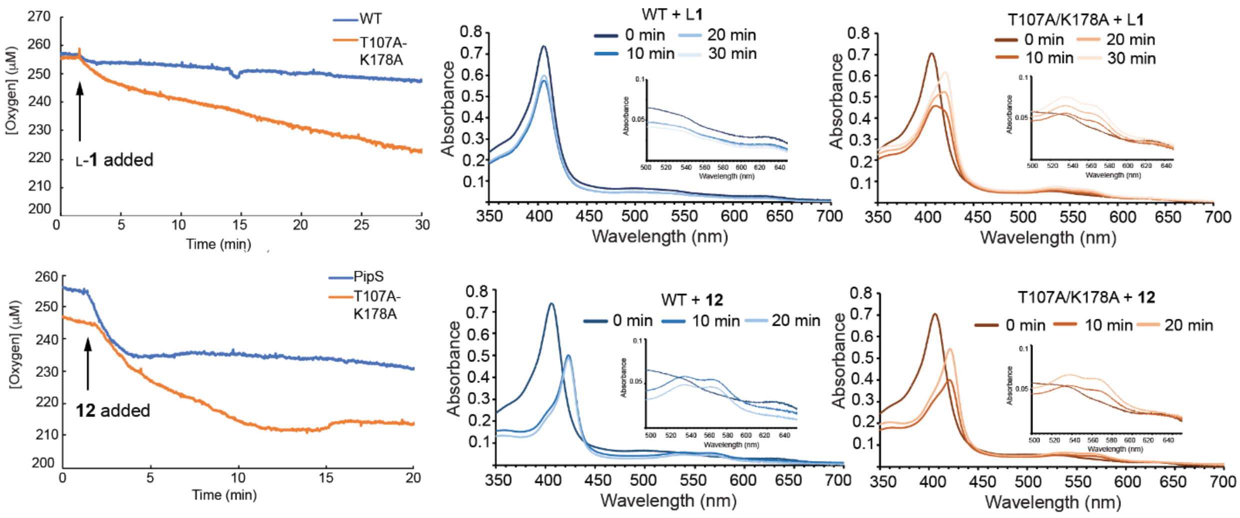

**b**

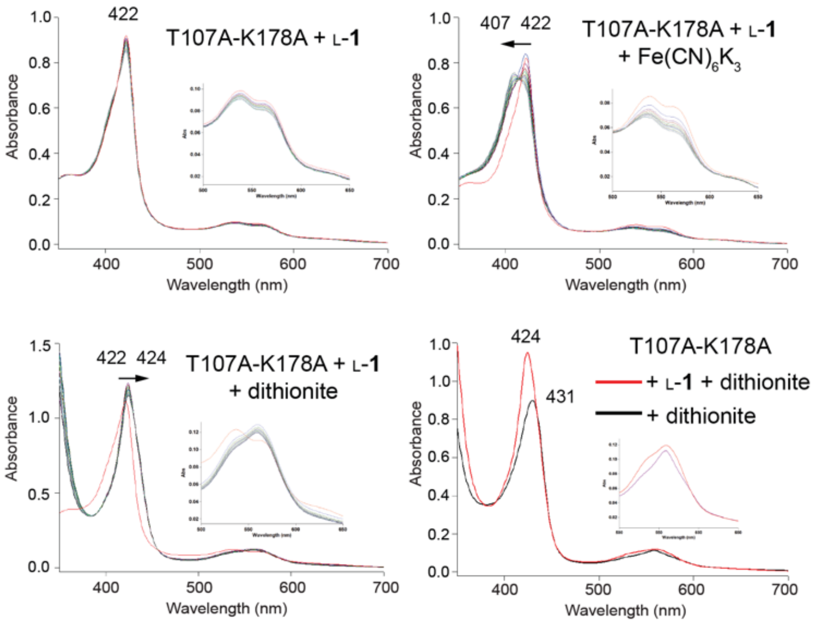
422
T107A-K178A + L-1
407 422
T107A-K178A + L-1
+ Fe(CN)6K3
422 424
T107A-K178A + L-1
+ dithionite
424
431
T107A-K178A
+ L-1 + dithionite
+ dithionite
Absorbance
Wavelength (nm)

**c**

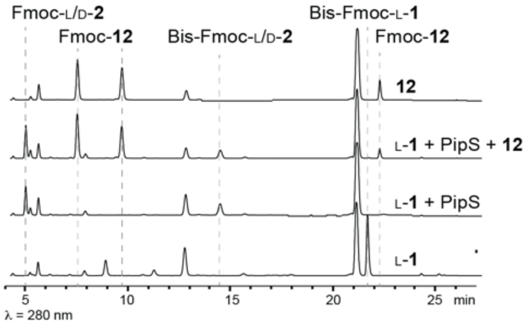
Fmoc-L/D-2
Fmoc-12
Bis-Fmoc-L/D-2
Bis-Fmoc-L-1
Fmoc-12
12
L-1 + PipS + 12
L-1 + PipS
L-1
5
10
15
20
25 min
λ = 280 nm

**Supplementary Figure 35.** rRaman (407 nm wavelength excitation, 20 mW power) spectrum of PipS T107A-K178A variant with **a**) L-**1** and **b**) isopropylhydroxylamine.

**a**

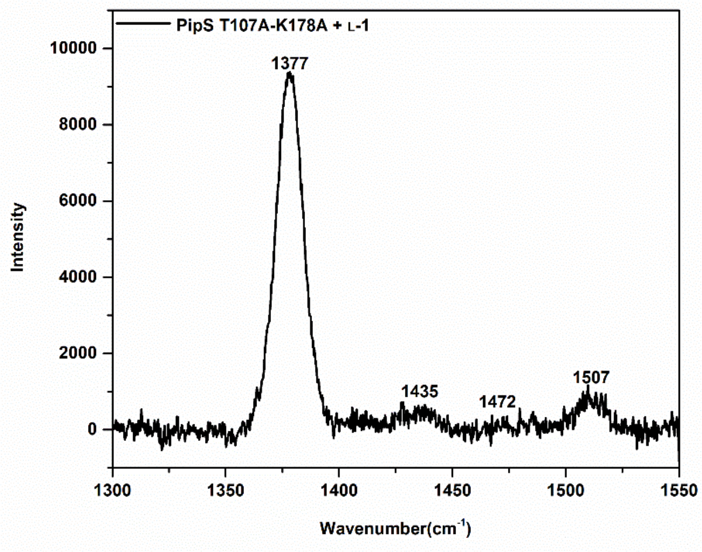

**b**

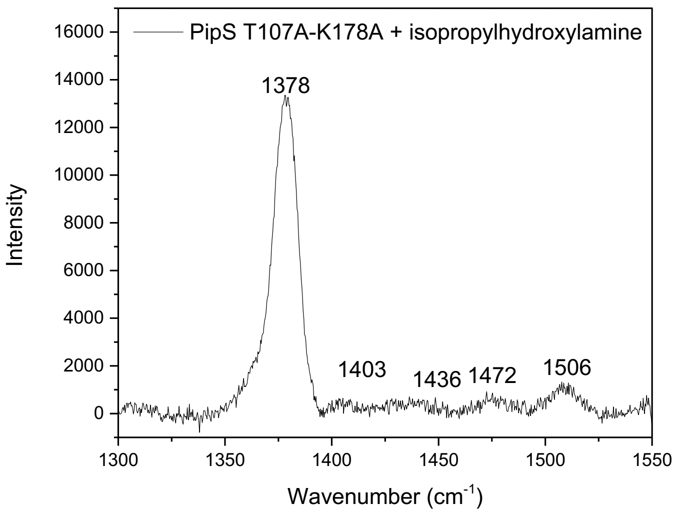

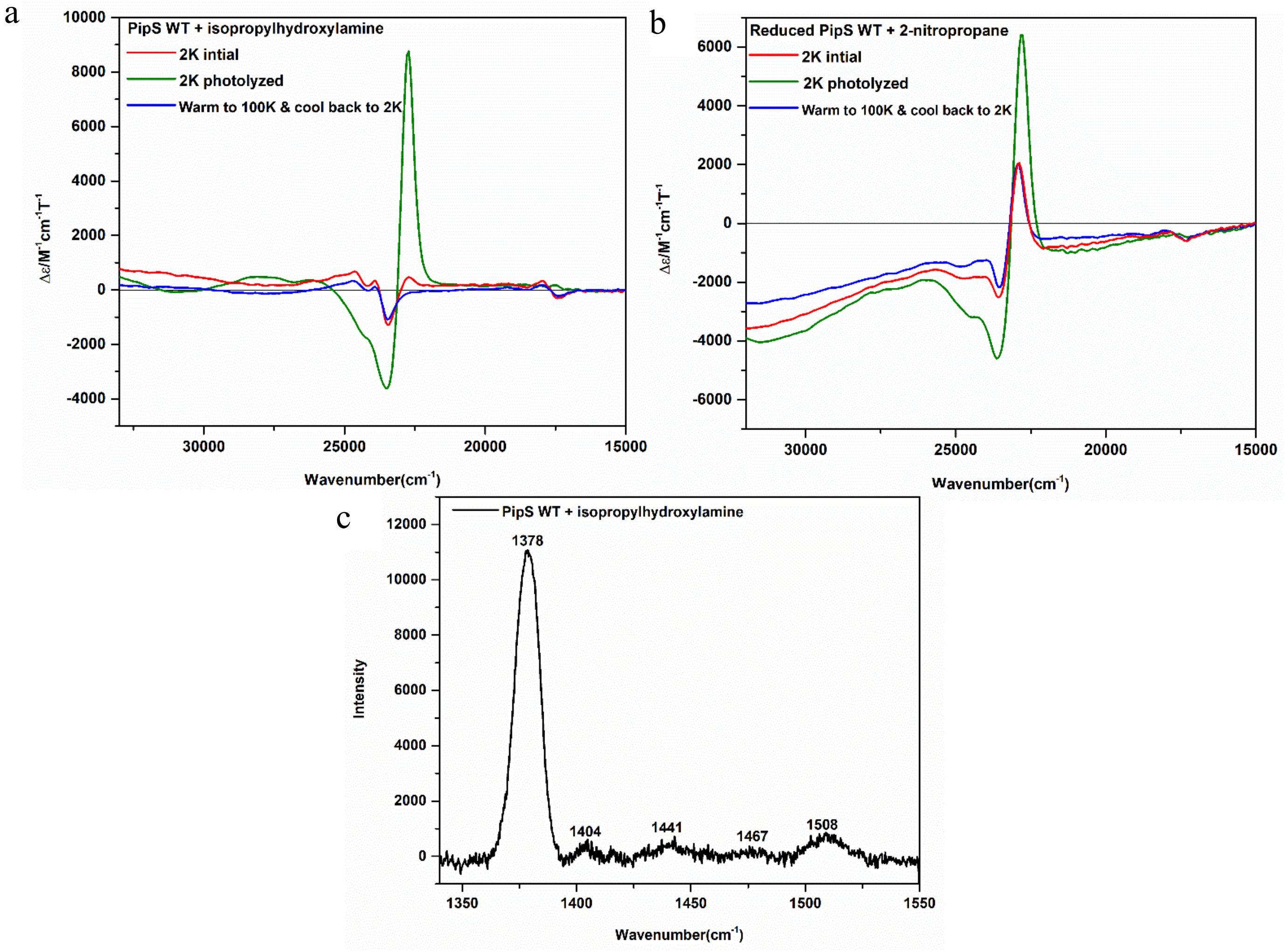

**Supplementary Figure 36.** MCD and rRaman analysis of PipS WT with isopropylhydroxylamine and 2-nitropropane. **(a)** MCD spectral overlay of PipS WT (50µM) with (+) isopropylhydroxylamine at 2K, 7T magnetic field (red), PipS WT + isopropylhydroxylamine photolyzed with a green laser laser (532nm, max power output 5mW) for 40min at 2K and 7T magnetic field (green), and PipS WT + isopropylhydroxylamine warmed to 100K and cooled back down to 2K at 7T magnetic field, after exposure to a green laser (blue). **(b)** MCD spectral overlay of PipS WT (50µM) reduced with excess dithionite and reacted with excess 2-nitropropane at 2K and 7T magnetic field (red), reduced PipS WT + 2-nitropropane photolyzed with a green laser (532nm, max power output 5mW) for 40min at 2K and 7T magnetic field (green), and reduced PipS WT + 2-nitropropane warmed to 100K and cooled back down to 2K at 7T magnetic field, after exposure to a green laser (blue). The data show that (a) and (b) lead to the formation of the same species, a ferrous heme alkyl-nitroso complex that is susceptible to photolysis, generating a five-coordinate HS ferrous heme. **(c)** rRaman (407 nm wavelength excitation, 20 mW power) spectrum of PipS WT + excess isopropylhydroxylamine. PipS WT was reduced with excess dithionite, which was confirmed by rRaman spectroscopy (Extended Data Figure 2) with the appearance of a ferrous marker band at 1352 $cm^{-1}$. The reduced PipS WT was reacted with excess 2-nitropropane to produce the RNO complex. The MCD spectrum of the RNO complex show similar spectral features as the PipS WT + isopropylhydroxylamine. Both PipS WT + isopropylhydroxylamine and reduced PipS WT + 2-nitropropane photolyze in the MCD. This is shown by the increase in intensity when exposed to a green laser over the course of 40 min at 2 K and 7 T magnetic field. rRaman spectra of Pips WT + isoHA show a major band 1378 $cm^{-1}$ which corresponds to the ferrous heme RNO complex, as described in the literature.[42–45]

### Part II

#### Computational Methods and Protocols

*Molecular Dynamics simulations*

The crystal structures for holo-PipS and substrate bound L-**1**-PipS in their dimeric form were used as starting points for MD simulations. Amino acid protonation states were predicted using the H++ server (http://biophysics.cs.vt.edu/H++) [46].

Molecular Dynamics simulations (MD) in explicit water were performed using the AMBER18 package [47].

Parameters for the substrate and the nitrenoid intermediate covalently bound to the Fe center were generated within the *antechamber* and MCPB.py [48] modules in AMBER18 package using the general AMBER force field (*gaff*) [49], with partial charges set to fit the electrostatic potential generated at the B3LYP/6-31G(d) level by the RESP model [50]. The charges were calculated according to the Merz–Singh–Kollman scheme [51,52] using the Gaussian 09 package [53].

The dimer was solvated in a pre-equilibrated cubic box with a 10-A buffer of TIP3P [54] water molecules using the AMBER18 leap module, resulting in the addition of ~14,000 solvent molecules. The systems were neutralized by addition of explicit counterions ($Na^+$ and $Cl^-$). All subsequent calculations were done using the AMBER force field 14 Stony Brook (ff14SB) [55]. A two-stage geometry optimization approach was performed. The first stage minimizes the positions of solvent molecules and ions imposing positional restraints on solute by a harmonic potential with a force constant of 500 kcal $mol^{-1}A^{-2}$, and the second stage is an unrestrained minimization of all the atoms in the simulation cell.

The systems were gently heated using six 50 ps steps, incrementing the temperature by 50 K for each step (0–300 K) under constant-volume and periodic-boundary conditions. Water molecules were treated with the SHAKE algorithm such that the angle between the hydrogen atoms was kept fixed. Long-range electrostatic effects were modelled using the particle-mesh-Ewald method [56]. An 8 Å cutoff was applied to Lennard–Jones and electrostatic interactions. Harmonic restraints of 30 kcal·$mol^{-1}$ were applied to the solute, and the Langevin equilibration scheme was used to control and equalize the temperature. The time step was kept at 1 fs during the heating stages, allowing potential inhomogeneities to self-adjust.

Each system was then equilibrated for 2 ns with a 2 fs time step at a constant pressure of 1 atm and temperature of 300 K without restraints. Once the systems were equilibrated in the NPT ensemble, production trajectories were then run for an additional 500 or 1000 ns (0.5 or 1 μs) under the NVT ensemble and periodic-boundary conditions. A total of 4 independent replicas of 500 or 1000 ns for each studied system were performed. Trajectories were processed and analyzed using the cpptraj [57] module from Ambertools utilities.

*Hybrid Quantum Mechanics/Molecular Mechanics (QM/MM) calculations*

QM/MM initial structures were manually selected from MD trajectories. All water molecules and counterions beyond 3 Å from any residue of the protein, cofactors or substrates were removed and the two resulting systems had 10371 and 10383 atoms for the substrate bound and nitrenoid intermediate bound, respectively.

The QM region included the heme porphyrin pyrrole core and a propionate group that directly interacts with the substrate, the His66' imidazole moiety, the iron center, Thr107 sidechain and the methylammonium moiety of Lys178, as well as the entire substrate or intermediate which lead to 83 or 80 QM atoms, 10 H-link atoms, and a neutral QM charge. All residues and water molecules outside a 12 Å shell from the QM region were keep frozen, thus resulting in more than 2.600 free MM atoms.

QM/MM calculations were carried out with ONIOM as implemented in Gaussian09 [53]. Geometry optimizations were performed with the hybrid (U)B3LYP [58–60] functional with an SDD basis set for iron and 6-31G(d) on all the other atoms, using an ultrafine integration grid and the QuadMacro optimization algorithm. The MM parameters were identical to those used in the MD simulations. A two-step sequential optimization protocol has been used: *i*) a first optimization using a Mechanical Embedding (ME) scheme was initially performed, and once optimized, *ii*) MM water molecules were kept frozen and a second optimization was performed within the Electrostatic Embedding scheme (EE). Stationary points were verified as minima or saddle point geometries after vibrational frequency analysis and thermal corrections were obtained at 1 atm and 298.15 K. Single point calculations from the latest optimized structure were performed at the (U)B3LYP/Def2TZVP theory level with EE scheme. MolUP VMD extension [61] was used for input preparation and output visualization.

The methodology employed in this study, based on the use of (U)B3LYP density functional, is very similar to the one used by Shaik group for the study of iron carbene porphyrins [62], and also by us in our recent studies on carbene transferases [63,64]. (U)B3LYP has also been extensively proved to accurately perform in the computational modeling of iron-oxo chemistry [65–69].

**Supplementary Figure 18. Conformational arrangement of K178-T107 catalytic dyad and L-1 in PipS active site from MD simulations.** Key distances explored by K178-T107 catalytic dyad and L-**1** substrate when bound in PipS and key L-**1** N-N distance, from four independent MD simulations and considering the two independent active sites from the PipS dimer. Distances are given in angstrom (Å).

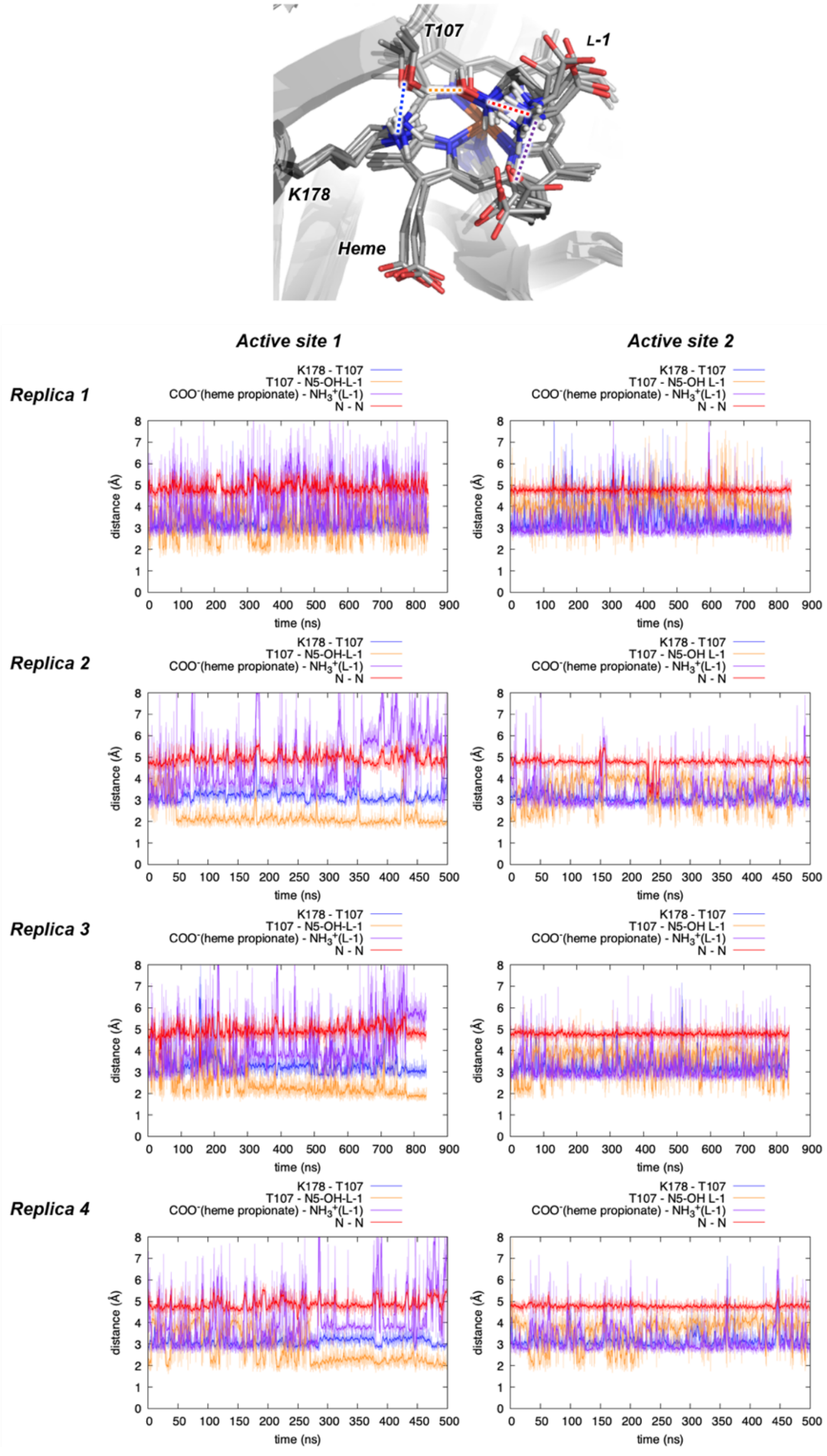

A total of 4 independent replicas starting from the X-ray structure with L-**1** bound in PipS (PipS-L-**1**). This implies independent minimizations, heatings and equilibrations during MD preparation protocols prior MD production runs (see Computational Methods), which ensure variability for the initiation of the different independent MD trajectories (production runs). Two different substrates, one at each active site of the PipS dimer, are considered. Production runs were propagated during 500 ns, which represent enough simulation time to capture relevant conformational changes of the substrate and the enzyme that might affect to the active site cavities (side chain rotations, flexible loop displacements). Two replicas (replica 1 and 3) were propagated up to 850 ns to ensure that longer MD trajectories would provide similar conclusions.

MD simulations starting from the X-ray structure with L-**1** bound in PipS (PipS-L-**1**) were performed to analyze the conformational arrangement of the substrate and the catalytic K178-T108. Simulations indicate that the distance between the two nitrogens from the α-amine and the N-OH group remains constant (ca. 5.0 Å) during the MD trajectories. This distance is equivalent to the one observed in the X-ray structure.

On the other hand, K178 and T107 residues establish a persistent H-bond that is maintained during the whole MD trajectory (distance ca. 3.0 Å), and the α-amino group also strongly interacts with the heme propionate (distance ca. 3.0 Å).

MD simulations also describe that T107 interacts with the hydroxyl group of the substrate during most of the simulation time, indicating a good preorganization for hydroxylamine activation.

A representative snapshot from these MD trajectories was used as starting point for further mechanistic interrogations using QM/MM calculations (see QM/MM calculations in **Supplementary Figure 26**). The snapshot was selected based on RMSD clustering of the protein backbone, and choosing a representative snapshot of the most populated cluster.

**Supplementary Figure 19. Interactions between L-1 and proximal residues in PipS active site explored by MD simulations.** Key distances between Y7, Y99, Y155’, E182, Q183, and K178 polar residues and L-**1** substrate when bound in PipS, explored in four independent MD simulations and considering the two independent active sites from the PipS dimer. Distances are given in angstrom (Å).

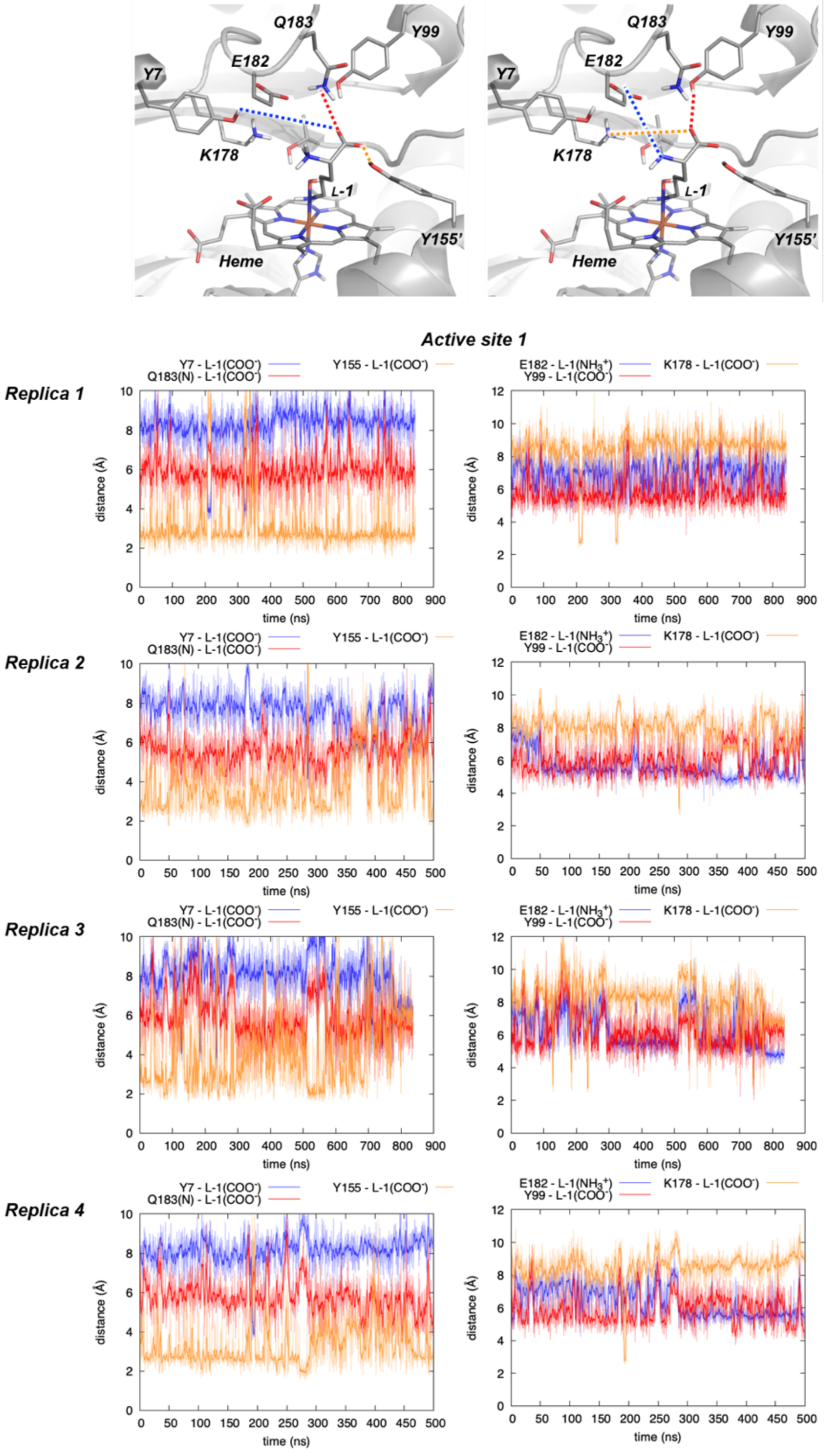

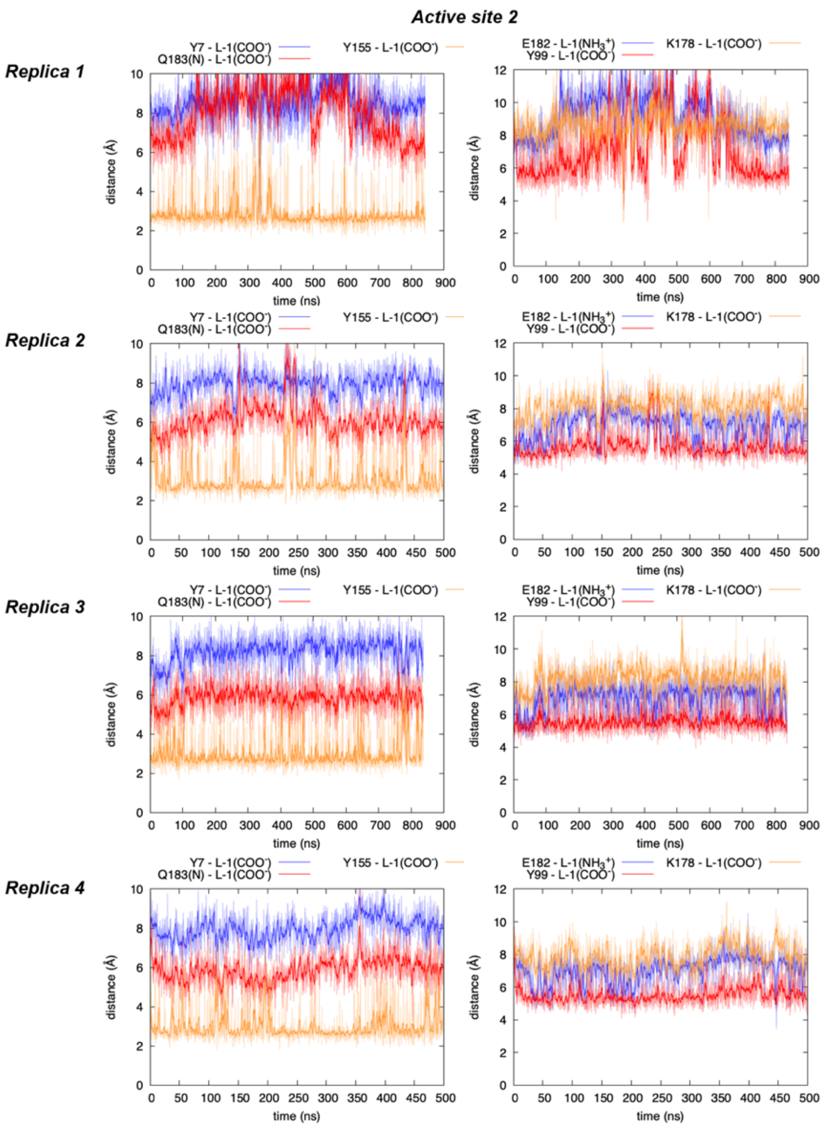

The same MD simulations reported in **Supplementary Figure 18** were used to analyze the principal interactions occurring between substrate L-**1** and PipS active site residues.

MD simulations showed that the major interaction is occurring between the carboxylate group of L-**1** and Y155 via H-bond (distance ca. 3.0 Å), which is maintained during most of the simulation time. This interaction is equivalent to the one observed in the X-ray structure.

**Supplementary Figure 20. Interactions between proximal residues in PipS-L-1 complex distal axial site explored by MD simulations**. Key distances between Y7, Y99, E182, Q183, T107 and K178 polar residues in PipS-L-**1** complex are explored in four independent MD simulations and considering the two independent active sites from the PipS dimer. Distances are given in angstrom (Å).

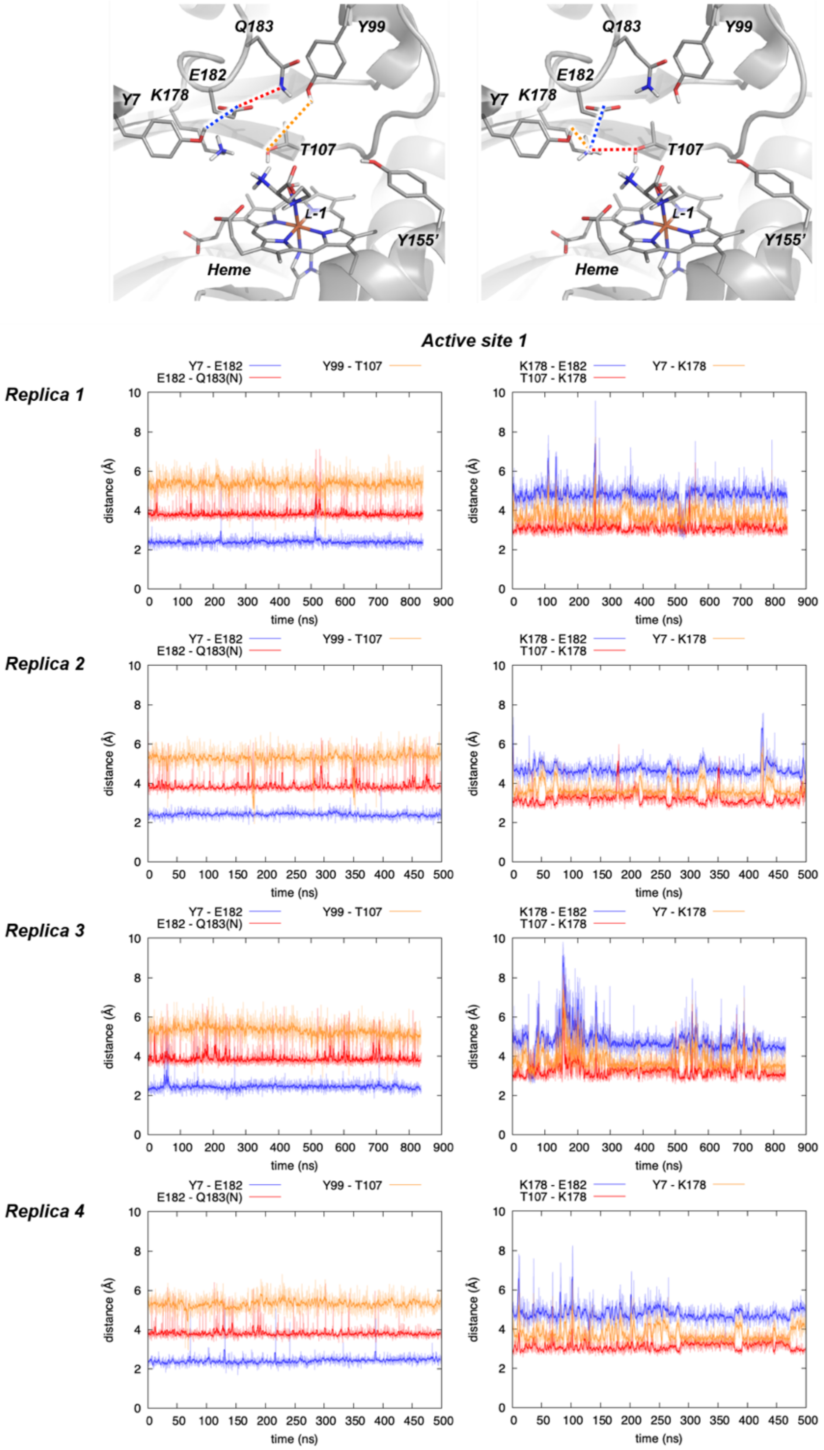

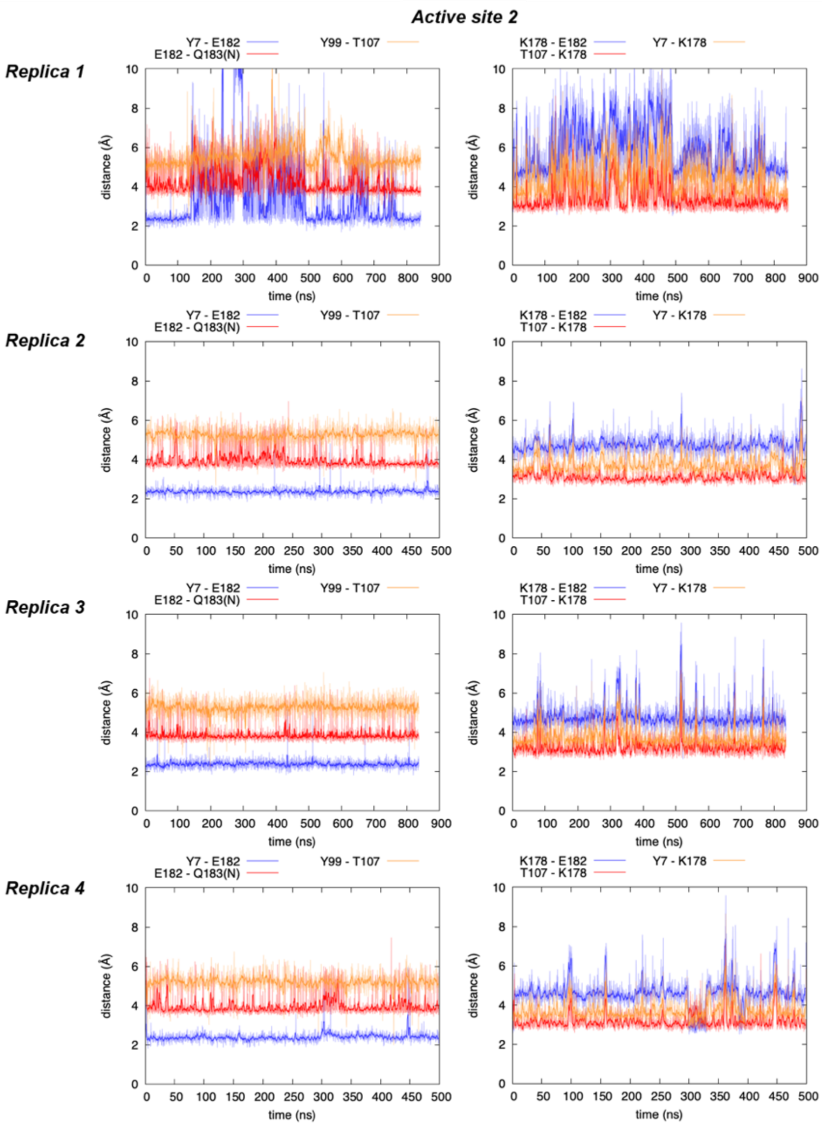

The same MD simulations reported in **Supplementary Figure 18** were used to analyze the hydrogen bond network established between PipS active site residues.

MD simulations described the existence of a persistent H-bond network that is maintained during the whole simulation time. This H-bond network involves the H-bond interactions established between Y7 and E182, and at the same time the interaction established between Y7 and K178, and E182 and Q183. As expected, K178 is H-bond interacting with T107 during the whole simulation time (see also **Supplementary Figure 18**).

**Supplementary Figure 21. Interactions between proximal residues and L-1 N5-OH and C5 positions in PipS-L-1 complex explored by MD simulations.** Key distances between polar residues capable of protonate/deprotonate L-**1** at N5-OH and C5 positions in PipS-L-**1** complex are explored in four independent MD simulations and considering the two independent active sites from the PipS dimer. Distances are given in angstrom (Å).

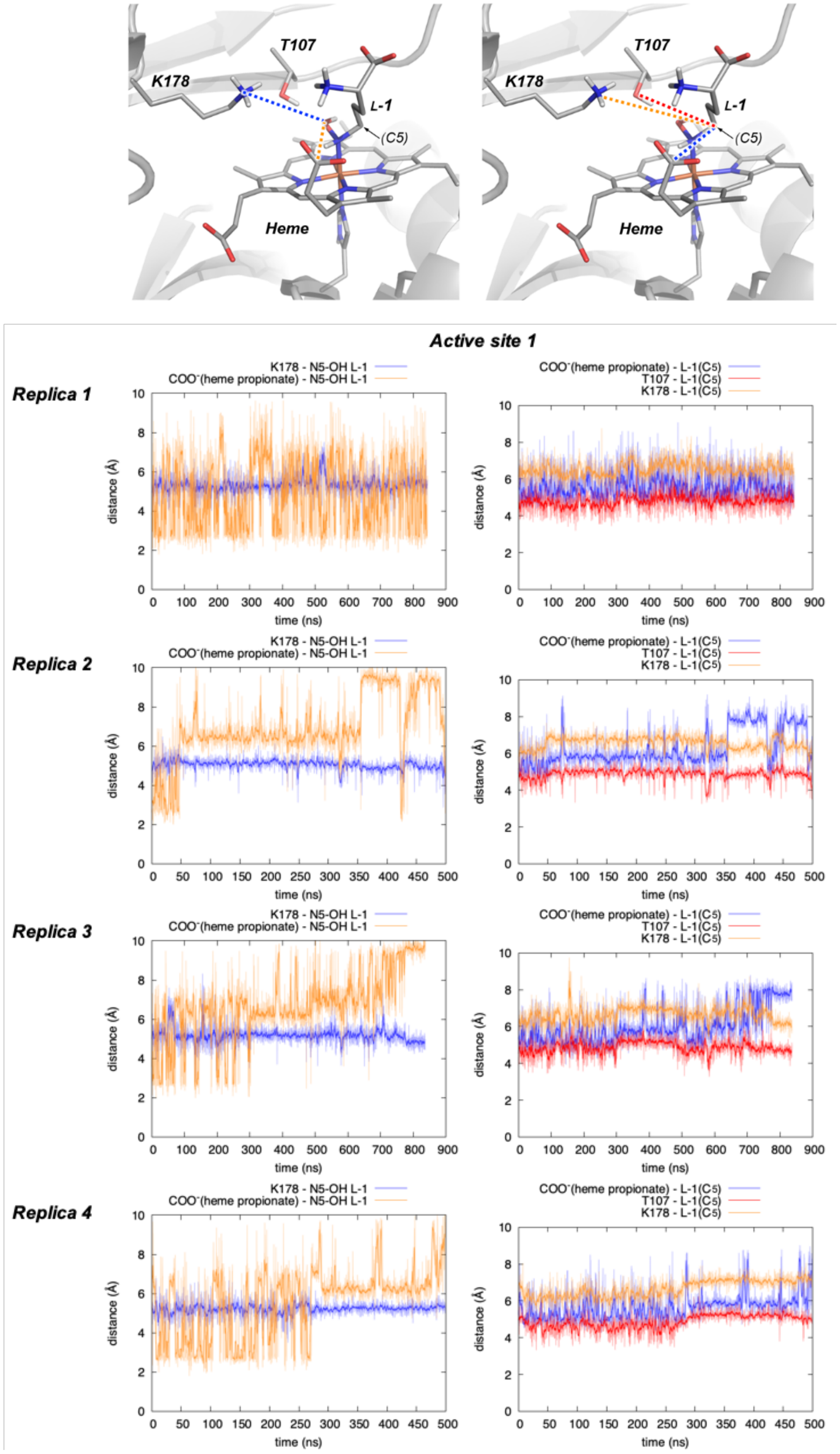

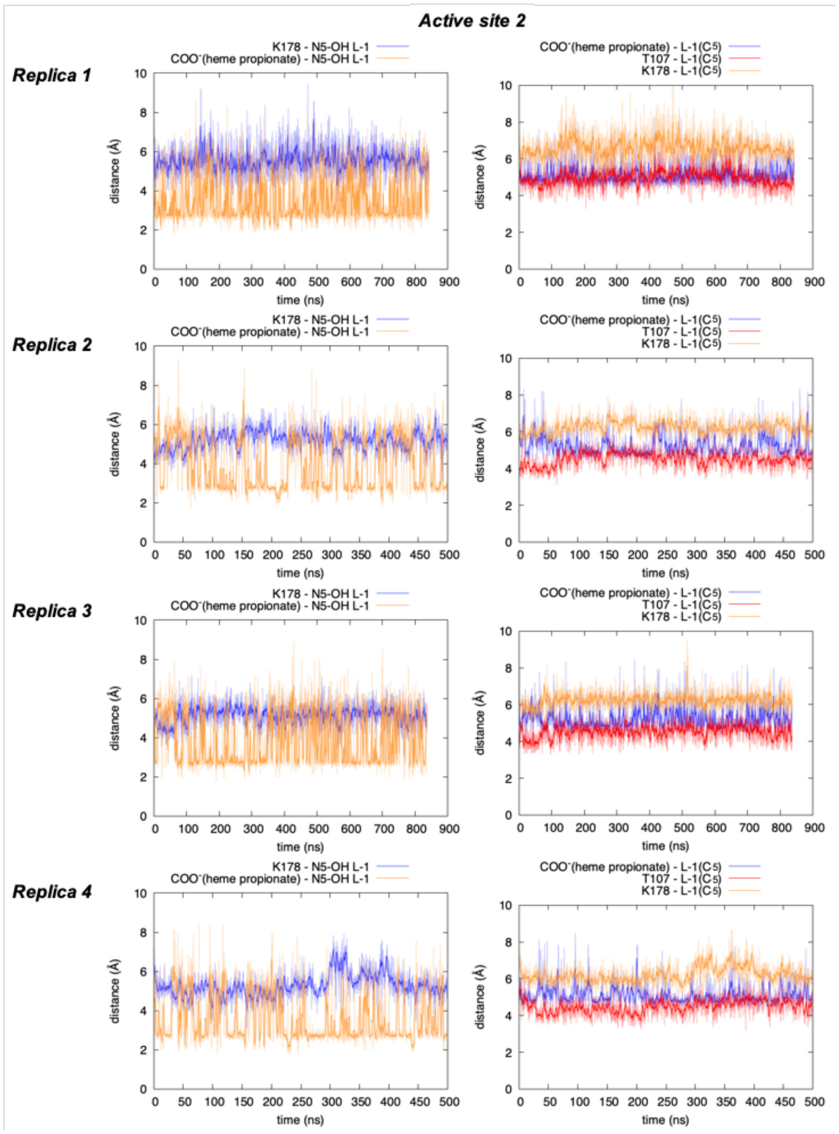

The same MD simulations reported in **Supplementary Figure 18** were used to analyze the potential interactions between OH and C5 positions from substrate L-**1** and PipS active site residues that could act as a base to deprotonate it.

MD simulations described that, in general, C5 position stays far (distances ca. > 5.0 Å) from the heme propionate group and T107, suggesting that these are not well oriented to interact and deprotonate at position C5. MD simulations also describe that the N-hydroxyl group can explore short distances with respect to the heme propionate group (distances ca. 3.0 Å), indicating that these two groups could interact. This is mainly due to a small rotation along the Fe-N bond. This is, however, not preventing the N-hydroxyl group to interact with the catalytic T107 residue, as shown in **Supplementary Figure 18**.

**Supplementary Figure 22. Conformational arrangement of key Fe-N intermediate in PipS active site (PipS-Int2) explored by MD simulations.** Distance between Fe-N nitrene intermediate (PipS-**Int2**) and heme propionate and key N-N bond forming distance explored in four independent MD simulations and considering the two independent active sites from the PipS dimer. Distances are given in angstrom (Å).

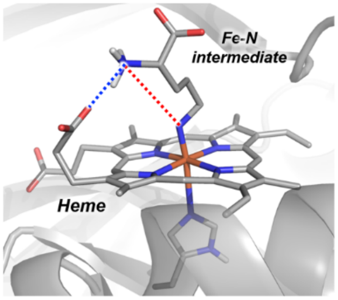

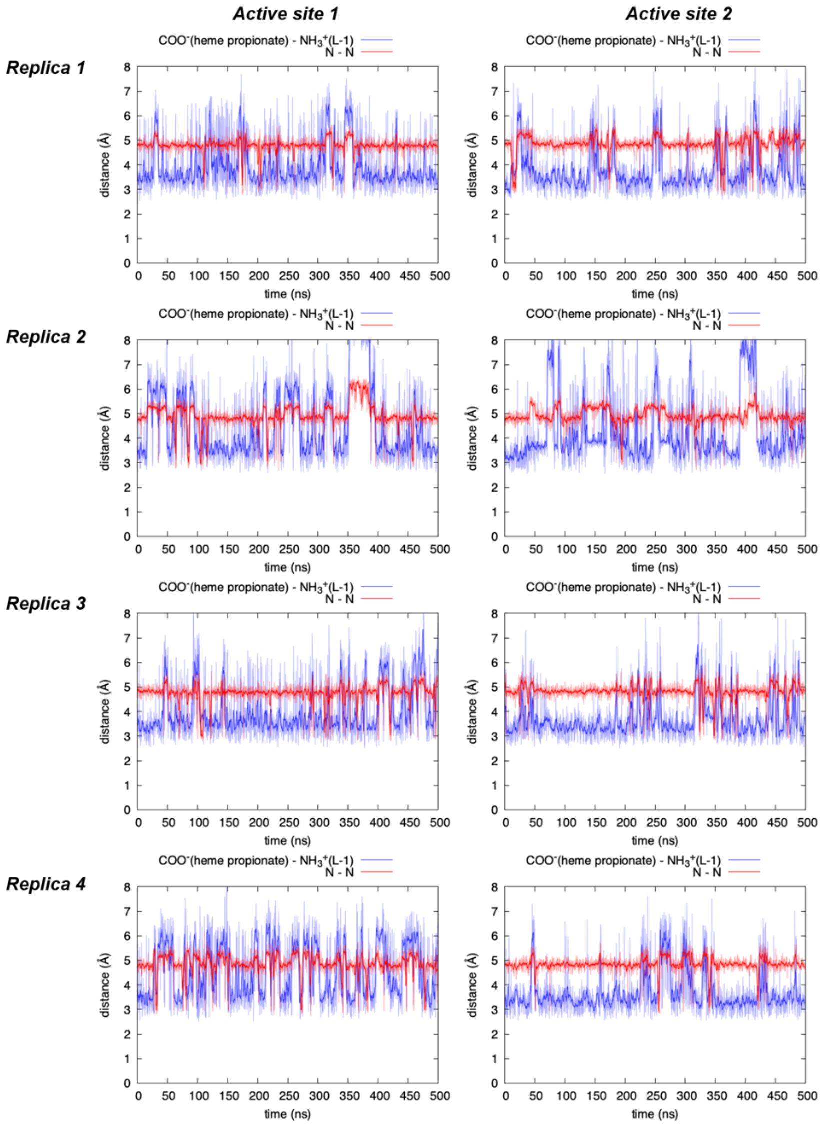

A total of 4 independent replicas were performed starting from the X-ray structure with the L-**1** bound substrate but it was manually changed for the key Fe-N intermediate PipS-**Int2** (see **Figure 3** in the main text for **Int2** description). This implies independent minimizations, heatings and equilibrations during MD preparation protocols prior MD production runs (see Computational Methods), which ensure variability for the initiation of the different independent MD trajectories (production runs). Two different Fe-N intermediates, one at each active site of the PipS dimer, are considered. Production runs were propagated during 500 ns, which represent enough simulation time to capture relevant conformational changes of the intermediate ant the enzyme that might affect to the active site cavities.

MD simulations for the key Fe-N intermediate bound in PipS active site (PipS-**Int2**) were performed to analyze the conformational arrangement and accessible conformations of the intermediate in the catalytic pocket.
Simulations indicate that the α-amino group interacts with the heme propionate (distance ca. 3.0 Å) during most of the simulation time, although larger distances can be explored. These larger distances for the heme propionate - α-amino group interaction allow the α-amino to explore shorter N-N distances (distance ca. 3.0 Å), which are described as potentially reactive conformations for N-N bond formation. A representative snapshot of these potentially reactive conformations was used as starting point for further mechanistic interrogations using QM/MM calculations (see QM/MM calculations in **Supplementary Figure 27**).

**Supplementary Figure 23. Interactions between key Fe-N intermediate (PipS-Int2) and proximal residues in PipS active site explored by MD simulations.** Key distances between Y7, Y99, Y155’, E182, Q183, and K178 polar residues and Fe-N nitrene intermediate (PipS-**Int2**) when formed in PipS, explored in four independent MD simulations and considering the two independent active sites from the PipS dimer. Distances are given in angstrom (Å).

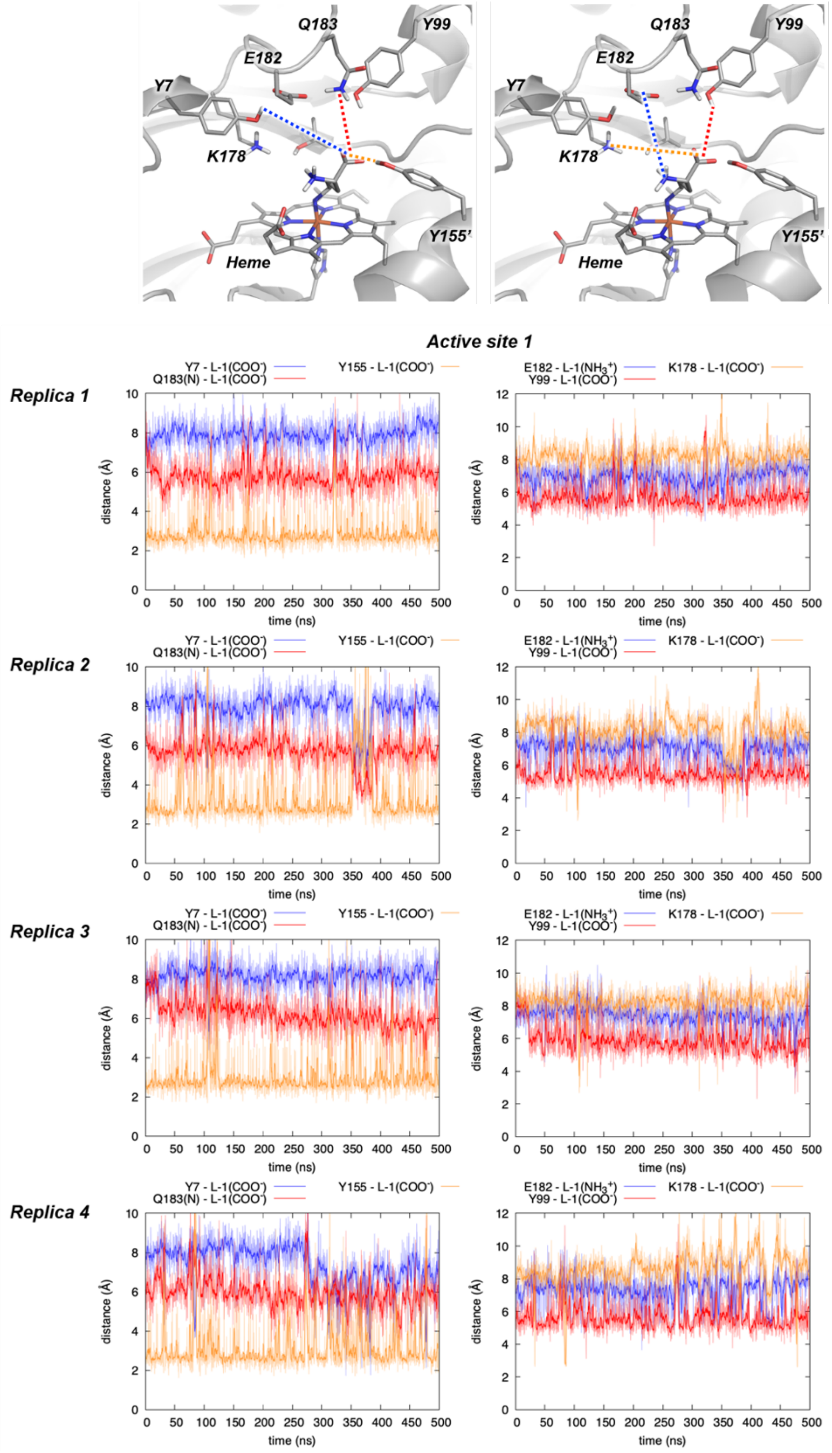

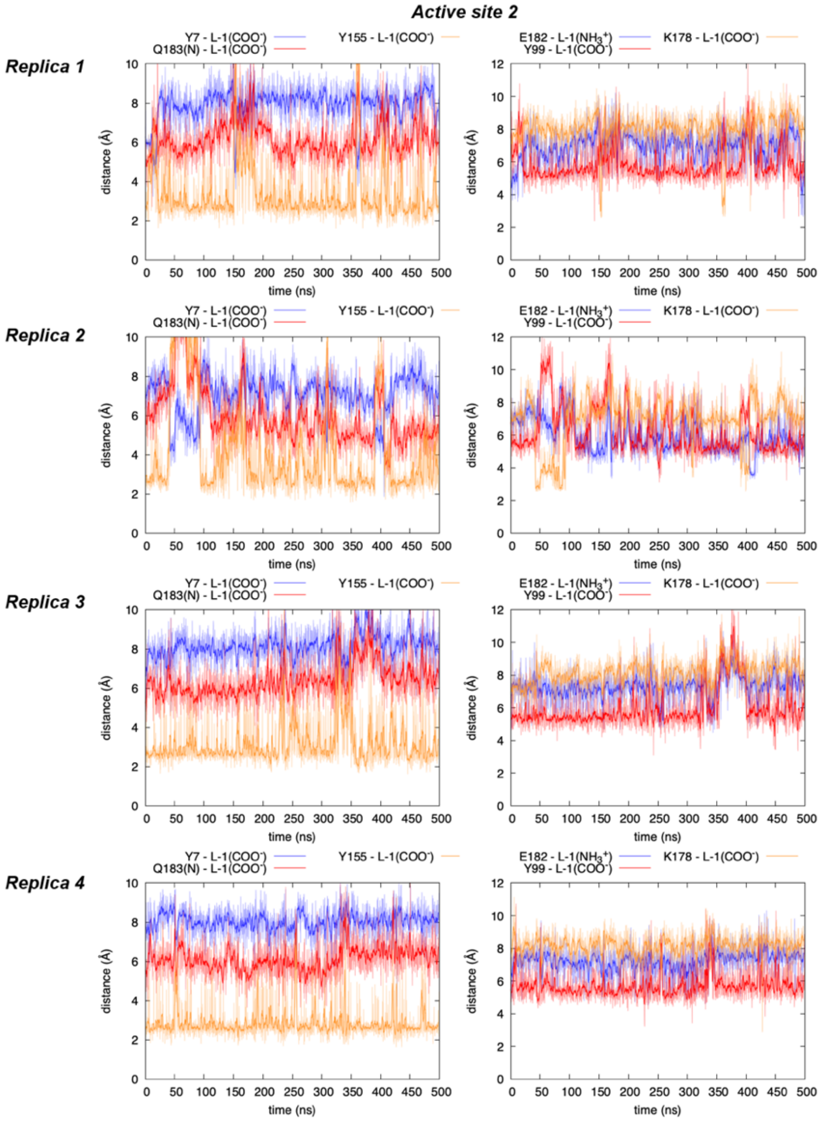

The same MD simulations reported in **Supplementary Figure 22** were used to analyze the principal interactions occurring between key intermediate Fe-N PipS-Int2 and PipS active site residues.

MD simulations showed that the major interaction is occurring between the carboxylate group of the former L-**1** and Y155 via H-bond (distance ca. 3.0 Å), which is maintained during most of the simulation time. This interaction is equivalent to the one observed in the X-ray structure, and in MD simulations with substrate L-**1** bound.

**Supplementary Figure 24. Interactions between proximal residues in distal axial site of Fe-N intermediate (PipS-Int2) complex explored by MD simulations.** Key distances between Y7, Y99, E182, Q183, T107 and K178 polar residues in Fe-N intermediate (PipS-**Int2**) complex are explored in four independent MD simulations and considering the two independent active sites from the PipS dimer. Distances are given in angstrom (Å).

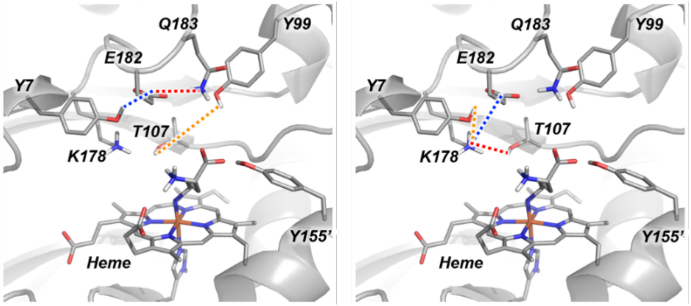

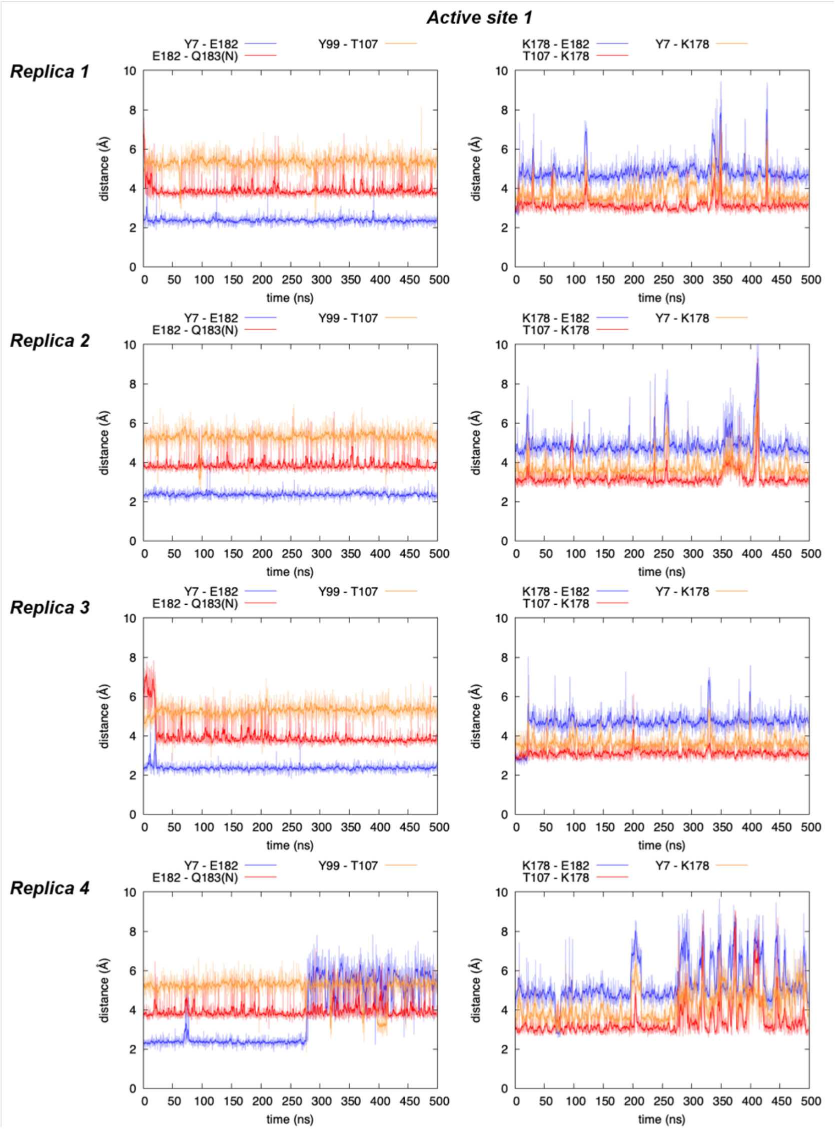

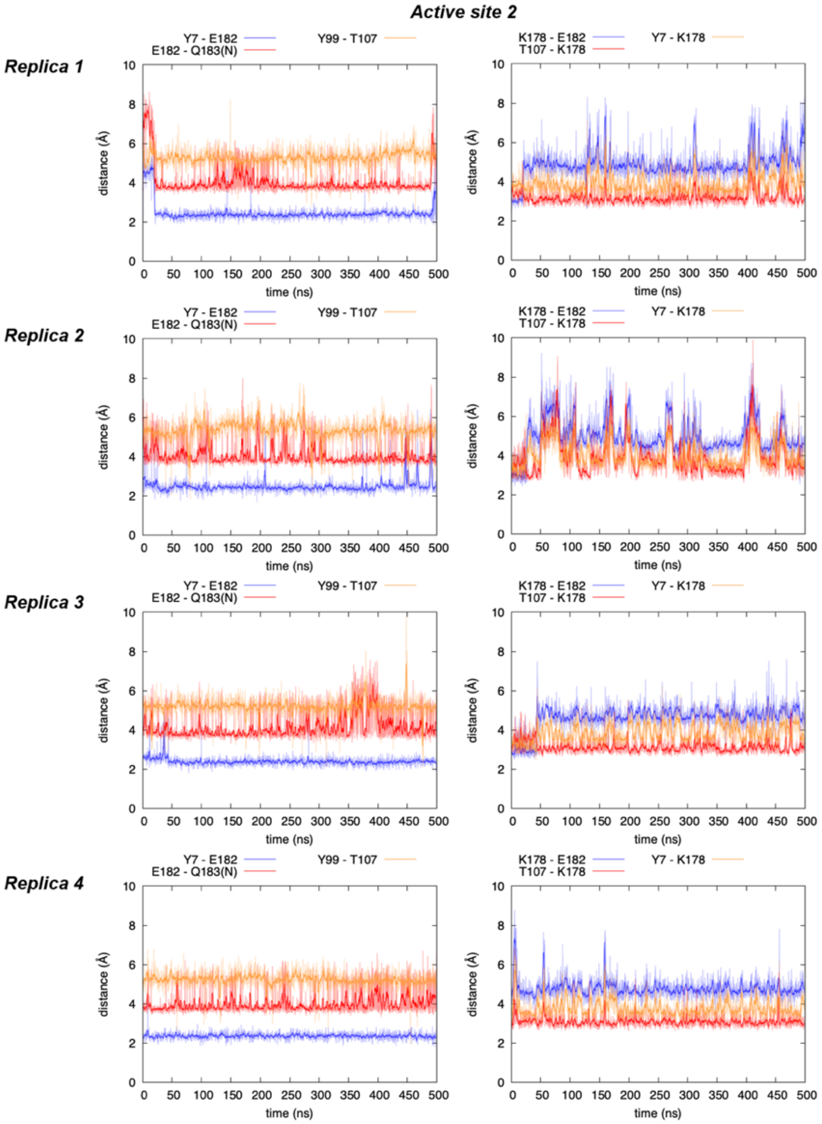

The same MD simulations reported in **Supplementary Figure 22** were used to analyze the hydrogen bond network established between PipS active site residues.

MD simulations described the existence of a persistent H-bond network that is maintained during the whole simulation time, similar to what is observed for substrate L-**1** bound MD simulations reported in **Supplementary Figure 20**. This H-bond network involves the H-bond interactions established between Y7 and E182, and at the same time the interaction established between Y7 and K178, and between E182 and Q183.

As expected, K178 is H-bond interacting with T107 during the whole simulation time.

**Supplementary Figure 25. Interactions between proximal residues and L-1 C5 position in Fe-N intermediate (PipS-Int2) complex explored by MD simulations.** Key distances between polar residues capable of deprotonate Fe-N nitrene intermediate (PipS-**Int2**) at the C5 position are explored in four independent MD simulations and considering the two independent active sites from the PipS dimer. Distances are given in angstrom (Å).

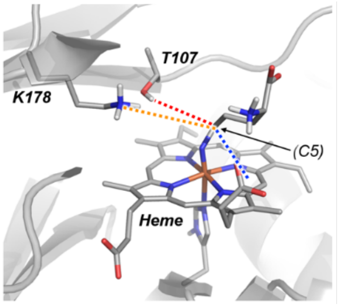

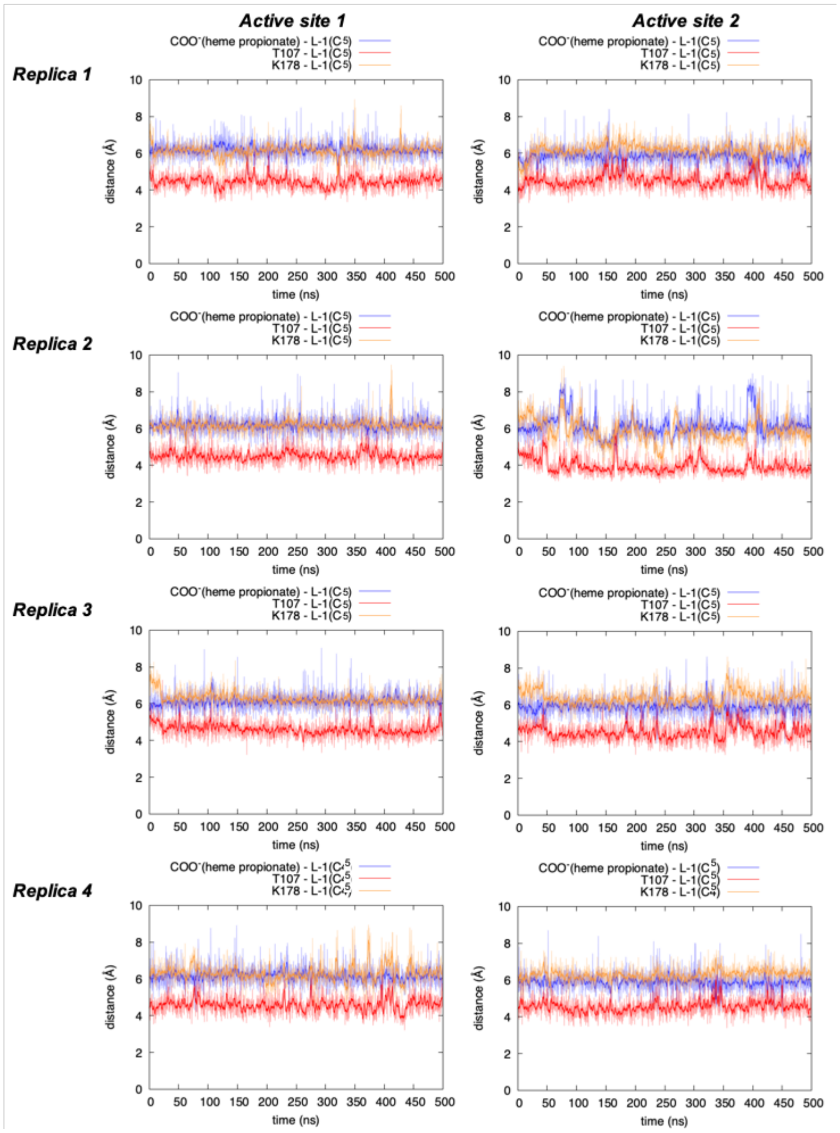

The same MD simulations reported in **Supplementary Figure 22** were used to analyze the potential interactions between the C5 position from key intermediate Fe-N PipS-**Int2** and PipS active site residues that could act as a base to deprotonate it.

MD simulations described that, in general, C5 position stays far from the heme propionate group (distances ca. > 6.0 Å) and T107 (distances ca. > 4.0 Å), suggesting that these are not well oriented to directly interact and deprotonate at position C5.

**Supplementary Figure 26. QM/MM exploration of PipS catalyzed hydroxylamine activation pathways.** QM/MM calculations were carried out starting from a selected representative snapshot extracted from MD simulations with PipS-L-**1** bound complex (**Supplementary Figure 18**).

(**a**) The proposed catalytic cycle of PipS corresponding to the hydroxylamine activation (in purple) and the formation of key nitrene intermediate (in green).

(**b**) Schematic Potential Energy Surface (PES) for the hydroxylamine activation. Energy values were obtained at the (U)B3LYP/Def2TZVP:AmberFF14SB//(U)B3LYP/6-31G(d)+SDD(Fe) :AmberFF14SB level, with the same MM parameters used in MD simulations (see SI, computational details). Relative Gibbs (ΔG) and electronic (ΔE) energies are reported for all electronic states. All energies are referred considering PipS-L-**1** (Q) structure as zero.

(**c**) Computed relative stabilities in terms of electronic QM energy ($\Delta E_{QM}$), electronic QM/MM energy (ΔE), enthalpy (ΔH), and Gibbs energy (ΔG) for the different species. All energies are referred considering PipS-L-**1** (Q) structure as zero.

(**d**) Optimized QM/MM structures for the different species involved in the studied reaction pathways. All atoms included in the QM-region are shown in ball and stick representation.

All energies and distances are given in kcal·mol$^{-1}$ and angstrom (Å), respectively.

**(a)**

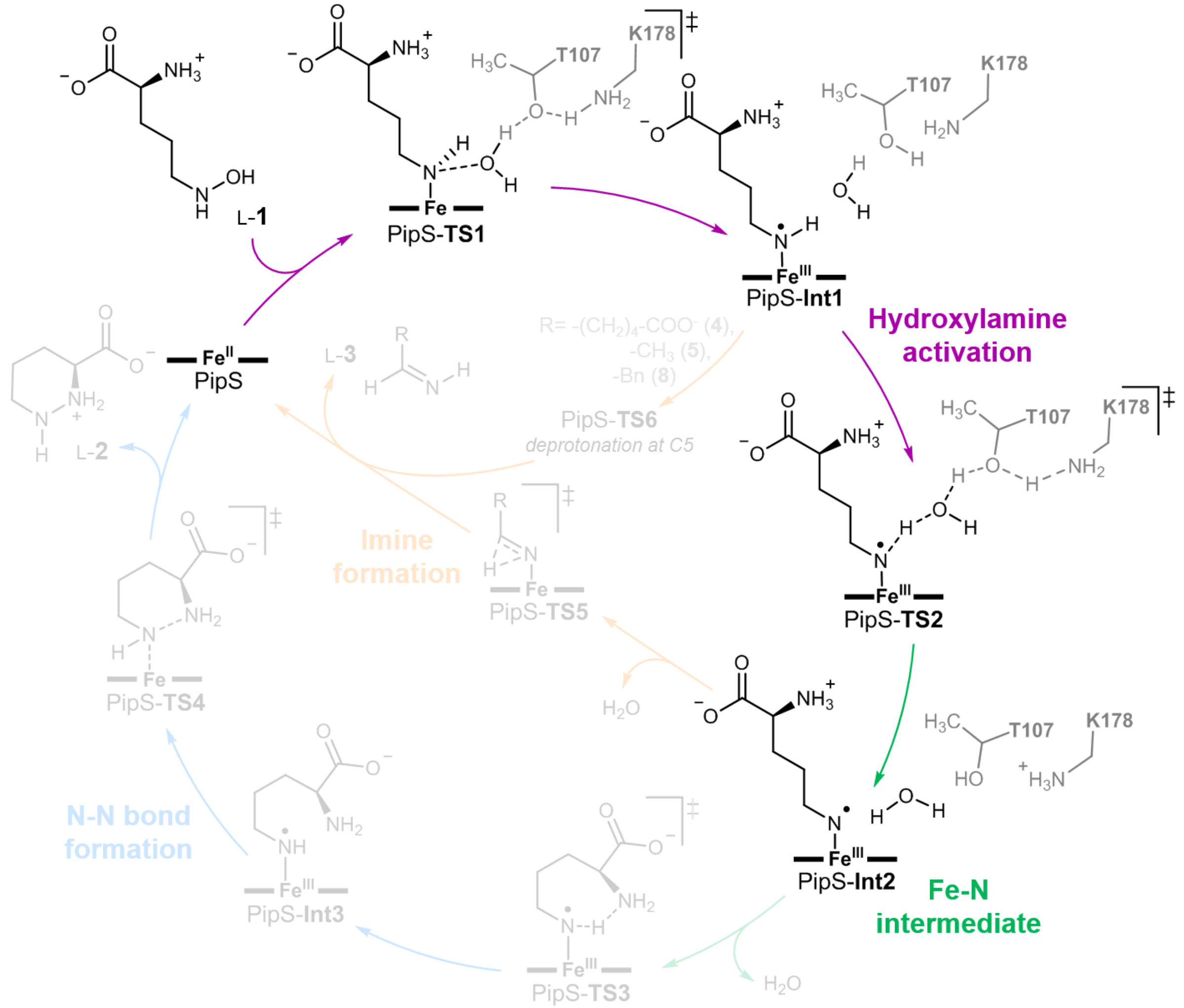

L-1
PipS-TS1
PipS-Int1
Hydroxylamine activation
PipS-TS2
PipS-Int2
Fe-N intermediate
PipS-TS3
PipS-Int3
N-N bond formation
PipS-TS4
L-2
PipS
PipS-TS5
Imine formation
PipS-TS6
deprotonation at C5
L-3
T107
K178

**(b)**

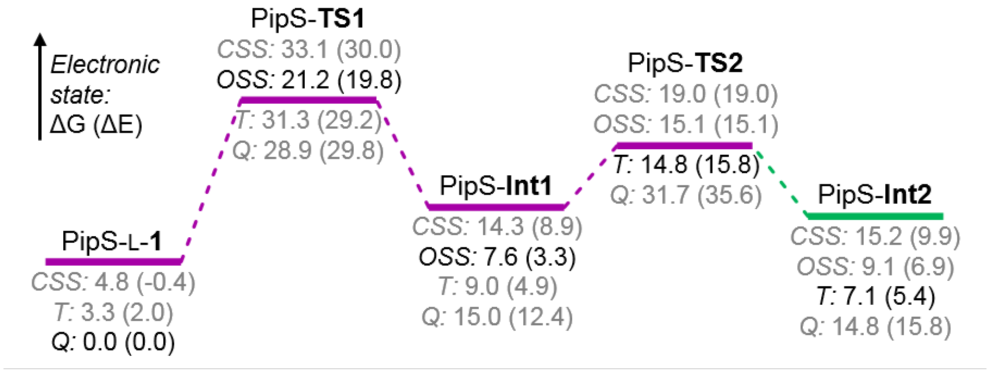

Electronic state: ΔG (ΔE)
PipS-L-1
CSS: 4.8 (-0.4)
T: 3.3 (2.0)
Q: 0.0 (0.0)
PipS-TS1
CSS: 33.1 (30.0)
OSS: 21.2 (19.8)
T: 31.3 (29.2)
Q: 28.9 (29.8)
PipS-Int1
CSS: 14.3 (8.9)
OSS: 7.6 (3.3)
T: 9.0 (4.9)
Q: 15.0 (12.4)
PipS-TS2
CSS: 19.0 (19.0)
OSS: 15.1 (15.1)
T: 14.8 (15.8)
Q: 31.7 (35.6)
PipS-Int2
CSS: 15.2 (9.9)
OSS: 9.1 (6.9)
T: 7.1 (5.4)
Q: 14.8 (15.8)

**(c)**

| Structure | Electronic State | $\Delta E_{QM}$ | ΔE | ΔH | ΔG |
|---|---|---|---|---|---|
| PipS-L-**1** | close-shell singlet (CSS) | -3.3 | -0.4 | 0.8 | 4.8 |
| | triplet (T) | 1.2 | 2.0 | 2.7 | 3.3 |
| | quintet (Q) | 0.0 | 0.0 | 0.0 | 0.0 |
| PipS-**TS1** | close-shell singlet (CSS) | 25.0 | 30.0 | 26.4 | 33.1 |
| | open-shell singlet (OSS) | 18.6 | 19.8 | 15.7 | 21.2 |
| | triplet (T) | 27.1 | 29.2 | 27.0 | 31.3 |
| | quintet (Q) | 28.5 | 29.8 | 25.8 | 28.9 |
| PipS-**Int1** | close-shell singlet (CSS) | -1.3 | 8.9 | 8.9 | 14.3 |
| | open-shell singlet (OSS) | -8.7 | 3.3 | 2.8 | 7.6 |
| | triplet (T) | -3.5 | 4.9 | 4.4 | 9.0 |
| | quintet (Q) | 2.6 | 12.4 | 11.9 | 15.0 |
| PipS-**TS2** | close-shell singlet (CSS) | 9.1 | 19.0 | 13.0 | 19.0 |
| | open-shell singlet (OSS) | 6.9 | 15.1 | 9.2 | 15.1 |
| | triplet (T) | 7.5 | 15.8 | 9.6 | 14.8 |
| | quintet (Q) | 27.0 | 35.6 | 28.3 | 31.7 |
| PipS-**Int2** | close-shell singlet (CSS) | -2.4 | 9.9 | 9.8 | 15.2 |
| | open-shell singlet (OSS) | -4.0 | 6.9 | 5.9 | 9.1 |
| | triplet (T) | -5.8 | 5.4 | 4.6 | 7.1 |
| | quintet (Q) | 4.8 | 15.8 | 14.1 | 14.8 |

**(d)**

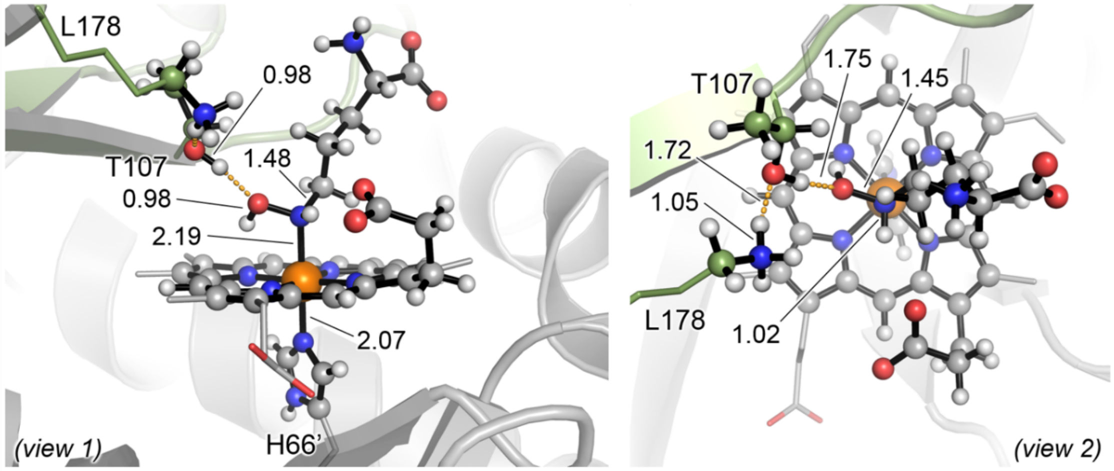

PipS-L-**1** (CSS)
ΔG = 4.8 (ΔE‡ = -0.4)

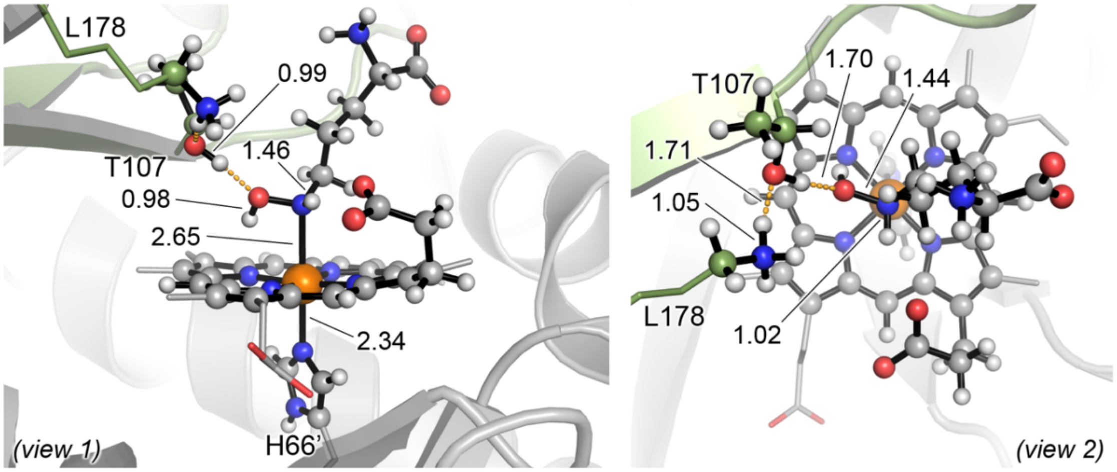

PipS-L-**1** (T)
ΔG‡ = 3.3 (ΔE‡ = 2.0)

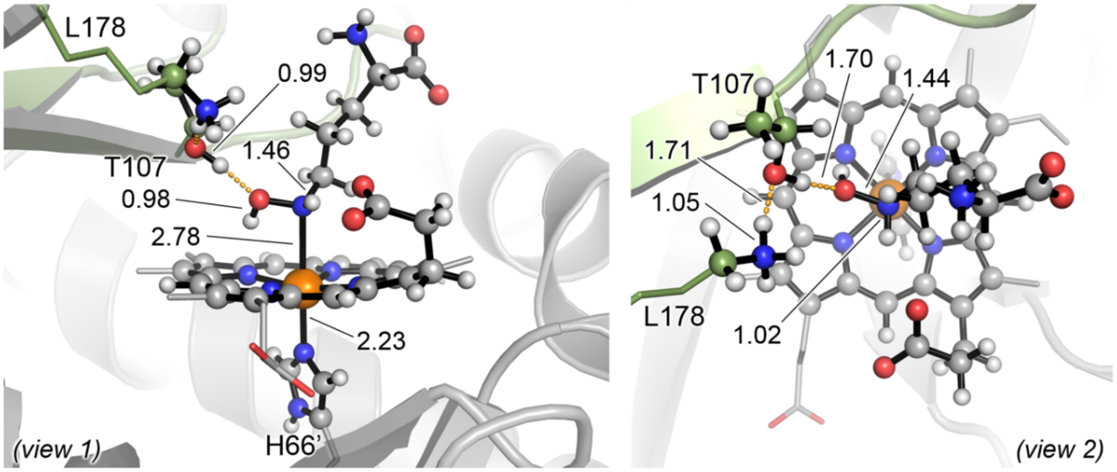

PipS-L-**1** (Q)
ΔG‡ = 0.0 (ΔE‡ = 0.0)

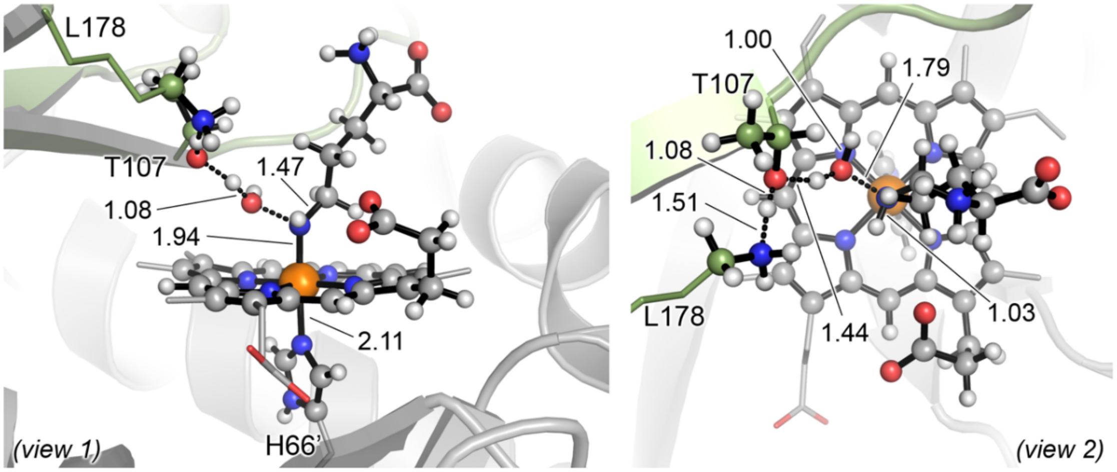

PipS-**TS1** (CSS)
ΔG‡ = 28.3 (ΔE‡ = 30.5)

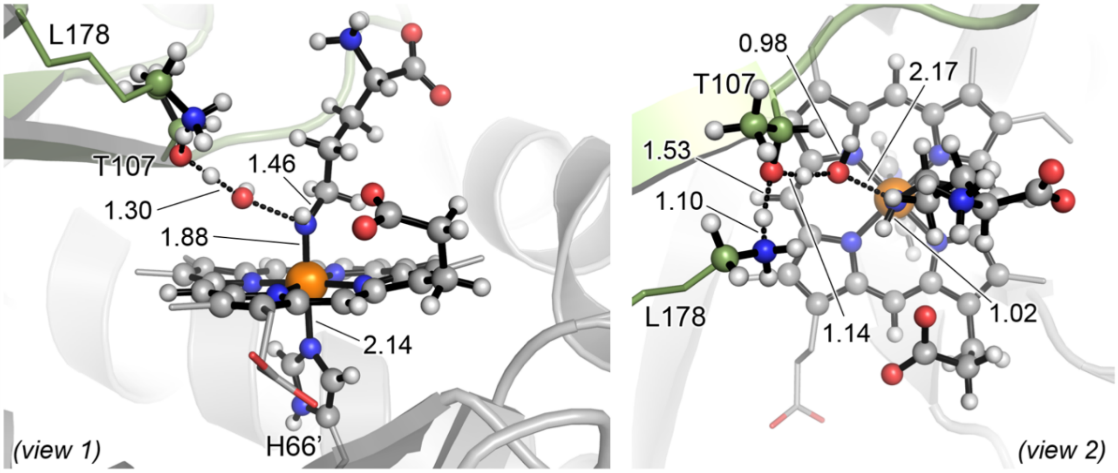

PipS-**TS1** (OSS)
ΔG‡ = 16.4 (ΔE‡ = 20.3)

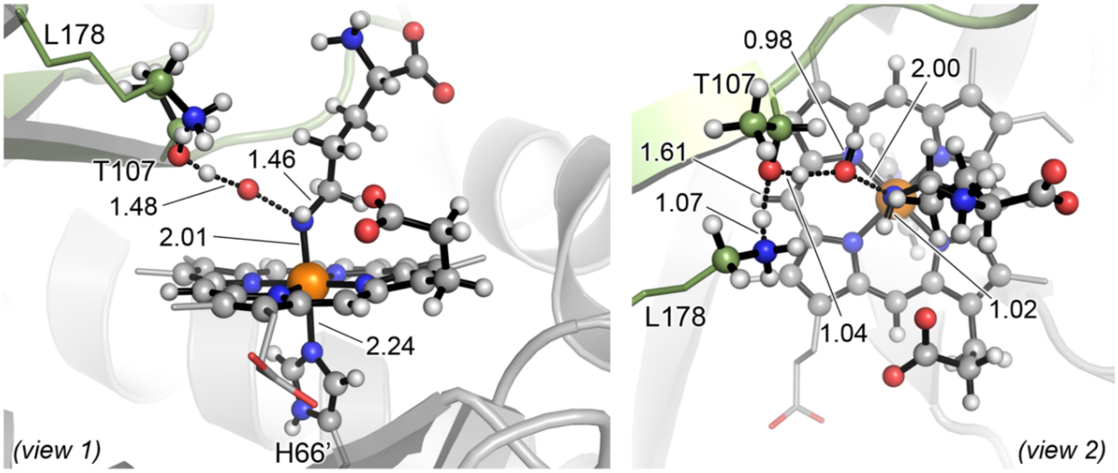

PipS-**TS1** (T)
ΔG‡ = 28.0 (ΔE‡ = 27.3)

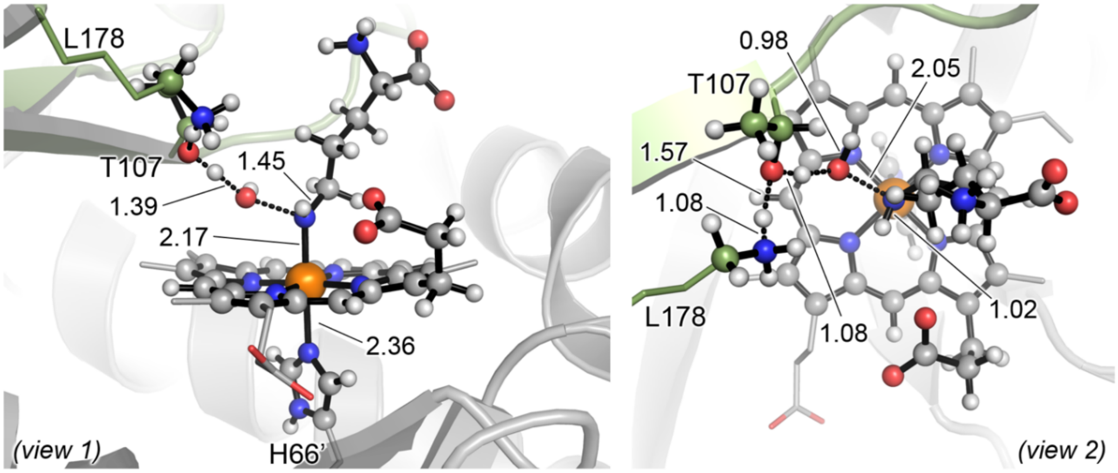

PipS-**TS1** (Q)
ΔG‡ = 28.9 (ΔE‡ = 29.8)

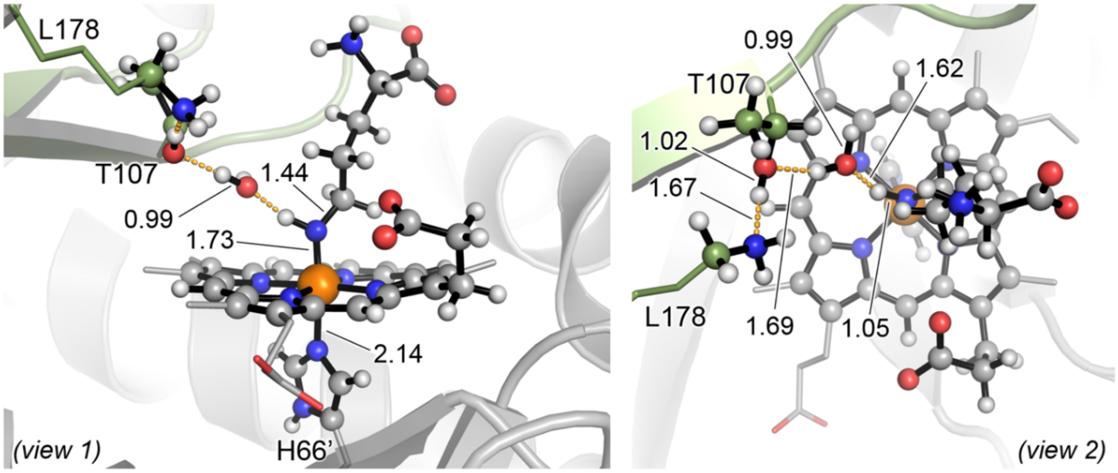

PipS-**Int1** (CSS)
$\Delta G_r = 9.5$ ($\Delta E_r = 9.3$)

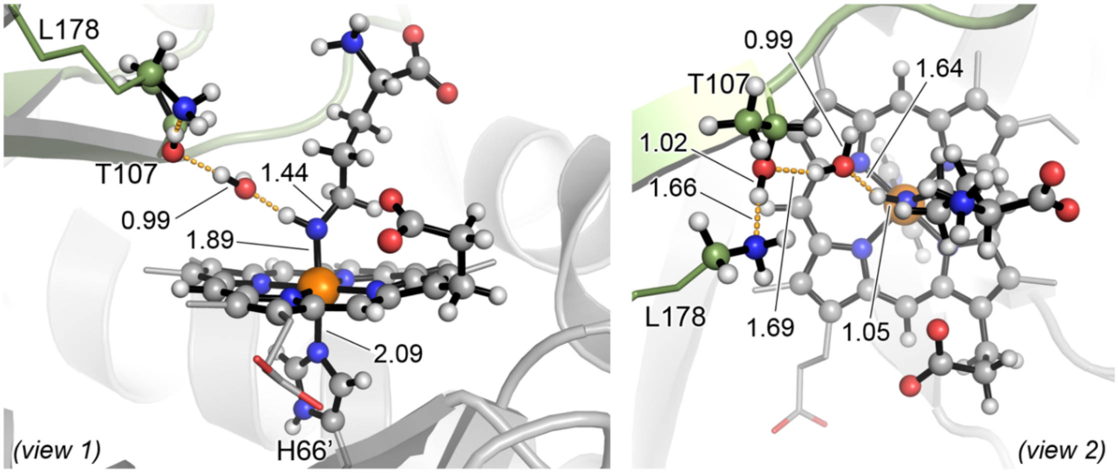

PipS-**Int1** (OSS)
$\Delta G_r = 2.8$ ($\Delta E_r = 3.7$)

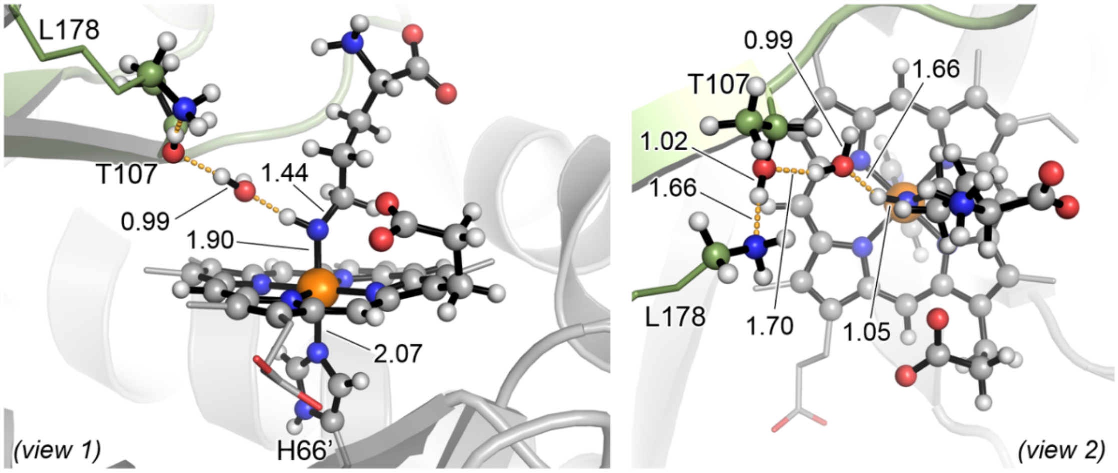

PipS-**Int1** (T)
$\Delta G_r = 5.7$ ($\Delta E_r = 2.9$)

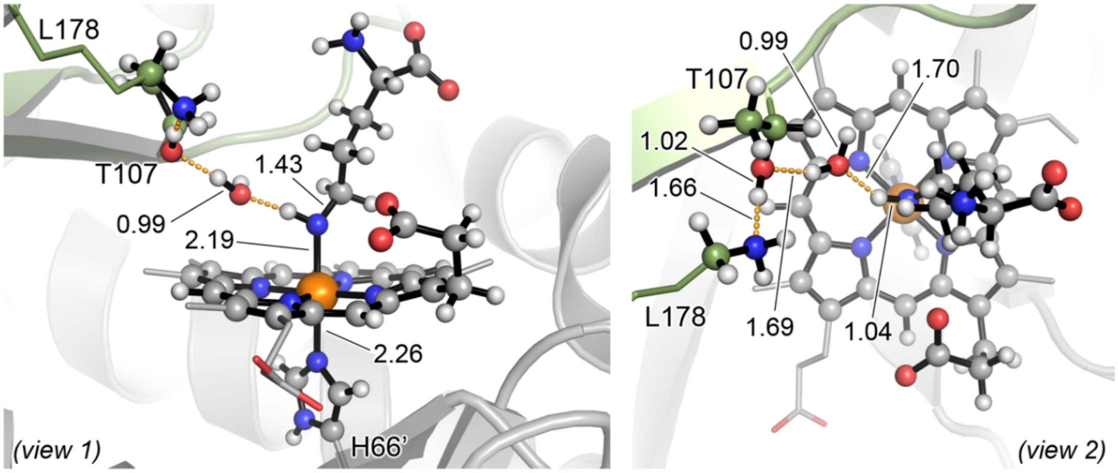

PipS-**Int1** (Q)
$\Delta G_r$ = 15.0 ($\Delta E_r$ = 12.4)

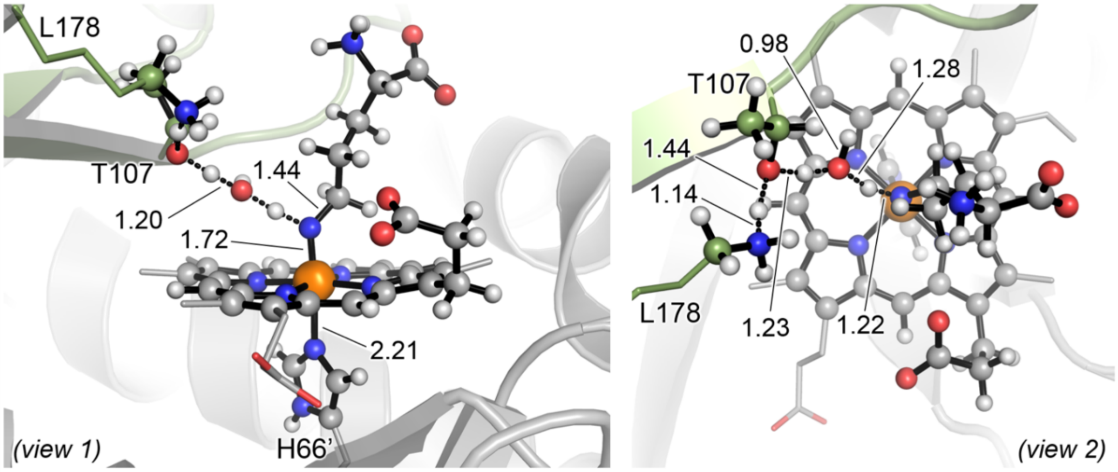

PipS-**TS2** (CSS)
$\Delta G^‡$ = 4.7 ($\Delta E^‡$ = 10.2)

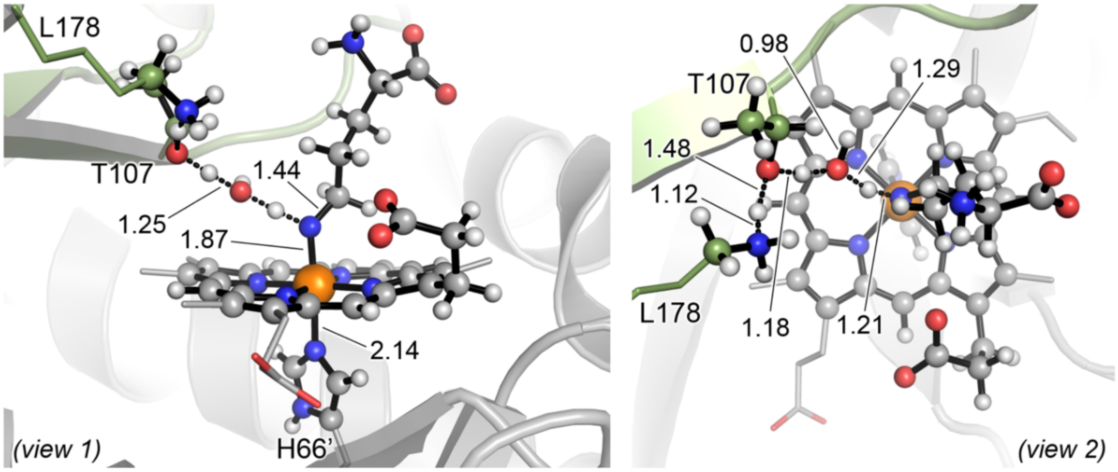

PipS-**TS2** (OSS)
$\Delta G^‡$ = 7.6 ($\Delta E^‡$ = 11.9)

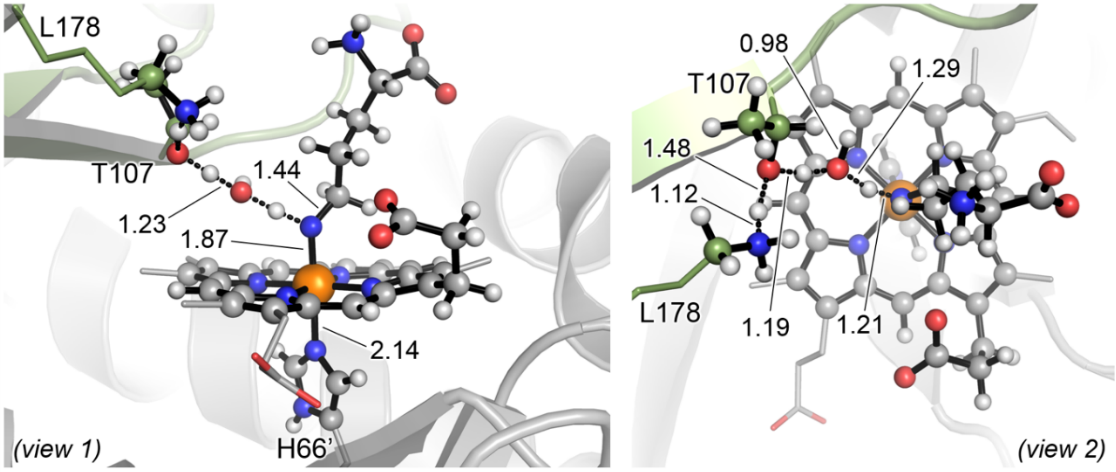

PipS-**TS2** (T)
ΔG‡ = 5.8 (ΔE‡ = 10.9)

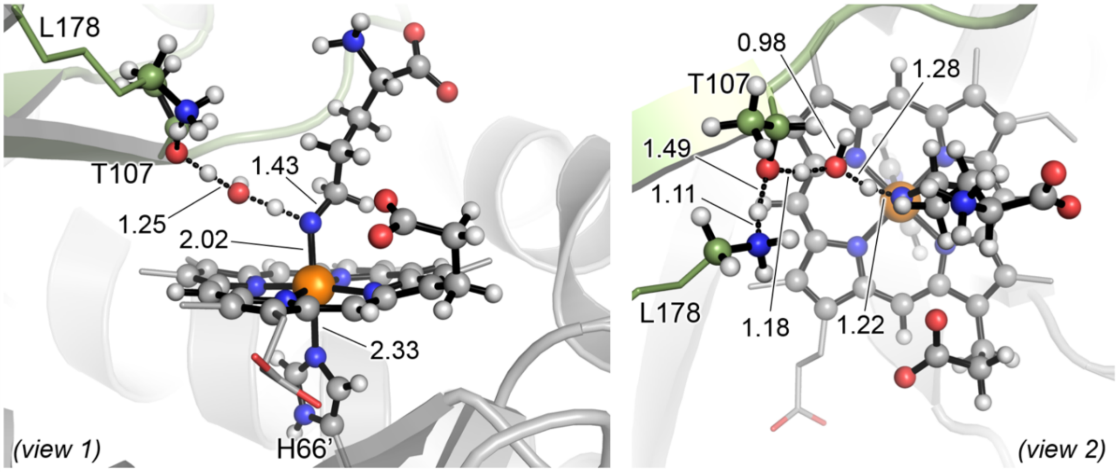

PipS-**TS2** (Q)
ΔG‡ = 16.7 (ΔE‡ = 23.1)

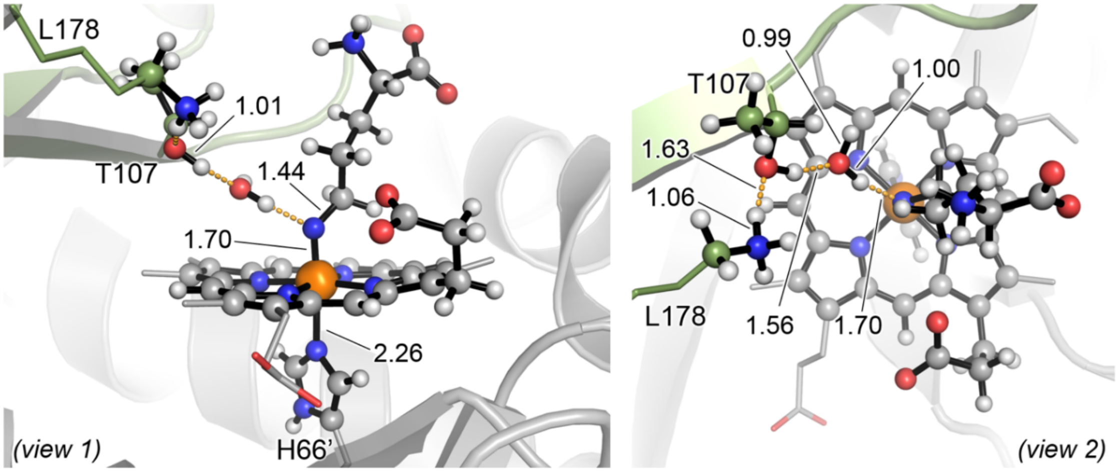

PipS-**Int2** (CSS)
$\Delta G_r$ = 0.8 ($\Delta E_r$ = 1.0)

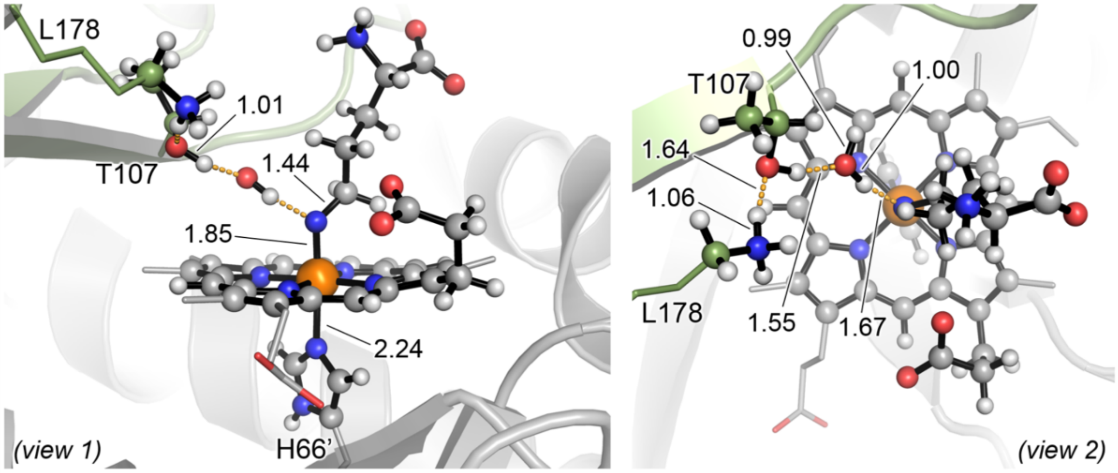

PipS-**Int2** (OSS)
$\Delta G_r = 1.6$ ($\Delta E_r = 3.6$)

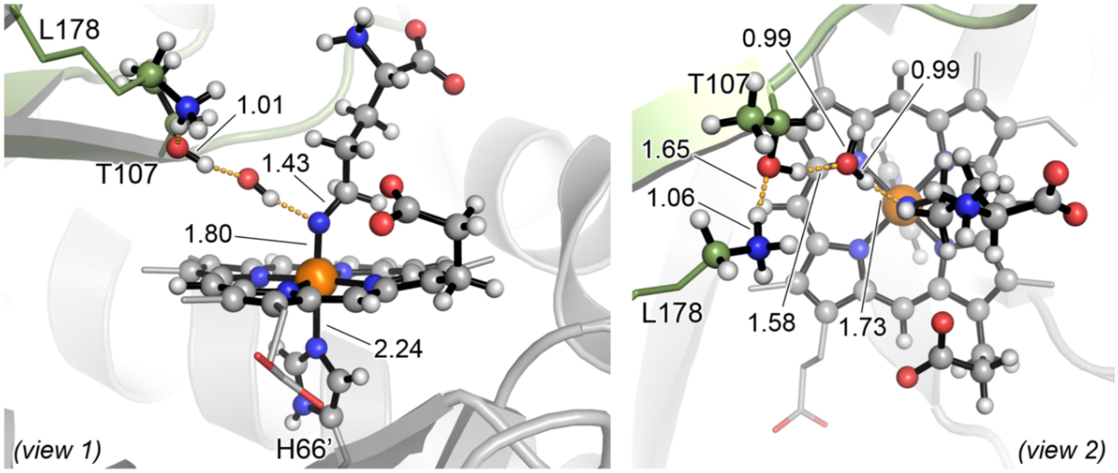

PipS-**Int2** (T)
$\Delta G_r = -1.9$ ($\Delta E_r = 0.5$)

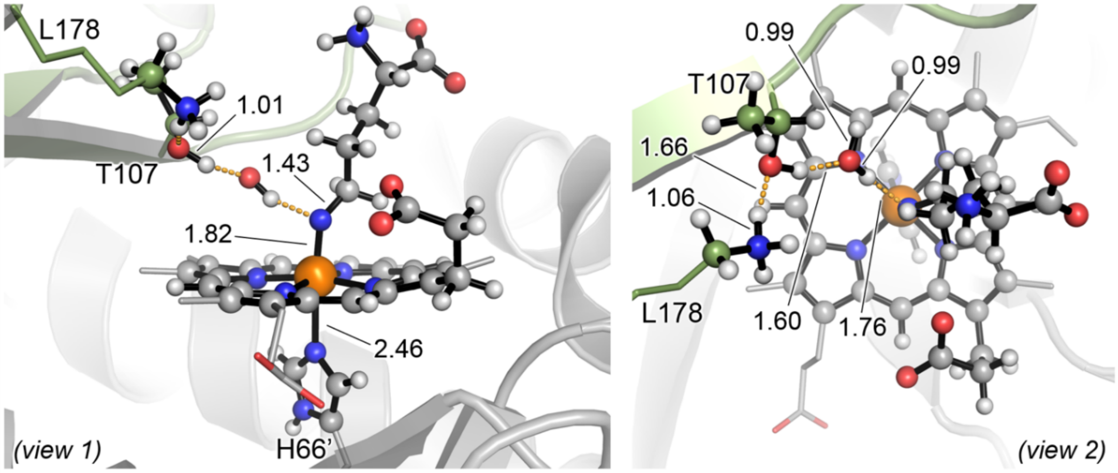

PipS-**Int2** (Q)
$\Delta G_r = -0.1$ ($\Delta E_r = 3.4$)

QM/MM calculations showed that PipS-L-**1** in its quintet (Q) electronic state is slightly more stable than triplet (T) by ca. 3 kcal·mol$^{-1}$. In the optimized PipS-L-**1** structures the catalytic dyad formed by K178 and T107 is well preorganized to protonate the hydroxylamine group (**Supplementary Figures 18** and **26b**). The corresponding transition state, PipS-**TS1**, has been optimized where K178-T107 catalytic dyad protonates the hydroxylamine group to release a water molecule to form a protonated Fe-nitrenoid species (PipS-**Int1**). The lowest in energy PipS-**TS1** is the open-shell singlet (OSS), with a free energy barrier of 21.1 kcal·mol$^{-1}$.

After the water molecule is released, the covalent radical intermediate PipS-**Int1** is formed. In this optimized QM/MM structure, a hydrogen bond network involving the two catalytic residues, the newly released water molecule and the protonated nitrenoid is shown to be conserved. PipS-**Int1** is 7.6 kcal·mol$^{-1}$ above the reactant PipS-L-**1** (Q) for the lowest in energy OSS electronic state.

In order to restore the original protonation state of the catalytic dyad, the nitrenoid intermediate PipS-**Int1** can be deprotonated by the released water molecule that transfers back a proton to the catalytic T107-K178 pair through PipS-**TS2** transition state. This leads to the formation of a nitrene intermediate PipS-**Int2**.This transition state is highly favored due to the preorganization of both the catalytic dyad and the water molecule, thanks to the H-bonding network that is established in PipS-**Int1** after hydroxylamine activation. PipS-**TS2** in its triplet electronic state has the lowest activation barrier $\Delta G^{\ddagger}$ = 7.3 kcal·mol$^{-1}$, with a significant imaginary frequency of 1220*i* cm$^{-1}$. The nitrene intermediate generated, PipS-**Int2**, has a triplet ground state (T) and it is $\Delta G$ = 7.1 kcal·mol$^{-1}$ less stable than PipS-L-**1** (Q). Nitrene intermediate PipS-**Int2** corresponds to the key reactive species from which it is proposed that both piperazic acid (L-**2**) and imine product (L-**3**) are formed.

To sum up, QM/MM calculations indicate that the activation of the hydroxylamine group to form the key nitrene radical intermediate is energetically accessible through a rate-limiting PipS-**TS1**. First the catalytic dyad protonates the hydroxylamine moiety of the substrate, releasing a water molecule and generating a covalent protonated nitrenoid species. Later, the same H-bond network is used to restore the protonation state of the catalytic dyad, by deprotonating the nitrenoid, thus leading to the key nitrene intermediate PipS-**Int2**.

**Supplementary Figure 27. QM/MM exploration of PipS catalyzed N-N bond formation and imine pathways from nitrene intermediate.** QM/MM calculations were carried out starting from a selected representative snapshot extracted from MD simulations with PipS-**Int2** bound complex (**Supplementary Figure 22**).

(**a**) The proposed catalytic steps for PipS-catalyzed N-N bond formation (in blue) and the imine formation pathway (in orange).
(**b**) Schematic Potential Energy Surface (PES) for the N-N bond formation and imine formation pathways. Energy values were obtained at the (U)B3LYP/Def2TZVP:AmberFF14SB// (U)B3LYP/6-31G(d)+SDD(Fe):AmberFF14SB level, with the same MM parameters used in MD simulations (see SI, computational details). Relative Gibbs (ΔG) and electronic (ΔE) energies are reported for all electronic states. All energies are referred considering PipS-**Int2** (T) structure as zero.
(**c**) Computed relative stabilities in terms of electronic QM energy ($\Delta E_{QM}$), electronic QM/MM energy (ΔE), enthalpy (ΔH), and Gibbs energy (ΔG) for the different species. All energies are referred considering PipS-**Int2** (T) structure as zero.
(**d**) Optimized QM/MM structures for the different species involved in the N-N bond formation pathway. All atoms included in the QM-region are shown in ball and stick representation.
(**e**) Relaxed PES scan along the N-N bond forming coordinate from nitrenoid intermediate PipS-**Int3** at (U)B3LYP/6-31G(d)+SDD(Fe):AmberFF14SB and closed shell singlet (CSS) electronic state (using an electrostatic embedding).
(**f**) Optimized QM/MM structures for the different species involved in the imine pathway. All atoms included in the QM-region are shown in ball and stick representation.

All energies and distances are given in $kcal \cdot mol^{-1}$ and angstrom (Å), respectively.

**(a)**

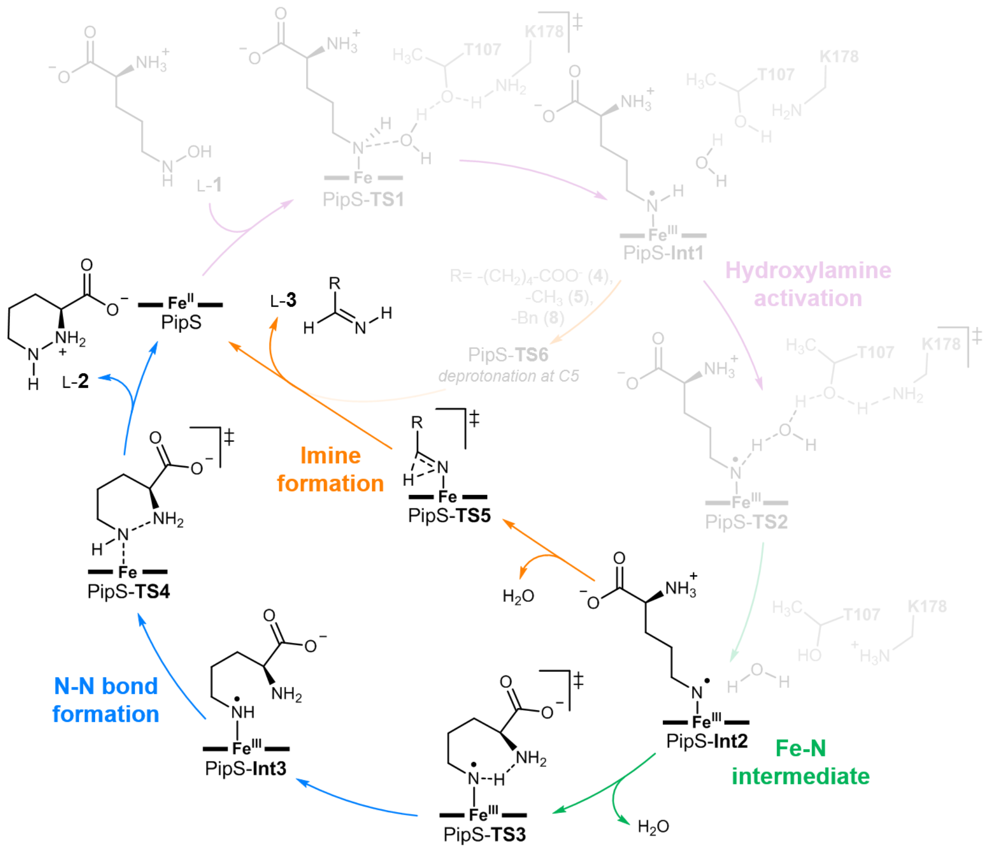

L-1
PipS-TS1
PipS-Int1
Hydroxylamine activation
PipS-TS2
PipS-Int2
Fe-N intermediate
$H_2O$
PipS-TS3
PipS-Int3
N-N bond formation
PipS-TS4
L-2
PipS
PipS-TS5
Imine formation
L-3
R= -(CH2)4-COO- (4), -CH3 (5), -Bn (8)
PipS-TS6
deprotonation at C5
T107
K178

**(b)**

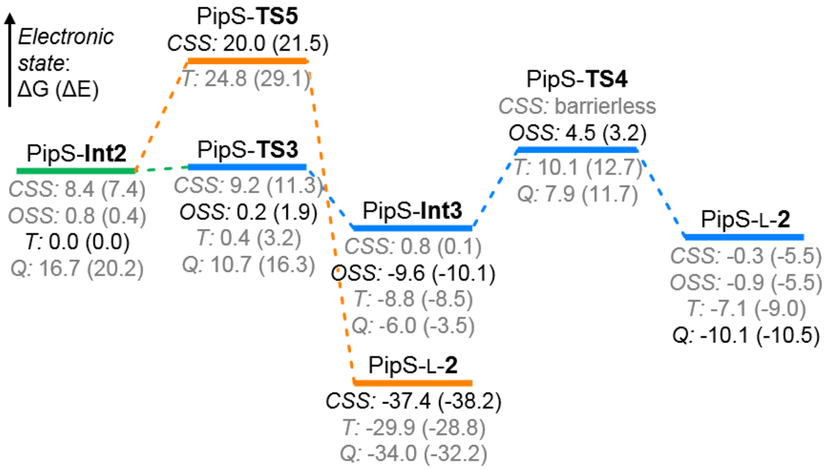

Electronic state: ΔG (ΔE)
PipS-Int2
CSS: 8.4 (7.4)
OSS: 0.8 (0.4)
T: 0.0 (0.0)
Q: 16.7 (20.2)
PipS-TS5
CSS: 20.0 (21.5)
T: 24.8 (29.1)
PipS-TS3
CSS: 9.2 (11.3)
OSS: 0.2 (1.9)
T: 0.4 (3.2)
Q: 10.7 (16.3)
PipS-Int3
CSS: 0.8 (0.1)
OSS: -9.6 (-10.1)
T: -8.8 (-8.5)
Q: -6.0 (-3.5)
PipS-L-2
CSS: -37.4 (-38.2)
T: -29.9 (-28.8)
Q: -34.0 (-32.2)
PipS-TS4
CSS: barrierless
OSS: 4.5 (3.2)
T: 10.1 (12.7)
Q: 7.9 (11.7)
PipS-L-2
CSS: -0.3 (-5.5)
OSS: -0.9 (-5.5)
T: -7.1 (-9.0)
Q: -10.1 (-10.5)

**(c)**

| Structure | Electronic State | ΔEQM | ΔE | ΔH | ΔG |
|---|---|---|---|---|---|
| PipS-**Int2** | close-shell singlet (CSS) | 7.7 | 7.4 | 7.5 | 8.4 |
| | open-shell singlet (OSS) | 0.6 | 0.4 | 0.1 | 0.8 |
| | triplet (T) | 0.0 | 0.0 | 0.0 | 0.0 |
| | quintet (Q) | 21.8 | 20.2 | 19.0 | 16.7 |
| PipS-**TS3** | close-shell singlet (CSS) | 10.5 | 11.3 | 8.2 | 9.2 |
| | open-shell singlet (OSS) | -0.7 | 1.9 | -1.2 | 0.2 |
| | triplet (T) | 0.3 | 3.2 | 0.0 | 0.4 |
| | quintet (Q) | 12.4 | 16.3 | 11.7 | 10.7 |
| PipS-**Int3** | close-shell singlet (CSS) | 50.1 | 0.1 | -1.0 | 0.8 |
| | open-shell singlet (OSS) | 40.4 | -10.1 | -11.4 | -9.6 |
| | triplet (T) | 42.0 | -8.5 | -9.8 | -8.8 |
| | quintet (Q) | 44.9 | -3.5 | -5.4 | -6.0 |
| PipS-**TS4** | close-shell singlet (CSS) | | *barrierless | | |
| | open-shell singlet (OSS) | 73.7 | 3.2 | 2.8 | 4.5 |
| | triplet (T) | 84.9 | 12.7 | 11.2 | 10.1 |
| | quintet (Q) | 83.9 | 11.7 | 9.3 | 7.9 |
| PipS-L-**2** | close-shell singlet (CSS) | 56.4 | -5.5 | -3.2 | -0.3 |
| | open-shell singlet (OSS) | 56.6 | -5.5 | -3.3 | -0.9 |
| | triplet (T) | 56.4 | -9.0 | -7.3 | -7.1 |
| | quintet (Q) | 54.9 | -10.5 | -9.5 | -10.1 |
| PipS-**TS5a** | close-shell singlet (CSS) | 22.1 | 21.5 | 18.5 | 20.0 |
| | triplet (T) | 32.2 | 29.1 | 25.6 | 24.8 |
| PipS-**TS5b** | close-shell singlet (CSS) | 22.0 | 31.9 | 28.5 | 29.5 |
| PipS-L-**3** | close-shell singlet (CSS) | -8.5 | -38.2 | -39.0 | -37.4 |
| | triplet (T) | 1.6 | -28.8 | -30.2 | -29.9 |
| | quintet (Q) | -2.0 | -32.2 | -34.0 | -34.0 |

**(d)**

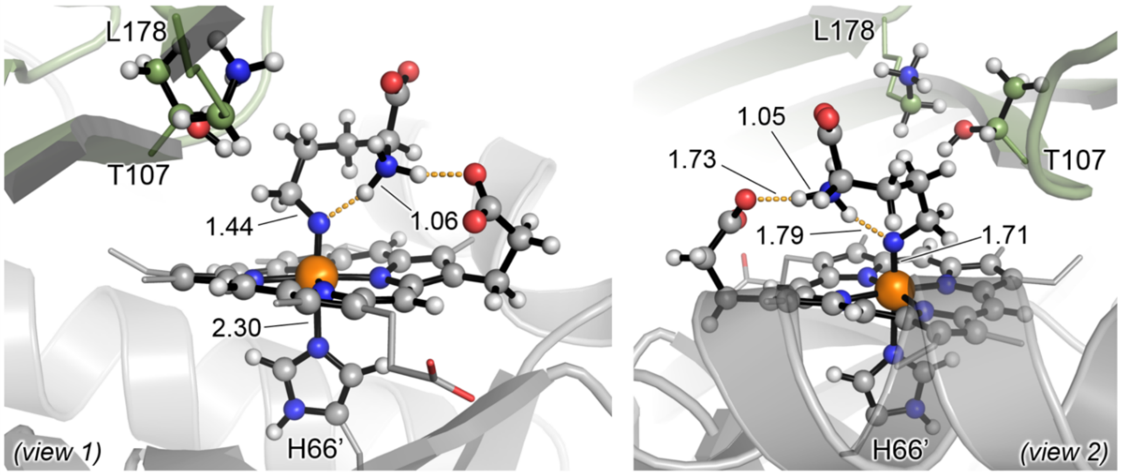

PipS-**Int2** (CSS)
ΔG = 8.4 (ΔE = 7.4)

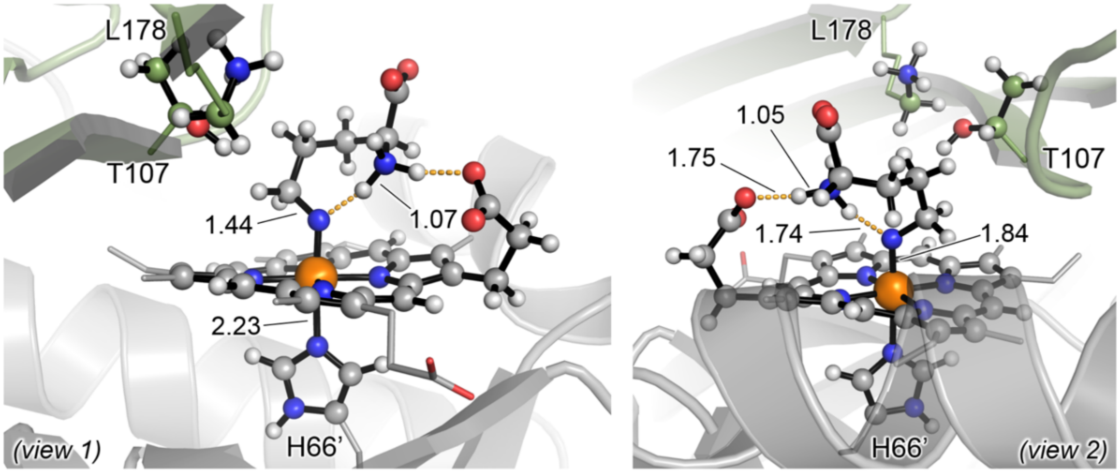

PipS-**Int2** (OSS)
ΔG = 0.8 (ΔE = 0.4)

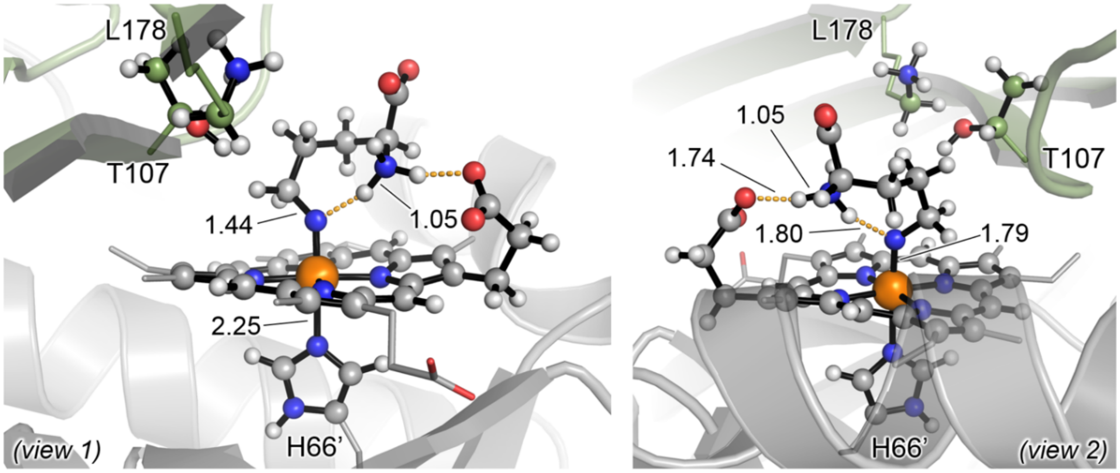

PipS-**Int2** (T)
ΔG = 0.0 (ΔE = 0.0)

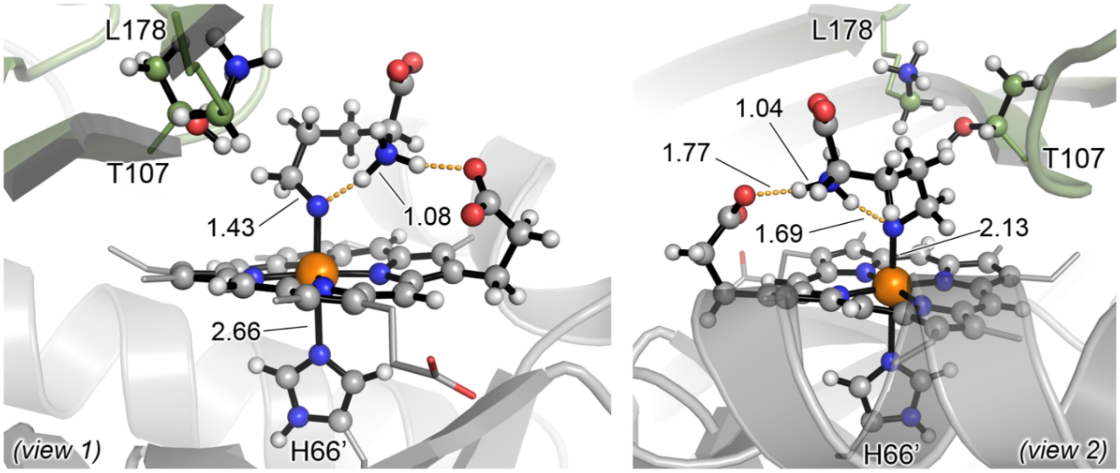

PipS-**Int2** (Q)
ΔG = 16.7 (ΔE = 20.2)

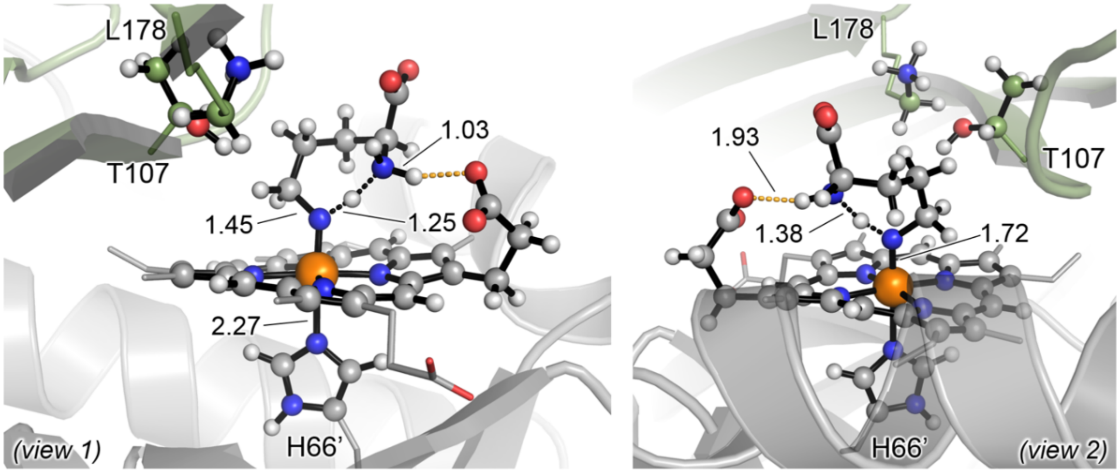

PipS-**TS3** (CSS)
ΔG‡ = 0.8 (ΔE‡ = 2.8)

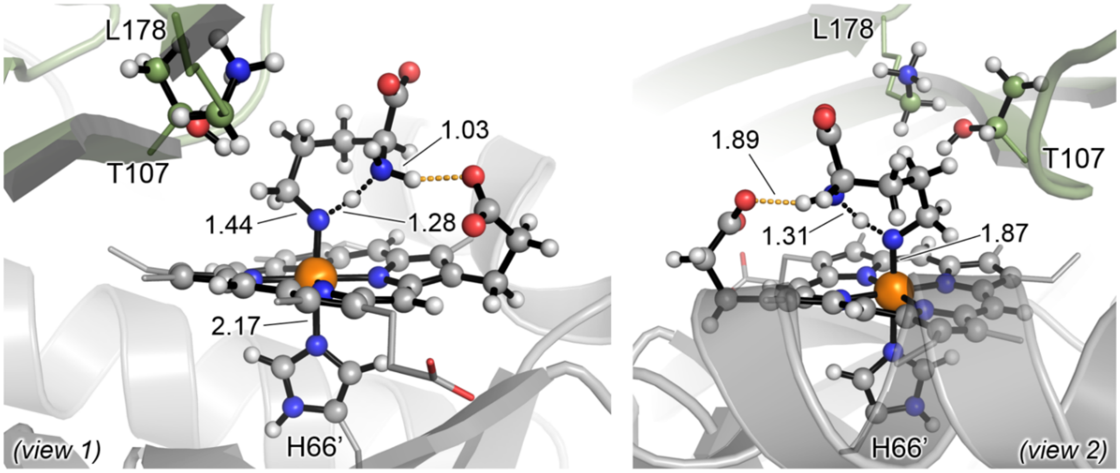

PipS-**TS3** (OSS)
ΔG‡ = -0.6 (ΔE‡ = 0.5)

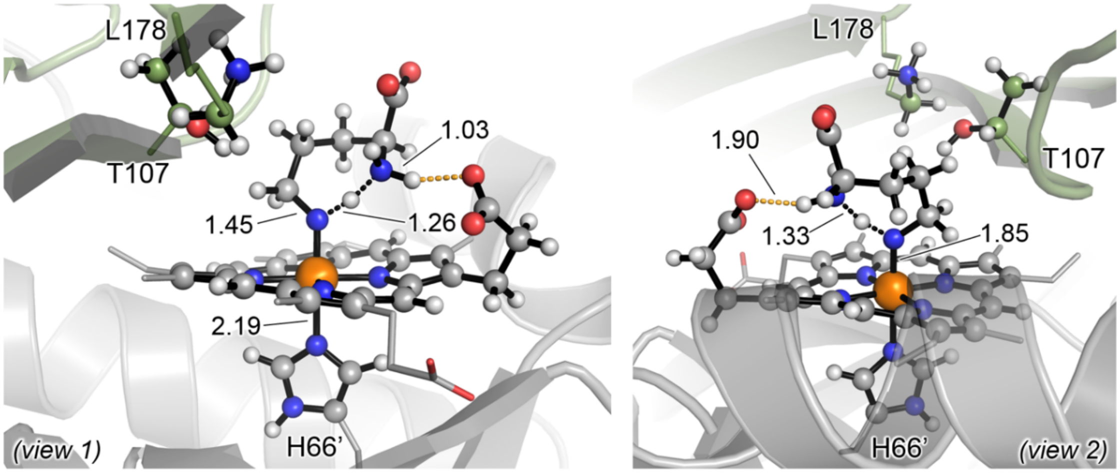

PipS-**TS3** (T)
ΔG‡ = 0.4 (ΔE‡ = 1.8)

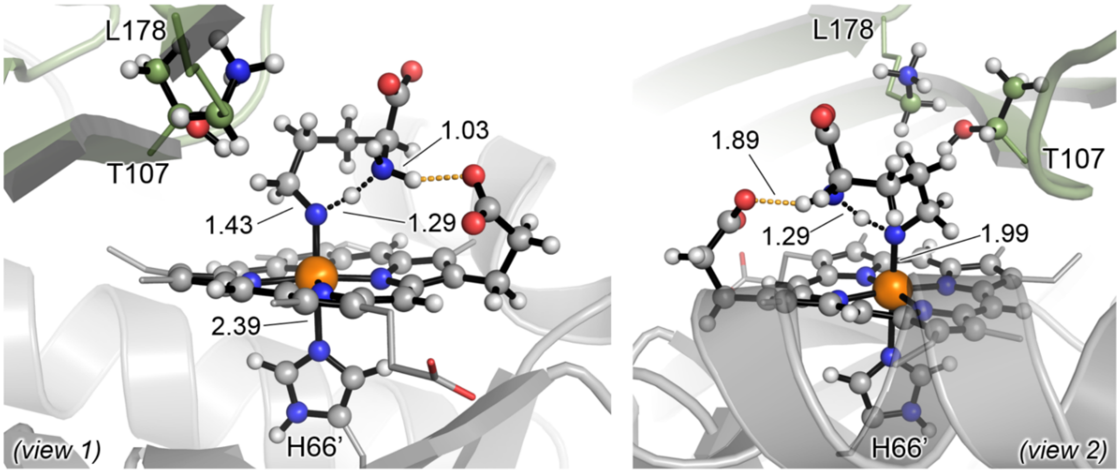

PipS-**TS3** (Q)
ΔG‡ = -6.0 (ΔE‡ = -3.0)

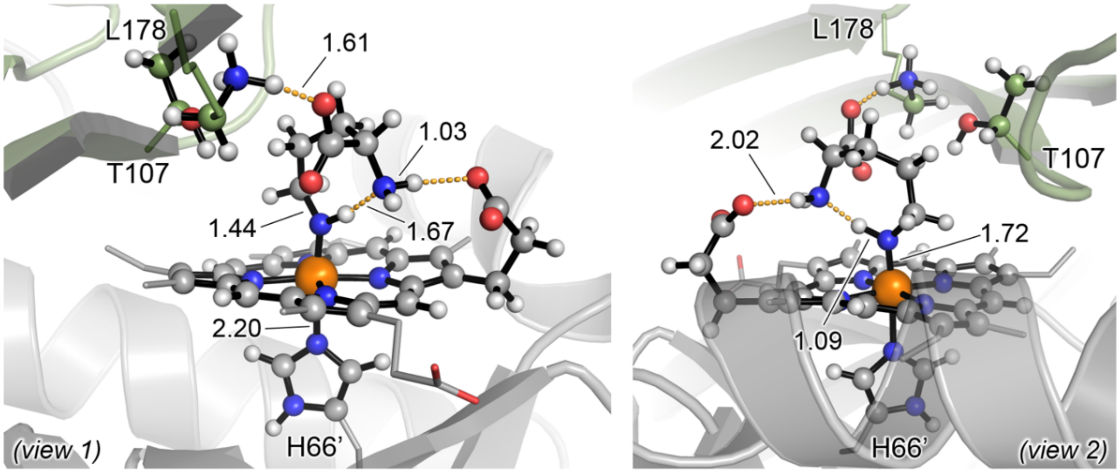

PipS-**Int3** (CSS)
$\Delta G_r$ = -7.6 ($\Delta E_r$ = -11.8)

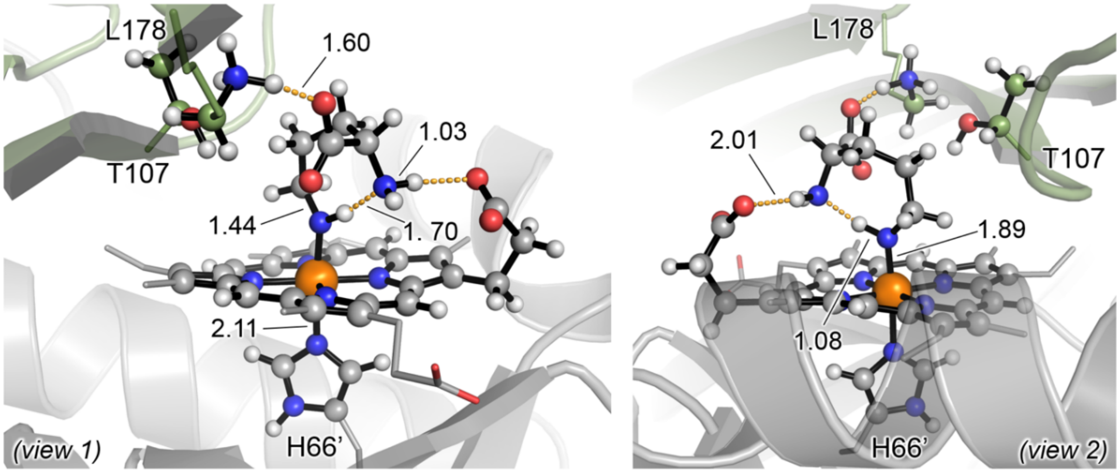

PipS-**Int3** (OSS)
$\Delta G_r$ = -10.4 ($\Delta E_r$ = -15.2)

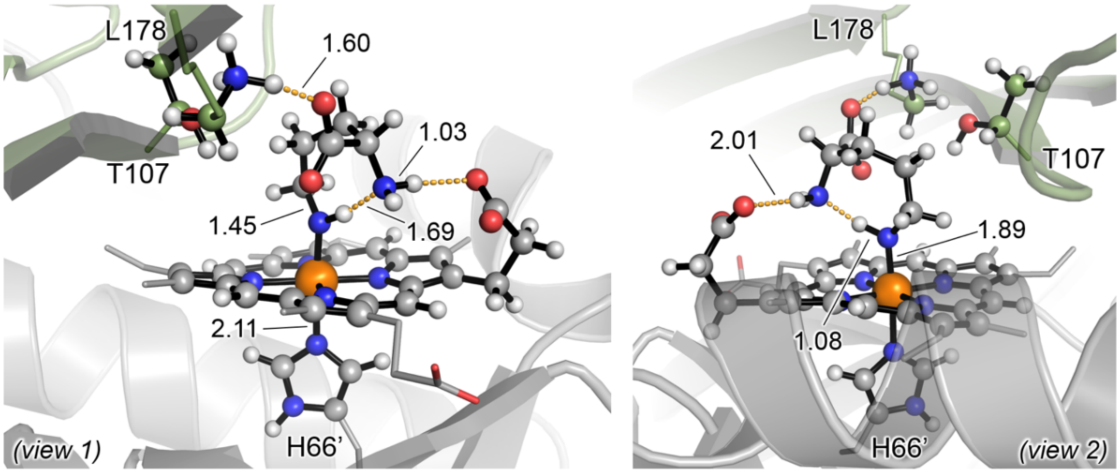

PipS-**Int3** (T)
$\Delta G_r$ = -8.8 ($\Delta E_r$ = -13.6)

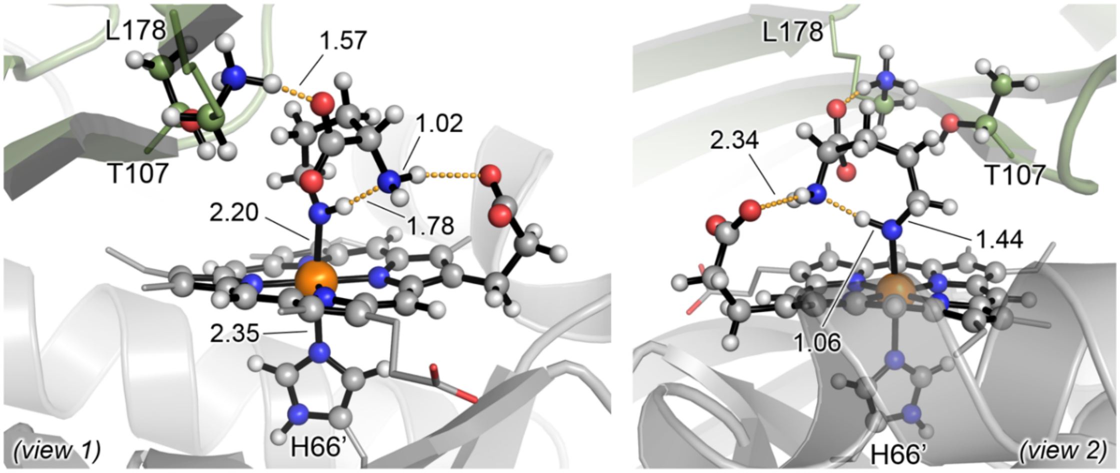

PipS-**Int3** (Q)
$\Delta G_r$ = -22.7 ($\Delta E_r$ = -21.0)

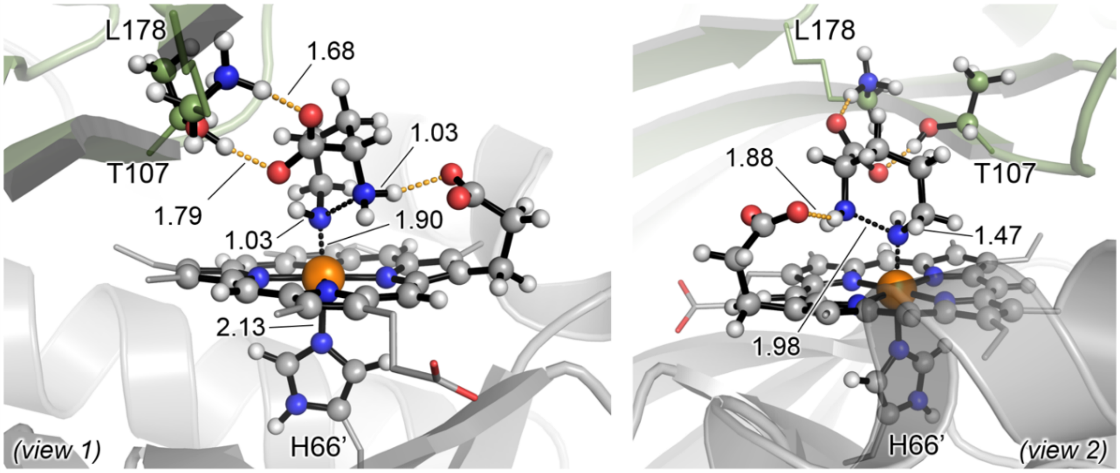

PipS-**TS4** (OSS)
$\Delta G^‡$ = 14.1 ($\Delta E^‡$ = 13.3)

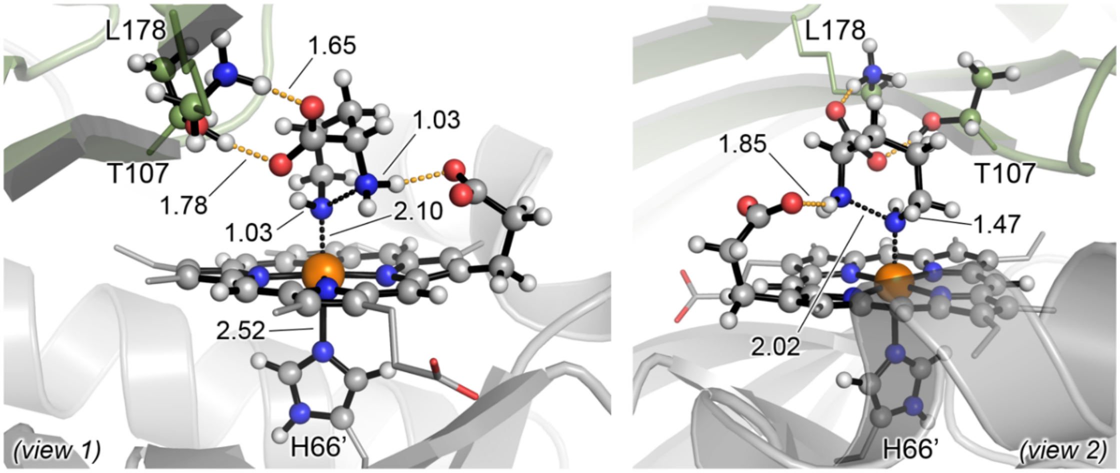

PipS-**TS4** (T)
$\Delta G^{\ddagger}$ = 18.9 ($\Delta E^{\ddagger}$ = 21.2)

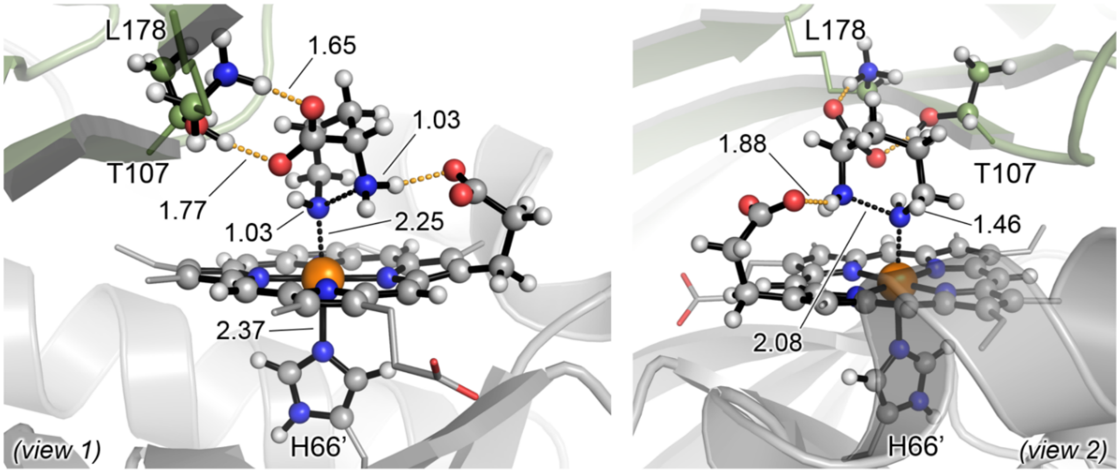

PipS-**TS4** (Q)
$\Delta G^{\ddagger}$ = 13.9 ($\Delta E^{\ddagger}$ = 15.2)

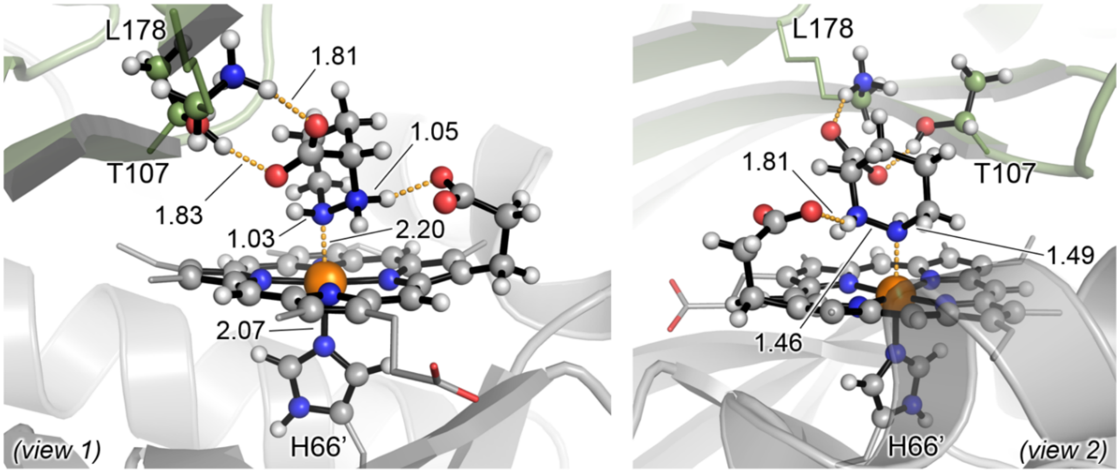

PipS-L-**2** (CSS)
$\Delta G_r$ = -1.1 ($\Delta E_r$ = -5.6)

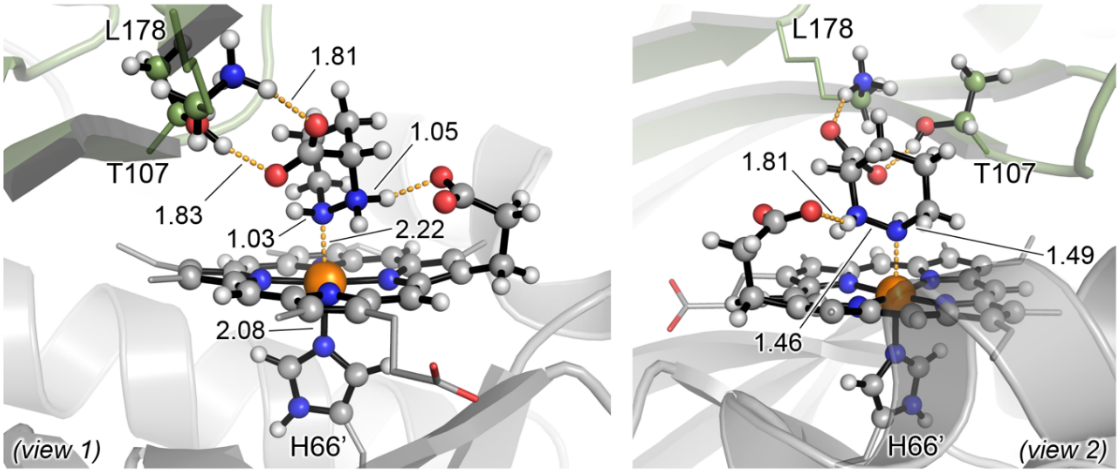

PipS-L-**2** (OSS)
$\Delta G_r$ = 8.7 ($\Delta E_r$ = 4.6)

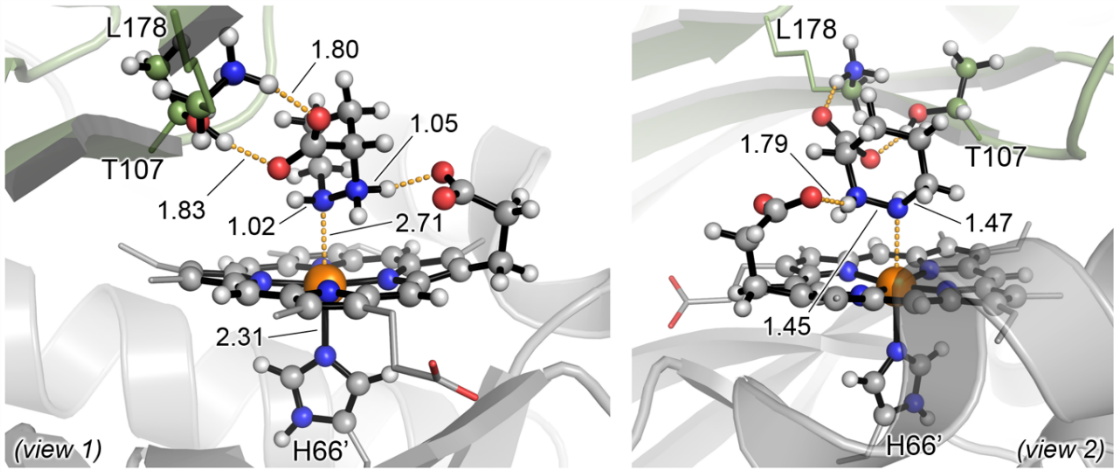

PipS-L-**2** (T)
$\Delta G_r$ = 1.7 ($\Delta E_r$ = -0.6)

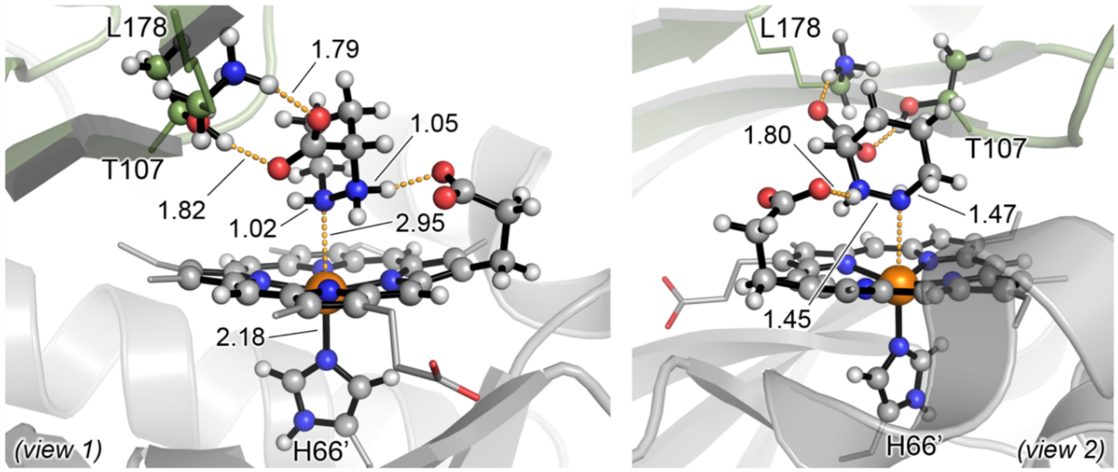

PipS-L-**2** (Q)
$\Delta G_r$ = -4.1 ($\Delta E_r$ = -7.0)

**(e)**

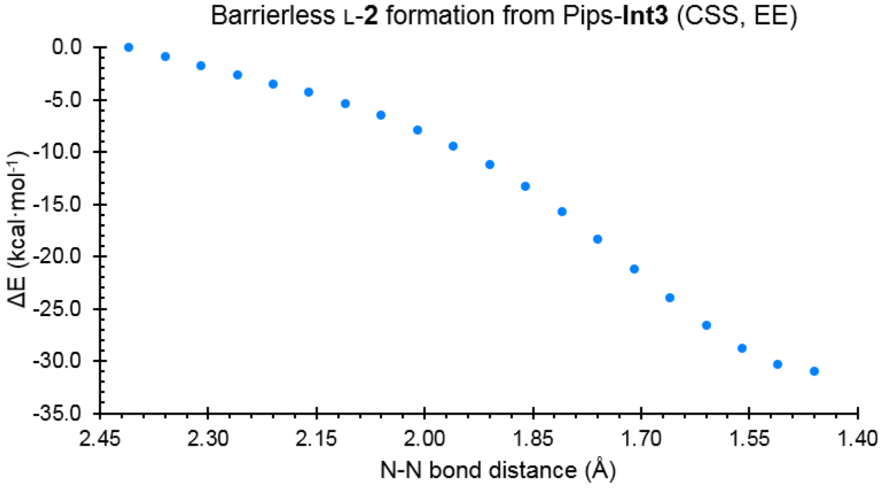

**(f)**

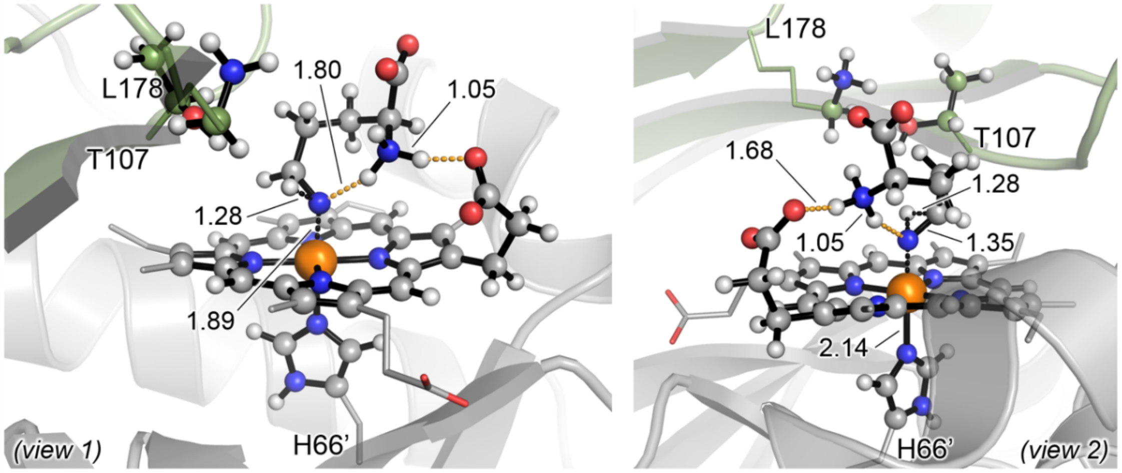

PipS-**TS5a** (CSS)
ΔG‡ = 19.2 (ΔE‡ = 22.2)

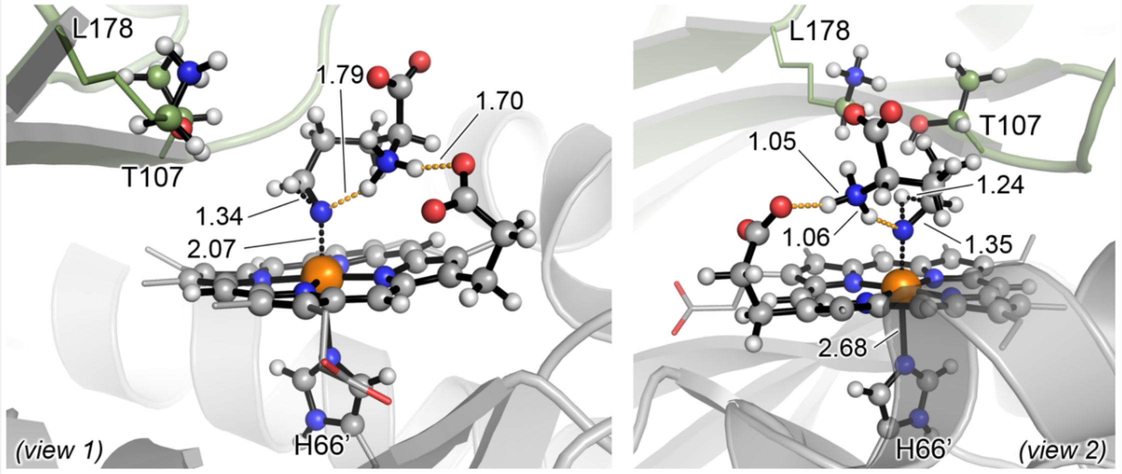

PipS-**TS5a** (T)
ΔG‡ = 24.8 (ΔE‡ = 29.1)

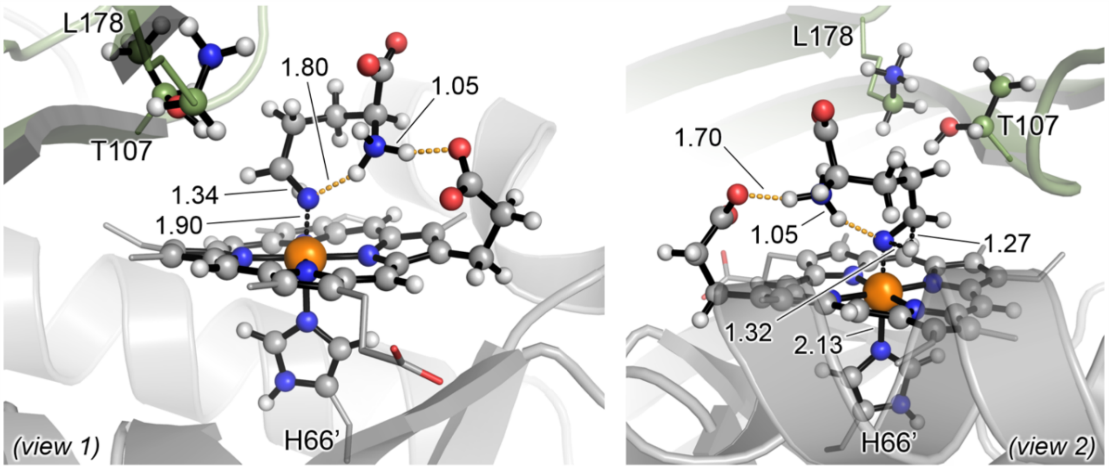

PipS-**TS5b** (CSS)
$\Delta G^{\ddagger}$ = 28.7 ($\Delta E^{\ddagger}$ = 31.2)

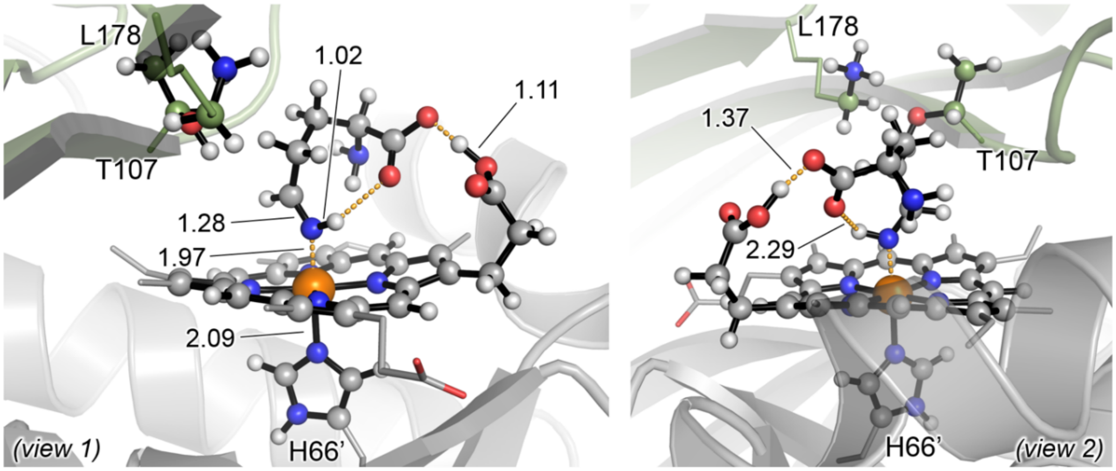

PipS-L-**3** (CSS)
$\Delta G_r$ = -45.8 ($\Delta E_r$ = -49.1)

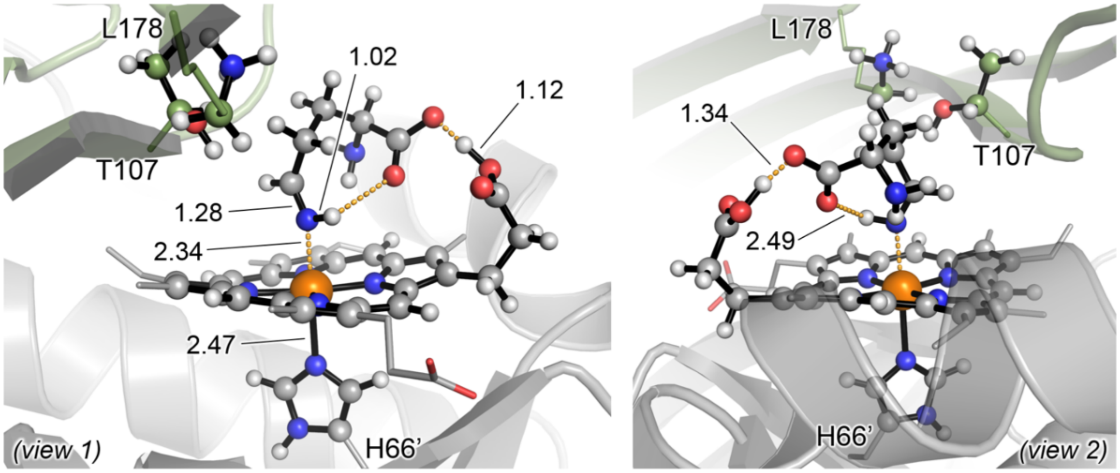

PipS-L-**3** (T)
$\Delta G_r$ = -29.9 ($\Delta E_r$ = -32.9)

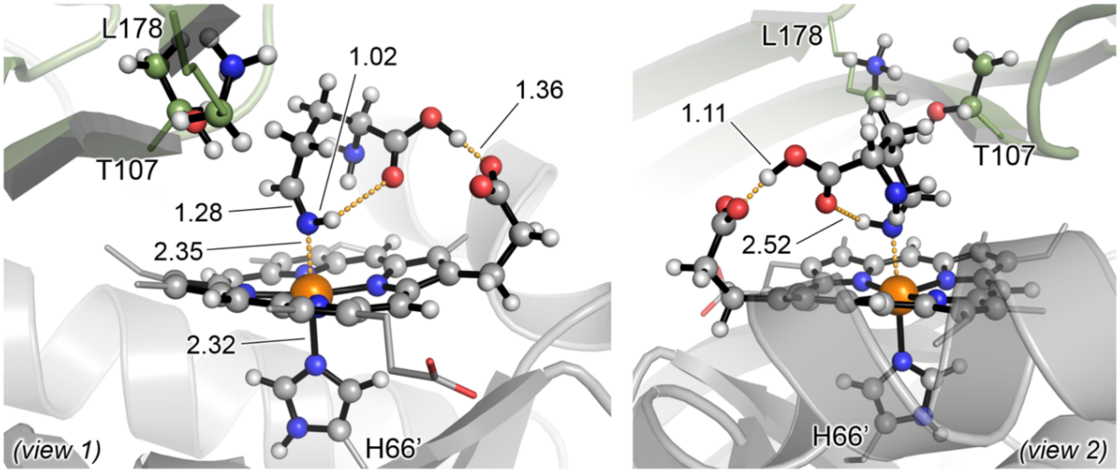

PipS-L-**3** (Q)
$\Delta G_r$ = -50.8 ($\Delta E_r$ = -50.7)

*PipS catalyzed N-N bond formation from Fe-nitrene to give L-**2** product*

Nitrene PipS-**Int2** optimized in its triplet ground state is set as the energy reference. Quintet electronic state for this intermediate was also explored but was found to be energetically much less stable (ΔG = 16.7 kcal·mol$^{-1}$, see **Supplementary Figure 27b**).

MD simulations (see **Supplementary Figures 22** to **25**) showed that the protonated α-amine in PipS-**Int2** can explore conformations in which it gets close to the nitrene N center (2.8 Å between nitrogen atoms in optimized PipS-**Int2**) thanks to its interaction with the Heme propionate group (1.7 Å). The distance between the α-amine proton and the N-nitrene is 1.8 Å in the optimized PipS-**Int2** intermediate (**Supplementary Figure 27c**). Considering this, a direct proton transfer from the protonated α-amine to the N-nitrene was modelled.

A transition state for this proton transfer step has been characterized, PipS-**TS3**, with a lowest in energy barrier of $\Delta G^{\ddagger}$ = 0.2 kcal·mol$^{-1}$ and an imaginary frequency of 1028*i* cm$^{-1}$ for OSS electronic state (**Supplementary Figure 27b**). The computed low Gibbs activation barriers indicate that the proton transfer can easily occur when the two nitrogen groups are close enough to react. The resulting intermediate, PipS-**Int3** (**Supplementary Figure 27a**), is $\Delta G_r$ = -9.6 kcal·mol$^{-1}$ (OSS, lowest in energy) more stable than nitrene PipS-**Int2** (T) intermediate. In the optimized PipS-**Int3** structure, the neutral α-amine is well preorganized to perform a nucleophilic attack to the protonated Fe-nitrenoid N center (N-N distance of 2.8 Å, **Supplementary Figure 27c**), which is in part facilitated by the H-bond established between the carboxylic group of the intermediate and K178 residue, which stabilizes this particular conformation. Charge distribution and spin density analysis of optimized PipS-**Int3** structures at all computed electronic states revealed that there is no spin density localized on the α-amine group (see **Supplementary Figure 27**), indicating that this reaction step corresponds to a proton transfer and not to an H-abstraction. This is in contrast to the recently reported Fe-nitrene insertion into C-H bonds, which involves an H-abstraction step (see for instance: Yang, Y.; Cho, I.; Qi, X.; Liu, P.; Arnold, F. H.. An enzymatic platform for the asymmetric amination of primary, secondary and tertiary C($sp^3$)–H bonds. *Nat. Chem*. **11**, 987–993 (2019). DOI: 10.1038/s41557-019-0343-5).

From PipS-**Int3**, a N-N bond forming transition state PipS-**TS4** has been optimized with an activation barrier of $\Delta G^{\ddagger}$ = 14.1 kcal·mol$^{-1}$ for lowest in energy OSS. Optimized PipS-**TS4** (OSS) has an N-N distance of 2.0 Å and an imaginary frequency of 470*i* cm$^{-1}$, as shown in **Supplementary Figure 27**. Electrostatic embedding relaxed scan calculations along the N-N reaction coordinate suggest that this is a barrierless process in the higher in energy CSS state (**Supplementary Figure 27e**). Spin density analysis revealed that slightly partial radical character on the α-amine is being generated in the PipS-**TS4** (in OSS, T, and Q electronic states, see **Supplementary Fig 28**).

The final PipS-L-**2** product complex (Q electronic state) has a relative Gibbs free energy of $\Delta G$ = -10.1 kcal·mol$^{-1}$ respect to PipS-**Int2** intermediate (Q).

*PipS catalyzed imine L-3 formation from Fe-nitrene via direct proton migration*

From PipS-**Int2** intermediate, imine product L-**3** can be formed through a direct H-migration from C5 to N-nitrene center. There are no polar residues in the active site of PipS close enough to the C5 position suitable to deprotonate it (**Supplementary Figures 22** to **25**).

Direct deprotonation at C5 position by substrate's α-carboxylic group is not possible because, when formed in PipS active site, intermediate PipS-**Int2** cannot explore conformations in which its carboxylic group gets close enough to C5 to deprotonate it. MD simulations showed that α-carboxylic group in the nitrene intermediate is mainly interacting with different surrounding residues that prevent it to explore conformations suitable for promoting C5 deprotonation.

The computed Gibbs energy barriers for the migrations of the two non-equivalent C5-Ha and C5-Hb are $\Delta G^{\ddagger}$ = 20.0 and 29.5 kcal·mol$^{-1}$ with imaginary frequencies of 978*i* and 926*i* cm$^{-1}$ for PipS-**TS5a** and PipS-**TS5b** in the lowest in energy CSS state, respectively. PipS-**TS5a** in triplet state is higher in energy, and triplet PipS-**TS5b** could not be optimized.

The final PipS-L-**3** product complex has a relative Gibbs free energy of ΔG = -37.4 kcal·mol$^{-1}$ (CSS) respect to PipS-**Int2** intermediate (Q).

Taking all these together, it is proposed that when the intermediate PipS-**Int2** is formed, it can explore conformations that bring the protonated α-amine close to the N-nitrene favored by the complementarity between L-**1** and PipS active site. This is driven by the interactions occurring between the amino acid group, the Heme propionate and K178 and other active site polar residues, together with the shape of the active site cavity. When the protonated α-amine and N-nitrene are closer enough, a proton transfer can quickly occur, activating the α-amine for a nucleophilic attack to the electrophilic protonated nitrene and forming a new N-N bond through an energetically accessible Pips-**TS4**. Although imine formation pathway is possible via a direct 1,2-H migration, it has a higher activation barrier than the proposed N-N bond forming pathway. However, this imine formation pathway becomes possible when no additional amine groups are present in the substrate molecule (as for substrates **4**, **5**, and **8**), or when the nucleophilic amine cannot be well positioned to effectively react with the N-Fe (as in the substrate analog D-**1** with inverted chiral α-amine).

Computational results, based on the combination of MD simulations and QM/MM calculations, are in good agreement with the experimental observations and elucidate the molecular basis of the reaction mechanisms by which PipS and other enzyme homologs uses to produce piperazic acid and imine derivatives.

**Supplementary Figure 28. Spin density and Mulliken charge distribution analysis in key intermediates and transition states for N-N bond formation.**

Atom-centered Mulliken spin densities and charges at key atoms (Fe, N1 and N2) in QM/MM optimized nitrenoid and nitrene intermediates, and N-N forming transition state (PipS-**Int2**, PipS-**Int3**, and PipS-**TS4**; see **Supplementary Figure 27**). Values obtained at (U)B3LYP/Def2TZVP:AmberFF14SB single point level.

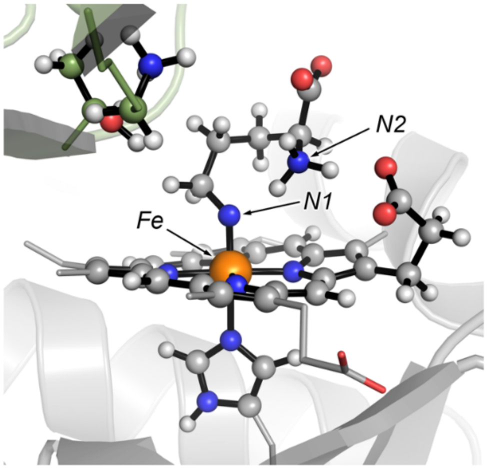

| Structure | Electronic State | Fe | | N1 | | N2 | |
|---|---|---|---|---|---|---|---|
| | | Charge | Spin Density | Charge | Spin Density | Charge | Spin Density |
| PipS-**Int2** | CSS | 0.04 | 0.00 | -0.26 | 0.00 | -0.28 | 0.00 |
| | OSS | 0.03 | -1.04 | -0.28 | 0.91 | -0.28 | 0.00 |
| | T | 0.08 | 1.07 | -0.31 | 0.93 | -0.27 | 0.01 |
| | Q | 0.18 | 2.72 | -0.44 | 1.30 | -0.29 | 0.02 |
| PipS-**Int3** | CSS | -0.15 | 0.00 | 0.02 | 0.00 | -0.58 | 0.00 |
| | OSS | -0.13 | -1.05 | -0.02 | 0.90 | -0.59 | 0.00 |
| | T | -0.14 | 1.18 | -0.06 | 0.82 | -0.58 | 0.01 |
| | Q | 0.15 | 2.87 | -0.29 | 1.03 | -0.62 | 0.01 |
| PipS-**TS4** | CSS | | | * barrierless | | | |
| | OSS | -0.03 | -0.57 | -0.04 | 0.27 | -0.29 | 0.23 |
| | T | 0.14 | 1.66 | -0.18 | 0.21 | -0.29 | 0.20 |
| | Q | 0.45 | 4.14 | -0.31 | -0.31 | -0.32 | -0.19 |

The negligible spin density on the neutral α-amine (N2) in PipS-**Int3** intermediate is consistent with a proton transfer mechanism (PipS-**TS3**) from the α-amine to the Fe-N nitrene center in PipS-**Int2** to give PipS-**Int3**. This also supports the subsequent nucleophilic attack of the neural α-amine to the electrophilic N-Fe center during the N-N bond formation step from PipS-**Int3** through PipS-**TS4**.

Partial spin density localized on the nucleophilic amine N2 atom is being generated at the N-N bond forming PipS-**TS4** transition state.

**Supplementary Figure 29. Exploration of alternative imine L-3 formation along the nitrene formation path using QM/MM calculations.** QM/MM exploration of deprotonation of nitrenoid intermediate PipS-**Int1** at C5 by the newly released water molecule from previous PipS-**TS1** and the catalytic dyad (K178, T107) to form the imine product PipS-L-**3**.

(**a**) The catalytic cycle of PipS corresponding to the imine formation pathway through deprotonation at PipS-**Int1** C5. (in orange).

(**b**) QM/MM relaxed PES scan along the C5 deprotonation coordinate in nitrenoid intermediate PipS-**Int1** to form imine product (PipS-L-**3**) at (U)B3LYP/6-31G(d)+SDD(Fe):AmberFF14SB (triplet (T) electronic state, mechanical embedding).

(**c**) Structure of relevant points along the previous scan calculation in **b**.

(**d**) Relative stabilities in terms of electronic energy (ΔE), enthalpy (ΔH), and Gibbs energy (ΔG) for different species were obtained at the (U)B3LYP/Def2TZVP:AmberFF14SB// (U)B3LYP/6-31G(d)+SDD(Fe):AmberFF14SB level, with the same MM parameters used in MD simulations (see SI, computational details). All energies are referred considering PipS-**Int1** (OSS) structure as zero.

(**e**) Optimized QM/MM structures for PipS-**TS6**. All atoms included in the QM-region are shown in ball and stick representation.

All energies are given in kcal·mol$^{-1}$. Distances are given in angstrom (Å).

(a)

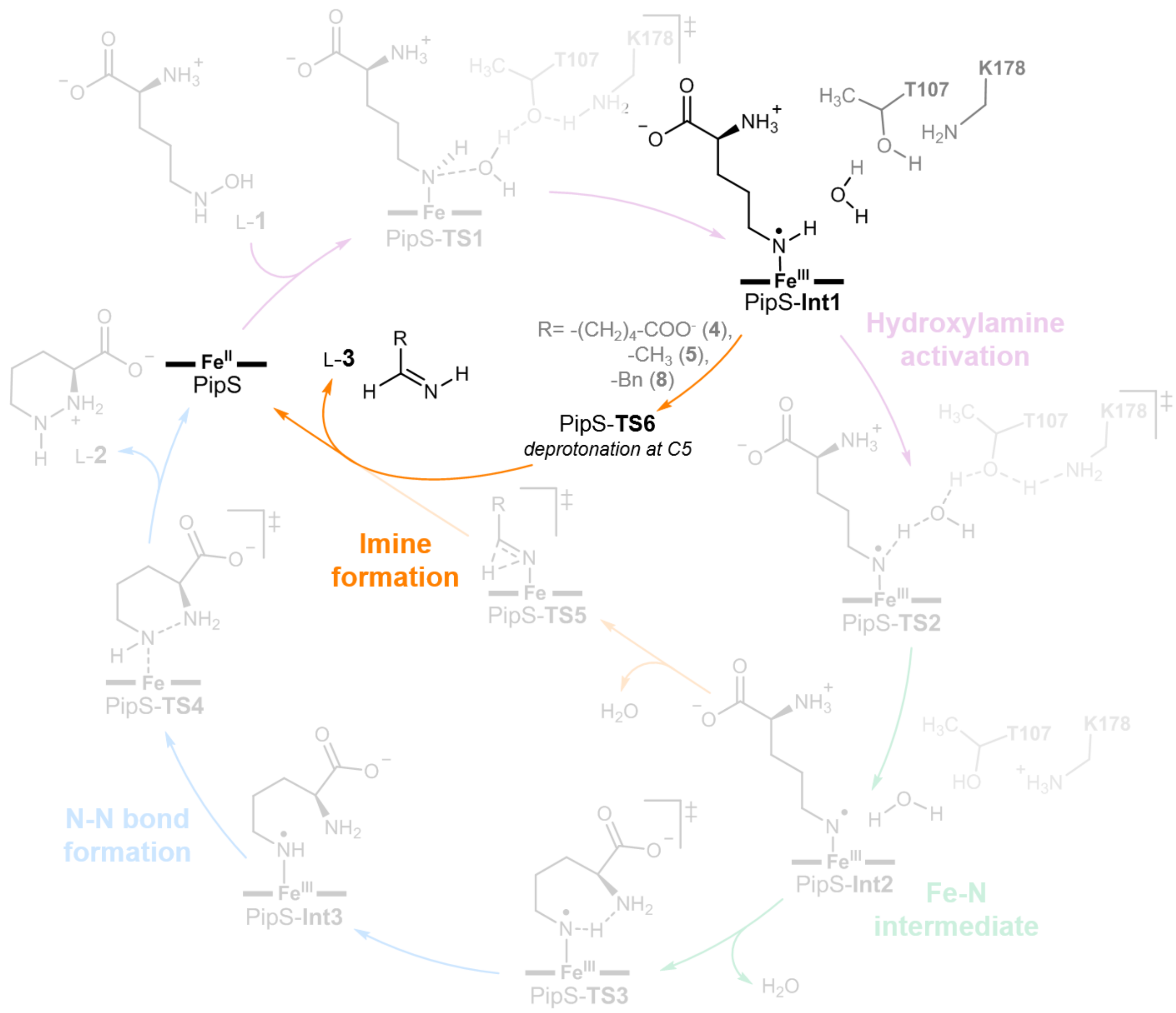

L-1
PipS-TS1
PipS-Int1
Hydroxylamine activation
R= -(CH2)4-COO- (4), -CH3 (5), -Bn (8)
PipS-TS6
deprotonation at C5
L-3
PipS
L-2
Imine formation
PipS-TS5
PipS-TS2
PipS-TS4
N-N bond formation
PipS-Int3
PipS-TS3
PipS-Int2
Fe-N intermediate
T107
K178

(b)

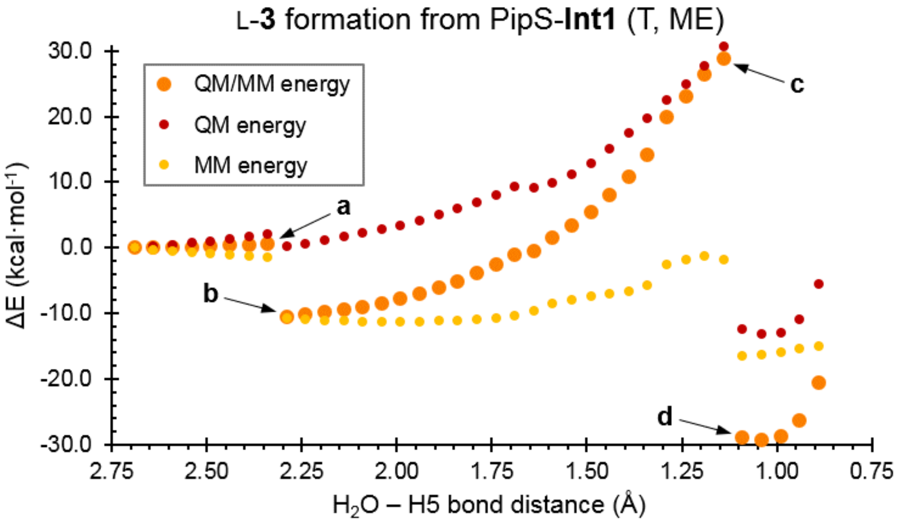

L-3 formation from PipS-Int1 (T, ME)
QM/MM energy
QM energy
MM energy
ΔE (kcal·mol-1)
H2O – H5 bond distance (Å)
a
b
c
d

**(c)**

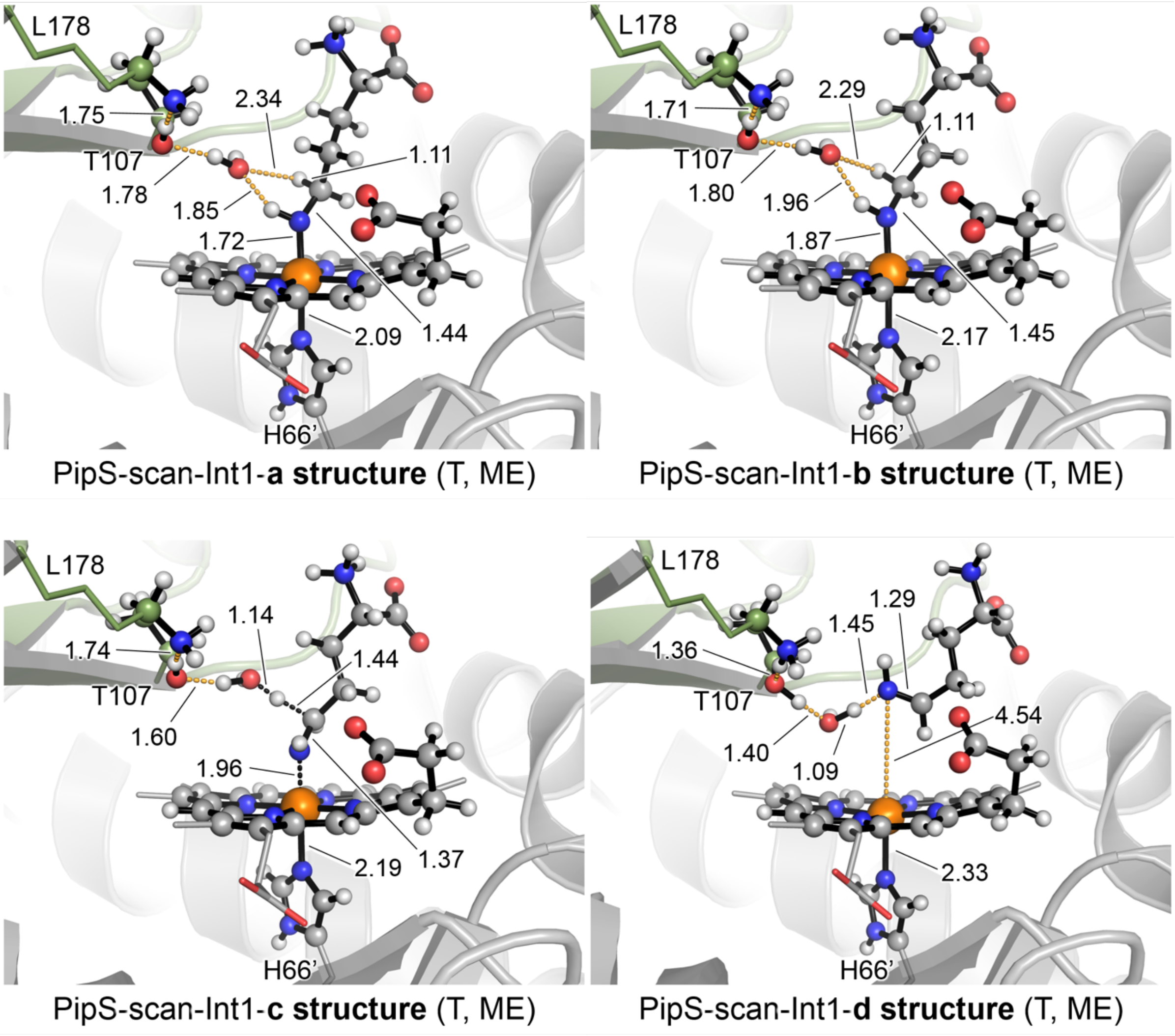

**(d)**

| Structure | Electronic State | $\Delta E_{QM}$ | ΔE | ΔH | ΔG |
|---|---|---|---|---|---|
| PipS-**Int1** | close-shell singlet (CSS) | 7.5 | 5.6 | 6.1 | 6.8 |
| | open-shell singlet (OSS) | 0.0 | 0.0 | 0.0 | 0.0 |
| | triplet (T) | 5.3 | 1.6 | 1.6 | 1.4 |
| | quintet (Q) | 11.4 | 9.2 | 9.1 | 7.4 |
| PipS-**TS6** | close-shell singlet (CSS)[a] | - | - | - | - |
| | open-shell singlet (OSS) | 1.5 | -0.4 | -3.3 | -0.9 |
| | triplet (T) | 9.9 | 8.4 | 5.0 | 6.5 |
| | quintet (Q) | 13.5 | 11.3 | 7.4 | 7.2 |
| PipS-**TS2**[b] | close-shell singlet (CSS) | 17.9 | 15.7 | 10.2 | 11.5 |
| | open-shell singlet (OSS) | 15.6 | 11.9 | 6.4 | 7.6 |
| | triplet (T) | 16.2 | 12.5 | 6.8 | 7.3 |
| | quintet (Q) | 35.8 | 32.3 | 25.5 | 24.1 |

[a] Could not be optimized. [b] From **Supplementary Figure 26**.

**(e)**

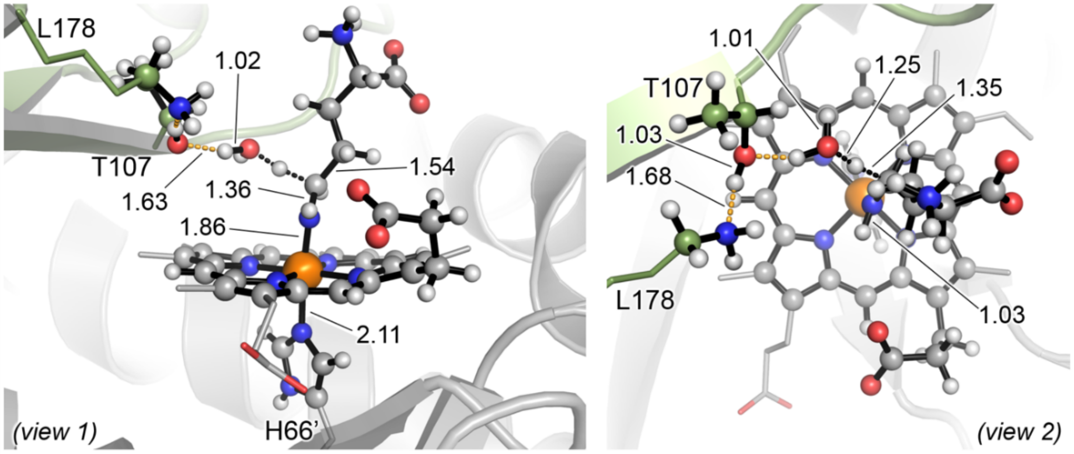

PipS-**TS6** (OSS)
ΔG‡ = -0.9 (ΔE‡ = -0.4)

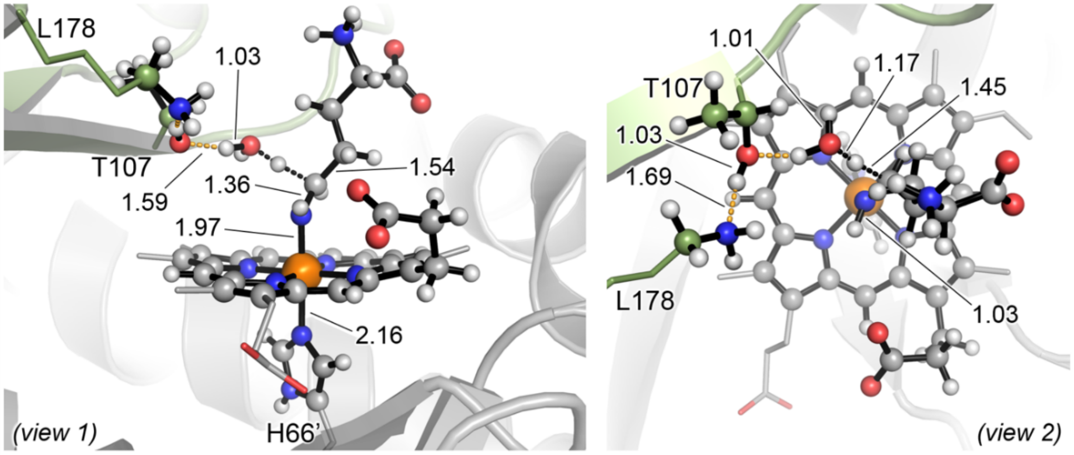

PipS-**TS6** (T)
ΔG‡ = 5.1 (ΔE‡ = 6.8)

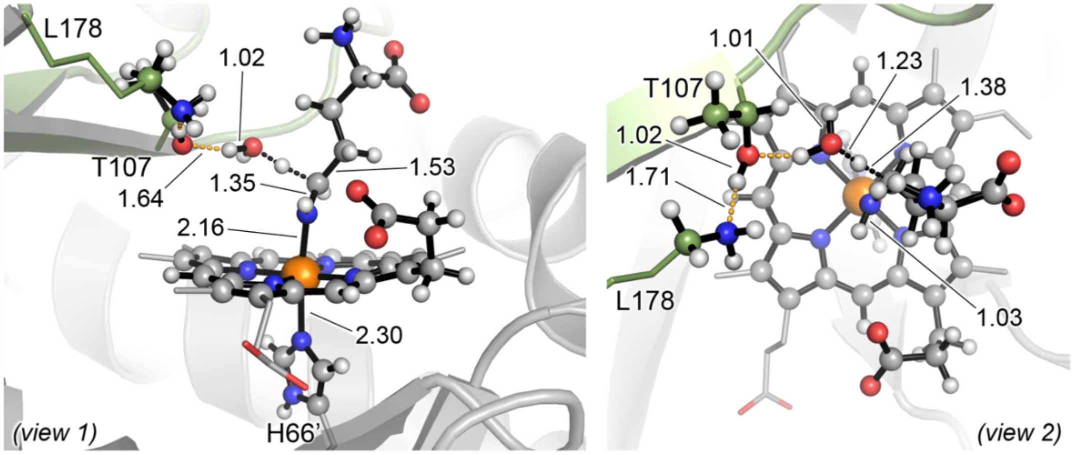

PipS-**TS6** (Q)
ΔG‡ = -0.2 (ΔE‡ = 2.1)

The formation of imine product L-**3** from nitrenoid intermediate PipS-**Int1** has been explored. PipS-**Int1** is an intermediate generated over the course of the formation of nitrene PipS-**Int2** pathway (described in **Supplementary Figure 26**)

Once nitrenoid intermediate PipS-**Int1** is formed (during the course of PipS-**Int2** formation), deprotonation at its δ-C5 position can take place through PipS-**TS6**. Deprotonation at δ-C5 position via PipS-**TS6** is an alternative to deprotonation at NH-Fe position through PipS-**TS2**. This deprotonation at δ-C5 position involves the released $H_2O$ molecule and the catalytic K178 and T107 residues, and it will restore the initial protonation state of K178.

In PipS-**Int1**, the distance between the oxygen atom of the $H_2O$ molecule and the H-C5 proton is relatively large (**Supplementary Figure 26**) as compared to the distance between the water molecule and the nitrenoid proton (3.0 Å versus 1.6 Å, respectively). Consequently, this optimal PipS-**Int1** geometry preorganizes the catalytic machinery towards nitrenoid deprotonation through PipS-**TS2** as early described. Consequently, in order to explore the deprotonation at δ-C5 position a conformational rearrangement of the intermediate PipS-**Int1** must take place. This conformational change has been explored using mechanical embedding relaxed scan calculations along the reaction coordinate defined as the distance between the oxygen atom of the $H_2O$ molecule and the H-C5 proton PipS-**Int1** (T) (**Supplementary Figure 29**).

Relaxed scan calculation starting from PipS-**Int1** (T) intermediate geometry, indicated that the intermediate first needs to reorient to explore a different conformation ($H_2O$ – H-C5 distance ca. 2.25 Å, local minima) in which the direct interaction between the H-C5 proton and the water molecule becomes possible (**Supplementary Figure 29b**). From there, the proton transfer step starts, with a constant increase of the QM and global QM/MM energy to give an approximate activation barrier of ca. 30 kcal·mol$^{-1}$ to finally deprotonate C5 position.

Starting from the new characterized conformation for PipS-**Int1**, the corresponding deprotonation transition state PipS-**TS6** has been optimized considering the OSS, T, and Q electronic states (**Supplementary Figure 29e**; it could not be optimized in the CSS state). The computed Gibbs activation barriers for PipS-**TS6** ($\Delta G^{\ddagger}$ = -0.9*, 6.5, and 7.2 kcal·mol$^{-1}$ with imaginary frequencies of 1063*i*, 1021*i*, and 1997*i* cm$^{-1}$ for OSS, T, and Q respectively; *barrierless) are lower than those obtained for PipS-**TS2** ($\Delta G^{\ddagger}$ = 7.3, 7.6, 11.5, and 24.1 for T, OSS, CSS, and Q respectively). These results indicate that both deprotonation pathways, PipS-**TS2** and PipS-**TS6**, have very low activation barriers and can directly compete. It is interesting to note that the optimized PipS-**TS6** do not directly restore the initial protonation state of K178, and the abstracted proton forms a transient hydroxonium ion (**Supplementary Figure 29e**), in contrast to PipS-**TS2**.

Taking all these together, it is proposed that geometric preorganization of PipS-**Int1** formed from PipS-**TS1** (**Supplementary Figure 26**) promotes the deprotonation at N-nitrenoid position through PipS-**TS2**. When formed in PipS active site, PipS-**Int1** has an appropriate geometric preorganization relative to the catalytic machinery to generate the nitrene PipS-**Int2** intermediate through the low in energy PipS-**TS2**. On the other hand, although deprotonation of PipS-**Int1** at C5 position to form the imine product L-**3** is energetically feasible through the low in energy PipS-**TS6**, a previous conformational rearrangement of intermediate PipS-**Int1** is required, which prevents it to occur.

**Supplementary Figure 30. Summary of QM/MM calculations on PipS catalytic cycle and reaction mechanism in the Fe(II) oxidation state.**

(**a**) The proposed complete catalytic cycle of PipS in the Fe(II) oxidation state. The hydroxylamine activation, the key Fe-N intermediate, N-N bond formation, and imine formation pathways are shown in purple, green, blue, and orange, respectively.

(**b**) Schematic Potential Energy Surface (PES) for PipS catalytic cycle starting from the hydroxylamine activation and leading to N-N bond formation and imine formation pathways. Energy values were obtained from **Supplementary Figure 26** and **27**.

**(a)**

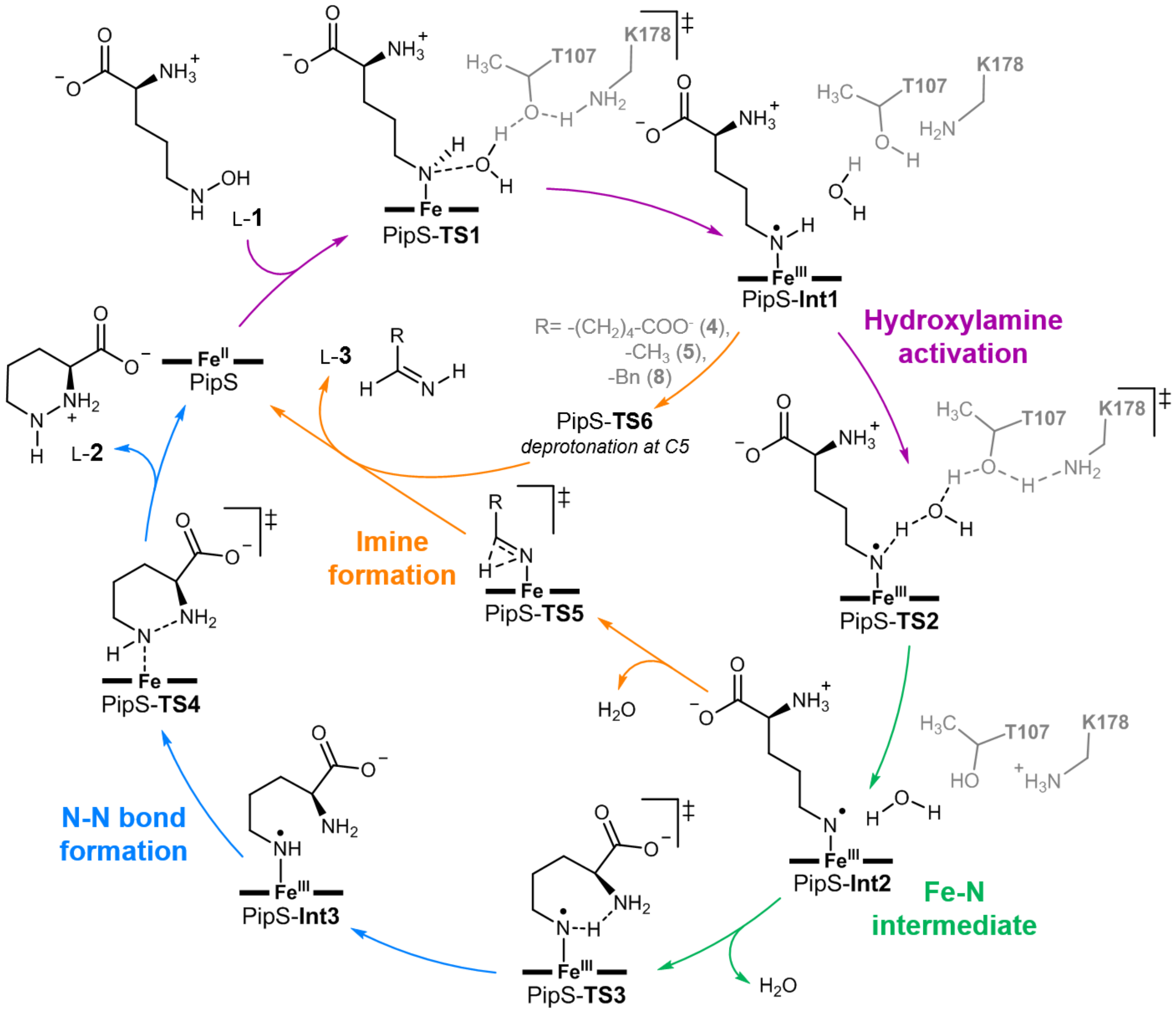

**(b)**

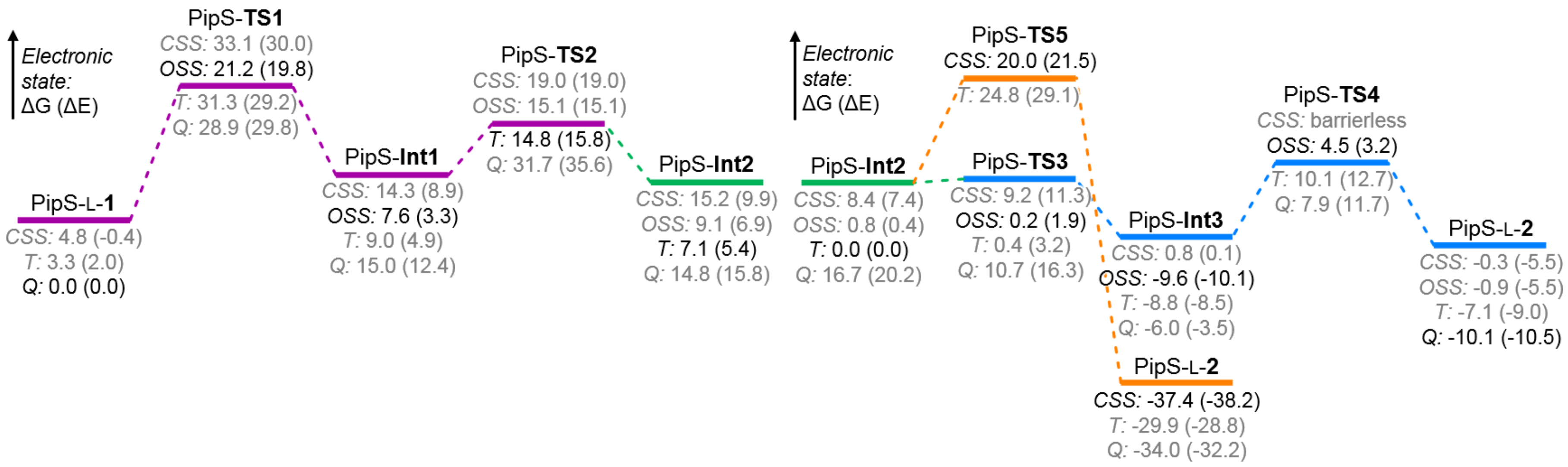

Electronic state: ΔG (ΔE)
PipS-L-1
CSS: 4.8 (-0.4)
T: 3.3 (2.0)
Q: 0.0 (0.0)
PipS-TS1
CSS: 33.1 (30.0)
OSS: 21.2 (19.8)
T: 31.3 (29.2)
Q: 28.9 (29.8)
PipS-Int1
CSS: 14.3 (8.9)
OSS: 7.6 (3.3)
T: 9.0 (4.9)
Q: 15.0 (12.4)
PipS-TS2
CSS: 19.0 (19.0)
OSS: 15.1 (15.1)
T: 14.8 (15.8)
Q: 31.7 (35.6)
PipS-Int2
CSS: 15.2 (9.9)
OSS: 9.1 (6.9)
T: 7.1 (5.4)
Q: 14.8 (15.8)
Electronic state: ΔG (ΔE)
PipS-Int2
CSS: 8.4 (7.4)
OSS: 0.8 (0.4)
T: 0.0 (0.0)
Q: 16.7 (20.2)
PipS-TS5
CSS: 20.0 (21.5)
T: 24.8 (29.1)
PipS-TS3
CSS: 9.2 (11.3)
OSS: 0.2 (1.9)
T: 0.4 (3.2)
Q: 10.7 (16.3)
PipS-Int3
CSS: 0.8 (0.1)
OSS: -9.6 (-10.1)
T: -8.8 (-8.5)
Q: -6.0 (-3.5)
PipS-L-2
CSS: -37.4 (-38.2)
T: -29.9 (-28.8)
Q: -34.0 (-32.2)
PipS-TS4
CSS: barrierless
OSS: 4.5 (3.2)
T: 10.1 (12.7)
Q: 7.9 (11.7)
PipS-L-2
CSS: -0.3 (-5.5)
OSS: -0.9 (-5.5)
T: -7.1 (-9.0)
Q: -10.1 (-10.5)

**Supplementary Figure 31. QM/MM exploration of PipS catalyzed hydroxylamine activation pathways in the Fe(III) oxidation state.** QM/MM calculations were carried out starting from the previously selected representative snapshot extracted from MD simulations with PipS-L-**1** bound complex (**Supplementary Figures 18** and **26**).

(**a**) The proposed catalytic cycle of PipS in the Fe(III) oxidation state corresponding to the hydroxylamine activation (in black) and the formation of key nitrene intermediate (in olive).
(**b**) Schematic Potential Energy Surface (PES) for the hydroxylamine activation. Energy values were obtained at the (U)B3LYP/Def2TZVP:AmberFF14SB//(U)B3LYP/6-31G(d)+SDD(Fe):AmberFF14SB level, with the same MM parameters used in MD simulations (see SI, computational details). Relative Gibbs (ΔG) and electronic (ΔE) energies are reported for all electronic states. All energies are referred considering PipS-L-**1** (Qu) structure as zero. The PES profile for the equivalent pathway for the Fe(II) oxidation state is also provided.
(**c**) Computed relative stabilities in terms of electronic QM energy ($\Delta E_{QM}$), electronic QM/MM energy (ΔE), enthalpy (ΔH), and Gibbs energy (ΔG) for the different species. All energies are referred considering PipS-L-**1** (Qu) structure as zero.
(**d**) Optimized QM/MM structures for the different species involved in the Fe(III) reaction pathway. All atoms included in the QM-region are shown in ball and stick representation.

All energies and distances are given in kcal·mol$^{-1}$ and angstrom (Å), respectively.

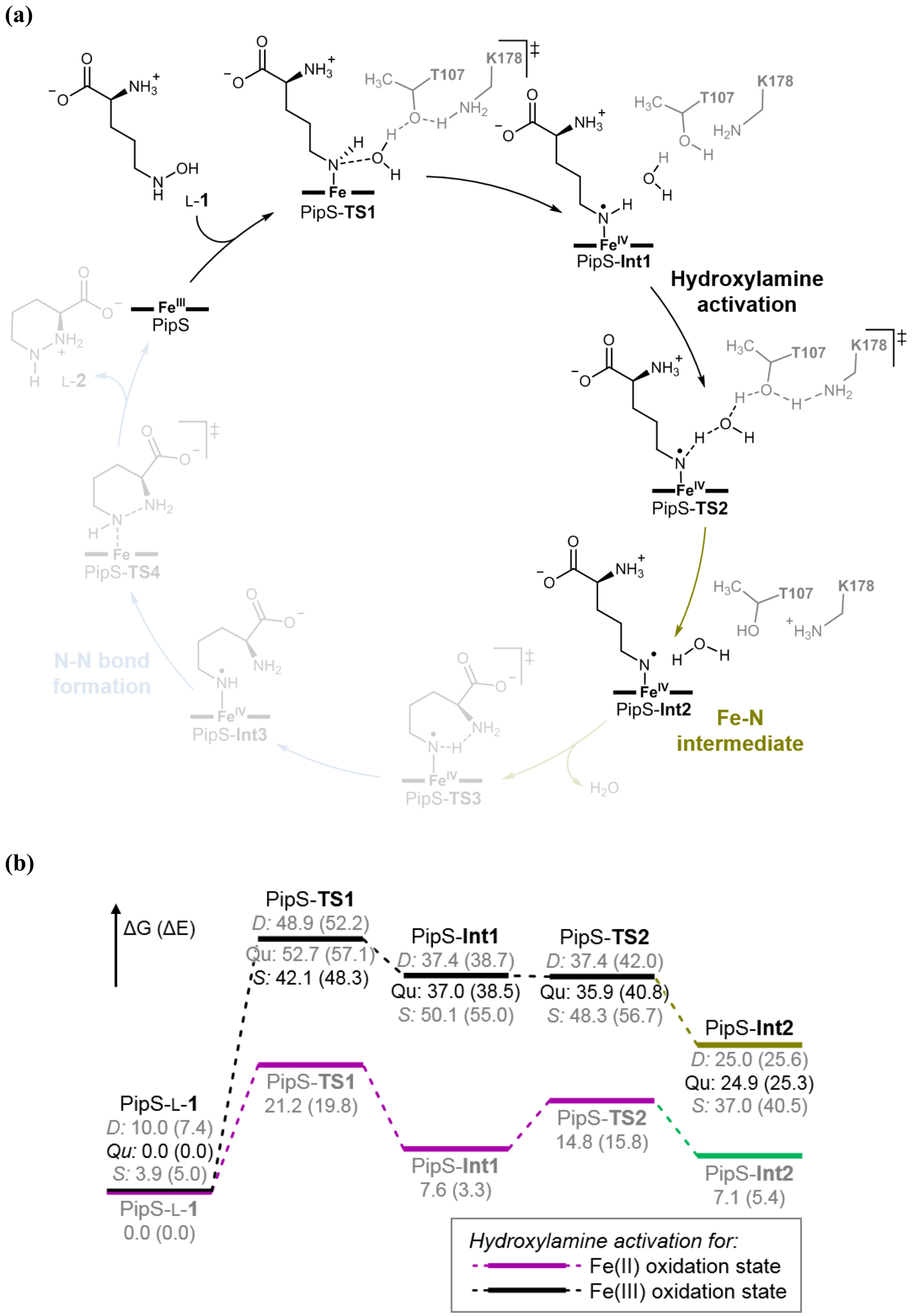
(a)
L-1
PipS-TS1
PipS-Int1
Hydroxylamine activation
PipS-TS2
PipS-Int2
Fe-N intermediate
PipS-TS3
$H_2O$
PipS-Int3
N-N bond formation
PipS-TS4
L-2
PipS
T107
K178
(b)
ΔG (ΔE)
PipS-TS1
D: 48.9 (52.2)
Qu: 52.7 (57.1)
S: 42.1 (48.3)
PipS-Int1
D: 37.4 (38.7)
Qu: 37.0 (38.5)
S: 50.1 (55.0)
PipS-TS2
D: 37.4 (42.0)
Qu: 35.9 (40.8)
S: 48.3 (56.7)
PipS-Int2
D: 25.0 (25.6)
Qu: 24.9 (25.3)
S: 37.0 (40.5)
PipS-L-1
D: 10.0 (7.4)
Qu: 0.0 (0.0)
S: 3.9 (5.0)
PipS-TS1
21.2 (19.8)
PipS-TS2
14.8 (15.8)
PipS-L-1
0.0 (0.0)
PipS-Int1
7.6 (3.3)
PipS-Int2
7.1 (5.4)
Hydroxylamine activation for:
Fe(II) oxidation state
Fe(III) oxidation state

**(c)**

| Structure | Electronic State | $\Delta E_{QM}$ | ΔE | ΔH | ΔG |
|---|---|---|---|---|---|
| PipS-L-**1** | doublet (D) | 11.1 | 7.4 | 7.9 | 10.0 |
| | quartet (Qu) | 0.0 | 0.0 | 0.0 | 0.0 |
| | sextet (S) | 4.5 | 5.0 | 4.1 | 3.9 |
| PipS-**TS1** | doublet (D) | 43.6 | 52.2 | 45.3 | 48.9 |
| | quartet (Qu) | 49.7 | 57.1 | 51.0 | 52.7 |
| | sextet (S) | 41.9 | 48.3 | 41.2 | 42.1 |
| PipS-**Int1** | doublet (D) | 28.6 | 38.7 | 34.9 | 37.4 |
| | quartet (Qu) | 28.6 | 38.5 | 34.7 | 37.0 |
| | sextet (S) | 46.2 | 55.0 | 49.5 | 50.1 |
| PipS-**TS2** | doublet (D) | 32.6 | 42.0 | 34.6 | 37.4 |
| | quartet (Qu) | 31.4 | 40.8 | 33.1 | 35.9 |
| | sextet (S) | 48.1 | 56.7 | 47.2 | 48.3 |
| PipS-**Int2** | doublet (D) | 2.8 | 25.6 | 23.6 | 25.0 |
| | quartet (Qu) | 5.9 | 25.3 | 22.5 | 24.9 |
| | sextet (S) | 21.4 | 40.5 | 36.5 | 37.0 |

**(d)**

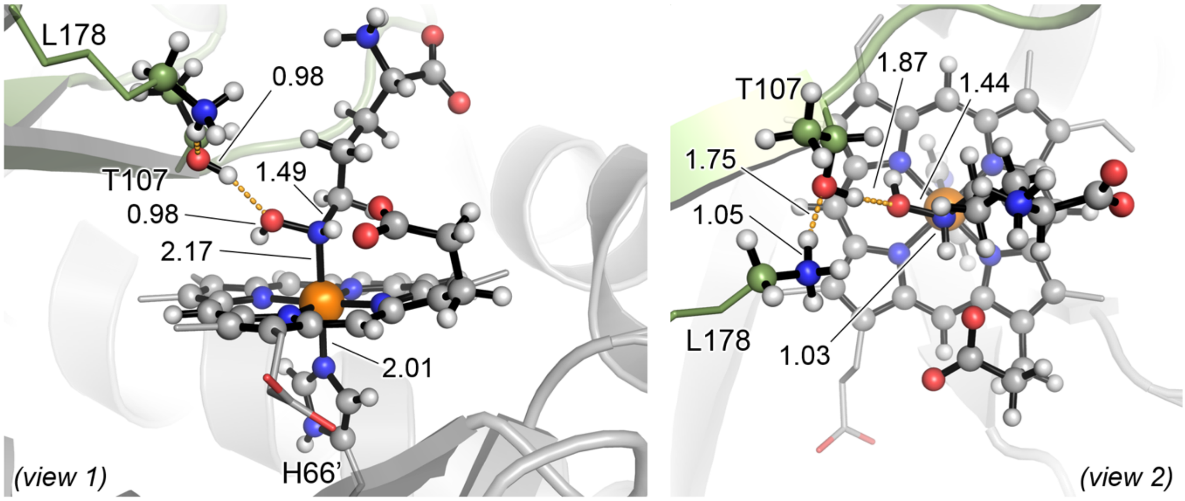

PipS-L-**1** (D)
ΔG = 10.0 (ΔE = 7.4)

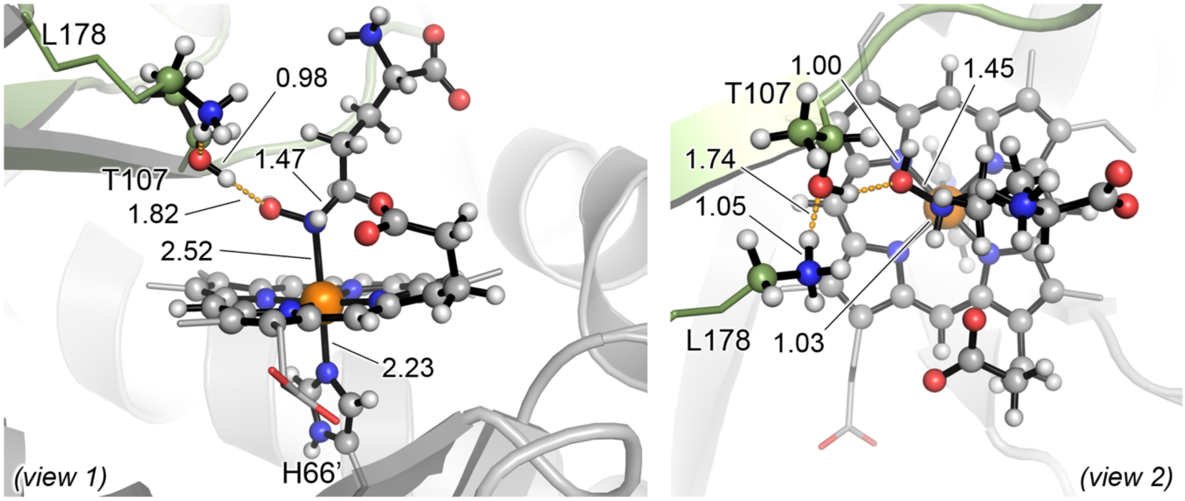

PipS-L-**1** (Qu)
ΔG = 0.0 (ΔE = 0.0)

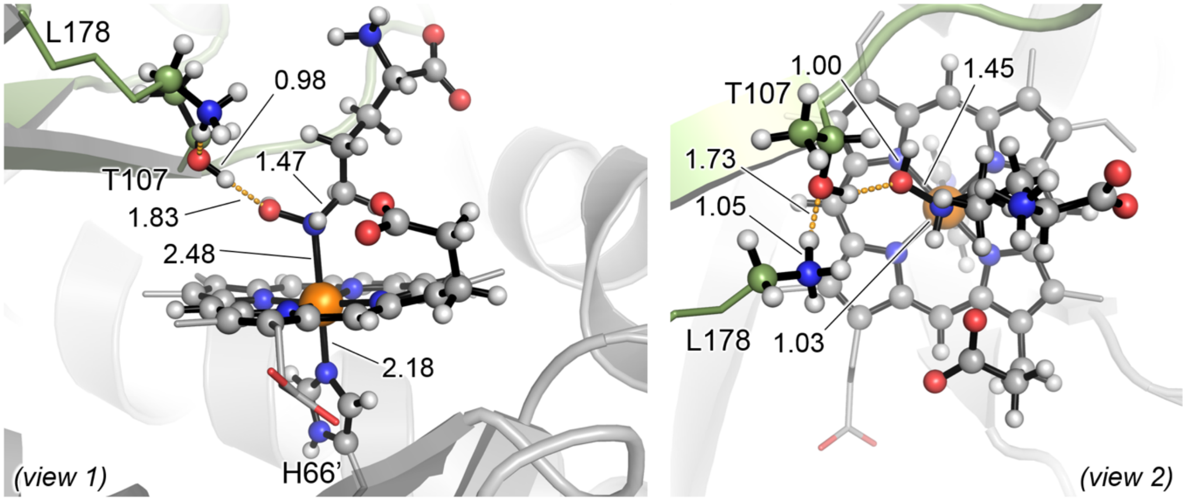

PipS-L-**1** (S)
ΔG = 3.9 (ΔE = 5.0)

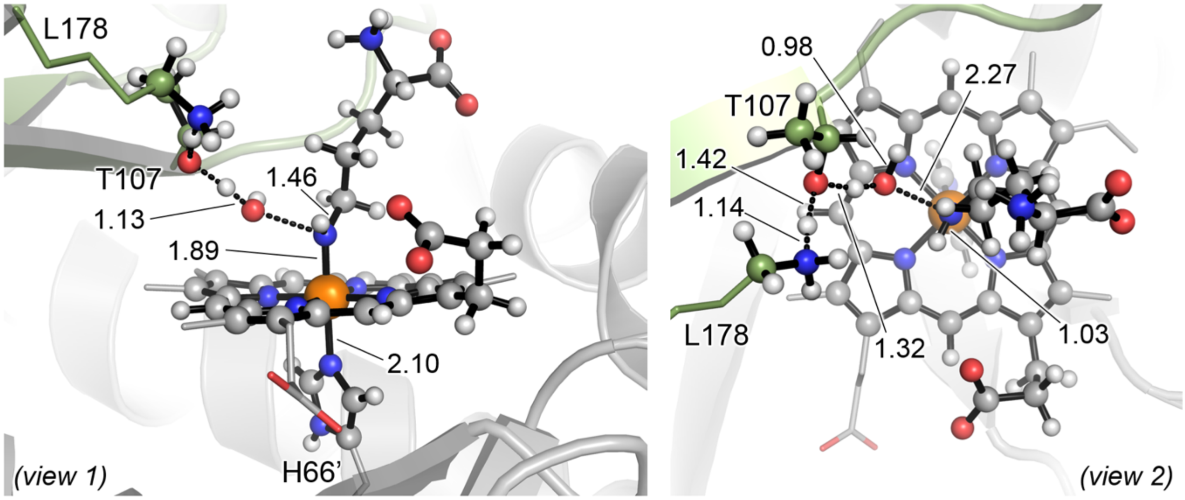

PipS-**TS1** (D)
$\Delta G^{\ddagger}$ = 38.9 ($\Delta E^{\ddagger}$ = 44.8)

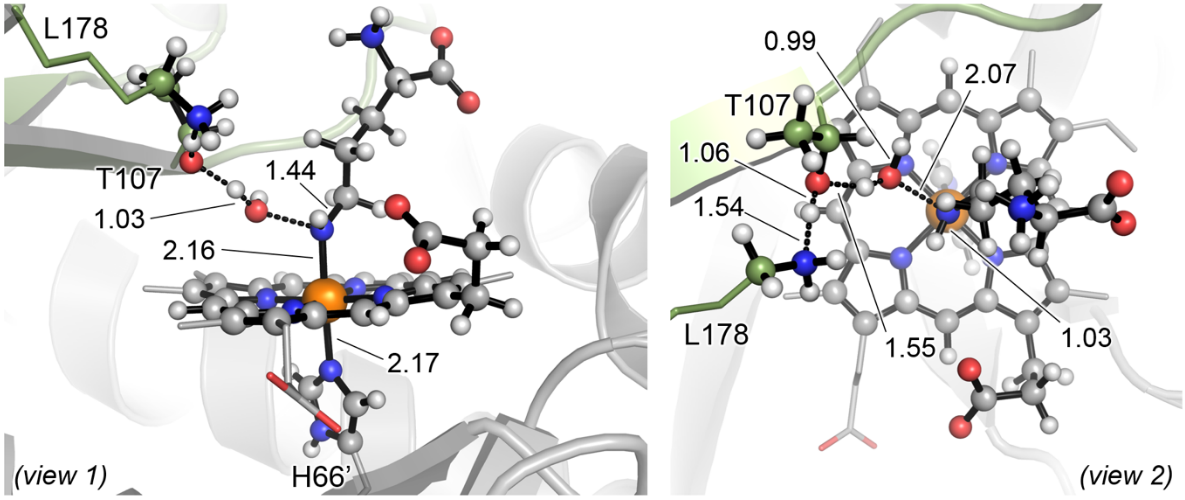

PipS-**TS1** (Qu)
$\Delta G^{\ddagger}$ = 52.7 ($\Delta E^{\ddagger}$ = 57.1)

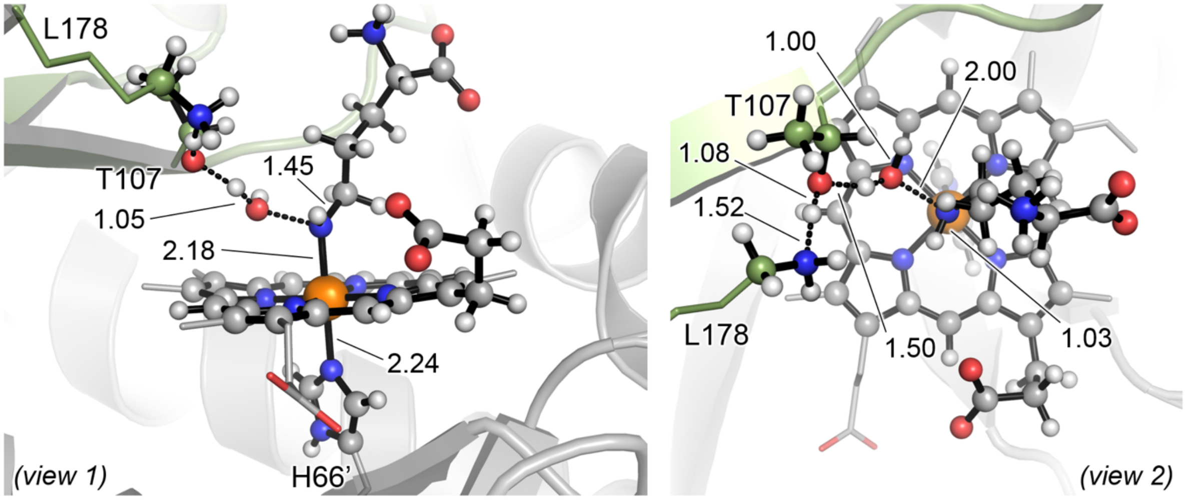

PipS-**TS1** (S)
ΔG‡ = 38.2 (ΔE‡ = 43.3)

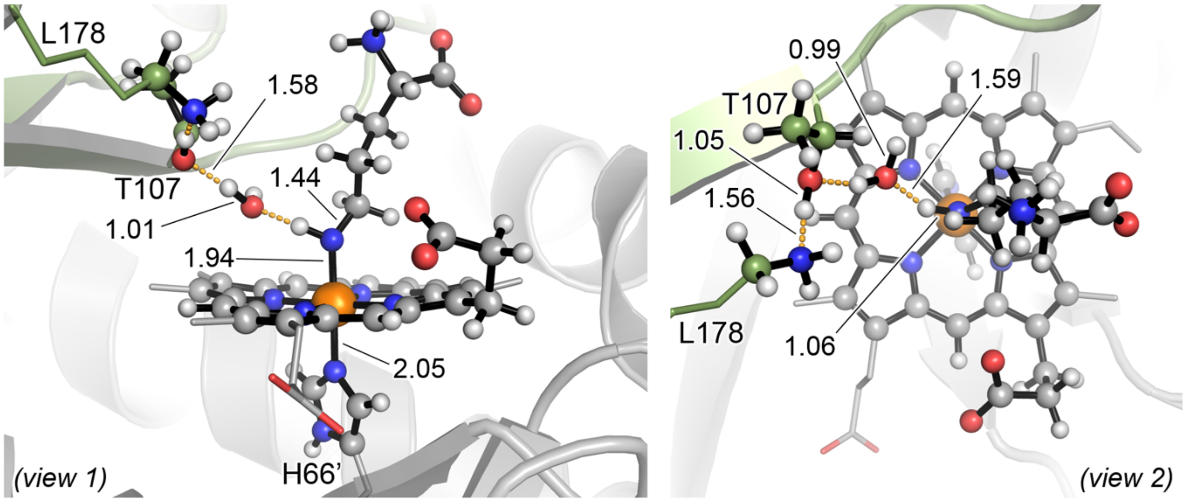

PipS-**Int1** (D)
$\Delta G_r$ = 27.4 ($\Delta E_r$ = 31.2)

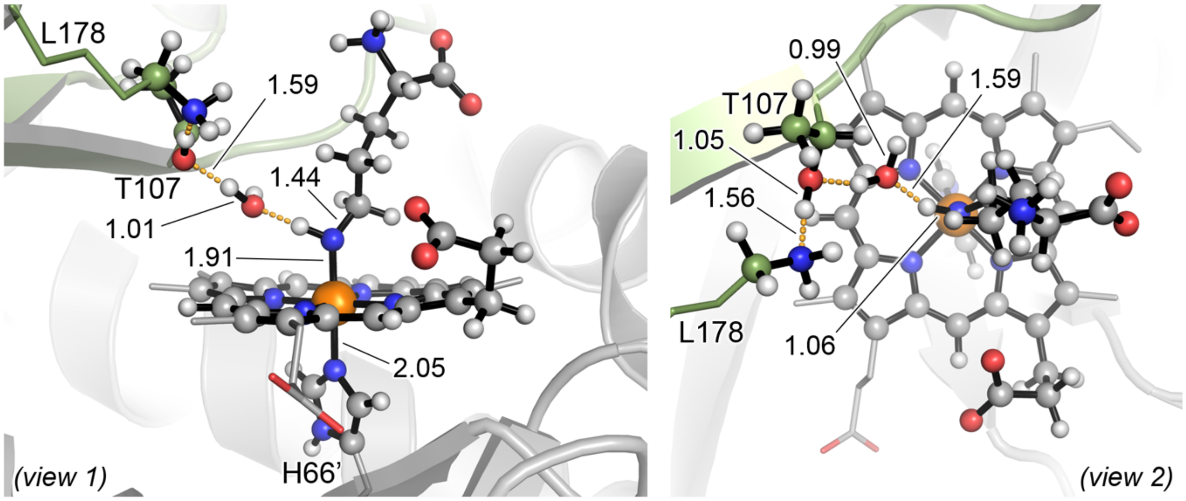

PipS-**Int1** (Qu)
$\Delta G_r$ = 37.0 ($\Delta E_r$ = 38.5)

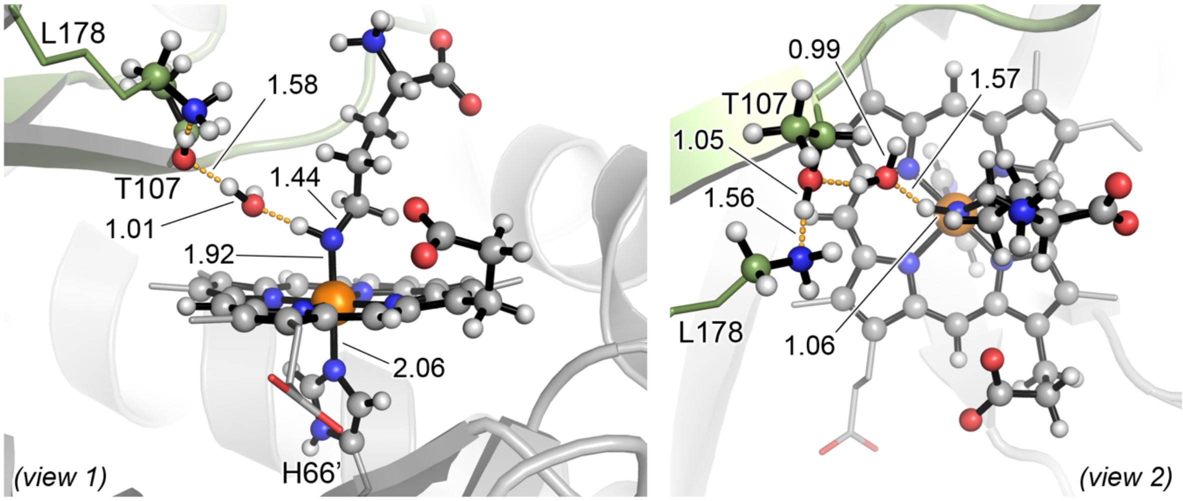

PipS-**Int1** (S)

$\Delta G_r = 46.2$ ($\Delta E_r = 50.0$)

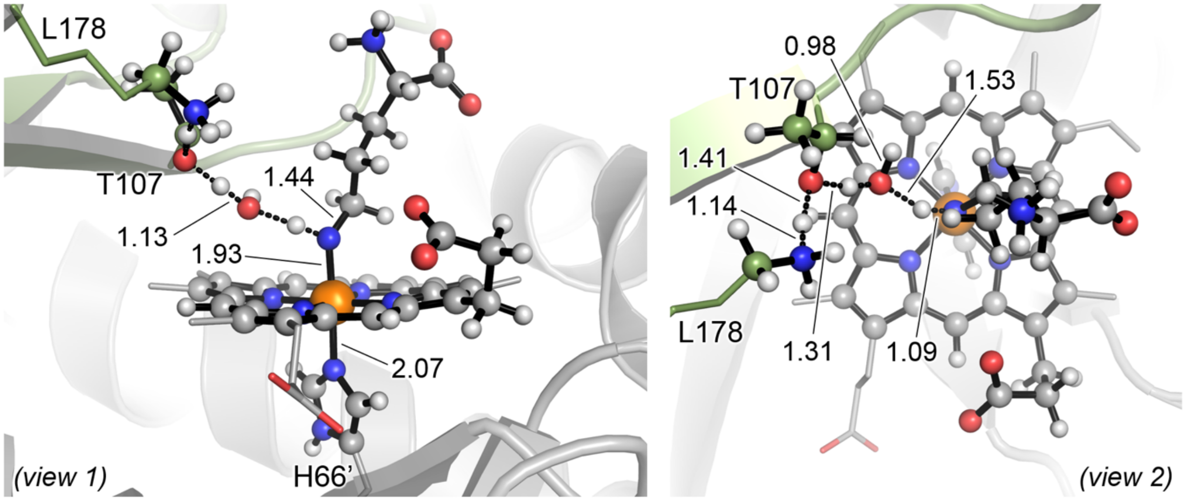

PipS-**TS2** (D)

$\Delta G^{\ddagger} = 0.0$ ($\Delta E^{\ddagger} = 3.3$)

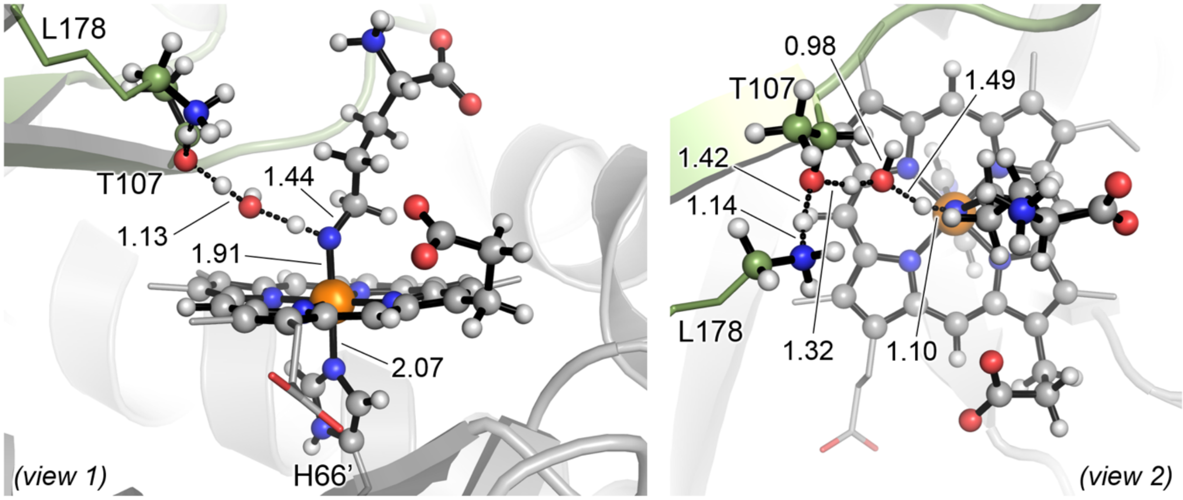

PipS-**TS2** (Qu)

$\Delta G^{\ddagger} = -1.1$ ($\Delta E^{\ddagger} = 2.2$)

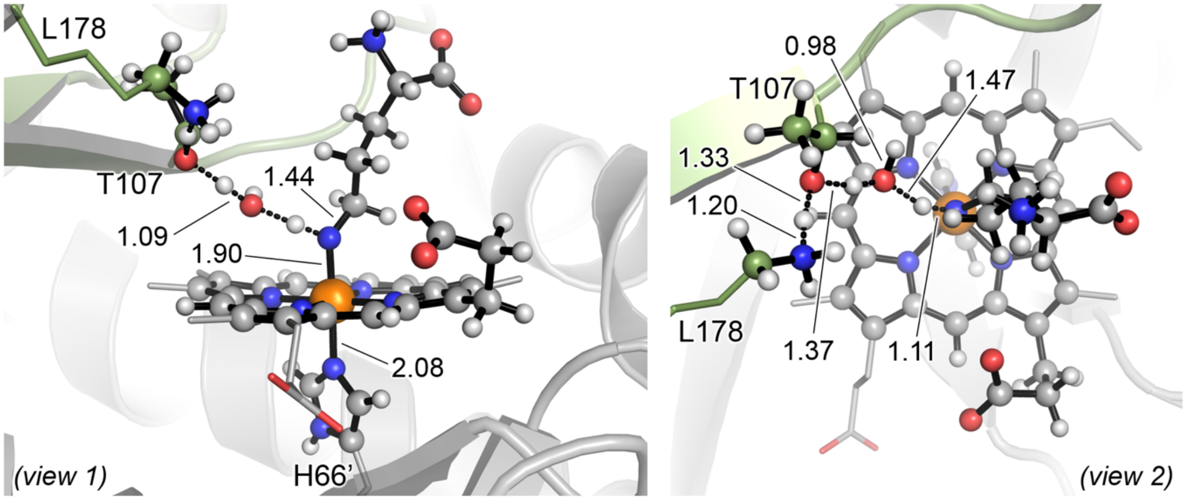

PipS-**TS2** (S)

$\Delta G^{\ddagger} = -1.8$ ($\Delta E^{\ddagger} = 1.7$)

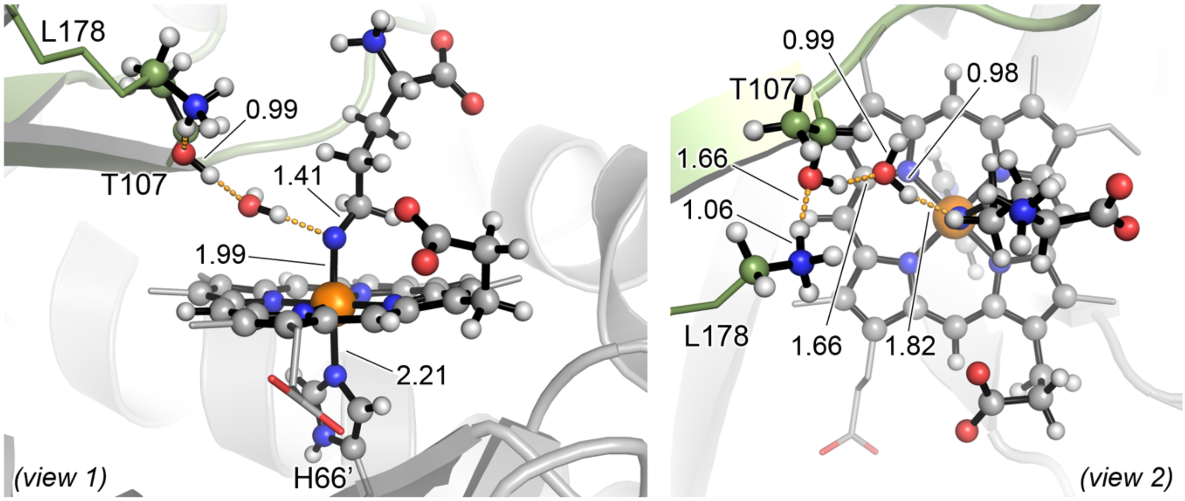

PipS-**Int2** (D)

$\Delta G_r = -12.4$ ($\Delta E_r = -13.1$)

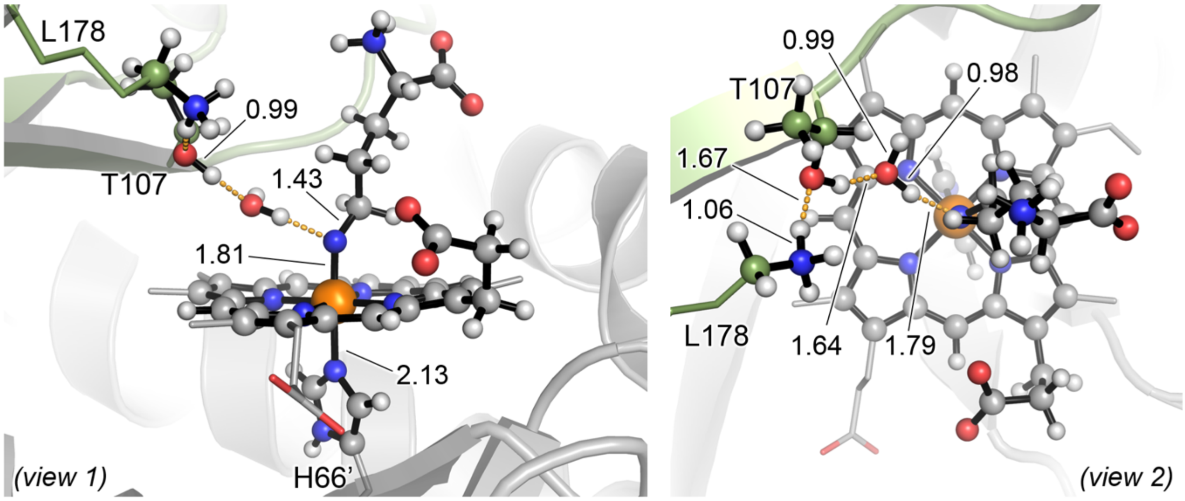

PipS-**Int2** (Qu)

$\Delta G_r = -12.0$ ($\Delta E_r = -13.2$)

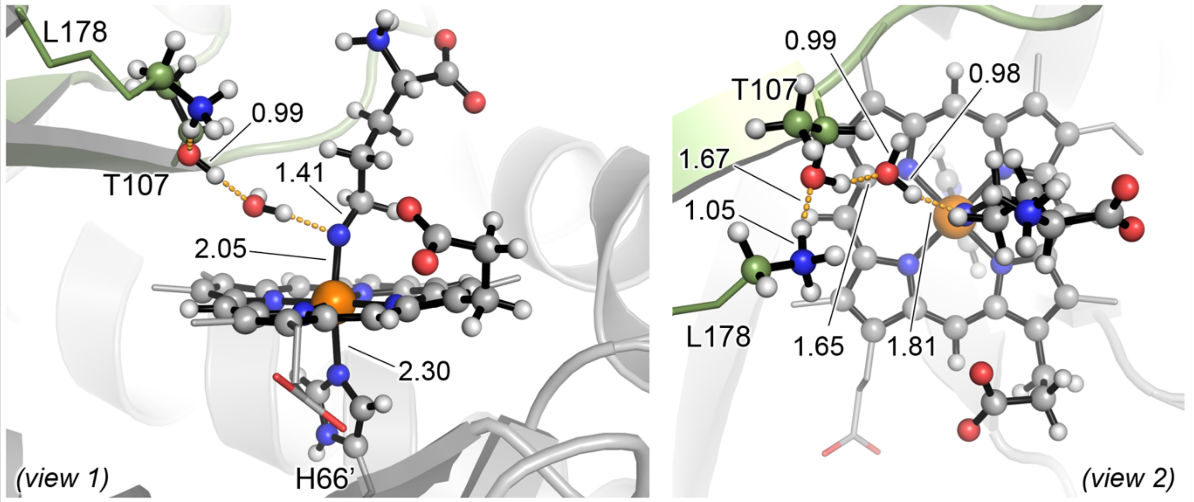

PipS-**Int2** (S)
$\Delta G_r$ = -13.1 ($\Delta E_r$ = -14.5)

The hydroxylamine activation pathway starting from a Fe(III) resting state is significantly higher in energy than the equivalent activation mechanism considering the Fe(II) oxidation state. The rate-limiting lowest in energy PipS-**TS1** optimized for Fe(III) oxidations tate is two times higher than the one calculated for Fe(II) (42.1 *versus* 21.2 kcal·mol$^{-1}$, see **Supplementary Figure 26**).

Once formed, the nitrenoid intermediate PipS-**Int1** quickly leads to intermediate PipS-**Int2** via PipS-**TS2**, restoring the initial protonation state of the catalytic T107-K178 dyad.

The resulting key nitrene intermediate PipS-**Int2** is *ca.* 25 kcal·mol$^{-1}$ higher in energy than the reactant complex PipS-L-**1**. This energy difference is *ca*. 18 kcal·mol$^{-1}$ higher than the energy difference between reactant complex PipS-L-**1** and PipS-**Int2** calculated for the Fe(II) oxidation state (**Supplementary Figure 26**).

The hydroxylamine activation following the proposed mechanism in the ferric oxidation state is much higher in energy than in the ferrous oxidation state. Consequently, it is proposed that PipS ferrous oxidation state is the one involved in catalysis. These results are in line with previous computational studies on aldoxime dehydratase Oxd (*J. Phys. Chem. B* **116**, 9396–9408 (2012)).

**Supplementary Figure 32. QM/MM exploration of PipS catalyzed N-N bond formation from nitrene intermediate in the Fe(III) oxidation state.** QM/MM calculations were carried out from the previously selected representative snapshot extracted from MD simulations with PipS-**Int2** bound complex (**Supplementary Figures 22** and **27**).

(**a**) The catalytic cycle of PipS in the Fe(III) oxidation state corresponding to the N-N bond formation (in dark blue).

(**b**) Schematic Potential Energy Surface (PES) for the N-N bond formation. Energy values were obtained at the (U)B3LYP/Def2TZVP:AmberFF14SB//(U)B3LYP/6-31G(d)+SDD(Fe): AmberFF14SB level, with the same MM parameters used in MD simulations (see SI, computational details). Relative Gibbs (ΔG) and electronic (ΔE) energies are reported for all electronic states. All energies are referred considering PipS-**Int2** (T) structure as zero. The PES profile for the equivalent pathway for the Fe(II) oxidation state is also provided.

(**c**) Computed relative stabilities in terms of electronic QM energy ($\Delta E_{QM}$), electronic QM/MM energy (ΔE), enthalpy (ΔH), and Gibbs energy (ΔG) for the different species. All energies are referred considering PipS-**Int2** (D) structure as zero.

(**d**) Optimized QM/MM structures for the different species involved in the N-N bond formation pathway. All atoms included in the QM-region are shown in ball and stick representation.

All energies and distances are given in $kcal{\cdot}mol^{-1}$ and angstrom (Å), respectively.

(a)

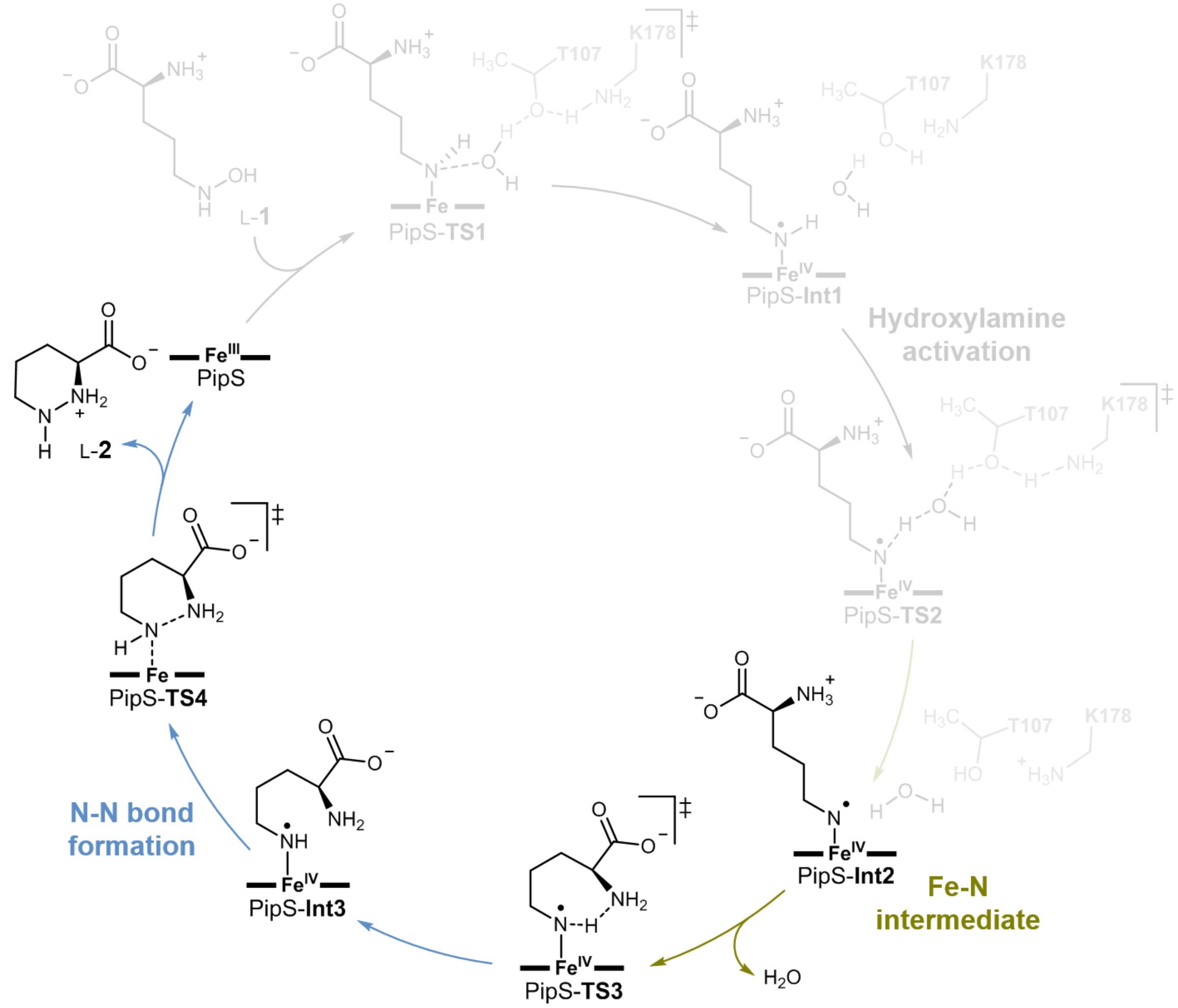
L-1
PipS-TS1
PipS-Int1
Hydroxylamine activation
PipS-TS2
T107
K178
PipS-Int2
Fe-N intermediate
$H_2O$
PipS-TS3
PipS-Int3
N-N bond formation
PipS-TS4
PipS
L-2

(b)

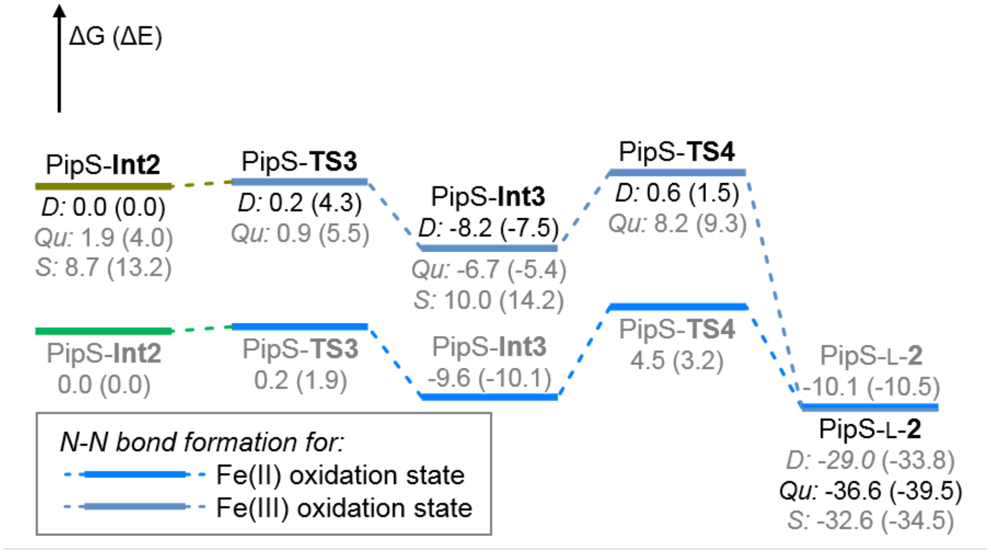
ΔG (ΔE)
PipS-Int2
D: 0.0 (0.0)
Qu: 1.9 (4.0)
S: 8.7 (13.2)
PipS-TS3
D: 0.2 (4.3)
Qu: 0.9 (5.5)
PipS-Int3
D: -8.2 (-7.5)
Qu: -6.7 (-5.4)
S: 10.0 (14.2)
PipS-TS4
D: 0.6 (1.5)
Qu: 8.2 (9.3)
PipS-Int2
0.0 (0.0)
PipS-TS3
0.2 (1.9)
PipS-Int3
-9.6 (-10.1)
PipS-TS4
4.5 (3.2)
PipS-L-2
-10.1 (-10.5)
PipS-L-2
D: -29.0 (-33.8)
Qu: -36.6 (-39.5)
S: -32.6 (-34.5)
N-N bond formation for:
Fe(II) oxidation state
Fe(III) oxidation state

**(c)**

| Structure | Electronic State | $\Delta E_{QM}$ | ΔE | ΔH | ΔG |
|---|---|---|---|---|---|
| PipS-**Int2** | doublet (D) | 0.0 | 0.0 | 0.0 | 0.0 |
| | quartet (Qu) | 3.5 | 4.0 | 3.5 | 1.9 |
| | sextet (S) | 11.9 | 13.2 | 10.9 | 8.7 |
| PipS-**TS3** | doublet (D) | 2.7 | 4.3 | -0.2 | 0.2 |
| | quartet (Qu) | 6.7 | 5.5 | 0.8 | 0.9 |
| PipS-**Int3** | doublet (D) | 44.5 | -7.5 | -9.4 | -8.2 |
| | quartet (Qu) | 46.8 | -5.4 | -7.3 | -6.7 |
| | sextet (S) | 64.1 | 14.2 | 10.6 | 10.0 |
| PipS-**TS4** | doublet (D) | 52.6 | 1.5 | -0.5 | 0.6 |
| | quartet (Qu) | 60.6 | 9.3 | 7.6 | 8.2 |
| PipS-L-**2** | doublet (D) | 40.5 | -33.8 | -30.7 | -29.0 |
| | quartet (Qu) | 35.4 | -39.5 | -37.0 | -36.6 |
| | sextet (S) | 40.7 | -34.5 | -32.9 | -32.6 |

**(d)**

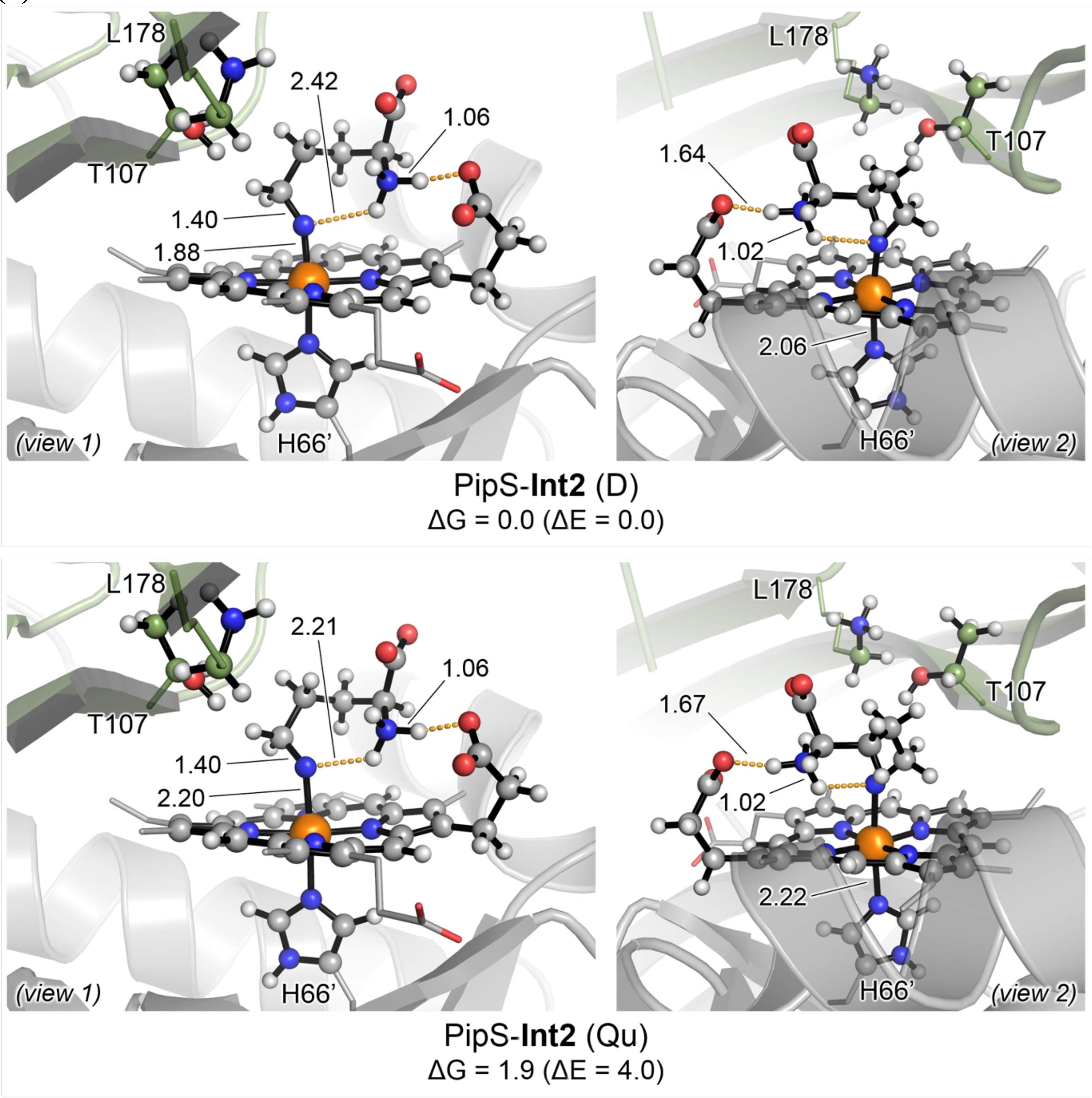

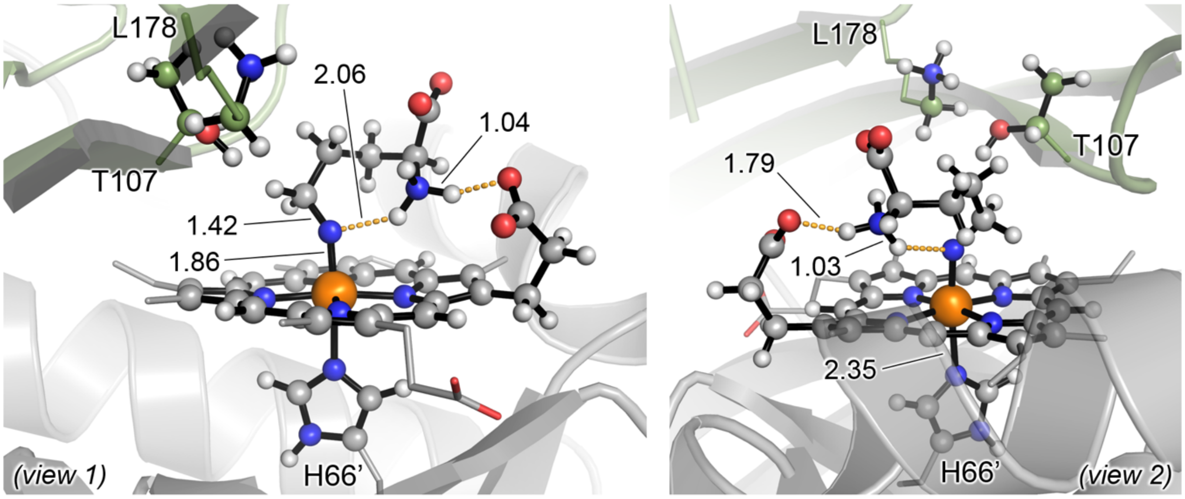

PipS-**Int2** (S)

$\Delta G_r$ = 8.7 ($\Delta E_r$ = 13.2)

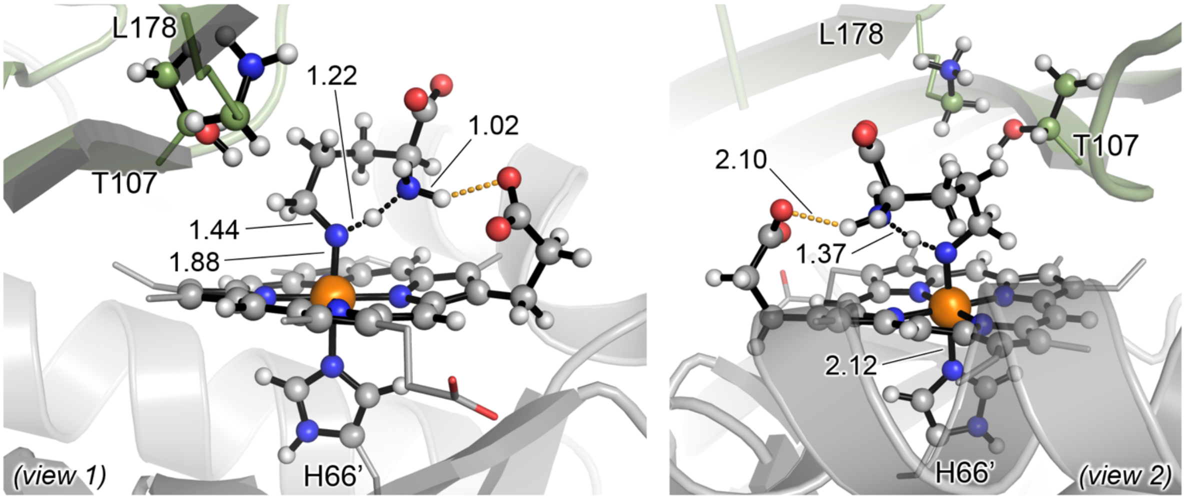

PipS-**TS3** (D)

$\Delta G^{\ddagger}$ = 0.2 ($\Delta E^{\ddagger}$ = 4.3)

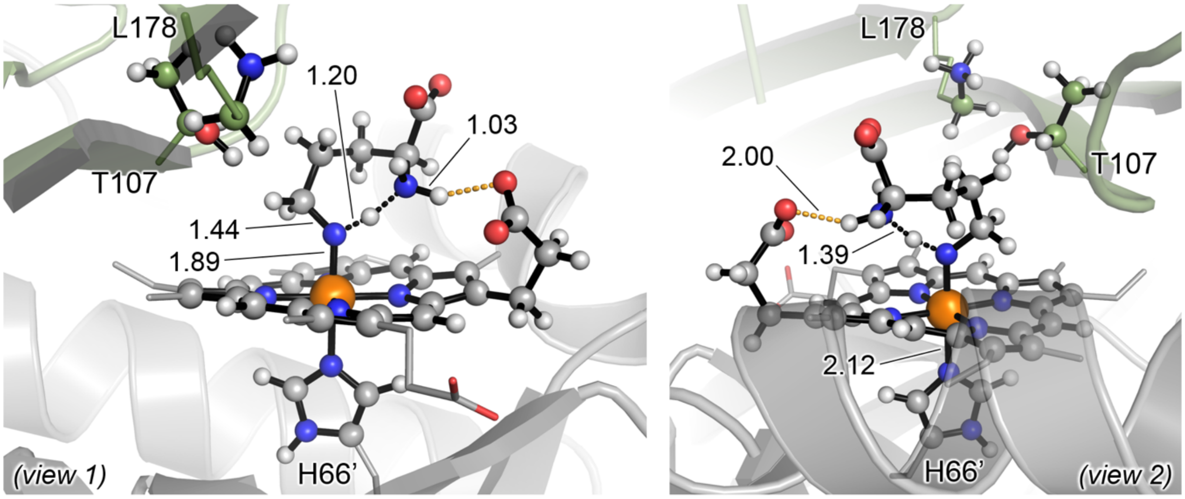

PipS-**TS3** (Qu)

$\Delta G^{\ddagger}$ = -1.1 ($\Delta E^{\ddagger}$ = 1.5)

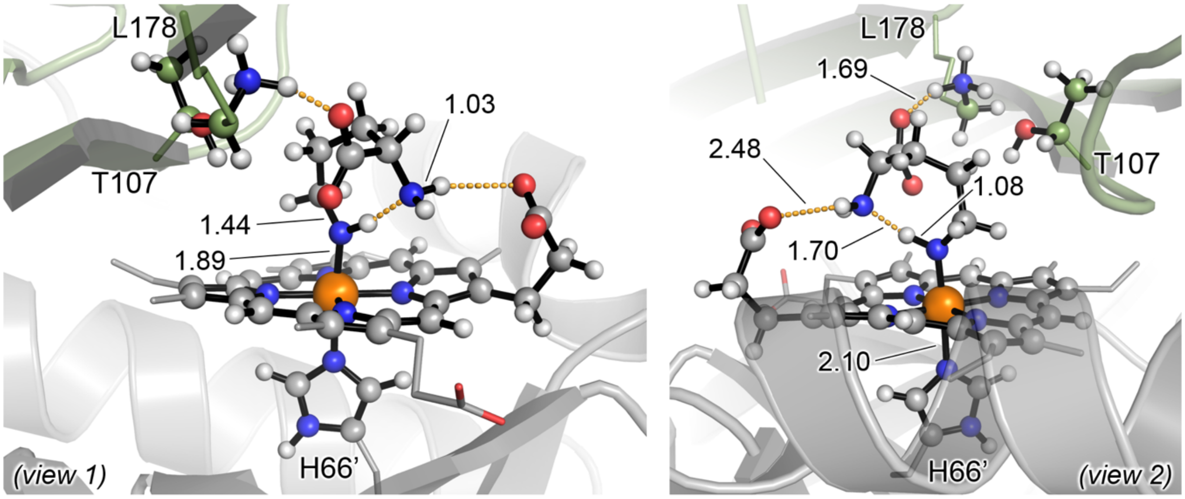

PipS-**Int3** (D)
$\Delta G_r$ = -8.2 ($\Delta E_r$ = -7.5)

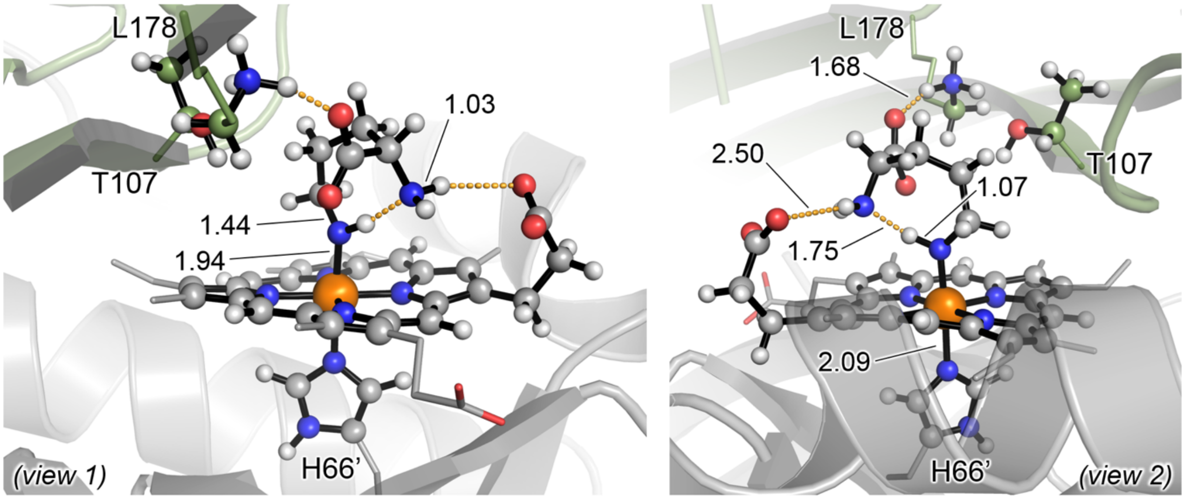

PipS-**Int3** (Qu)
$\Delta G_r$ = -8.6 ($\Delta E_r$ = -9.4)

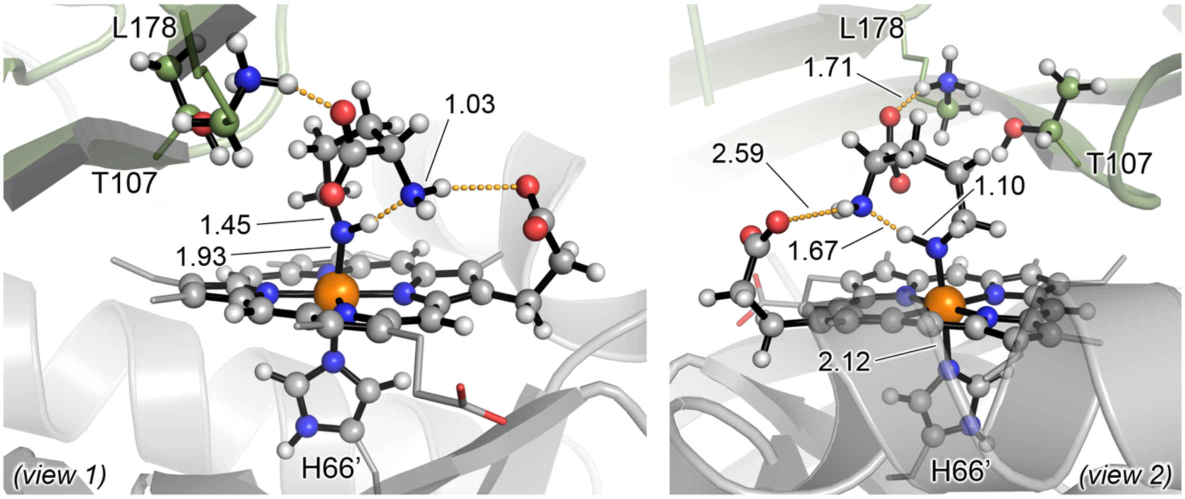

PipS-**Int3** (S)
$\Delta G_r$ = 1.4 ($\Delta E_r$ = 1.0)

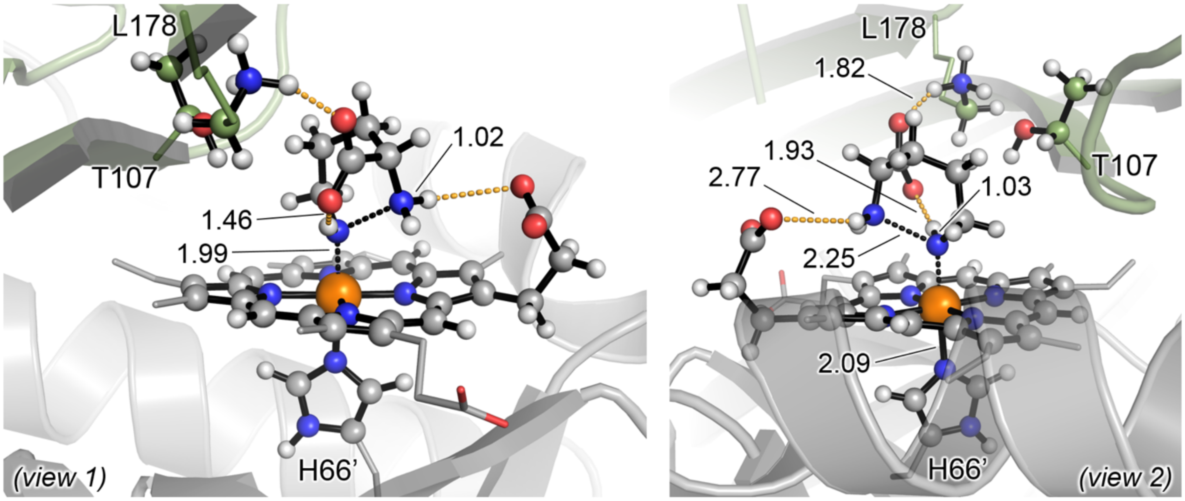

PipS-**TS4** (D)

$\Delta G^{\ddagger}$ = 8.9 ($\Delta E^{\ddagger}$ = 9.0)

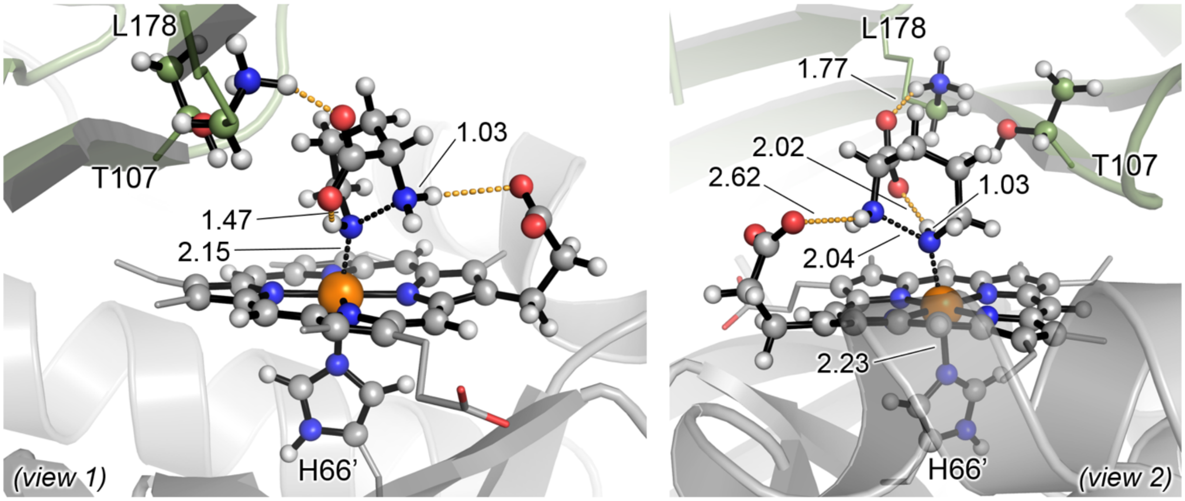

PipS-**TS4** (Qu)

$\Delta G^{\ddagger}$ = 14.9 ($\Delta E^{\ddagger}$ = 14.7)

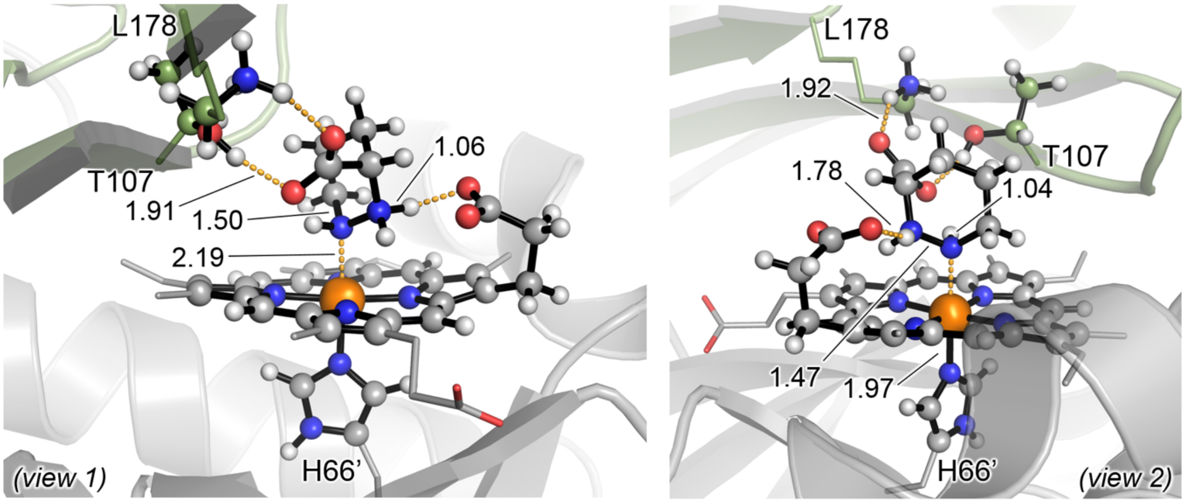

PipS-L-**2** (D)

$\Delta G_r$ = -20.7 ($\Delta E_r$ = -26.2)

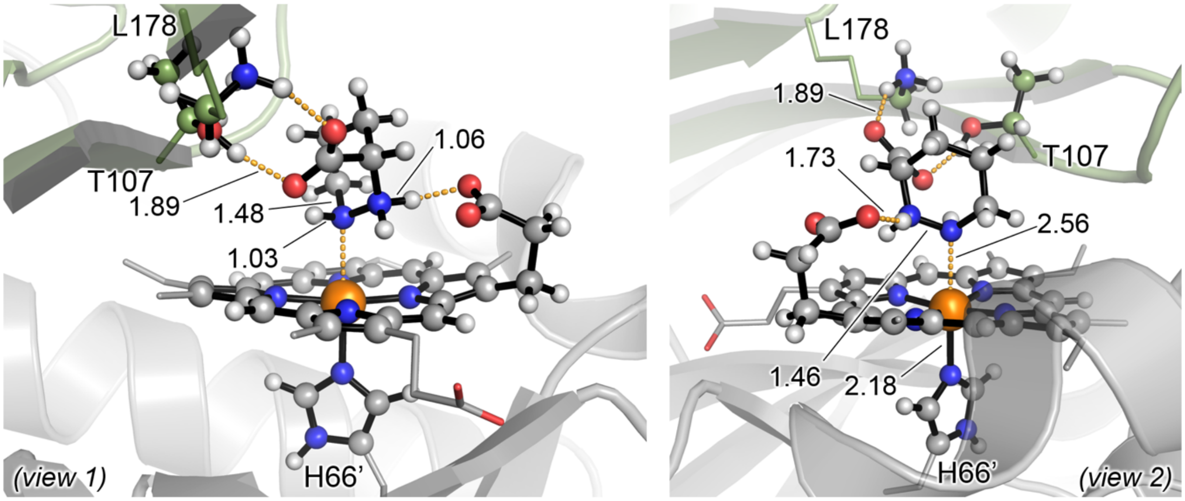

PipS-L-**2** (Qu)
$\Delta G_r$ = -29.9 ($\Delta E_r$ = -34.1)

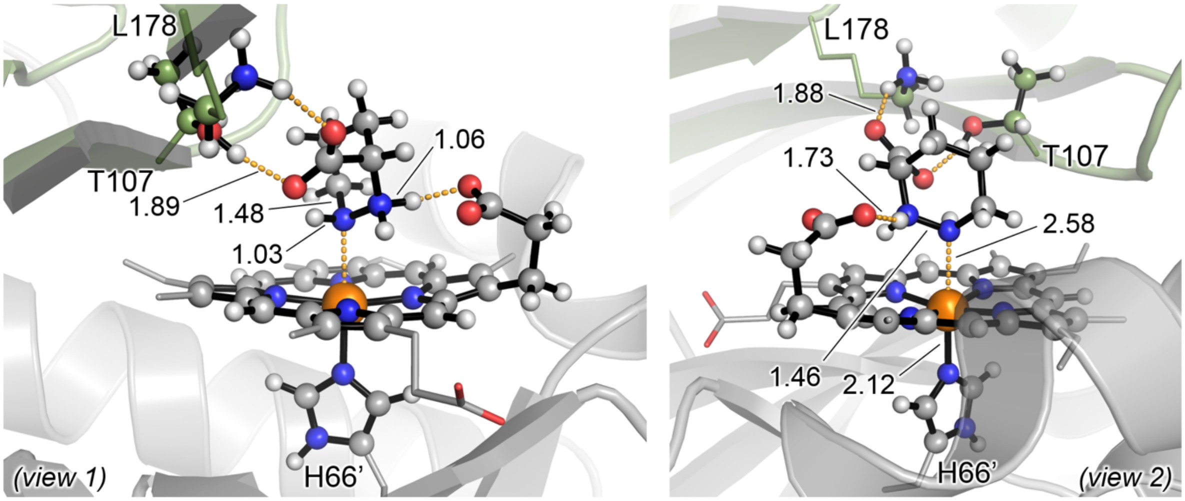

PipS-L-**2** (S)
$\Delta G_r$ = -42.6 ($\Delta E_r$ = -48.7)

The N-N bond formation pathway in the ferric oxidation state is energetically similar to the equivalent pathway for the ferrous oxidation state (see **Supplementary Figures 32b** and **27**).

The bottleneck for the ferric PipS catalytic cycle, as discussed earlier in **Supplementary Figure 31**, is the hydroxylamine activation step and the formation of the nitrene intermediate PipS-**Int2**. This hydroxylamine activation mechanism is shown to be very high in energy when considering the Fe(III) oxidation state, much higher than the equivalent mechanistic steps calculated for the Fe(II) oxidation state (**Supplementary Figure 31**)

**Supplementary Figure 33. Summary of QM/MM calculations on PipS catalytic cycle and its reaction mechanism in the Fe(III) oxidation state.**

(**a**) The catalytic cycle of PipS in the Fe(III) oxidation state. The hydroxylamine activation, the key Fe-N intermediate, and N-N bond formation are shown in black, olive, and dark blue, respectively.

(**b**) Schematic Potential Energy Surface (PES) for PipS catalytic cycle starting from the hydroxylamine activation and leading to N-N bond formation. Energy values were obtained from **Supplementary Figures 31** and **32**. The PES profile for the equivalent pathway but in the Fe(II) oxidation state is also provided.

**(a)**

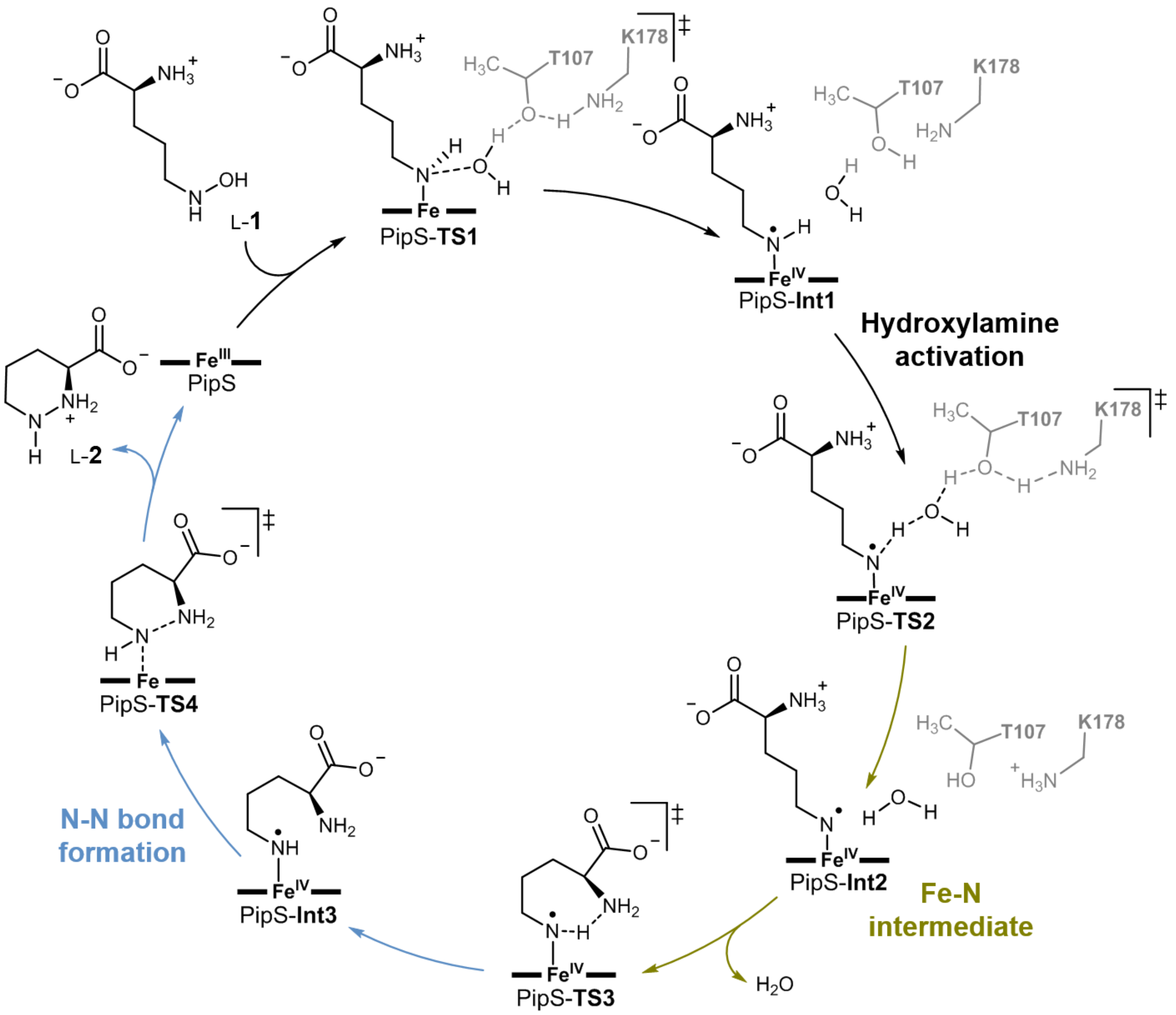

(b)

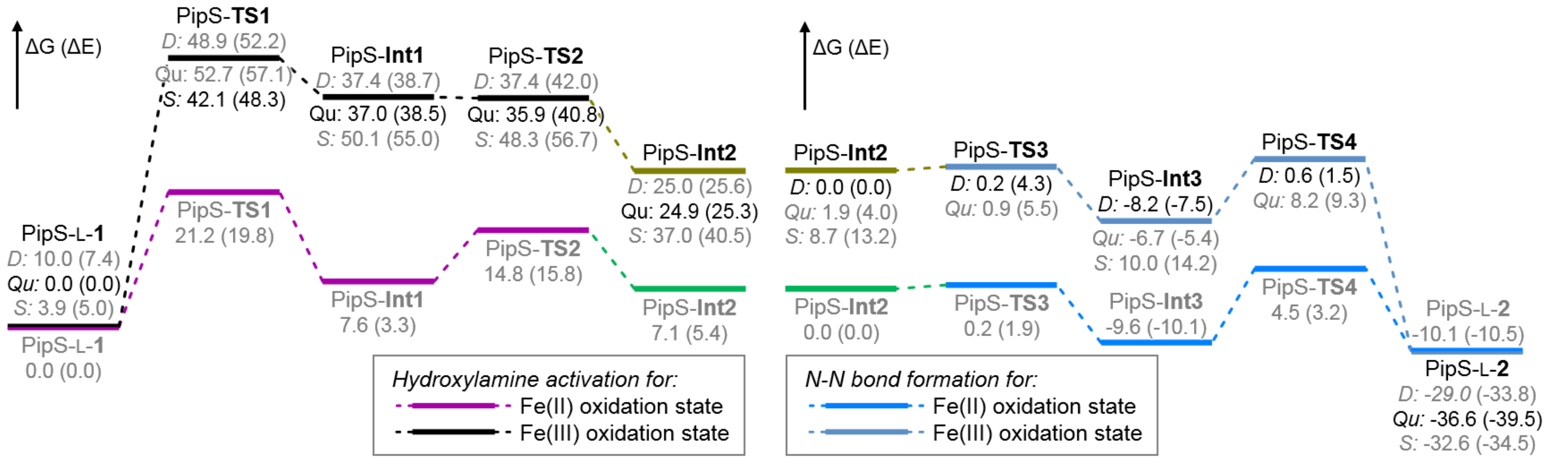

ΔG (ΔE)
PipS-TS1
D: 48.9 (52.2)
Qu: 52.7 (57.1)
S: 42.1 (48.3)
PipS-Int1
D: 37.4 (38.7)
Qu: 37.0 (38.5)
S: 50.1 (55.0)
PipS-TS2
D: 37.4 (42.0)
Qu: 35.9 (40.8)
S: 48.3 (56.7)
PipS-Int2
D: 25.0 (25.6)
Qu: 24.9 (25.3)
S: 37.0 (40.5)
PipS-L-1
D: 10.0 (7.4)
Qu: 0.0 (0.0)
S: 3.9 (5.0)
PipS-TS1
21.2 (19.8)
PipS-TS2
14.8 (15.8)
PipS-Int1
7.6 (3.3)
PipS-Int2
7.1 (5.4)
PipS-L-1
0.0 (0.0)
Hydroxylamine activation for:
Fe(II) oxidation state
Fe(III) oxidation state
ΔG (ΔE)
PipS-Int2
D: 0.0 (0.0)
Qu: 1.9 (4.0)
S: 8.7 (13.2)
PipS-TS3
D: 0.2 (4.3)
Qu: 0.9 (5.5)
PipS-Int3
D: -8.2 (-7.5)
Qu: -6.7 (-5.4)
S: 10.0 (14.2)
PipS-TS4
D: 0.6 (1.5)
Qu: 8.2 (9.3)
PipS-Int2
0.0 (0.0)
PipS-TS3
0.2 (1.9)
PipS-Int3
-9.6 (-10.1)
PipS-TS4
4.5 (3.2)
PipS-L-2
-10.1 (-10.5)
PipS-L-2
D: -29.0 (-33.8)
Qu: -36.6 (-39.5)
S: -32.6 (-34.5)
N-N bond formation for:
Fe(II) oxidation state
Fe(III) oxidation state

**Supplementary Table 4.** Energies and thermochemistry parameters (at T = 298.15 K and P = 1 atm) of all computationally characterized stationary points reported in **Supplementary Figures 26** and **29**: QM Electronic energies ($E_{QM}$), MM energies ($E_{MM}$), enthalpy corrections (H correction), free energy corrections (G correction), QM electronic energies from high level single point calculations ($E_{QM}$ (SP)), and single imaginary frequencies for transitions state. All energies and frequencies are given in a.u. and $cm^{-1}$, respectively.

| Structure | Electronic State | $E_{QM}$ | $E_{MM}$ | H correction | G correction | $E_{QM}$ (SP) | Imag. Freq. |
|---|---|---|---|---|---|---|---|
| PipS-L-**1** | close-shell singlet (CSS) | -2389.841138 | -34.253519 | 22.453591 | 20.087608 | -3530.625904 | - |
| | triplet (T) | -2389.832330 | -34.256885 | 22.452743 | 20.081428 | -3530.618722 | - |
| | quintet (Q) | -2389.837503 | -34.258039 | 22.451638 | 20.079276 | -3530.620695 | - |
| PipS-**TS1** | close-shell singlet (CSS) | -2389.792601 | -34.250074 | 22.445856 | 20.084171 | -3530.580792 | 357.2i |
| | open-shell singlet (OSS) | -2389.798271 | -34.256018 | 22.444999 | 20.081470 | -3530.591103 | 522.7i |
| | triplet (T) | -2389.786824 | -34.254727 | 22.448076 | 20.082544 | -3530.577440 | 397.9i |
| | quintet (Q) | -2389.776509 | -34.255959 | 22.445283 | 20.077793 | -3530.575242 | 892.0i |
| PipS-**Int1** | close-shell singlet (CSS) | -2389.819746 | -34.241922 | 22.451724 | 20.088016 | -3530.622691 | - |
| | open-shell singlet (OSS) | -2389.842158 | -34.238891 | 22.450871 | 20.086097 | -3530.634616 | - |
| | triplet (T) | -2389.833763 | -34.244797 | 22.450931 | 20.085885 | -3530.626204 | - |
| | quintet (Q) | -2389.824726 | -34.242430 | 22.450817 | 20.083281 | -3530.616475 | - |
| PipS-**TS2** | close-shell singlet (CSS) | -2389.803197 | -34.242286 | 22.441967 | 20.079300 | -3530.606136 | 1229.5i |
| | open-shell singlet (OSS) | -2389.817408 | -34.244848 | 22.442121 | 20.079256 | -3530.609760 | 1156.7i |
| | triplet (T) | -2389.815657 | -34.244871 | 22.441774 | 20.077783 | -3530.608742 | 1219.8i |
| | quintet (Q) | -2389.804042 | -34.244420 | 22.440032 | 20.073019 | -3530.577594 | 1171.2i |
| PipS-**Int2** | close-shell singlet (CSS) | -2389.824731 | -34.238440 | 22.451540 | 20.087668 | -3530.624531 | - |
| | open-shell singlet (OSS) | -2389.835927 | -34.240739 | 22.450111 | 20.082900 | -3530.627041 | - |
| | triplet (T) | -2389.837952 | -34.240263 | 22.450411 | 20.082049 | -3530.629896 | - |
| | quintet (Q) | -2389.824855 | -34.240473 | 22.448896 | 20.077724 | -3530.613053 | - |
| PipS-**TS6** | open-shell singlet (OSS) | -2389.838233 | -34.242000 | 22.446219 | 20.085300 | -3530.632172 | 1063.3i |
| | triplet (T) | -2389.824551 | -34.241305 | 22.445587 | 20.083181 | -3530.618873 | 1021.1i |
| | quintet (Q) | -2389.812383 | -34.242430 | 22.444615 | 20.079575 | -3530.613098 | 1997.4i |

**Supplementary Table 5.** Energies and thermochemistry parameters (at T = 298.15 K and P = 1 atm) of all computationally characterized stationary points reported in **Supplementary Figure 27**: QM Electronic energies ($E_{QM}$), MM energies ($E_{MM}$), enthalpy corrections (H correction), free energy corrections (G correction), QM electronic energies from high level single point calculations ($E_{QM}$ (SP)), and single imaginary frequencies for transitions state. All energies and frequencies are given in a.u. and $cm^{-1}$, respectively.

| Structure | Electronic State | $E_{QM}$ | $E_{MM}$ | H correction | G correction | $E_{QM}$ (SP) | Imag. Freq. |
|---|---|---|---|---|---|---|---|
| PipS-**Int2** | close-shell singlet (CSS) | -2313.420631 | -32.684454 | 20.373195 | 18.230670 | -3454.165061 | - |
| | open-shell singlet (OSS) | -2313.433843 | -32.684529 | 20.372621 | 18.229832 | -3454.176252 | - |
| | triplet (T) | -2313.433859 | -32.684083 | 20.373040 | 18.229111 | -3454.177287 | - |
| | quintet (Q) | -2313.408572 | -32.686605 | 20.371037 | 18.223501 | -3454.142527 | - |
| PipS-**TS3** | close-shell singlet (CSS) | -2313.412911 | -32.682805 | 20.368121 | 18.225824 | -3454.160562 | 989.3i |
| | open-shell singlet (OSS) | -2313.434144 | -32.679908 | 20.368062 | 18.226375 | -3454.178406 | 1028.0i |
| | triplet (T) | -2313.431159 | -32.679484 | 20.367923 | 18.224744 | -3454.176826 | 1016.6i |
| | quintet (Q) | -2313.418669 | -32.677916 | 20.365731 | 18.220160 | -3454.157511 | 1018.6i |
| PipS-**Int3** | close-shell singlet (CSS) | -2313.359784 | -32.763841 | 20.371300 | 18.230353 | -3454.097440 | - |
| | open-shell singlet (OSS) | -2313.378179 | -32.764572 | 20.371029 | 18.229952 | -3454.112904 | - |
| | triplet (T) | -2313.374503 | -32.764524 | 20.370873 | 18.228588 | -3454.110331 | - |
| | quintet (Q) | -2313.369185 | -32.761179 | 20.369980 | 18.225170 | -3454.105797 | - |
| PipS-**TS4** | close-shell singlet (CSS) | | | barrierless | | | |
| | open-shell singlet (OSS) | -2313.335543 | -32.796487 | 20.372366 | 18.231232 | -3454.059802 | 469.8i |
| | triplet (T) | -2313.323074 | -32.799193 | 20.370593 | 18.225044 | -3454.041938 | 386.5i |
| | quintet (Q) | -2313.312450 | -32.799096 | 20.369138 | 18.223038 | -3454.043580 | 716.6i |
| PipS-L-**2** | close-shell singlet (CSS) | -2313.367709 | -32.782776 | 20.376739 | 18.237405 | -3454.087368 | - |
| | open-shell singlet (OSS) | -2313.367428 | -32.783130 | 20.376528 | 18.236540 | -3454.087057 | - |
| | triplet (T) | -2313.365923 | -32.788295 | 20.375809 | 18.232236 | -3454.087469 | - |
| | quintet (Q) | -2313.373991 | -32.788390 | 20.374705 | 18.229789 | -3454.089734 | - |
| PipS-**TS5a** | close-shell singlet (CSS) | -2313.394940 | -32.684888 | 20.368191 | 18.226709 | -3454.142141 | 978.3i |
| | triplet (T) | -2313.378888 | -32.689074 | 20.367510 | 18.222349 | -3454.125968 | 1092.5i |
| PipS-**TS5b** | close-shell singlet (CSS) | -2313.395849 | -32.668256 | 20.367535 | 18.225290 | -3454.142269 | 925.7i |
| PipS-L-**3** | close-shell singlet (CSS) | -2313.451707 | -32.731370 | 20.371802 | 18.230394 | -3454.190854 | - |
| | triplet (T) | -2313.436057 | -32.732600 | 20.370775 | 18.227469 | -3454.174712 | - |
| | quintet (Q) | -2313.446431 | -32.732239 | 20.370184 | 18.226220 | -3454.180499 | - |

**Supplementary Table 6.** Energies and thermochemistry parameters (at T = 298.15 K and P = 1 atm) of all computationally characterized stationary points reported in **Supplementary Figure 31**: QM Electronic energies ($E_{QM}$), MM energies ($E_{MM}$), enthalpy corrections (H correction), free energy corrections (G correction), QM electronic energies from high level single point calculations ($E_{QM}$ (SP)), and single imaginary frequencies for transitions state. All energies and frequencies are given in a.u. and $cm^{-1}$, respectively.

| Structure | Electronic State | $E_{QM}$ | $E_{MM}$ | H correction | G correction | $E_{QM}$ (SP) | Imag. Freq. |
|---|---|---|---|---|---|---|---|
| | doublet (D) | -2389.886815 | -34.261521 | 22.456341 | 20.089154 | -3530.661065 | - |
| PipS-L-**1** | quartet (Qu) | -2389.901999 | -34.255755 | 22.455650 | 20.085097 | -3530.678678 | - |
| | sextet (S) | -2389.899306 | -34.254962 | 22.454242 | 20.083364 | -3530.671498 | - |
| | doublet (D) | -2389.845304 | -34.242034 | 22.444562 | 20.079688 | -3530.609137 | 469.9i |
| PipS-**TS1** | quartet (Qu) | -2389.817908 | -34.243933 | 22.445867 | 20.078014 | -3530.599457 | 422.8i |
| | sextet (S) | -2389.835717 | -34.245436 | 22.444346 | 20.075246 | -3530.611983 | 414.8i |
| | doublet (D) | -2389.849094 | -34.239621 | 22.449631 | 20.083055 | -3530.633173 | - |
| PipS-**Int1** | quartet (Qu) | -2389.850435 | -34.239970 | 22.449564 | 20.082650 | -3530.633097 | - |
| | sextet (S) | -2389.825224 | -34.241775 | 22.446955 | 20.077434 | -3530.605079 | - |
| | doublet (D) | -2389.843516 | -34.240785 | 22.443809 | 20.077867 | -3530.626745 | 393.7i |
| PipS-**TS2** | quartet (Qu) | -2389.845110 | -34.240786 | 22.443411 | 20.077366 | -3530.628696 | 396.1i |
| | sextet (S) | -2389.821402 | -34.242154 | 22.440533 | 20.071771 | -3530.601947 | 491.7i |
| | doublet (D) | -2389.888745 | -34.219450 | 22.452585 | 20.084191 | -3530.674234 | - |
| PipS-**Int2** | quartet (Qu) | -2389.886959 | -34.224890 | 22.451203 | 20.084560 | -3530.669248 | - |
| | sextet (S) | -2389.870326 | -34.225279 | 22.449347 | 20.079570 | -3530.644644 | - |

**Supplementary Table 7.** Energies and thermochemistry parameters (at T = 298.15 K and P = 1 atm) of all computationally characterized stationary points reported in **Supplementary Figure 32**: QM Electronic energies ($E_{QM}$), MM energies ($E_{MM}$), enthalpy corrections (H correction), free energy corrections (G correction), QM electronic energies from high level single point calculations ($E_{QM}$ (SP)), and single imaginary frequencies for transitions state. All energies and frequencies are given in a.u. and $cm^{-1}$, respectively.

| Structure | Electronic State | $E_{QM}$ | $E_{MM}$ | H correction | G correction | $E_{QM}$ (SP) | Imag. Freq. |
|---|---|---|---|---|---|---|---|
| PipS-**Int2** | doublet (D) | -2313.532760 | -32.668431 | 20.373337 | 18.227835 | -3454.259807 | - |
| | quartet (Qu) | -2313.522425 | -32.667629 | 20.372541 | 18.224500 | -3454.254201 | - |
| | sextet (S) | -2313.514580 | -32.666367 | 20.369603 | 18.220545 | -3454.240780 | - |
| PipS-**TS3** | doublet (D) | -2313.520897 | -32.665887 | 20.366138 | 18.221307 | -3454.255483 | 966.5i |
| | quartet (Qu) | -2313.514336 | -32.670234 | 20.365823 | 18.220403 | -3454.249196 | 935.9i |
| PipS-**Int3** | doublet (D) | -2313.461245 | -32.751428 | 20.370374 | 18.226709 | -3454.188826 | - |
| | quartet (Qu) | -2313.443475 | -32.751667 | 20.370349 | 18.225756 | -3454.185184 | - |
| | sextet (S) | -2313.432491 | -32.747968 | 20.367625 | 18.221183 | -3454.157618 | - |
| PipS-**TS4** | doublet (D) | -2313.451264 | -32.749869 | 20.370108 | 18.226503 | -3454.176005 | 448.9i |
| | quartet (Qu) | -2313.436759 | -32.750298 | 20.370721 | 18.226082 | -3454.163187 | 509.0i |
| PipS-L-**2** | doublet (D) | -2313.483035 | -32.786699 | 20.378145 | 18.235471 | -3454.195331 | - |
| | quartet (Qu) | -2313.490478 | -32.787776 | 20.377213 | 18.232427 | -3454.203369 | - |
| | sextet (S) | -2313.486461 | -32.788263 | 20.375861 | 18.230909 | -3454.194949 | - |
